# Correlations reveal the hierarchical organization of networks with latent binary variables

**DOI:** 10.1101/2023.07.27.550891

**Authors:** Stefan Häusler

## Abstract

Deciphering the functional organization of large biological networks is a major challenge for current mathematical methods. A common approach is to decompose networks into largely independent functional modules, but inferring these modules and their organization from network activity is difficult, given the uncertainties and incompleteness of measurements. Typically, some parts of the overall functional organization, such as intermediate processing steps, are latent. We show that the hidden structure can be determined from the statistical moments of observable network components alone, as long as the functional relevance of the network components lies in their mean values and the mean of each latent variable maps onto a scaled expectation of a binary variable. Whether the function of biological networks permits a hierarchical modularization can be falsified by a correlation-based statistical test that we derive. We apply the test to three biological networks at different spatial scales, i.e., gene regulatory networks, dendrites of pyramidal neurons, and networks of spiking neurons.

## 1 Introduction

Modern recording techniques in neuroscience and cell biology are generating datasets of rapidly increasing dimensionality, posing a major challenge to current analytical methods for deciphering the function of the underlying biological systems [1, 2]. A promising approach in graph theory [3–5] is to decompose and organize complex networks [6] into largely autonomous functional modules [7], as found at all levels of biological organization [3, 8]. Various heuristic algorithms have been proposed to detect functional modularity, many of them based on hierarchical clustering [9], but a more rigorous analysis requires exact probabilistic inference [10]. Following this approach, a functional module can be conveniently formalized as a subnetwork that communicates or interacts with the rest of the network only through a particular variable, which we call interface variable. This interface variable may represent, for example, the firing rate of a population of sensory neurons that encodes all information about stimuli relevant to downstream areas. If the value of this interface variable is known, the internal and external components of a module are conditionally independent (Fig. 1A).

**Figure 1:**
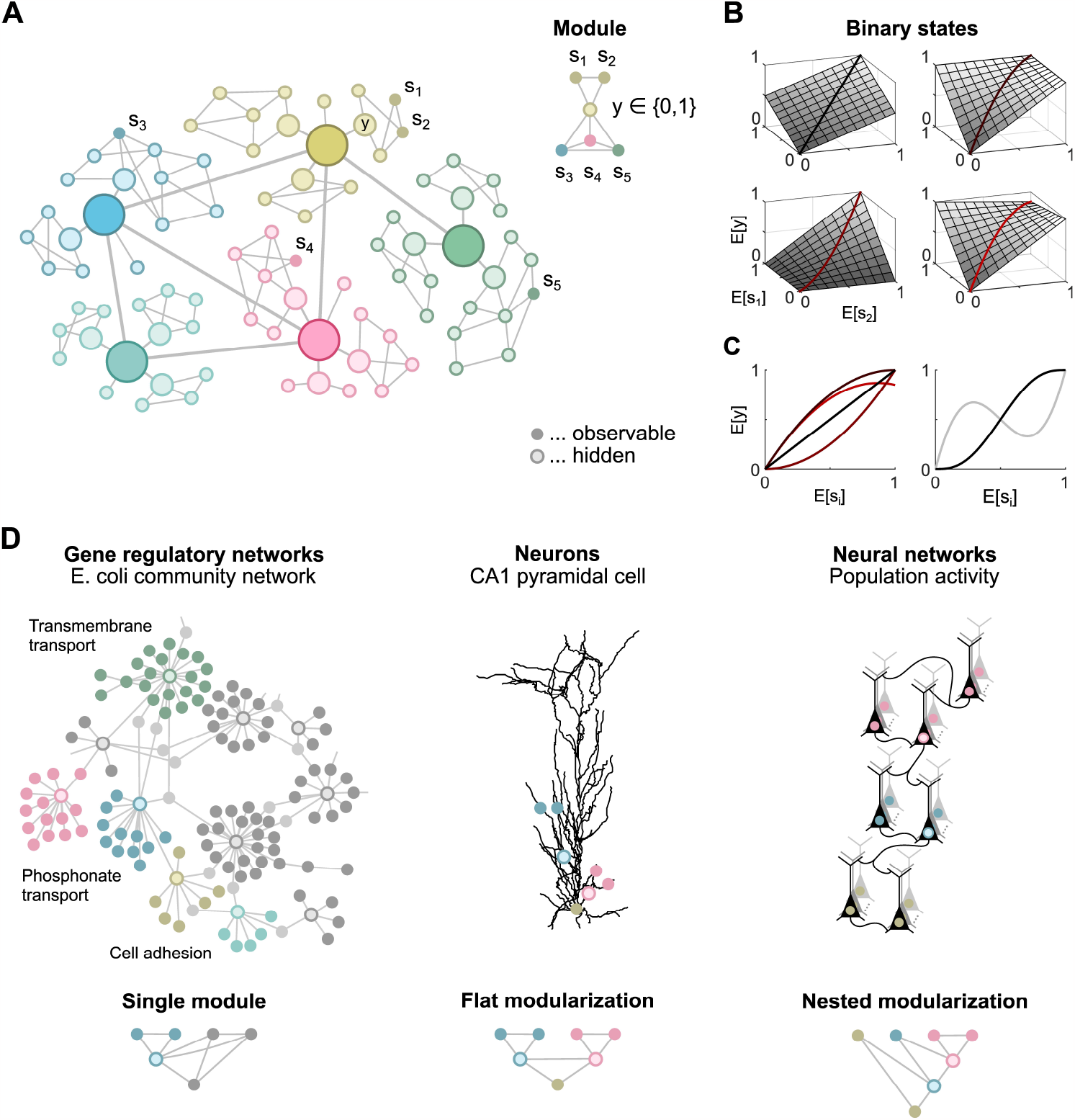
**A:** Undirected graphical model representing dependencies between network components (left). Right: Functional module consisting of the observable components *s*_1_ and *s*_2_, which are independent of all other observable components given the interface variable *y*. **B:** Examples of interface rate functions for the functional module shown in A. **C:** Left: Three of the interface rates functions shown in B are nonlinear, as illustrated for *E*[*s*_1_] = *E*[*s*_2_]. Right: Examples of nonlinear interface rate functions of three (black line) and five (gray line) arguments, illustrated for identical arguments *E*[*s*_*i*_]. **D:** Three scenarios used for statistical testing.

However, probabilistic inference of functional modules in large biological networks is challenging. It is often not possible to record from the entire network, and interface variables may be inaccessible. Moreover, these variables can be abstract quantities such as sensory, associative, motor, or cognitive information encoded in the activity of cell populations [11]. And even if the interface variables are recorded, their identification for large networks is computationally intractable for combinatorial reasons. Here, we bypass these problems and investigate whether it is possible to infer functional modularizations from the distribution of observable network components alone, without information about the organization and values of interface variables.

Remarkably, an arbitrary scalar interface variable does not impose any experimentally testable conditions for continuous network states. This is easy to see, as each of the finitely many samples in a dataset can always be mapped to different values of a scalar variable, thus allowing any functional modularization. To avoid this trivial solution, we constrain the interface variables and focus on the simplest case of binary variables.

We show that the functional organization of networks with latent binary interface variables can be inferred from the statistical moments of observable network components alone, and derive a statistical test for hierarchical modularizations. Importantly, this test can also be applied to refute functional modularizations of networks consisting of continuous scalar interface variables if the following two conditions are met.

First, only the mean values of the continuous scalar interface variables and observable components are relevant for the function of the network and thus for its modularization. The actual distribution of the network states conditioned on these mean values is arbitrary as long as it is consistent with the modularization. For many stochastic biological systems, this condition is assumed to be satisfied, e.g., in molecular biology by the rate of gene transcription [12] and in neuroscience by the instantaneous firing rate of neurons [13]. Second, the mean of a variable downstream of an interface variable depends only linearly on the mean of that interface variable, where downstream refers to any sampling scheme (Fig. 1B). The specific shape of this linear function may depend on other variables, allowing for distributed nonlinear computing (Fig. 1C). In particular for modularizations where a subnetwork depends on the interface variables of several disjoint functional modules, this assumption is satisfied for arbitrary continuous interface variables as long as each of the interface variables contributes only linearly to the mean of each subnetwork component.

Although these assumptions limit the applicability of the method, it is relevant for a number of biological networks. Both assumptions are met by probabilistic Boolean networks, where uncertainties about binary network states are encoded by mean values. Moreover, these assumptions are reasonable when network components are well connected such that a single input has only a small, approximately linear effect on the overall nonlinear activity of a component. Here, we show that the statistical test for modularization is applicable to three biological networks at different spatial scales, and evaluate key hypotheses about their underlying functional organization (Fig. 1D).

## 2 Results

We describe the observable network components by random vectors **s** = (*s*_1_, …, *s*_*d*_) in ℝ^*d*^ and functional modules by sets *S* _*n*_ for *n* = 1, 2, … that contain the indices of all observable components within a module. Associated with each functional module *S* _*n*_ is a potentially hidden binary interface variable *y*_*n*_ that separates its internal components from all other components such that all internal components indexed by *S* _*n*_ are conditionally independent of all other components given *y*_*n*_ (Fig. 1A). A modularization consists of several functional modules and is described by a set ℳ = {*S* _1_, *S* _2_, …}.

To allow the construction of an efficient statistical test, we consider only hierarchically organized modularizations that are either flat or nested. We call a modularization ℳ flat if all functional modules contained in ℳ are disjoint (Fig. 1D). And we call a modularizationℳ nested if all functional modules contained in ℳ are either disjoint or a subset of another functional module in ℳ (Fig. 1D). To clearly distinguish between flat and non-flat nested modularizations, we single out one component, denoted as *s*_ref_, that is not part of any functional module. In the following, *s*_ref_ refers to *s*_*d*_.

The key question here is how to infer flat or nested modularizations from samples of **s** without information about the underlying interface variables **y**, which prevents direct testing of the corresponding conditional independencies.

### Moment ratios indicate functional modules

We show that moments of **s** uniquely determine whether the observable states form a particular modularization or not. As a precondition, we assume that the observable states are bounded and, thus, the probability distribution of **s** is uniquely determined by its moments. We consider raw mixed moments defined as an expectation of the corresponding monomials in **s**, e.g. *E*[*s*_1_ *s*_2_].

The test is based on the following property of single functional modules. Let *P*_*n*_ for *n* = 1, 2, denote an infinite sequence of all monomials in the observable components within a given functional module, e.g. for the module in Fig. 1A, the sequence *P*_1_ = 1, *P*_2_ = *s*_1_, *P*_3_ = *s*_1_ *s*_2_, … . Moreover, let *Q*_*m*_ for *m* = 1, 2, denote the correspond-ing sequence of all monomials in the observable components outside of the functional module, e.g. for the module in Fig. 1A, the sequence *Q*_1_ = 1, *Q*_2_ = *s*_3_, *Q*_3_ = *s*_3_ *s*_4_, …. If the 2×2 matrix *M*_*i j*_ = *E*[*P*_*i*_*Q*_*j*_] for *i, j* = 1, 2 is invertible, then a necessary and sufficient condition for the existence of a binary interface variable *y* is

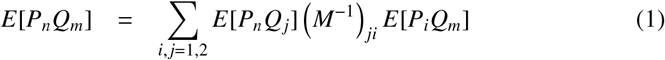

for all *n, m*. Intuitively, the expectation *E*[*P*_*n*_*Q*_*m*_] can be decomposed into multiple expectations, each conditioned on a particular value of the interface variable. As *P*_*n*_ and *Q*_*m*_ are conditionally independent given the interface variable, each of these conditional expectations factorizes into the product of two expectations and the moment corresponds to a scalar product of two vectors. Analogously, a necessary and sufficient condition for the existence of a nested modularization is that this condition holds for each functional module in the modularization.

Although a precise identification of modularizations requires an infinite number of moments, these conditions can be tested for finitely many moments to potentially falsify modularizations. If all observable states have zero mean, the test can be simplified to the condition that certain ratios of moments of **s**, if finite, have identical values. The specific choice of moments and ratios depends on the modularization. To test for a single functional module *S*, we use the correlations *E*[*s*_*k*_ *s*_ref_], *E*[*s*_*l*_ *s*_ref_] and *E*[*s*_*k*_ *s*_*l*_ *s*_ref_], where *k* and *l* index all observable components inside and outside of *S*, respectively, and *s*_*l*_ never refers to *s*_ref_. To enable efficient testing, we introduce a matrix **B** with elements

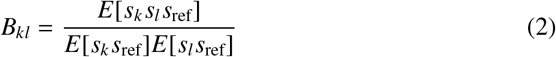

for *k, l ≠ d*. Observable components **s** with (finite) moments as above, a functional module *S* and a binary interface variable exist if and only if for each *l* indexing an observable component outside of the functional module *S*, the moment ratios *B*_*kl*_ have the same value for all *k* in *S* (Fig. 2AB).

**Figure 2:**
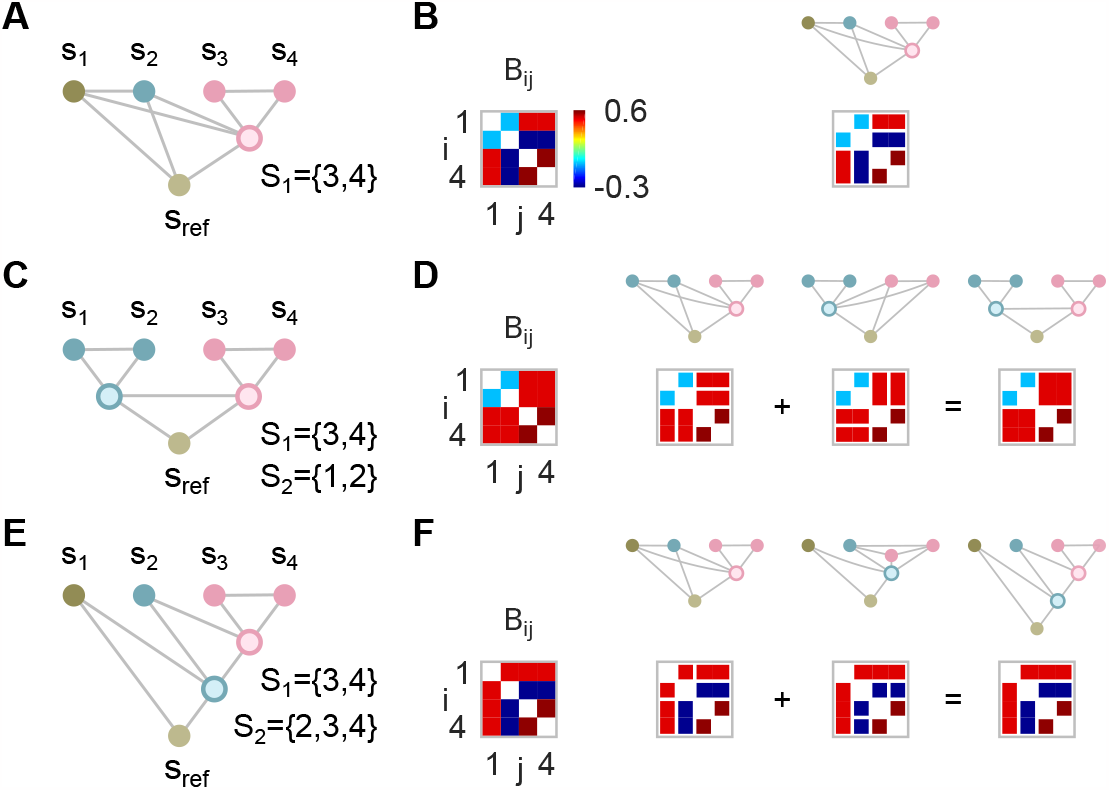
**A:** Single functional module. **B:** Example of a moment ratio matrix for the modularization shown in A (left). Right: Matrix elements with the same value due to the single functional module are shown connected. **C:** Flat modularization. **D:** Example of a moment ratio matrix for the modularization shown in C (left). Right: Each of the two modules implies that different elements of the moment ratio matrix have the same value. The conditions for the moment ratio matrix resulting from the overall modularization are obtained by combining the conditions for the individual modules. **E:** Nested modularization. **F:** The conditions for the moment ratio matrix resulting from the nested modularization shown in E are obtained by combining the conditions for the individual modules.

Moreover, for flat and nested modularizations consisting of several functional modules, the combined conditions can be derived directly from the conditions of the individual functional modules (Fig. 2CD). Observable components **s** with moments analogous to those above, a nested (flat) modularization ℳ and a binary interface variable exist only if (if and only if) the corresponding combined conditions are met. For single functional modules, an even simpler test can be derived using only pairwise correlations, as described in the next section.

Based on estimators 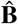 of the moment ratio matrix **B** and estimators of the covariance matrix of the elements of 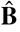, we derive an asymptotic test for modularizations that is valid even if some moment ratios do not exist (see Methods). Furthermore, we consolidate numerically that this test is not only asymptotically correct but holds for sufficiently many samples such that the moment estimates are approximately joint normal. For the investigated datasets, a few hundred samples turn out to be sufficient to fulfill these requirements.

### Inferring functional modules in gene regulatory networks

As a first example, we infer gene regulatory networks from gene expression data, as their function has already been successfully modeled and analyzed by probabilistic Boolean networks [14, 15]. In order to test in a competitive environment, we retroactively participate in the DREAM5 Challenge [16], a comprehensive evaluation of 35 network inference methods on various datasets. The task is to predict direct regulatory interactions between transcription factors (TFs) and their target genes (TGs), where all predicted interactions form the gene regulatory network and the expression values of TFs and TGs constitute the network components. Here, the interface variables of functional modules are non-binary and not hidden, but correspond to recorded expression values of TFs, allowing a comparison with inference methods that rely on this information.

The reconstructed networks are compared to experimentally established gold standards for two datasets, Escherichia coli and an in-silico benchmark. The submission format of the DREAM5 Challenge is a ranked list of predicted regulatory interactions. The performance is evaluated using the area under the precision-recall curve (AUPR), the receiver operating characteristic curve (AUROC) and an overall score that summarizes the performance across networks. Because TF-TF interactions are not organized hierarchically, we restrict the reconstruction to TF-TG interactions, which represent more than 94% of the gold standard. For a fair comparison with previous results, we evaluate all performance measures against the full gold standard.

To apply the test, we start with a ranked list of TF-TG interactions, ordered by the absolute value of their Pearson correlation coefficient, and investigate whether some of the correlations can be explained by indirect interactions through other TFs. More specifically, we investigate all subnetworks consisting of two TFs and two TGs and test for a functional module *S* containing both TFs. Based on the test, the rank of each TF-TG interaction is re-evaluated in such a way that evidence against a functional module, i.e. against an indirect interaction, shifts the rank towards more likely interactions, and reduced evidence shifts the rank in the opposite direction (see Methods).

For each subnetwork (Fig. 3A), we denote the two TFs as *s*_1_ and *s*_2_, the two TGs as *s*_3_ and *s*_ref_, and use the correlations *E*[*s*_*k*_ *s*_*l*_] to test for the functional module *S* = {1, 2}, where *k* and *l* index the observable components inside and outside of the functional module, respectively. For finite moment ratios, observable components with these correlations, a functional module *S* and a binary interface variable *y* exist if and only if

**Figure 3:**
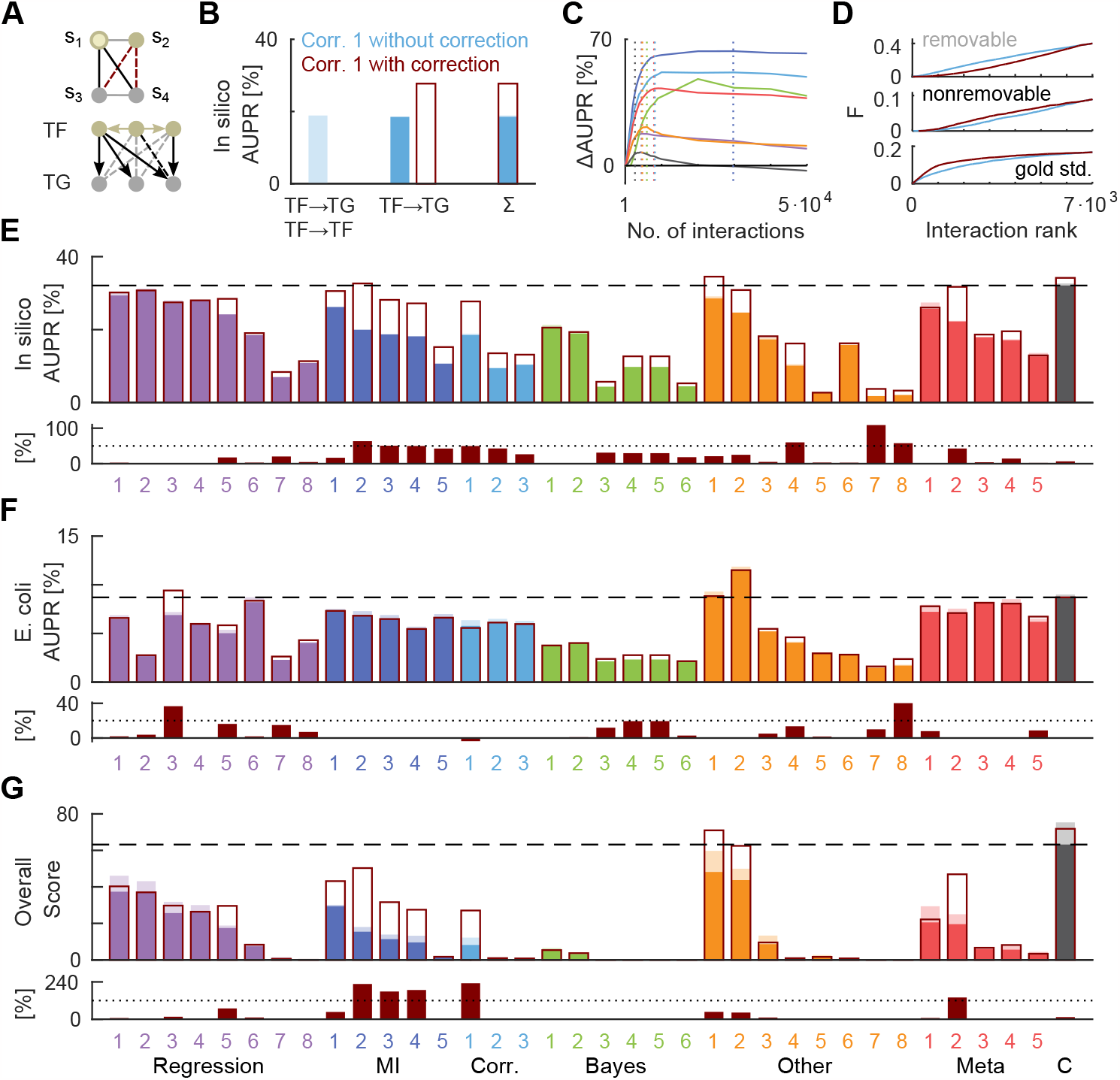
**A:** Undirected graphical model of a subnetwork consisting of four genes (top). Black edges represent putative direct interactions, which are elements of the set of most likely interactions. Red edges represent putative indirect interactions to be tested. Bottom: Example of a directed graphical model representing causal interactions between genes (arrows). Dashed lines indicate removable (gray) and nonremovable (black) indirect interactions. **B:** Performance of the uncorrected method based on Pearson correlation coefficients (Corr. 1) for all interactions (light blue) or only for TF-TG interactions (blue) of the in-silico benchmark. The correction based on the test improves the performance by 50% (dark red rectangle). **C:** Performance as a function of the size of the set of most likely interactions for the network inference methods Regression 5, MI 2, Corr. 1, Bayes 4, Other 1, Meta 2 and the community network (color code as in E). Dotted lines indicate the selected sizes of the sets, determined on a disjoint holdout set. **D:** Cumulative distributions of removable, nonremovable and gold standard interactions as a function of the rank in the list of most likely interactions for the corrected (blue line) and uncorrected (dark red line) inference method Corr. 1. **E:** Performance of all 36 inference methods of the DREAM5 challenge before (colored bars) and after their correction based on the test (dark red rectangles) for the in-silico benchmark (top). The dashed line indicates the performance of the community network (C). Bottom: Relative performance improvement in %. The dotted line indicates an improvement of 50%. **F:** Same as E, but for the Escherichia coli dataset. **G:** Overall score summarizing the performance across networks and performance measures.

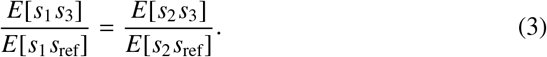

The test is only applied if a subnetwork is sufficiently connected such that at least three of the four TF-TG interactions, including both with the putative interface variable, are in the set of the most likely interactions. The size of this set is a free parameter. The method tests for indirect TF-TG interactions caused by a single functional module. Thus, there are no constraints on the interface rate functions as long as the interdependence of the co-regulated TGs *s*_3_ and *s*_ref_ is linear.

Fig. 3B shows the performance (AUPR) of the uncorrected reconstruction of the in-silico network based on all TF-TF and TF-TG interactions ordered by the absolute value of their Pearson correlation coefficients. Omitting all TF-TF interactions from the reconstruction results in about the same performance, while correcting according to the test improves the AUPR by 50%. The test doesn’t require fine-tuning of its only free parameter, the number of most likely interactions that determine whether a subnetwork is sufficiently connected, which is optimized on a holdout set (Fig. 3C).

The effect of the correction can be analyzed in terms of the rates of true negatives and false positives (type I errors). If a TG is regulated by several interdependent TFs, the test might fail to refute a false direct TF-TG interaction because the corresponding functional module has not one but several interface variables. If this is the case according to the gold standard, we call indirect TF-TG interactions nonremovable (Fig. 3A), which account for less than 30% of all TF-TG dependencies. Fig. 3D shows that the improvement in performance is due to a majority of removable indirect interactions, whose rank distribution is correctly shifted towards less likely interactions. In contrast, the rank distribution of nonremovable indirect interactions is shifted towards more likely interactions, introducing more likely false positives (type I errors). Overall, the rank distribution of the gold standard is shifted toward more likely interactions.

As the correction only requires a ranked list of predicted regulatory interactions, we apply it to each of the inference methods reported in [16] for the in-silico (Fig. 3E) and E. coli microarray data (Fig. 3F). In general, the correction improves most of the 36 inference methods, suggesting that it takes advantage of otherwise unexploited information. In particular, the correction improves the overall score of the community network, which is about the same as that of the single corrected inference method *Genie3* [17], denoted as *Other 1*. However, our aim is not to develop a single best inference method for gene regulatory networks, which will probably be a combination of different inference methods. Rather, we show that this inference method is generally suitable for reconstructing gene regulatory networks and propose it for datasets with missing or unknown regulatory TFs.

### Inferring flat modularizations in dendrites of pyramidal neurons

Pyramidal neurons exhibit complex morphologies and spatially modulated distributions of ion channels [18, 19] that generate regenerative events, such as Na^+^ or NMDA spikes, localized to specific branches or subtrees [20]. Previous work has suggested that these branches act as independent functional modules [21–25], whose responses to local synaptic inputs are linearly summed at the soma. Simulation studies have confirmed that the resulting flat modularization is indeed an accurate description for computations on firing rates [26, 27], where the input and the response are encoded by the rate of synaptic inputs and somatic action potentials, respectively. However, these studies were limited to paired branch stimulation and required complete information about synaptic inputs.

Here, we simultaneously infer multiple functional modules in a neuron model driven by arbitrary synaptic input. More specifically, we simulate a detailed multi-compartment model of a CA1 pyramidal neuron [22] and stimulate excitatory synapses at terminal branches at a constant rate. The network components correspond to sub-threshold membrane potentials at 26 locations in the proximal and oblique apical dendrites, recorded at 50 ms intervals over 20 or 60 minutes (Fig. 4AB) to investigate the statistical power of the test for different sample sizes. The interface variables are non-binary, latent and correspond to membrane potentials within functionally independent dendritic compartments downstream of the recording sites that are linearly summed at the soma. As the somatic module is linear and we are testing for flat modularizations, there are no constraints on the interface rate functions.

**Figure 4:**
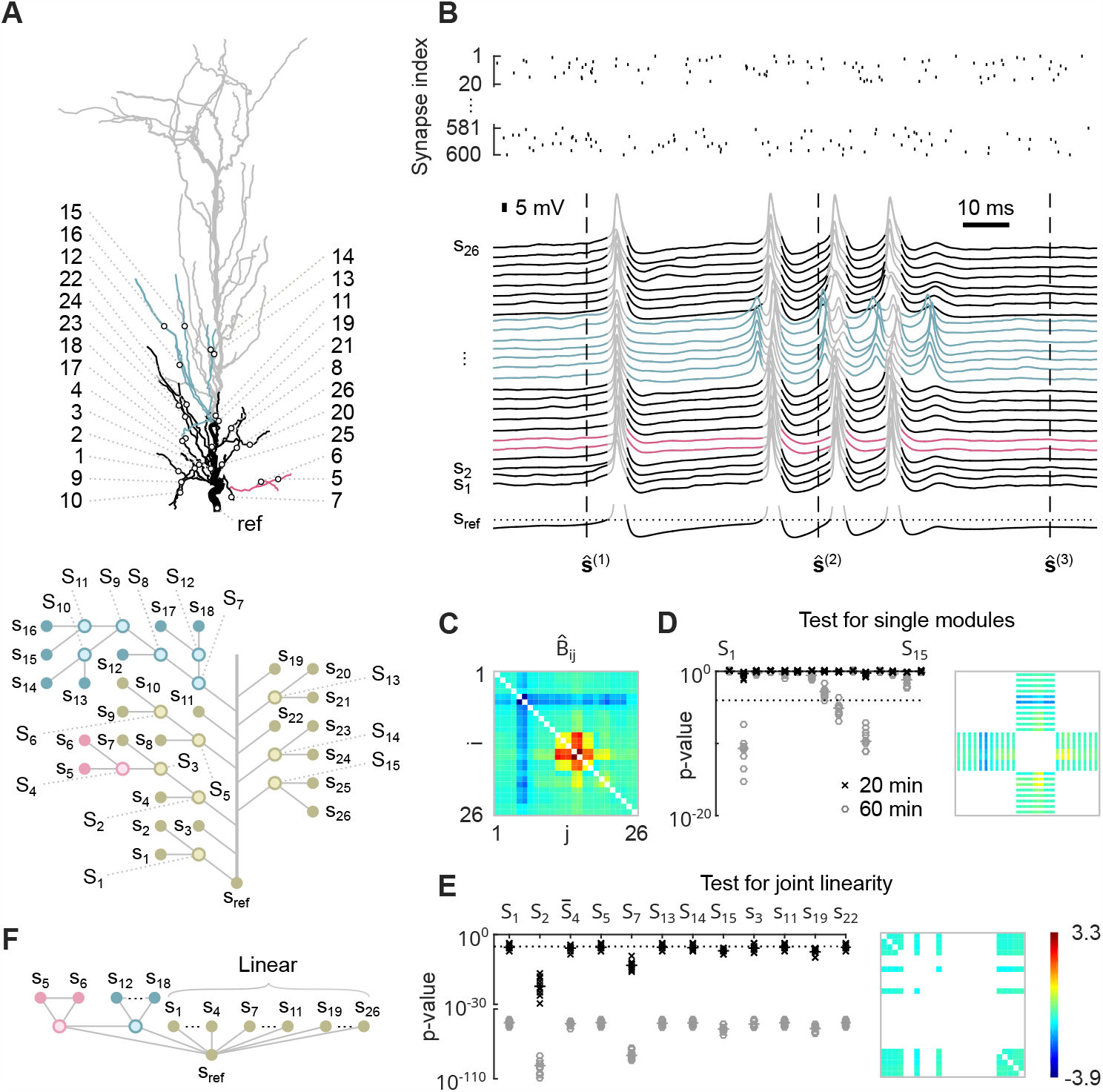
**A:** Model of the apical dendrites of a CA1 pyramidal neuron (top). The investigated subtree is shown in color and black. Black circles indicate recording locations. Bottom: Undirected graphical model of the largest possible modularization consistent with the morphology of the investigated subtree. **B:** Synaptic input to terminal branches (top). Bottom: Observable states are membrane potentials recorded at the corresponding locations shown in A, sampled every 50 ms. Spiking activity at the soma (gray traces) is excluded from the analysis. **C:** Estimated moment ratio matrix. **D:** *p*-values for testing each of the 15 functional modules shown in A on a log scale (left). Right: Elements of the moment ratio matrix expected to have the same value according to the single module *S* _7_, represented as contiguous blocks. **E:** *p*-values for testing whether individual functional modules or dendritic branches originating from the trunk participate in a large linear somatic module (on log scales). All tests are repeated 10 times on independent datasets obtained from 20 and 60 min recordings. Dotted lines represent overall significance levels of 0.01. Horizontal bars indicate medians. Right: Elements of the moment ratio matrix expected to have the same value according to a linear module consisting of *S* _1_, *S* _14_, *S* _15_, *s*_3_, *s*_11_, *s*_22_ and 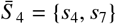 . **F:** Approximate flat modularization of the proximal apical and oblique dendrites.

The estimated moment ratio matrix 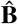 (Equ. 2), obtained from 60 min recordings, shows three nonlinear functional modules, i.e. *S* _4_, *S* _7_ and *S* _11_ (Fig. 4C). The test rejects only three individual functional modules (Fig. 4D) at an overall significance level of 0.01. The first, *S* _2_, is rejected because its two most proximal observable components, *s*_4_ and *s*_7_, are part of a large linear somatic module, while its two distal observable components, *s*_5_ and *s*_6_, form the nonlinear module *S* _4_. The other two rejected modules, *S* _9_ and *S* _11_, are part of the large nonlinear module *S* _7_.

In addition, we investigate which of the functional modules are purely linear, as these can be integrated into a large somatic module. For a purely linear module *S*, the square submatrix of **B** indexed by *S* has identical off-diagonal elements. Then, any functional modularization within *S* is possible, reflecting the commutative property of addition, as multiplication can be excluded for somatic integration. For each functional module, we test whether it is part of a larger linear module consisting of 11 observable components. The constraint ensures the same degrees of freedom for all tests. For 20 min recordings, only modules 2 and 7 are not part of the large linear somatic module, in contrast to the proximal part of module 2 labeled 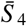 (Fig. 4E).

The resulting flat functional modularization of the proximal apical dendrite consists of a linear somatic module and the two nonlinear modules *S* _4_ and *S* _7_ (Fig. 4F).

However, the large linear somatic module is only approximate and rejected for large enough sample sizes.

### Inferring nested modularizations in neural networks

A classical hypothesis in neuroscience is that neurons convey information only through their instantaneous firing rates, typically characterized by the spike count within a certain time window or population [13, 28, 29]. As a final example, we infer the hierarchical organization of a neural network from simultaneously recorded spiking activity. For this purpose, we simulate five populations of 10 neurons each that interact according to the undirected graphical model shown in Fig. 5A. The corresponding interface rate functions are shown in Fig. 1B (see Methods) and can be implemented by a probabilistic Boolean network.

**Figure 5:**
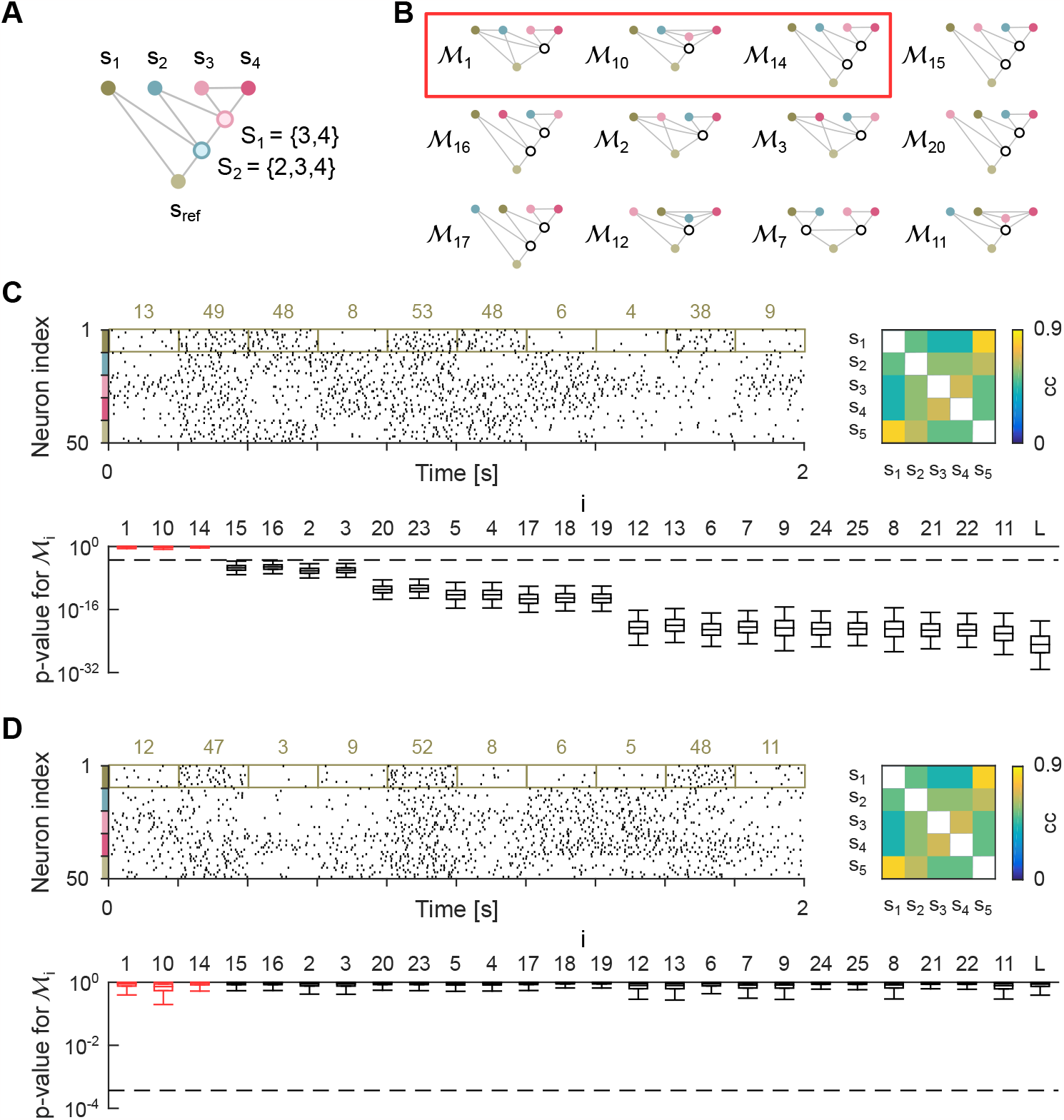
**A:** Nested modularization consisting of five populations of neurons. **B:** 12 out of 26 potential modularizations. Modularizations consistent with the modularization shown in A are marked with a red rectangle. **C:** Spiking activity of all neurons and correlation matrix of observable components (top). Observable states are spike counts within populations and time intervals. Colored numbers represent samples of the observable component *s*_1_. Bottom: Boxplot showing the distribution of *p*-values for testing each of the 26 modularizations. Boxes represent the first through third quartiles, and whiskers indicate the 2.5 and 97.5 percentiles. **D:** Same as C, but for a linear modularization with the same correlation matrix. Dashed lines represent overall significance levels of 0.01.

The task is to infer all modularizations that are consistent with the data (Fig. 5B), given recorded spiking activity over 15 min. For simplicity, we follow a classical population coding approach and define the observable network components as the total number of spikes of all neurons recorded within a population during consecutive 200 ms time intervals (Fig. 5C). Likewise, the interface variables are spike counts for latent populations. The corresponding moment ratio matrix 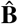 (Equ. 2) is shown in Fig. 2F, which is consistent with the three modularizations ℳ_1_, ℳ_10_ andℳ_14_. In contrast, all other modularizations, including a purely linear model ℳ_L_, are rejected by the test at an overall significance level of 0.01 (Fig. 5C).

The test relies on a small number of correlations between observable components to efficiently investigate potential nonlinear dependencies with *s*_ref_. For this approach, pairwise correlations alone are not sufficient to refute any nonlinear modularization. There always exists a random vector with the same pairwise correlations as the observable states that allows any modularization (Fig. 5D).

## 3 Discussion

Unraveling the functional organization of large biological networks is challenged by incomplete information and combinatorial problems. We present an asymptotic test for hierarchical functional organization of network components based on observable correlations alone, which requires no information about latent network components.

The method presented here differs significantly from previous approaches to inferring functional structure in large networks because it is not based on optimization. In neuroscience, network connectivity and hidden variables are traditionally inferred from neuronal activity based on principles such as maximum a posteriori estimation, Bayesian inference, or information theory [2, 30, 31]. However, such optimization paradigms require regularization in the form of prior information or otherwise prefer more complex structures due to overfitting. Moreover, there is no indication of whether unexplained network activity is due to noise or an inadequate (hierarchical) structure of the latent variables.

In contrast, constraint-based approaches, which first test for conditional independence in the data and then find appropriate network structures, provide intuitive results such as *p*-values, but cannot be applied to incomplete data [10]. The method presented here combines the advantages of constraint-based approaches and probabilistic models by providing a statistical test for partially latent network structures. In particular, the method tests not only necessary conditions, potentially refuting any false nested modularization, but also sufficient conditions for flat modularizations of observable network components with the same correlations as used for the test.

In molecular biology, probabilistic Boolean network models have been successfully applied to infer gene regulatory networks [32–34], but these methods aim at a complete network reconstruction including all logical relationships between genes (logic gates). In contrast, the statistical test presented here does not estimate model parameters. For flat modularizations, only a minimal set of sufficient and necessary conditions is considered, in the sense that omitting a single correlation estimate renders the test inconclusive. We therefore believe it is particularly well suited for small sample sizes, or equivalently, for inferring large functional organizations from datasets of a given size. In particular, the method is useful as a first step in the analysis process to gain an initial understanding of the functional organization of a network.

The test is based on correlations between network components, which are usually reflected in correlated moment ratios. The stronger these correlations are, the lower is the statistical power of the test. For moderately correlated network components, as in the neural network inference example, recording durations of about 15 min are sufficient to achieve adequate statistical power. However, the highly correlated network components of the pyramidal neuron model require recording durations of 60 min. In particular, for synaptic integration in the tuft dendrites, the statistical power is too low to refute any functional module for the chosen recording durations.

Probabilistic Boolean networks with univariate interface variables can not capture the complex dependencies between components of many biological neural networks. However, the assumptions may be well satisfied for sensory systems if neuronal populations implement optimal coding schemes for information processing on short time scales [35, 36]. Then, the optimal neuronal response functions are binary and intermediate rates reflect states of uncertainty. In particular, binary response functions have been shown to be reasonable approximations for various sensory domains [37].

Univariate binary interface variables allow information to flow in only one direction. Interface variables with a larger number of values can capture more complex network dependencies that exhibit bidirectional information flow, multidimensional interface variables or noise correlations [38]. In particular, the lower performance in reconstructing the E. coli gene regulatory network compared to the in-silico benchmark may be due to noise correlations of expression levels caused by sample preparation, array fabrication, and array processing [39, 40]. An extension to interface variables with four or more values seems feasible and promising, since the necessary conditions for corresponding functional modularizations are already derived in the Suppl. Material.

In general, exact probabilistic inference is intractable in large biological networks. We provide a hypothesis-driven statistical method that efficiently tests for selected functional modularizations and does not require complete information about the entire network. With recent advances in high-throughput single-cell technology [41], multi-electrode array technology [42, 43], two-photon microscopy [44, 45] and genetically encoded voltage indicators [46, 47], our mathematical framework can be applied to a wide range of datasets to facilitate the analysis of complex biological systems.

## 4 Methods

### A statistical test for modularizations

Let *N*^(t)^ denote the total number of i.i.d. samples of **s** and let **ŝ**^(*n*)^ for *n* ∈{1, …, *N*^(t)^} denote the *n*-th sample. We assume *N*^(t)^ is even and the samples are normalized to zero mean. In this case, the test can be further simplified to the condition that certain ratios of moments of **s** have identical values if they are finite.

Let *d*^(χ0^) denote the total number of moment ratios used for the test and let the components of 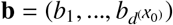 denote these ratios. For a given modularizationℳ, we introduce the index sets 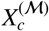 for *c* ∈{1, …, *d*^(χ)^} such that all components of **b** indexed by a set have the same value. The specific choice of moments, their ratios used for the test and the definition of the index sets 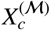 depend on the type of modularization, i.e., single functional module, flat or nested modularization, and is described in the sections below (see Suppl. Material for details).

For each sample moment ratio 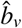 for *v* ∈{ 1, …, *d*^(χ0^)} we define a vector 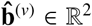 containing the numerator and the denominator of 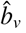. We estimate the moment ratio *b*_*v*_ by

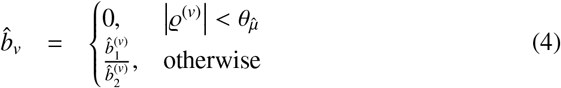

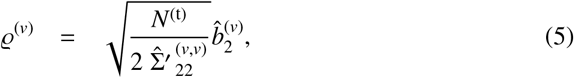

where a cutoff 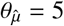 ensures finite expectations of 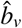 for joint normal 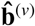.

The simplest possible moments for testing nested modularizations are 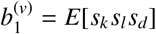and 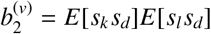 for *k≠ l*, which are estimated by the sample moments

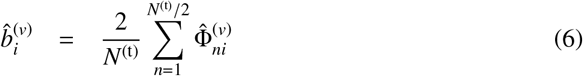

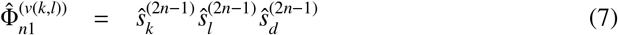

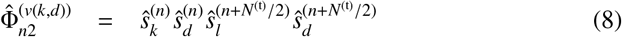

for *i* ∈ {1, 2}, *k* ∈ {1, …, *l* − 1}, *l* ∈ {2, …, *d* − 1} and vectorization

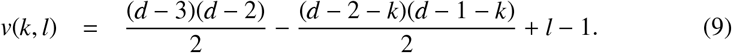

The corresponding sample covariance matrices are

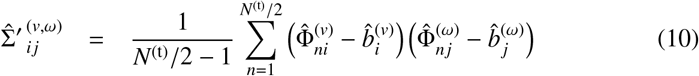

for *i, j* ∈ {1, 2} and *v, ω* ∈ {1, …, *d*^(χ0^)}, where *ω*(*k, l*) = *v*(*k, l*). The covariance matrix of the sample moment ratios 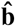 can be approximated by

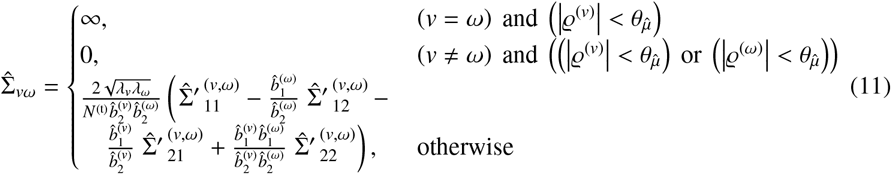

for *v, ω*∈{1, …, *d*^(χ0^)}.

The simplest possible moments for testing single functional modules (in gene regulatory networks) are 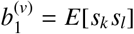 and 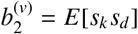 for *k≠ l* estimated by

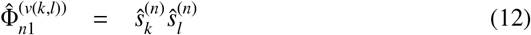

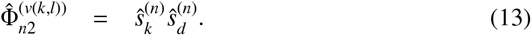

We show that the statistic

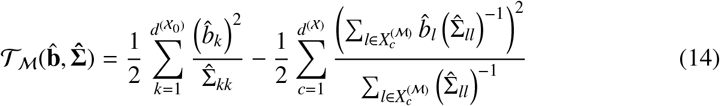

can be used to test for modularization ℳ even if some moment ratios don’t exist because of zero denominators. The test is asymptotically correct for *λ*_*v*_ = 1. However, we consolidate numerically that the test also applies to finitely many samples if the moment estimates are approximately joint normal, 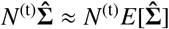 and

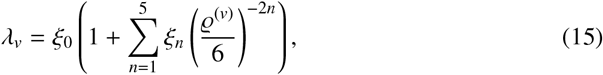

where (*ξ*_0_, …, *ξ*_5_) = (1.367, 2.047, 4.735, −1.923, −1.231, 2.790).

For the most general test, we introduce a scaling factor *λ*^(max)^, which corrects for correlated components of 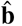, and constrain the nominal significance level *α*^(*)^ to be larger than a minimal nominal significance level *α*^(min)^, which corrects for potential zero correlations between *s*_ref_ and other observable components.

Let ℳ_*i*_ denote the *i*-th of *d*^(ℋ)^ modularizations tested on the same samples **ŝ**. If the observable states form the modularization ℳ_*i*_, the probability of sampling 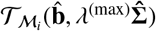 at least as extreme as observed is less than *α*^(*)^ if

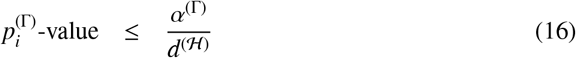

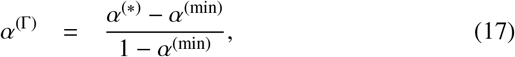

where 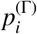-value denotes the probability of sampling 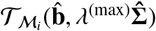 at least as extreme as observed when distributed according to the gamma distribution 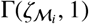 for shape parameter 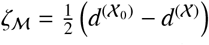 and scale parameter 1. For *ζ*_M_ = 0, 𝒯_M_ = 0.

For potentially correlated components of the sample moment ratios 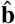, we set *λ*^(max)^ such that the statistical test remains conservative. The smallest possible *λ*^(max)^ with this property is the largest eigenvalue of a submatrix of 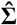 normalized to unit diagonal, i.e.,

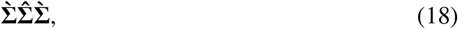

where the matrix 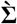 has the same size as 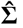, is diagonal and 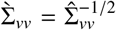 if *v* is element of an index set of size larger than one and 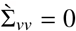, otherwise. A more power full test is derived in Section 2.4 of the Suppl. Material. The minimal nominal significance level

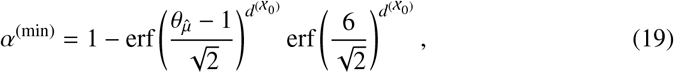

where erf denotes the error function.

If all moment rations are finite, the test is asymptotically consistent for flat modularizations, i.e., its power for any incorrect modularization converges asymptotically to one. If in addition all moment ratios are uncorrelated, the test statistic is asymptotically minimal sufficient, i.e., it most efficiently captures all information about a modularization contained in the sample moment ratios. More details and the derivative of the statistical test can be found in the Suppl. Material.

Finally, if both conditions in the introduction are met, the inference for modularizations with binary and continuous interface variables is equivalent. Given that *s*_ref_ is considered as a single module, a modularization with continuous interface variables exists if and only if a corresponding modularization with binary interface variables and the same correlations as used for the test exists.

### Inference in gene regulatory networks

We participated in the transcriptional network inference challenge from DREAM5 that compares 35 methods for inference of gene regulatory networks: 29 submitted by participants and additional 6 “off-the-shelf” methods classified into six categories: Regression, Mutual information, Correlation, Bayesian networks, Meta (combinations of several different approaches), and Other (methods not belonging to any of the previous categories). The design of the challenge, detailed methods and results are published in [16].

We evaluate the network reconstruction from gene expression microarray datasets for E. coli and an in-silico benchmark using the area under the precision-recall curve (AUPR), the area under the receiver operating characteristic curve (AUROC), and an overall score defined as the mean of the (log-transformed) network specific p-values (obtained by simulating a null distribution for 25000 random networks),

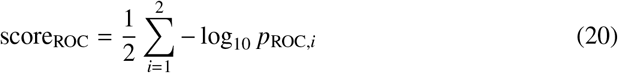

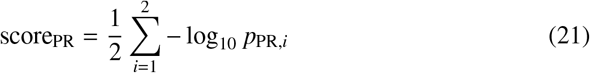

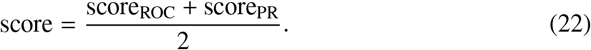

We omit the third dataset of the challenge for S. cerevisiae due to technical reasons, i.e. the size of the network is too large for the algorithm and computer hardware in use.

The Escherichia coli dataset consists of 4511 genes (334 TF) and a gold standard of 3766 TF-TG interactions (94% of 4012 total), according to which 89% of all indirect TF-TG interactions are removable. The in-silico datasets consists of 1643 genes (195 TF) and a gold standard of 1923 TF-TG interactions (94% of 2066 total), according to which 72% of all indirect TF-TG interactions are removable.

Given a sorted list of regulatory interaction *p*-values in ascending order, we apply the test to any four-node subnetwork consisting of two TFs and two TGs. We call a certain number of regulatory interactions with the lowest ranks in the list the set of most likely interactions. We call the TF with the most likely interaction in a four-node sub-network the putative interface variable. We call a subnetwork sufficiently connected if at least three of the four TF-TG interactions, including both with the putative interface variable, are in the set of the most likely interactions. If a subnetwork is sufficiently connected, then each *p*-value of interactions with the other TF (not the putative interface variable) is changed by

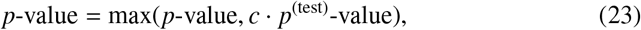

where *p*^(test)^-value denotes the *p*-value of the test. For a scaling factor *c* = 1, the test is conservative for the null hypothesis (no direct interaction), which corresponds to the hypothesis ((no dependency) or (functional module)). We heuristically set *c* to the smallest *p*-value of any interaction in the set of most likely interactions, ensuring that missing evidence against a functional module is not weighted more heavily than the evidence for interactions that determine whether the test is performed at all. The size of the set of most likely interactions is determined by a holdout set consisting of every 8th sample.

To combine tests for the same interaction in different subnetworks, we again take the maximum of the individual *p*-values, which corresponds to a combined conservative test for the logic or operation of the individual hypotheses. If the gene regulatory network is i) nested such that each TG can only by reached from any TF via a single interface variable, ii) all TF-TG interactions are essential, i.e., the removal of a single TF-TG interaction results in additional independencies in **s**, and iii) indirect TF-TG interactions are less correlated than direct TF-TG interactions, then the network reconstruction is asymptotically correct in the sense that the most likely inferred interactions are all true TF-TG interactions.

More precisely, we test for the modularization ℳ = {*S*} consisting of the single functional module *S* = {1, 2} (Fig. 3A) resulting in a single set *X*^(ℳ)^ = {1, 2} that indexes the only two components of the moment ratio vector *b*_*k*_ = *E*[*s*_*k*_ *s*_3_]*/E*[*s*_*k*_ *s*_ref_] for *k* ∈{1, 2}. To avoid corrections for correlated components of the moment ratio vector, we estimate both *b*_*k*_ using disjoint sets of samples. Furthermore, we use the uncorrected asymptotic version of the test, where *λ*_*v*_ = 1, and a small cutoff 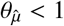.

To apply the method to a ranked list of *N* regulatory interactions, we artificially assign the *p*-values *rank/N*. For the sorted list of Pearson correlation coefficients, this procedure results in roughly the same AUPR, with even a slight performance improvement of 0.29%.

### Inference in pyramidal neurons

We simulated the detailed compartmental model of a CA1 pyramidal neuron developed by Poirazi et al. [22] in the simulation environment NEURON. The model includes various active and passive membrane mechanisms, such as sodium and potassium currents, A-type potassium currents, m-type potassium currents, a hyperpolarization-activated h-current, voltage-dependent calcium currents, and Ca^2+^-dependent potassium currents. The densities and distributions of these currents are based on published data. We are interested in subthreshold synaptic integration and block all spike-generating currents at the soma.

Synaptic inputs consist of an NMDA and an AMPA-type conductance with a ratio of their peak values of 2.5. Each of the 60 terminal branches contains 10 synapses, with equal distances between adjacent synapses or branch ends. Each synapse is stimulated by a Poisson process at a constant rate of 32 Hz. The dendritic spike rate is approximately 28 Hz, which is in the range of values observed experimentally in neocortical pyramidal neurons from freely behaving rats [20].

The datasets consist of the membrane potentials at the soma and the centers of the 26 most proximal terminal branches of the apical dendrites. Samples are recorded for 20 or 60 min at 50 ms time intervals, ensuring that their normalized autocovariance is less than 0.05.

The moment ratio vector 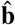 and the index sets *X*^(ℳ)^ are derived from the corresponding elements above the diagonal of 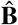 (see Fig. 2D and Suppl. Material). To test for a purely linear functional module *S*, we define an additional set *X*^(ℳ)^ that indexes all elements above the diagonal of the square submatrix of 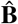 that is indexed by *S* . For each module or branch originating from the trunk, the largest *p*-value is calculated over all combinations of modules and branches containing that module and consisting of 11 observable components to ensure identical degrees of freedoms.

The resulting flat modularization, which consists of the nonlinear functional modules *S* _4_ and *S* _7_ and a complementary linear somatic module (Fig. 4F), cannot be rejected at an overall significance level of 0.01. In contrast, the flat modularization consisting of the functional modules *S* _2_ and *S* _7_ and a complementary linear somatic module can be rejected at an overall significance level of 0.01.

### Inference in neural networks

The neural network consists of *d* = 5 populations, each with 10 neurons. For every 200 ms time interval, the firing rates of all neurons in the *i*-th population are chosen according to a binary random variable *x*_*i*_ ∈{1, 2} for *I* ∈{ 1, …, *d*} such that they are set to 5 Hz if *x*_*i*_ = 1 or 25 Hz, otherwise. The total number of spikes in the i-th population and the n-th time interval defines the observable component 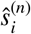 (normalized to zero mean).

We implement the modularization shown in Fig. 5A, which consists of *d*^(ℳ)^ = 2 functional modules, where *s*_ref_ refers to *s*_5_. The binary random vector **x** and the binary interface variables *y*_*c*_ ∈ {1, 2} for *c* ∈ {1, 2} are distributed according to

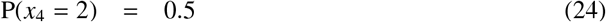

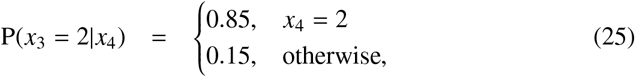

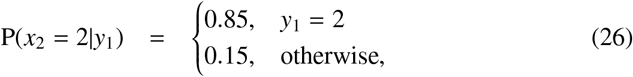

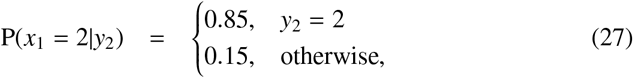

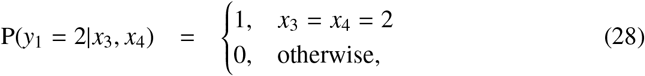

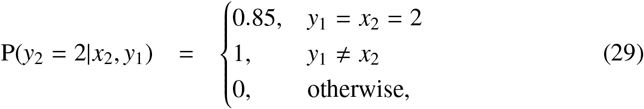

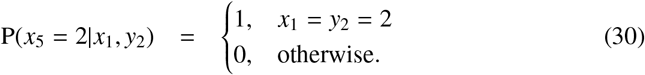

The rate functions for *y*_1_ and *x*_5_ are shown in the bottom left panel and the rate function for *y*_2_ is shown in the bottom right panel of Fig. 1B. An optimal linear model predicting *x*_5_ from the other components of **x** has a coefficient of determination of *R*^2^ = 0.79.

The moment ratio vector 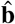 and the index sets *X*^(ℳ)^ are derived from the corresponding elements above the diagonal of 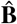(see Fig. 2F and Suppl. Material). To test for an arbitrary linear modularization ℳ_L_, we define an additional set *X*^(ℳ L^) that indexes all elements above the diagonal of 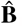 .

All components of **ŝ** are correlated with Pearson correlation coefficients greater than 0.41 (Fig. 5C). To generate a linear modularization with the covariance matrix 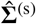 of **ŝ**, we apply the linear transformation **L** obtained from the Cholesky decomposition 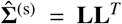 to the time-shuffled observable states (shuffled independently for each population). All tests are repeated 10^4^ times on independent datasets. All 25 modularizations are shown in Fig. S3 in the Suppl. Material.

## Code availability

An implementation of the statistical test in MATLAB is available at Zenodo [48].

## References

[1] Anne E Urai, Brent Doiron, Andrew M Leifer, and Anne K Churchland. Largescale neural recordings call for new insights to link brain and behavior. Nature neuroscience, 25(1):11–19, 2022.

[2] Liam Paninski and John P Cunningham. Neural data science: accelerating the experiment-analysis-theory cycle in large-scale neuroscience. Current opinion in neurobiology, 50:232–241, 2018.

[3] Ed Bullmore and Olaf Sporns. Complex brain networks: graph theoretical analysis of structural and functional systems. Nature reviews neuroscience, 10(3):186–198, 2009.

[4] Edward T Bullmore and Danielle S Bassett. Brain graphs: graphical models of the human brain connectome. Annual review of clinical psychology, 7:113–140, 2011.

[5] Danielle S Bassett and Olaf Sporns. Network neuroscience. Nature neuroscience, 20(3):353–364, 2017.

[6] Mark EJ Newman. The structure and function of complex networks. SIAM review, 45(2):167–256, 2003.

[7] Mark EJ Newman. Communities, modules and large-scale structure in networks. Nature physics, 8(1):25–31, 2012.

[8] GüAnter P Wagner, Mihaela Pavlicev, and James M Cheverud. The road to modularity. Nature Reviews Genetics, 8(12):921–931, 2007.

[9] Michelle Girvan and Mark EJ Newman. Community structure in social and biological networks. Proceedings of the national academy of sciences, 99(12):7821–7826, 2002.

[10] Daphne Koller and Nir Friedman. Probabilistic graphical models: principles and techniques. MIT press, 2009.

[11] Stefano Panzeri, Monica Moroni, Houman Safaai, and Christopher D Harvey. The structures and functions of correlations in neural population codes. Nature Reviews Neuroscience, 23(9):551–567, 2022.

[12] David S Latchman. Transcription factors: an overview. The international journal of biochemistry & cell biology, 29(12):1305–1312, 1997.

[13] Wulfram Gerstner and Werner M Kistler. Spiking neuron models: Single neurons, populations, plasticity. Cambridge university press, 2002.

[14] Guy Karlebach and Ron Shamir. Modelling and analysis of gene regulatory networks. Nature reviews Molecular cell biology, 9(10):770–780, 2008.[1]

[15] Nicolas Le Novere. Quantitative and logic modelling of molecular and gene networks. Nature Reviews Genetics, 16(3):146–158, 2015.

[16] Daniel Marbach, James C Costello, Robert KüAffner, Nicole M Vega, Robert J Prill, Diogo M Camacho, Kyle R Allison, Manolis Kellis, James J Collins, et al. Wisdom of crowds for robust gene network inference. Nature methods, 9(8):796–804, 2012.

[17] Vaân Anh Huynh-Thu, Alexandre Irrthum, Louis Wehenkel, and Pierre Geurts. Inferring regulatory networks from expression data using tree-based methods. PloS one, 5(9):e12776, 2010.

[18] Greg J Stuart and Nelson Spruston. Dendritic integration: 60 years of progress. Nature neuroscience, 18(12):1713–1721, 2015.

[19] Greg Stuart, Nelson Spruston, and Michael HäAusser. Dendrites. Oxford University Press, 2016.

[20] Jason J Moore, Pascal M Ravassard, David Ho, Lavanya Acharya, Ashley L Kees, Cliff Vuong, and Mayank R Mehta. Dynamics of cortical dendritic membrane potential and spikes in freely behaving rats. Science, 355(6331):eaaj1497, 2017.

[21] Kevin A Archie and Bartlett W Mel. A model for intradendritic computation of binocular disparity. Nature neuroscience, 3(1):54–63, 2000.

[22] Panayiota Poirazi, Terrence Brannon, and Bartlett W Mel. Pyramidal neuron as two-layer neural network. Neuron, 37(6):989–999, 2003.

[23] Alon Polsky, Bartlett W Mel, and Jackie Schiller. Computational subunits in thin dendrites of pyramidal cells. Nature neuroscience, 7(6):621–627, 2004.

[24] Yael Katz, Vilas Menon, Daniel A Nicholson, Yuri Geinisman, William L Kath, and Nelson Spruston. Synapse distribution suggests a two-stage model of dendritic integration in ca1 pyramidal neurons. Neuron, 63(2):171–177, 2009.

[25] Tiago Branco, Beverley A Clark, and Michael HäAusser. Dendritic discrimination of temporal input sequences in cortical neurons. Science, 329(5999):1671–1675, 2010.

[26] Bardia F Behabadi and Bartlett W Mel. Mechanisms underlying subunit independence in pyramidal neuron dendrites. Proceedings of the National Academy of Sciences, 111(1):498–503, 2014.

[27] Florian Eberhardt, Andreas VM Herz, and Stefan HäAusler. Tuft dendrites of pyramidal neurons operate as feedback-modulated functional subunits. PLoS Computational Biology, 15(3):e1006757, 2019.

[28] R Christopher Decharms and Anthony Zador. Neural representation and the cortical code. Annual review of neuroscience, 23(1):613–647, 2000.

[29] Frédéric Theunissen and John P Miller. Temporal encoding in nervous systems: a rigorous definition. Journal of computational neuroscience, 2:149–162, 1995.

[30] Greg Ver Steeg and Aram Galstyan. Discovering structure in high-dimensional data through correlation explanation. Advances in Neural Information Processing Systems, 27, 2014.

[31] Concha Bielza and Pedro Larrañaga. Bayesian networks in neuroscience: a survey. Frontiers in computational neuroscience, 8:131, 2014.

[32] Melanie Grieb, Andre Burkovski, J Eric SträAng, Johann M Kraus, Alexander Groß, GüAnther Palm, Michael KüAhl, and Hans A Kestler. Predicting variabilities in cardiac gene expression with a boolean network incorporating uncertainty. PloS one, 10(7):e0131832, 2015.

[33] Seyed Amir Malekpour, Amir Reza Alizad-Rahvar, and Mehdi Sadeghi. Logic-net: probabilistic continuous logics in reconstructing gene regulatory networks. BMC bioinformatics, 21:1–21, 2020.

[34] Seyed Amir Malekpour, Maryam Shahdoust, Rosa Aghdam, and Mehdi Sadeghi. wplogicnet: logic gate and structure inference in gene regulatory networks. Bioinformatics, 39(2):btad072, 2023.

[35] M Bethge, D Rotermund, and K Pawelzik. Optimal neural rate coding leads to bimodal firing rate distributions. Network: Computation in Neural Systems, 14(2):303, 2003.

[36] Alexander P Nikitin, Nigel G Stocks, Robert P Morse, and Mark D McDonnell. Neural population coding is optimized by discrete tuning curves. Physical review letters, 103(13):138101, 2009.

[37] Julijana Gjorgjieva, Markus Meister, and Haim Sompolinsky. Functional diversity among sensory neurons from efficient coding principles. PLoS computational biology, 15(11):e1007476, 2019.

[38] Bruno B Averbeck, Peter E Latham, and Alexandre Pouget. Neural correlations, population coding and computation. Nature reviews neuroscience, 7(5):358–366, 2006.

[39] Alexander J Hartemink, David K Gifford, Tommi S Jaakkola, and Richard A Young. Maximum-likelihood estimation of optimal scaling factors for expression array normalization. In Microarrays: Optical technologies and informatics, volume 4266, pages 132–140. SPIE, 2001.

[40] Benjamin M Bolstad, Rafael A Irizarry, Magnus Åstrand, and Terence P. Speed. A comparison of normalization methods for high density oligonucleotide array data based on variance and bias. Bioinformatics, 19(2):185–193, 2003.

[41] Pau Badia-i Mompel, Lorna Wessels, Sophia MüAller-Dott, Rémi Trimbour, Ricardo O Ramirez Flores, Ricard Argelaguet, and Julio Saez-Rodriguez. Gene regulatory network inference in the era of single-cell multi-omics. Nature Reviews Genetics, pages 1–16, 2023.

[42] Jeffrey Abbott, Tianyang Ye, Keith Krenek, Rona S Gertner, Steven Ban, Youbin Kim, Ling Qin, Wenxuan Wu, Hongkun Park, and Donhee Ham. A nanoelectrode array for obtaining intracellular recordings from thousands of connected neurons. Nature biomedical engineering, 4(2):232–241, 2020.

[43] Angelique C Paulk, Yoav Kfir, Arjun R Khanna, Martina L Mustroph, Eric M Trautmann, Dan J Soper, Sergey D Stavisky, Marleen Welkenhuysen, Barundeb Dutta, Krishna V Shenoy, et al. Large-scale neural recordings with single neuron resolution using neuropixels probes in human cortex. Nature Neuroscience, 25(2):252–263, 2022.

[44] Hillel Adesnik and Lamiae Abdeladim. Probing neural codes with two-photon holographic optogenetics. Nature neuroscience, 24(10):1356–1366, 2021.

[45] Christine Grienberger, Andrea Giovannucci, William Zeiger, and Carlos Portera-Cailliau. Two-photon calcium imaging of neuronal activity. Nature Reviews Methods Primers, 2(1):67, 2022.

[46] Yuki Bando, Michael Wenzel, and Rafael Yuste. Simultaneous two-photon imaging of action potentials and subthreshold inputs in vivo. Nature Communications, 12(1):7229, 2021.

[47] Victor Hugo Cornejo, Netanel Ofer, and Rafael Yuste. Voltage compartmentalization in dendritic spines in vivo. Science, 375(6576):82–86, 2022.

[48] Stefan HäAusler. Zenodo, 10.5281/zenodo.8190172, July 2023.

