## Supplementary material for "Correlations reveal the hierarchical organization of networks with latent binary variables"

Stefan Häusler

Faculty of Biology and Bernstein Center for Computational Neuroscience,
Ludwig-Maximilians-Universität München, Germany

September 25, 2023

### 1 Summary

#### Observable states

We use the standard notations $\mathbb{N} = \{1, 2, ...\}$ and $\mathbb{N}_0 = \{0, 1, 2, ...\}$. Suppose that $(\Omega, \mathcal{F}, \mathbf{P})$ is a probability space on which all random variables and vectors are defined. Let $\mathbf{s} = (s_1, ..., s_d)$ denote a random vector in $\mathbb{R}^d$ of dimension $d \in \mathbb{N}$, where $d > 1$. Corresponding to $\mathbf{s}$ is a probability distribution $\mathrm{P}_{\mathbf{s}}$, or just P, which is a measure on the Borel space $(\mathbb{R}^d, \mathcal{B}^d)$. We refer to $\mathbf{s}$ as *observable states*. We assume all observable states are bounded random variables. The expectation of $\mathbf{s}$ with respect to probability $\mathbf{P}$ is denoted by $E[\mathbf{s}]$.

Superscripts in brackets are either text strings, e.g., $\mathbf{s}^{(\mathrm{i})}$ and $\mathbf{s}^{(\mathrm{j})}$, where $\mathbf{s}^{(\mathrm{i})}$ and $\mathbf{s}^{(\mathrm{j})}$ always denote different variables, or variables, e.g., $\mathbf{s}^{(i)}$ and $\mathbf{s}^{(j)}$ for $i, j \in \mathbb{N}$, where $\mathbf{s}^{(i)}$ and $\mathbf{s}^{(j)}$ denote the same variable if and only if $i = j$. Analogously, $\mathbf{s}$, $\mathbf{s}^{(i)}$ and $\mathbf{s}^{(i,j)}$ denote different variables. Let $N^{(\mathrm{t})}$ denote the total number of i.i.d. samples of $\mathbf{s}$ and let $\hat{\mathbf{s}}^{(n)}, n \in \{1, ..., N^{(\mathrm{t})}\}$ denote the $n$-th sample. We assume $N^{(\mathrm{t})}$ is even and the samples are normalized to zero mean, i.e., $E[s_k] = 0$ for $k \in \{1, ..., d\}$.

#### Functional modules and modularizations

A *functional module* is defined by a non-empty index set $S \subsetneq \{1, ..., d-1\}$. We say the components of $\mathbf{s}$ indexed by $S$ form a functional module if there exists an *interface variable* $y$ such that all components of $\mathbf{s}$ indexed by $S$ are independent of all other components of $\mathbf{s}$ given $y$. In the following we consider functional modules with stochastic binary interface variables $y \in \{1, 2\}$, which potentially encode rates.[1] Let $d^{(\mathcal{M})} \in \mathbb{N}$ denote the total number of functional modules and let $S_c$ for $c \in \{1, ..., d^{(\mathcal{M})}\}$ denote the corresponding index sets.

We refer to all $d^{(\mathcal{M})}$ functional modules as *modularization*, which we define by the set $\mathcal{M} = \{S_1, ..., S_{d^{(\mathcal{M})}}\}$. We call a modularization *flat* if for all $c_1, c_2 \in \{1, ..., d^{(\mathcal{M})}\}$, $c_1 \neq c_2$ implies $S_{c_1} \cap S_{c_1} = \{\}$. Furthermore, we call a modularization *nested* if for all $c_1, c_2 \in \{1, ..., d^{(\mathcal{M})}\}$, $c_1 \neq c_2$ implies ($S_{c_1} \subsetneq S_{c_2}$ or $S_{c_2} \subsetneq S_{c_1}$ or $S_{c_1} \cap S_{c_2} = \{\}$) (see Fig. S1). Any modularization consisting of a single functional module is a flat modularization and any flat modularization is a nested modularization. Furthermore, if the components of $\mathbf{s}$ form a modularization defined by $\mathcal{M}$, they also form all modularizations defined by non-empty subsets of $\mathcal{M}$ (e.g. see Fig. S1A and B).

The component $s_d$ is not part of any functional module, which enables the construction of an efficient statistical test for flat and nested modularizations as shown below. Although this does not constrain the set of testable modularizations,[2] it affects the moments used for the statistical test.

#### Statistical test for modularizations

We show that whether or not observable states form a particular modularization $\mathcal{M}$ (with binary interface variables) can be uniquely determined from the moments of $\mathbf{s}$. Although a precise identification involves an infinite number of moments, the following two null hypotheses can be falsified given a finite number of moments.

(H1) The observable states $\mathbf{s}$ form a nested modularization $\mathcal{M}$.

[1] Lemma 1 applies also to non-binary interface variables with a finite number of states.

[2] The type of modularization might depend on the choice of $s_d$. E.g., the flat modularization shown in Fig. S1B turns into the nested modularization shown in Fig. S1C if $s_2$ and $s_5$ are exchanged.

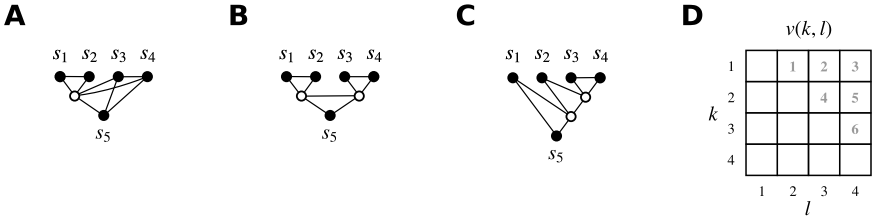


Figure S1: Types of modularizations. Undirected graphical models (Markov networks) representing the dependencies between the components of $\mathbf{s}$ for the single functional module $S = \{1, 2\}$ (**A**), the flat modularization $\mathcal{M} = \{\{1, 2\}, \{3, 4\}\}$ (**B**) and the nested modularization $\mathcal{M} = \{\{3, 4\}, \{2, 3, 4\}\}$ (**C**). Observable components and potentially hidden interface variables are shown as filled and open circles, respectively. The component $s_5$ is not part of any functional module and serves as a reference to uniquely define flat and nested modularizations. **D:** Index $v$ (bold gray numbers) as a function of the indices $k$ and $l$ for $d = 5$.

(H2) There exists a model of $\mathbf{s}$, i.e., another random vector in $\mathbb{R}^d$, with flat modularization $\mathcal{M}$ and the same moments as used for the test.

Under both null hypotheses, certain ratios of population moments of $\mathbf{s}$ have identical values.[3] Let $d^{(\mathcal{X}_0)}$ denote the total number of ratios used for the statistical test and let the components of $\mathbf{b} \in \mathbb{R}^{d^{(\mathcal{X}_0)}}$ denote these ratios. In order to specify the components of $\mathbf{b}$ that have identical values for given $\mathcal{M}$, we partition the index set $\{1, ..., d^{(\mathcal{X}_0)}\}$ into $d^{(\mathcal{X})}$ disjoint non-empty sets, denoted by $X_c^{(\mathcal{M})}$, such that for each $c \in \{1, ..., d^{(\mathcal{X})}\}$ all $b_v$ indexed by $v \in X_c^{(\mathcal{M})}$ have identical values. We refer to $X_c^{(\mathcal{M})}$ as *concise index sets*. The set of all concise index sets is uniquely determined by $\mathcal{M}$ and vice versa.[4]

We define below an estimator of the moment ratios $\mathbf{b}$, denoted by $\hat{\mathbf{b}}$, and an estimator of the covariance matrix of $\hat{\mathbf{b}}$, denoted by $\hat{\mathbf{\Sigma}}$. Furthermore, we assume all sample moments used for the test are linearly independent, which implies that the covariance matrix of $\hat{\mathbf{b}}$, denoted by $\mathbf{\Sigma}$, is invertible. Then, the statistic

$$\mathcal{T}_{\mathcal{M}}(\hat{\mathbf{b}}, \hat{\mathbf{\Sigma}}) = \frac{1}{2} \sum_{k=1}^{d^{(\mathcal{X}_0)}} \frac{\left(\hat{b}_k\right)^2}{\hat{\Sigma}_{kk}} - \frac{1}{2} \sum_{c=1}^{d^{(\mathcal{X})}} \frac{\left(\sum_{l_1 \in X_c^{(\mathcal{M})}} \hat{b}_{l_1} \left(\hat{\Sigma}_{l_1 l_1}\right)^{-1}\right)^2}{\sum_{l_2 \in X_c^{(\mathcal{M})}} \left(\hat{\Sigma}_{l_2 l_2}\right)^{-1}} \tag{1}$$

tests asymptotically for both null hypotheses if the covariance matrix $\mathbf{\Sigma}$ is diagonal and the components of $\mathbf{b}$ are defined (all moment ratios have non-zero denominators).

In order to extend the test to arbitrary $\mathbf{\Sigma}$ and potentially zero denominators of $\mathbf{b}$, we introduce a scaling factor $\lambda^{(\max)}$ that corrects for correlated components of $\hat{\mathbf{b}}$ by rescaling the covariance matrix $\hat{\mathbf{\Sigma}}$. In addition, we constrain the nominal significance level, denoted by $\alpha^{(*)}$, to be larger than a minimal nominal significance level $\alpha^{(\min)}$.

More precisely, let $d^{(\mathcal{H})}$ denote the total number of modularizations tested on the same samples $\hat{\mathbf{s}}$ and let $\mathcal{M}_i$ denote the $i$-th modularization. Under both null hypotheses, the probability of sampling $\mathcal{T}_{\mathcal{M}_i}(\hat{\mathbf{b}}, \lambda^{(\max)}\hat{\mathbf{\Sigma}})$ for $i \in \{1, ..., d^{(\mathcal{H})}\}$ at least as extreme as observed is less than $\alpha^{(*)}$ if

$$p_i^{(\Gamma)}\text{-value} \leq \frac{\alpha^{(\Gamma)}}{d^{(\mathcal{H})}} \tag{2}$$

$$\alpha^{(\Gamma)} = \frac{\alpha^{(*)} - \alpha^{(\min)}}{1 - \alpha^{(\min)}}, \tag{3}$$

[3]The statistical test applies even if some or all of these ratios have zero denominator and are undefined.
[4]Up to irrelevant singletons.

where $p_i^{(\Gamma)}$-value denotes the probability of sampling $\mathcal{T}_{\mathcal{M}_i}(\hat{\mathbf{b}}, \lambda^{(\max)}\hat{\mathbf{\Sigma}})$ at least as extreme as observed when distributed according to the gamma distribution $\Gamma(\zeta_{\mathcal{M}_i}, 1)$ for shape parameter $\zeta_{\mathcal{M}} = \frac{1}{2}\left(d^{(X_0)} - d^{(X)}\right)$ and scale parameter 1. For $\zeta_{\mathcal{M}} = 0$, $\mathcal{T}_{\mathcal{M}} = 0$.

Furthermore, we consolidate numerically that this statement holds not only asymptotically but also for sufficiently many samples such that the moment estimates are approximately joint normal and $N^{(t)}\hat{\mathbf{\Sigma}} \approx N^{(t)}\mathbf{\Sigma}$. For the investigated datasets, a few hundred samples turn out to be sufficient to fulfill these requirements.

Finally, if all moment rations $\mathbf{b}$ are finite, the test is asymptotically consistent for H2, i.e., its power for any false hypothesis converges asymptotically to one. If in addition all components of $\hat{\mathbf{b}}$ are uncorrelated, the test statistic $\mathcal{T}_{\mathcal{M}_i}(\hat{\mathbf{b}}, \hat{\mathbf{\Sigma}})$ is asymptotically minimal sufficient, i.e., it most efficiently captures all information about a modularization $\mathcal{M}_i$ contained in the sample moment ratios $\hat{\mathbf{b}}$.

In the remainder of this section, we define the concise index sets $X_c^{(\mathcal{M})}$ (see Equ. 7), the estimators $\hat{\mathbf{b}}$ and $\hat{\mathbf{\Sigma}}$ (see Equ. 9 and 15), the scaling factor $\lambda^{(\max)}$ (see Equ. 152), and the minimal nominal significance level $\alpha^{(\min)}$ (see Equ. 20).

##### Construction of concise index sets

Concise index sets can be constructed by first extending a modularization $\mathcal{M}$ by including all (missing) singletons $S_k = \{k\}, k \in \{1, ..., d-1\}$, subsequently, constructing *redundant index sets*, one for each unordered pair of functional modules, and, finally, removing some of the redundant index sets to obtain concise index sets.

For overlapping pairs of functional modules, the corresponding redundant index sets are empty. For disjoint pairs of functional modules $S_{c_1}$ and $S_{c_2}$, the corresponding redundant index set, denoted by $Y_u$, consists of the vectorization $v(k, l)$ of all elements $(k, l)$ of the Cartesian product $S_{c_1} \times S_{c_2}$, where $k$ and $l$ are exchanged if $k > l$. More precisely,[5]

$$\begin{aligned} Y_{u(c_1,c_2)} &= \{v(k,l) | k, l \in \{1, ..., d-1\} \text{ and } (k < l) \text{ and } (S_{c_1} \cap S_{c_2} = \{\}) \text{ and} \\ &\quad [((s_k \in S_{c_1}) \text{ and } (s_l \in S_{c_2})) \text{ or } ((s_l \in S_{c_1}) \text{ and } (s_k \in S_{c_2}))]\} \end{aligned} \tag{4}$$

for $c_1 \in \{1, ..., c_2 - 1\}, c_2 \in \{2, ..., d^{(\mathcal{M})}\}$ and vectorizations (see Fig. S1D)

$$v(k,l) = \frac{(d-3)(d-2)}{2} - \frac{(d-2-k)(d-1-k)}{2} + l - 1 \tag{5}$$

$$u(c_1, c_2) = \frac{(d^{(\mathcal{M})}-2)(d^{(\mathcal{M})}-1)}{2} - \frac{(d^{(\mathcal{M})}-1-c_1)(d^{(\mathcal{M})}-c_1)}{2} + c_2 - 1, \tag{6}$$

where $k \in \{1, ..., l-1\}$, $l \in \{2, ..., d-1\}$, $u \in \{1, ..., d^{(u)}\}$ and $d^{(u)} = \frac{d^{(\mathcal{M})}(d^{(\mathcal{M})}-1)}{2}$.[6]

Finally, only the redundant sets that are not subsets of other redundant sets are selected as concise index sets, i.e., the elements of

$$X^{(\mathcal{M})} = \left\{Y_{c_1} \middle| c_1 \in \left\{1, ..., d^{(u)}\right\} \text{ and } Y_{c_1} \not\subseteq Y_{c_2} \text{ for all } c_2 \neq c_1, c_2 \in \left\{1, ..., d^{(u)}\right\}\right\} \tag{7}$$

$$d^{(X)} = |X^{(\mathcal{M})}| \tag{8}$$

sorted by their smallest element define the concise index sets $X_c^{(\mathcal{M})}$ for $c \in \{1, ..., d^{(X)}\}$. Detailed examples for the construction of $Y_u$ and $X_c^{(\mathcal{M})}$ are shown in Table 1.

[5]Using set-builder notation, where $\{x | x \in X \text{ and } \Theta(x)\}$ denotes the set of all $x \in X$ such that $x$ satisfies the condition $\Theta(x)$, e.g., $\{x | x \in \mathbb{R} \text{ and } (x \geq 0)\}$ defines the set of all non-negative real numbers.

[6]$u$ and $v$ are similar (see Fig. S1D). I.e., if $d^{(\mathcal{M})} = d-1$, $u(k,l) = v(k,l)$ for $k \in \{1, ..., l-1\}$, $l \in \{2, ..., d-1\}$.

**A:** Construction of $\mathcal{X}^{(\mathcal{M})}$ for the modularization $\mathcal{M} = \{S_1\}$ consisting of a single functional module $S_1 = \{1,2\}$ (see Fig. S1A). For the construction, the modularization is extended to $\mathcal{M} = \{S_1, S_2, S_3, S_4, S_5\}$, where $S_1 = \{1\}$, $S_2 = \{2\}$, $S_3 = \{3\}$, $S_4 = \{4\}$ and $S_5 = \{1,2\}$. $\mathcal{X}^{(\mathcal{M})} = \{X_1^{(\mathcal{M})}, X_2^{(\mathcal{M})}, X_3^{(\mathcal{M})}, X_4^{(\mathcal{M})}\}$.

| $(c_1, c_2)$ | $S_{c_1}$ | $S_{c_2}$ | $Y_{u(c_1,c_2)} =$ | $Y_u$ | | | |
|---|---|---|---|---|---|---|---|
| (1,2) | {1} | {2} | $Y_{u(1,2)} =$ | $Y_1 =$ | $\{v(1,2)\} =$ | {1} | $= X_1^{(\mathcal{M})}$ |
| (1,3) | {1} | {3} | $Y_{u(1,3)} =$ | $Y_2 =$ | $\{v(1,3)\} =$ | {2} | |
| (1,4) | {1} | {4} | $Y_{u(1,4)} =$ | $Y_3 =$ | $\{v(1,4)\} =$ | {3} | |
| (1,5) | {1} | {1,2} | $Y_{u(1,5)} =$ | $Y_4 =$ | $\{\} =$ | {} | |
| (2,3) | {2} | {3} | $Y_{u(2,3)} =$ | $Y_5 =$ | $\{v(2,3)\} =$ | {4} | |
| (2,4) | {2} | {4} | $Y_{u(2,4)} =$ | $Y_6 =$ | $\{v(2,4)\} =$ | {5} | |
| (2,5) | {2} | {1,2} | $Y_{u(2,5)} =$ | $Y_7 =$ | $\{\} =$ | {} | |
| (3,4) | {3} | {4} | $Y_{u(3,4)} =$ | $Y_8 =$ | $\{v(3,4)\} =$ | {6} | $= X_4^{(\mathcal{M})}$ |
| (3,5) | {3} | {1,2} | $Y_{u(3,5)} =$ | $Y_9 =$ | $\{v(1,3), v(2,3)\} =$ | {2,4} | $= X_2^{(\mathcal{M})}$ |
| (4,5) | {4} | {1,2} | $Y_{u(4,5)} =$ | $Y_{10} =$ | $\{v(1,4), v(2,4)\} =$ | {3,5} | $= X_3^{(\mathcal{M})}$ |

**B:** Construction of $\mathcal{X}^{(\mathcal{M})}$ for the flat modularization $\mathcal{M} = \{S_1, S_2\}$ consisting of $S_1 = \{1,2\}$ and $S_2 = \{3,4\}$ (see Fig. S1B). For the construction, the modularization is extended to $\mathcal{M} = \{S_1, S_2, S_3, S_4, S_5, S_6\}$, where $S_1 = \{1\}$, $S_2 = \{2\}$, $S_3 = \{3\}$, $S_4 = \{4\}$, $S_5 = \{1,2\}$ and $S_6 = \{3,4\}$. $\mathcal{X}^{(\mathcal{M})} = \{X_1^{(\mathcal{M})}, X_2^{(\mathcal{M})}, X_3^{(\mathcal{M})}\}$.

| $(c_1, c_2)$ | $S_{c_1}$ | $S_{c_2}$ | $Y_{u(c_1,c_2)} =$ | $Y_u$ | | | |
|---|---|---|---|---|---|---|---|
| (1,2) | {1} | {2} | $Y_{u(1,2)} =$ | $Y_1 =$ | $\{v(1,2)\} =$ | {1} | $= X_1^{(\mathcal{M})}$ |
| (1,3) | {1} | {3} | $Y_{u(1,3)} =$ | $Y_2 =$ | $\{v(1,3)\} =$ | {2} | |
| (1,4) | {1} | {4} | $Y_{u(1,4)} =$ | $Y_3 =$ | $\{v(1,4)\} =$ | {3} | |
| (1,5) | {1} | {1,2} | $Y_{u(1,5)} =$ | $Y_4 =$ | $\{\} =$ | {} | |
| (1,6) | {1} | {3,4} | $Y_{u(1,6)} =$ | $Y_5 =$ | $\{v(1,3), v(1,4)\} =$ | {2,3} | |
| (2,3) | {2} | {3} | $Y_{u(2,3)} =$ | $Y_6 =$ | $\{v(2,3)\} =$ | {4} | |
| (2,4) | {2} | {4} | $Y_{u(2,4)} =$ | $Y_7 =$ | $\{v(2,4)\} =$ | {5} | |
| (2,5) | {2} | {1,2} | $Y_{u(2,5)} =$ | $Y_8 =$ | $\{\} =$ | {} | |
| (2,6) | {2} | {3,4} | $Y_{u(2,6)} =$ | $Y_9 =$ | $\{v(2,3), v(2,4)\} =$ | {4,5} | |
| (3,4) | {3} | {4} | $Y_{u(3,4)} =$ | $Y_{10} =$ | $\{v(3,4)\} =$ | {6} | $= X_3^{(\mathcal{M})}$ |
| (3,5) | {3} | {1,2} | $Y_{u(3,5)} =$ | $Y_{11} =$ | $\{v(1,3), v(2,3)\} =$ | {2,4} | |
| (3,6) | {3} | {3,4} | $Y_{u(3,6)} =$ | $Y_{12} =$ | $\{\} =$ | {} | |
| (4,5) | {4} | {1,2} | $Y_{u(4,5)} =$ | $Y_{13} =$ | $\{v(1,4), v(2,4)\} =$ | {3,5} | |
| (4,6) | {4} | {3,4} | $Y_{u(4,6)} =$ | $Y_{14} =$ | $\{\} =$ | {} | |
| (5,6) | {1,2} | {3,4} | $Y_{u(5,6)} =$ | $Y_{15} =$ | $\{v(1,3), v(2,3), v(1,4), v(2,4)\} =$ | {2,3,4,5} | $= X_2^{(\mathcal{M})}$ |

**C:** Construction of $\mathcal{X}^{(\mathcal{M})}$ for the nested modularization $\mathcal{M} = \{S_1, S_2\}$ consisting of $S_1 = \{3,4\}$ and $S_2 = \{2,3,4\}$ (see Fig. S1C). For the construction, the modularization is extended to $\mathcal{M} = \{S_1, S_2, S_3, S_4, S_5, S_6\}$, where $S_1 = \{1\}$, $S_2 = \{2\}$, $S_3 = \{3\}$, $S_4 = \{4\}$, $S_5 = \{3,4\}$ and $S_6 = \{2,3,4\}$. $\mathcal{X}^{(\mathcal{M})} = \{X_1^{(\mathcal{M})}, X_2^{(\mathcal{M})}, X_3^{(\mathcal{M})}\}$.

| $(c_1, c_2)$ | $S_{c_1}$ | $S_{c_2}$ | $Y_{u(c_1,c_2)} =$ | $Y_u$ | | | |
|---|---|---|---|---|---|---|---|
| (1,2) | {1} | {2} | $Y_{u(1,2)} =$ | $Y_1 =$ | $\{v(1,2)\} =$ | {1} | |
| (1,3) | {1} | {3} | $Y_{u(1,3)} =$ | $Y_2 =$ | $\{v(1,3)\} =$ | {2} | |
| (1,4) | {1} | {4} | $Y_{u(1,4)} =$ | $Y_3 =$ | $\{v(1,4)\} =$ | {3} | |
| (1,5) | {1} | {3,4} | $Y_{u(1,5)} =$ | $Y_4 =$ | $\{v(1,3), v(1,4)\} =$ | {2,3} | |
| (1,6) | {1} | {2,3,4} | $Y_{u(1,6)} =$ | $Y_5 =$ | $\{v(1,2), v(1,3), v(1,4)\} =$ | {1,2,3} | $= X_1^{(\mathcal{M})}$ |
| (2,3) | {2} | {3} | $Y_{u(2,3)} =$ | $Y_6 =$ | $\{v(2,3)\} =$ | {4} | |
| (2,4) | {2} | {4} | $Y_{u(2,4)} =$ | $Y_7 =$ | $\{v(2,4)\} =$ | {5} | |
| (2,5) | {2} | {3,4} | $Y_{u(2,5)} =$ | $Y_8 =$ | $\{v(2,3), v(2,4)\} =$ | {4,5} | $= X_2^{(\mathcal{M})}$ |
| (2,6) | {2} | {2,3,4} | $Y_{u(2,6)} =$ | $Y_9 =$ | $\{\} =$ | {} | |
| (3,4) | {3} | {4} | $Y_{u(3,4)} =$ | $Y_{10} =$ | $\{v(3,4)\} =$ | {6} | $= X_3^{(\mathcal{M})}$ |
| (3,5) | {3} | {3,4} | $Y_{u(3,5)} =$ | $Y_{11} =$ | $\{\} =$ | {} | |
| (3,6) | {3} | {2,3,4} | $Y_{u(3,6)} =$ | $Y_{12} =$ | $\{\} =$ | {} | |
| (4,5) | {4} | {3,4} | $Y_{u(4,5)} =$ | $Y_{13} =$ | $\{\} =$ | {} | |
| (4,6) | {4} | {2,3,4} | $Y_{u(4,6)} =$ | $Y_{14} =$ | $\{\} =$ | {} | |
| (5,6) | {3,4} | {2,3,4} | $Y_{u(5,6)} =$ | $Y_{15} =$ | $\{\} =$ | {} | |

Table 1: Construction of concise index sets.

##### Simplest possible moments

For each sample moment ratio $\hat{b}_v$, $v \in \{1, ..., d^{(\mathcal{X}_0)}\}$, $d^{(\mathcal{X}_0)} = \frac{(d-2)(d-1)}{2}$ we define a vector $\hat{\mathbf{b}}^{(v)} \in \mathbb{R}^2$ containing the numerator and the denominator of $\hat{b}_v$. We estimate the moment ratio $b_v$ by

$$\hat{b}_v = \begin{cases} 0, & \left|\varrho^{(v)}\right| < \theta_{\hat{\mu}} \\ \frac{\hat{b}_1^{(v)}}{\hat{b}_2^{(v)}}, & \text{otherwise} \end{cases} \tag{9}$$

$$\varrho^{(v)} = \sqrt{\frac{N^{(\mathrm{t})}}{2\,\hat{\Sigma}'^{(v,v)}_{22}}}\hat{b}_2^{(v)}, \tag{10}$$

where a cutoff $\theta_{\hat{\mu}} > 0$ ensures finite expectations of $\hat{b}_v$ for joint normal $\hat{\mathbf{b}}^{(v)}$.

The simplest possible moments for testing nested modularizations are $b_1^{(v)} = E[s_k s_l s_d]$ and $b_2^{(v)} = E[s_k s_d]E[s_l s_d]$ for $k \neq l$, which are estimated by the sample moments

$$\hat{b}_i^{(v)} = \frac{2}{N^{(\mathrm{t})}} \sum_{n=1}^{N^{(\mathrm{t})}/2} \hat{\Phi}_{ni}^{(v)} \tag{11}$$

$$\hat{\Phi}_{n1}^{(v(k,l))} = \hat{s}_k^{(2n-1)} \hat{s}_l^{(2n-1)} \hat{s}_d^{(2n-1)} \tag{12}$$

$$\hat{\Phi}_{n2}^{(v(k,d))} = \hat{s}_k^{(n)} \hat{s}_d^{(n)} \hat{s}_l^{(n+N^{(\mathrm{t})}/2)} \hat{s}_d^{(n+N^{(\mathrm{t})}/2)} \tag{13}$$

for $i \in \{1, 2\}$, $k \in \{1, ..., l-1\}$, $l \in \{2, ..., d-1\}$. The corresponding sample covariance matrices are

$$\hat{\Sigma}'^{(v,\omega)}_{ij} = \frac{1}{N^{(\mathrm{t})}/2 - 1} \sum_{n=1}^{N^{(\mathrm{t})}/2} \left(\hat{\Phi}_{ni}^{(v)} - \hat{b}_i^{(v)}\right)\left(\hat{\Phi}_{nj}^{(\omega)} - \hat{b}_j^{(\omega)}\right) \tag{14}$$

for $i, j \in \{1, 2\}$ and $v, \omega \in \{1, ..., d^{(\mathcal{X}_0)}\}$, where $\omega(k, l) = v(k, l)$.

The covariance matrix of the sample moment ratios $\hat{\mathbf{b}}$ can be approximated by

$$\hat{\Sigma}_{v\omega} = \begin{cases} \infty, & (v = \omega) \text{ and } \left(\left|\varrho^{(v)}\right| < \theta_{\hat{\mu}}\right) \\ 0, & (v \neq \omega) \text{ and } \left(\left(\left|\varrho^{(v)}\right| < \theta_{\hat{\mu}}\right) \text{ or } \left(\left|\varrho^{(\omega)}\right| < \theta_{\hat{\mu}}\right)\right) \\ \frac{2\sqrt{\lambda_v \lambda_\omega}}{N^{(\mathrm{t})}\hat{b}_2^{(v)}\hat{b}_2^{(\omega)}} \left(\hat{\Sigma}'^{(v,\omega)}_{11} - \frac{\hat{b}_1^{(\omega)}}{\hat{b}_2^{(\omega)}}\hat{\Sigma}'^{(v,\omega)}_{12} - \right. & \\ \quad \left. \frac{\hat{b}_1^{(v)}}{\hat{b}_2^{(v)}}\hat{\Sigma}'^{(v,\omega)}_{21} + \frac{\hat{b}_1^{(v)}\hat{b}_1^{(\omega)}}{\hat{b}_2^{(v)}\hat{b}_2^{(\omega)}}\hat{\Sigma}'^{(v,\omega)}_{22}\right), & \text{otherwise} \end{cases} \tag{15}$$

for $v, \omega \in \{1, ..., d^{(\mathcal{X}_0)}\}$.[7] All results hold asymptotically for $\lambda_v = 1$. However, we consolidate numerically that the test applies to sufficiently many samples such that the moment estimates are approximately joint normal, $\hat{\mathbf{\Sigma}} \approx \mathbf{\Sigma}$ and

$$\lambda_v = \xi_0 \left(1 + \sum_{n=1}^{5} \xi_n \left(\frac{\varrho^{(v)}}{6}\right)^{-2n}\right), \tag{16}$$

where the parameters $\xi_n$, $n \in \{0, ..., 5\}$ are chosen in dependence on the cutoff $\theta_{\hat{\mu}} \in \{5, 6, 7\}$ according to Table 2.

Finally, the simplest possible moments for testing single functional modules are $b_1^{(v)} = E[s_k s_l]$ and $b_2^{(v)} = E[s_k s_d]$ for $k \neq l$, which are estimated by

$$\hat{\Phi}_{n1}^{(v(k,l))} = \hat{s}_k^{(n)} \hat{s}_l^{(n)} \tag{17}$$

$$\hat{\Phi}_{n2}^{(v(k,l))} = \hat{s}_k^{(n)} \hat{s}_d^{(n)}. \tag{18}$$

[7] $\hat{\Sigma}_{v\omega} = \infty$ denotes $\lim_{\hat{\Sigma}_{v\omega} \to \infty} \mathcal{T}_{\mathcal{M}}$.

| $\theta_{\hat{\mu}}$ | $\xi_0$ | $\xi_1$ | $\xi_2$ | $\xi_3$ | $\xi_4$ | $\xi_5$ |
|---|---|---|---|---|---|---|
| 5 | 1.367 | 2.047 | 4.735 | -1.923 | -1.231 | 2.790 |
| 6 | 1.601 | 1.176 | 18.912 | -19.224 | 0.022 | 26.680 |
| 7 | 1.916 | 0.446 | 39.582 | -65.234 | 42.532 | 120.680 |

Table 2: Parameters $\xi_n$ for different values of $\theta_{\hat{\mu}}$.

**Correction for correlated moment ratios and minimal nominal significance level**

For potentially correlated components of the sample moment ratios $\hat{\mathbf{b}}$, we define a multiplicative factor $\lambda^{(\max)} \geq 1$ that rescales $\hat{\boldsymbol{\Sigma}}$ such that the statistical test remains conservative. The smallest possible $\lambda^{(\max)}$ with this property is the largest eigenvalue of a submatrix of $\hat{\boldsymbol{\Sigma}}$ normalized to unit diagonal, i.e.,

$$\grave{\boldsymbol{\Sigma}}\hat{\boldsymbol{\Sigma}}\grave{\boldsymbol{\Sigma}}, \tag{19}$$

where the matrix $\grave{\boldsymbol{\Sigma}}$ has the same size as $\hat{\boldsymbol{\Sigma}}$, is diagonal and $\grave{\Sigma}_{vv} = \hat{\Sigma}_{vv}^{-1/2}$ if $v$ is element of an index set of size larger than one and $\grave{\Sigma}_{vv} = 0$, otherwise. A more power full test is derived in Section 2.4.

Finally, we define the minimal nominal significance level

$$\alpha^{(\min)} = 1 - \operatorname{erf}\left(\frac{\theta_{\hat{\mu}} - 1}{\sqrt{2}}\right)^{d^{(X_0)}} \operatorname{erf}\left(\frac{6}{\sqrt{2}}\right)^{d^{(X_0)}}, \tag{20}$$

where erf denotes the error function. Table 3 shows $\alpha^{(\min)}$ in dependence on the cutoff $\theta_{\hat{\mu}}$ and $d^{(X_0)}$.

| $\theta_{\hat{\mu}}$ | $d^{(X_0)}(d)$ : | 6 (5) | 10 (6) | 45 (11) | 105 (16) | 153 (19) | 190 (21) |
|---|---|---|---|---|---|---|---|
| 5 | | $3.8 \cdot 10^{-4}$ | $6.3 \cdot 10^{-4}$ | $28.5 \cdot 10^{-4}$ | $66.3 \cdot 10^{-4}$ | $96.5 \cdot 10^{-4}$ | $119.6 \cdot 10^{-4}$ |
| 6 | | $3.5 \cdot 10^{-6}$ | $5.8 \cdot 10^{-6}$ | $25.9 \cdot 10^{-6}$ | $60.4 \cdot 10^{-6}$ | $88.0 \cdot 10^{-6}$ | $109.3 \cdot 10^{-6}$ |
| 7 | | $2.4 \cdot 10^{-8}$ | $3.9 \cdot 10^{-8}$ | $17.8 \cdot 10^{-8}$ | $41.4 \cdot 10^{-8}$ | $60.4 \cdot 10^{-8}$ | $75.0 \cdot 10^{-8}$ |

Table 3: $\alpha^{(\min)}$ for different values of $\theta_{\hat{\mu}}$ and $d^{(X_0)}$.

### 2 Results

In the next section, we show that the existence of single functional modules with binary interface variables can be uniquely determined by the moments of the observable states. In Section 2.2 we extend this result to multiple functional modules and derive necessary conditions for nested modularizations and sufficient and necessary conditions for models of flat modularizations. In Section 2.3 we define a test statistic for nested and flat modularizations given joint normal and uncorrelated estimates of moment ratios. In Section 2.4 we drop this assumption and derive an asymptotically conservative test that requires only an invertible covariance matrix of sample moments (Equ. 153). We conclude by demonstrating numerically that the statistical tests can be applied for finitely many samples if the moment estimates are approximately joint normal and the corresponding sample covariance matrix is close to its asymptotic value.

#### 2.1 Identification of single functional modules

##### Moments

For singe functional modules, we divide $\mathbf{s}$ into $\mathbf{x} \in \mathbb{R}^{d^{(x)}}$ and $\mathbf{z} \in \mathbb{R}^{d^{(z)}}$ such that $\mathbf{x}$ consists of the components of $\mathbf{s}$ indexed by $S$, $d^{(x)} + d^{(z)} = d$ and $0 < d^{(x)} < d$. We refer to $\mathbf{x}$ as *observable states within a functional module* and $\mathbf{z}$ as *observable states outside of a functional module*. The conditional expectation of $\mathbf{x}$ relative to the $\sigma$-algebra generated by $\mathbf{z}$ is denoted by $E[\mathbf{x}|\mathbf{z}]$.

Furthermore, we denote by $\mathbb{N}_0^d$ the set of all multi-indices $\mathbf{q} = (q_1, ..., q_d)$, $q_i \in \mathbb{N}_0$, $i \in \{1, ..., d\}$. For multiple multi-indices $\mathbf{q}^{(n)} \in \mathbb{N}_0^{d^{(x)}}$, $n \in \mathbb{N}$, we denote the monomials $\prod_{i=1}^{d^{(x)}} x_i^{q_i^{(n)}}$ by $P_n(\mathbf{x})$.[8] Likewise, for multiple multi-indices $\mathbf{r}^{(m)} \in \mathbb{N}_0^{d^{(z)}}$, $m \in \mathbb{N}$, we denote the monomials $\prod_{i=1}^{d^{(z)}} z_i^{r_i^{(m)}}$ by $Q_m(\mathbf{z})$. All moments $E[P_n(\mathbf{x})Q_m(\mathbf{z})]$ exist and uniquely determine the distribution of $\mathbf{s}$ as all observable states are bounded (Kleiber and Stoyanov, 2013).

##### Sufficient and necessary conditions for functional modules

Whether or not functional modules with binary interface variables exist can be uniquely determined by the moments of the distribution of $\mathbf{x}$ and $\mathbf{z}$. Assume a matrix of mixed moments $M_{ij} = E[P_i(\mathbf{x})Q_j(\mathbf{z})]$, $i, j \in \{1, 2\}$ with arbitrary monomials $P_i(\mathbf{x})$ and $Q_j(\mathbf{z})$ has full rank. Then, according to Theorem 6, there exists a functional module with a binary interface variable $y$ such that $\mathbf{x}$ is independent of $\mathbf{z}$ given $y$ if and only if

$$E[P_n(\mathbf{x})Q_m(\mathbf{z})] = \sum_{j=1}^{2} C_{mj} E[P_n(\mathbf{x})Q_j(\mathbf{z})] \tag{21}$$

for all $n, m \in \mathbb{N}$ and $C_{mj} \in \mathbb{R}$, where $P_n(\mathbf{x})$ and $Q_m(\mathbf{z})$ denote all monomials in $\mathbf{x}$ and $\mathbf{z}$.

These conditions can be tested empirically for finite sets of moments to potentially

[8]The symbol P denotes probabilities, whereas $P$ denotes monomials and $p$ denotes probability densities.

falsify functional modules. For single functional modules we choose

$$P_n(\mathbf{x}) \quad = \quad \begin{cases} 1, & n = 1 \\ x_{n-1}, & \text{otherwise} \end{cases} \tag{22}$$

$$Q_m(\mathbf{z}) \quad = \quad \begin{cases} 1, & m = 1 \\ z_{m-1}, & \text{otherwise} \end{cases} \tag{23}$$

and for modularizations we choose

$$P_n(\mathbf{x}) \quad = \quad \begin{cases} 1, & n = 1 \\ x_{n-1}, & \text{otherwise} \end{cases} \tag{24}$$

$$Q_m(\mathbf{z}) \quad = \quad \begin{cases} 1, & m = 1 \\ z_1, & m = 2 \\ z_1 z_{m-1}, & \text{otherwise,} \end{cases} \tag{25}$$

where $n \in \{1, ..., d^{(x)} + 1\}$ and $m \in \{1, ..., d^{(z)} + 1\}$.

Analogous conditions can be derived for discrete interface variables $y$ with more than two values.[9] Again, these can be tested empirically for finite sets of moments to potentially falsify functional modules. However, for interface variables with more than two values the existence of functional modules can not be guaranteed even if all conditions hold.

##### Sufficient and necessary conditions for models of functional modules

In this section, we assume all observable components are correlated with $z_1$, i.e., $E[x_k z_1] \neq 0$ and $E[z_l z_1] \neq 0$ for all $k \in \{1, ..., d^{(x)}\}$ and all $l \in \{2, ..., d^{(z)}\}$. Furthermore, without loss of generality, we normalize the data to zero mean, i.e., $E[x_k] = 0$ and $E[z_l] = 0$ for all $k \in \{1, ..., d^{(x)}\}$ and all $l \in \{1, ..., d^{(z)}\}$. This normalization preserves functional modules, i.e., $\mathbf{x}$ are observable states within a functional module after the normalization if and only if they are observable states within a functional module before. Then, the conditions for functional modules in Equ. 21 can not only be used to falsify a functional module, but also to falsify any model of a functional module that matches a finite set of moments.

More precisely, for the choice $P_1(\mathbf{x}) = Q_1(\mathbf{z}) = 1$ the matrix $\mathbf{M}$ is diagonal and has full rank. Then, the conditions in Equ. 21 hold for all $P_n(\mathbf{x})$ and all $Q_m(\mathbf{z})$ as defined for single functional modules (Equ. 22 and 23) if and only if the observable states $\mathbf{x}$ and $\mathbf{z}$ can be modeled by random variables $\tilde{\mathbf{x}} \in \mathbb{R}^{d^{(x)}}$, $\tilde{\mathbf{z}} \in \mathbb{R}^{d^{(z)}}$ and $\tilde{y} \in \{1, 2\}$ such that (i) $\tilde{\mathbf{x}}$ is independent of $\tilde{\mathbf{z}}$ given $\tilde{y}$ and (ii)

$$E[P_n(\mathbf{x}) Q_m(\mathbf{z})] \quad = \quad E[P_n(\tilde{\mathbf{x}}) Q_m(\tilde{\mathbf{z}})] \tag{26}$$

for all $n \in \{1, ..., d^{(x)} + 1\}$ and all $m \in \{1, ..., d^{(z)} + 1\}$ (Lemma 7).

We simplify these conditions by defining

$$B^{(s)}_{kl} = \frac{E[x_k z_l]}{E[x_k z_1]}, \quad k \in \{1, ..., d^{(x)}\}, \quad l \in \{1, ..., d^{(z)}\}, \tag{27}$$

which always exist because $E[x_k z_1] \neq 0$ for all $k \in \{1, ..., d^{(x)}\}$. Then, the conditions in Equ. 21 hold for all $n \in \{1, ..., d^{(x)} + 1\}$ and all $m \in \{1, ..., d^{(z)} + 1\}$ if and only if

$$B^{(s)}_{kl} = b^{(s)}_l \tag{28}$$

[9] See Lemma 1.

for all $k \in \{1, ..., d^{(\mathrm{x})}\}$, all $l \in \{1, ..., d^{(\mathrm{z})}\}$ and $\mathbf{b}^{(\mathrm{s})} \in \mathbb{R}^{d^{(\mathrm{z})}}$.[10] For non-zero correlations between $z_1$ and all other observable components, these are sufficient and necessary conditions that a model of a functional module with a binary interface variable and the same correlations as implied by Equ. 22 and 23 exists.

Likewise, given $\mathbf{M}$ is diagonal and has full rank the conditions in Equ. 21 hold for all $P_n(\mathbf{x})$ and all $Q_m(\mathbf{z})$ as defined for modularizations (Equ. 24 and 25) if and only if the observable states $\mathbf{x}$ and $\mathbf{z}$ can be modeled by random variables $\tilde{\mathbf{x}} \in \mathbb{R}^{d^{(\mathrm{x})}}$, $\tilde{\mathbf{z}} \in \mathbb{R}^{d^{(\mathrm{z})}}$ and $\tilde{y} \in \{1, 2\}$ such that (i) $\tilde{\mathbf{x}}$ is independent of $\tilde{\mathbf{z}}$ given $\tilde{y}$ and (ii)

$$E[P_n(\mathbf{x})Q_m(\mathbf{z})] \quad = \quad E[P_n(\tilde{\mathbf{x}})Q_m(\tilde{\mathbf{z}})] \tag{29}$$

for all $n \in \{1, ..., d^{(\mathrm{x})} + 1\}$ and all $m \in \{1, ..., d^{(\mathrm{z})} + 1\}$ (Lemma 8).

We simplify these conditions by defining

$$B^{(\mathrm{m})}_{kl} = \frac{E[x_k z_1 z_l]}{E[x_k z_1]E[z_1 z_l]}, \quad k \in \{1, ..., d^{(\mathrm{x})}\}, \quad l \in \{2, ..., d^{(\mathrm{z})}\}, \tag{30}$$

which always exist because $E[x_k z_1] \neq 0$ for all $k \in \{1, ..., d^{(\mathrm{x})}\}$ and $E[z_l z_1] \neq 0$ for all $l \in \{2, ..., d^{(\mathrm{z})}\}$. Then, the conditions in Equ. 21 hold for all $n \in \{1, ..., d^{(\mathrm{x})} + 1\}$ and all $m \in \{1, ..., d^{(\mathrm{z})} + 1\}$ if and only if

$$B^{(\mathrm{m})}_{kl} = b^{(\mathrm{m})}_l \tag{31}$$

for all $k \in \{1, ..., d^{(\mathrm{x})}\}$, all $l \in \{2, ..., d^{(\mathrm{z})}\}$ and $b^{(\mathrm{m})}_l \in \mathbb{R}$.[11] Again, for non-zero correlations between $z_1$ and all other observable components, these are sufficient and necessary conditions that a model of a functional module with a binary interface variable and the same moments as implied by Equ. 24 and 25 exists.

As a side note, analogous statements can be derived for less constrained monomials (see Lemma 9).

#### 2.2 Identification of nested modularizations

In this section we derive necessary and sufficient conditions for the existence of nested modularizations and necessary and sufficient conditions for the existence of models of flat modularizations.

Let $\mathbf{x}^{(c)}$, $\mathbf{z}^{(c)}$ and $y^{(c)}$ for $c \in \{1, ..., d^{(\mathcal{M})}\}$ denote all observable components indexed by $S_c$, all other observable components and the interface variable of the $c$-th functional module, respectively. Let $z^{(c)}_1$ denote $s_d$ for all $c \in \{1, ..., d^{(\mathcal{M})}\}$. Furthermore, let $d^{(\mathrm{x},c)} \in \mathbb{N}$ and $d^{(\mathrm{z},c)} \in \mathbb{N}$ denote the total number of elements of $\mathbf{x}^{(c)}$ and $\mathbf{z}^{(c)}$, respectively. According to Theorem 6, there exist binary interface variables $y^{(c)}$ such that $\mathbf{x}^{(c)}$ is independent of $\mathbf{z}^{(c)}$ given $y^{(c)}$ if and only if

$$E[P_n(\mathbf{x}^{(c)})Q_m(\mathbf{z}^{(c)})] = \sum_{j=1}^{2} C^{(c)}_{mj} E[P_n(\mathbf{x}^{(c)})Q_j(\mathbf{z}^{(c)})] \tag{32}$$

[10] According to Lemma 2. For $M_{11} = 1$ and $M_{22} = E[x_1 z_1]$ (see Equ. 177) we obtain $E[x_k z_l] = \frac{E[x_k]E[z_l]}{M_{11}} + \frac{E[x_k z_1]E[x_1 z_l]}{M_{22}} = \frac{E[x_k z_1]E[x_1 z_l]}{E[x_1 z_1]}$ and $\frac{E[x_{k_1} z_l]}{E[x_{k_1} z_1]} = \frac{E[x_1 z_l]}{E[x_1 z_1]} = \frac{E[x_{k_2} z_l]}{E[x_{k_2} z_1]}$. All other conditions defined by Equ. 177 involving either $P_i(\mathbf{x})$ or $Q_j(\mathbf{z})$ are always true.

[11] According to Lemma 2. For $M_{11} = 1$ and $M_{22} = E[x_1 z_1]$ (see Equ. 177) we obtain $E[x_k z_1 z_l] = \frac{E[x_k]E[z_1 z_l]}{M_{11}} + \frac{E[x_k z_1]E[x_1 z_1 z_l]}{M_{22}} = \frac{E[x_k z_1]E[x_1 z_1 z_l]}{E[x_1 z_1]}$ and $\frac{E[x_{k_1} z_1 z_l]}{E[x_{k_1} z_1]} = \frac{E[x_1 z_1 z_l]}{E[x_1 z_1]} = \frac{E[x_{k_2} z_1 z_l]}{E[x_{k_2} z_1]}$. All other conditions defined by Equ. 177 involving either $P_i(\mathbf{x})$ or $Q_j(\mathbf{z})$ are always true.

for all $n, m \in \mathbb{N}$ and all $c \in \{1, ..., d^{(\mathcal{M})}\}$, where $P_n(\mathbf{x}^{(c)})$ and $Q_m(\mathbf{z}^{(c)})$ denote all monomials in $\mathbf{x}^{(c)}$ and $\mathbf{z}^{(c)}$ and $C^{(c)}_{mj} \in \mathbb{R}$. As shown in the previous section, the conditions in Equ. 32 can again be further simplified for correlated and normalized components of $\mathbf{s}$, i.e.,

$$E[s_d] = 0, \quad E[s_k s_d] \neq 0 \text{ and } E[s_k] = 0 \text{ for all } k \in \{1, ..., d-1\}, \tag{33}$$

and moments as defined for modularizations (Equ. 24 and 25) such that they hold if and only if

$$b^{(c)}_l = \frac{E[x^{(c)}_k z^{(c)}_1 z^{(c)}_l]}{E[x^{(c)}_k z^{(c)}_1] E[z^{(c)}_1 z^{(c)}_l]} \tag{34}$$

for all $k \in \{1, ..., |S_c|\}$, all $l \in \{2, ..., d - |S_c|\}$, all $c \in \{1, ..., d^{(\mathcal{M})}\}$ and $b^{(c)}_l \in \mathbb{R}$.

If a modularization is nested, the corresponding interface variables $y^{(c)}$ in the associated combined undirected graphical model must be organized hierachically as well. Otherwise, each additional link in the combined graph introduces potential dependencies between observable components that violate the individual independence statements of single modules. The conditions above can be tested empirically to potentially falsify any nested modularization.[12]

Moreover, the conditions in Equ. 34 can not only be used to falsify a modularizations, but also to falsify any model of a flat modularization with the same moments as defined in Equ. 24 and 25. More precisely, according to Theorem 10, there exist bounded random variables $\tilde{\mathbf{s}} \in \mathbb{R}^d$ and $\tilde{y}^{(c)} \in \{1, 2\}, c \in \{1, ..., d^{(\mathcal{M})}\}$ such that (i) the elements of $\tilde{\mathbf{s}}$ inside of the $c$-th functional module of $\mathcal{M}$, denoted by $\tilde{\mathbf{x}}^{(c)}$, are independent of the elements of $\tilde{\mathbf{s}}$ outside of the $c$-th functional module of $\mathcal{M}$, denoted by $\tilde{\mathbf{z}}^{(c)}$, given $\tilde{y}^{(c)}$ for all $c \in \{1, ..., d^{(\mathcal{M})}\}$ and (ii)

$$E[\tilde{s}_k \tilde{s}_d] = E[s_k s_d], \quad E[\tilde{s}_k] = E[s_k], \quad E[\tilde{s}_d] = E[s_d] \tag{35}$$

for all $k \in \{1, ..., d-1\}$ and

$$E[\tilde{x}^{(c)}_k \tilde{z}^{(c)}_1 \tilde{z}^{(c)}_l] = E[x^{(c)}_k z^{(c)}_1 z^{(c)}_l] \tag{36}$$

for all $k \in \{1, ..., |S_c|\}$, all $l \in \{2, ..., d - |S_c|\}$ and all $c \in \{1, ..., d^{(\mathcal{M})}\}$ if and only if

$$b^{(c)}_l = \frac{E[x^{(c)}_k z^{(c)}_1 z^{(c)}_l]}{E[x^{(c)}_k z^{(c)}_1] E[z^{(c)}_1 z^{(c)}_l]} \tag{37}$$

for all $k \in \{1, ..., |S_c|\}$, all $l \in \{2, ..., d - |S_c|\}$, all $c \in \{1, ..., d^{(\mathcal{M})}\}$ and $b^{(c)}_l \in \mathbb{R}$.

For nested modularizations some of the conditions in Equ. 37 are redundant. In the remainder of this section, we determine the most concise conditions as used in the next section and introduce an order of modularizations based on these concise conditions.

To simplify notation and without loss of generality,[13] we include each observable $s_k, k \in \{1, ..., d-1\}$ as an additional functional module (if not already present, see

[12]Analogous conditions can be derived for discrete interface variables with more than two values (see Lemma 1).

[13]In general, functional modules consisting of single observables don't impose any conditions and are only applied to simplify notation. Conceptually, any modularization is rejected if and only if the corresponding extended modularization (with additional single observable modules) is rejected and, likewise, there exists a flat modularization with the same moments as defined in Equ. 35 and 36 if and only if the corresponding extended modularization exists.

Fig. S2A). Then, the conditions in Equ. 37 are equivalent to

$$B^{(c_1,c_2)} = \frac{E[x_k^{(c_1)} s_d x_l^{(c_2)}]}{E[x_k^{(c_1)} s_d] E[s_d x_l^{(c_2)}]} \tag{38}$$

for all $k \in \{1, ..., |S_{c_1}|\}$, all $l \in \{1, ..., |S_{c_2}|\}$, all $c_1, c_2 \in \{1, ..., d^{(\mathcal{M})}\}$ with $S_{c_1} \cap S_{c_2} = \{\}$ and $B^{(c_1,c_2)} \in \mathbb{R}$. To avoid the double index $(c_1, c_2)$, we first define

$$B_{kl}^{(\text{all})} = \frac{E[s_k s_d s_l]}{E[s_k s_d] E[s_d s_l]}, \quad k, l \in \{1, ..., d-1\} \tag{39}$$

$$\mathbf{b} = \text{vec}(\mathbf{B}^{(\text{all})}), \tag{40}$$

where $\text{vec}(\mathbf{B}^{(\text{all})})$ denotes the vectorization of $\mathbf{B}^{(\text{all})}$ such that

$$b_{v(k,l)} = B_{kl}^{(\text{all})}, \quad k \in \{1, ..., l-1\}, \ l \in \{2, ..., d-1\} \tag{41}$$

$$v(k, l) = \frac{(d-3)(d-2)}{2} - \frac{(d-2-k)(d-1-k)}{2} + l - 1 \tag{42}$$

and $\mathbf{b} \in \mathbb{R}^{\frac{(d-2)(d-1)}{2}}$ contains all elements above the diagonal of the symmetric matrix $\mathbf{B}^{(\text{all})}$ (see Fig. S2B). Secondly, we define redundant index sets (see Fig. S2C)

$$\begin{aligned} Y_{u(c_1,c_2)} = \ & \{v(k,l) | k, l \in \{1, ..., d-1\} \text{ and } (k < l) \text{ and } (S_{c_1} \cap S_{c_2} = \{\}) \text{ and} \\ & [((s_k \in S_{c_1}) \text{ and } (s_l \in S_{c_2})) \text{ or } ((s_l \in S_{c_1}) \text{ and } (s_k \in S_{c_2}))]\} \end{aligned} \tag{43}$$

$$u(c_1, c_2) = \frac{(d^{(\mathcal{M})} - 2)(d^{(\mathcal{M})} - 1)}{2} - \frac{(d^{(\mathcal{M})} - 1 - c_1)(d^{(\mathcal{M})} - c_1)}{2} + c_2 - 1 \tag{44}$$

for $c_1 \in \{1, ..., c_2 - 1\}, c_2 \in \{2, ..., d^{(\mathcal{M})}\}$, such that the conditions in Equ. 38 are equivalent to

$$b_{q_1} = b_{q_2} \tag{45}$$

for all $q_1, q_2 \in Y_{u(c_1,c_2)}$ and all $c_1 \in \{1, ..., c_2 - 1\}, c_2 \in \{2, ..., d^{(\mathcal{M})}\}$. If all $S_c$, $c \in \{1, ..., d^{(\mathcal{M})}\}$ consist of consecutive indices, then all $Y_u$, $u \in \{1, ..., d^{(\text{u})}\}$, $d^{(\text{u})} = \frac{d^{(\mathcal{M})}(d^{(\mathcal{M})}-1)}{2}$ consist of elements of submatrices of $\mathbf{B}^{(\text{all})}$ (see Fig. S2C).

Finally, we define the set

$$\mathcal{X}^{(\mathcal{M})} = \left\{ Y_{c_1} \middle| c_1 \in \left\{1, ..., d^{(\text{u})}\right\} \text{ and } Y_{c_1} \not\subseteq Y_{c_2} \text{ for all } c_2 \neq c_1, c_2 \in \left\{1, ..., d^{(\text{u})}\right\} \right\} \tag{46}$$

$$d^{(\mathcal{X})} = |\mathcal{X}^{(\mathcal{M})}|. \tag{47}$$

The $d^{(\mathcal{X})}$ elements of $\mathcal{X}^{(\mathcal{M})}$ sorted by their smallest element are denoted by $X_c^{(\mathcal{M})}$ for $c \in \{1, ..., d^{(\mathcal{X})}\}$ and referred to as concise index sets (see Fig. S2D). Then, the conditions in Equ. 38 are equivalent to

$$b_{q_1} = b_{q_2} \tag{48}$$

for all $q_1, q_2 \in X_c^{(\mathcal{M})}$ and all $c \in \{1, ..., d^{(\mathcal{X})}\}$. For correlated observable components of $\mathbf{s}$ these are necessary conditions for the existence of nested modularizations and sufficient and necessary conditions for the existence of models of flat modularizations with the moments defined in Equ. 35 and 36.

We introduce the following partial order over the set of modularizations. Let $\mathcal{M}_1$ and $\mathcal{M}_2$ denote different nested modularizations associated with $d^{(\mathcal{X}_1)}$ and $d^{(\mathcal{X}_2)}$ concise index sets, respectively. Let $X_{c_1}^{(\mathcal{M}_1)}, c_1 \in \{1, ..., d^{(X_1)}\}$ and $X_{c_2}^{(\mathcal{M}_2)}, c_2 \in \{1, ..., d^{(X_2)}\}$ denote the corresponding concise index sets. Then,

$$\mathcal{M}_1 \leq \mathcal{M}_2 \tag{49}$$

if and only if for all $c_1 \in \{1, ..., d^{(X_1)}\}$ the index set $X_{c_1}^{(\mathcal{M}_1)} \subseteq X_{c_2}^{(\mathcal{M}_2)}$ for some $c_2 \in \{1, ..., d^{(X_2)}\}$.[14] Thus, each concise index set associated with $\mathcal{M}_2$ consists of one or multiple concise index sets associated with $\mathcal{M}_1$. For nested modularizations, this is the case if $\mathcal{M}_1 \subseteq \mathcal{M}_2$.[15]

We define two particular modularizations. First, a modularization consisting of all functional modules with one observable component (see Fig. S2E), i.e.

$$\begin{aligned}
\mathcal{M}_0 &= \{S_1, ..., S_{d-1}\}, \quad S_c = \{s_c\}, \quad c \in \{1, ..., d-1\} && (50)\\
X_k^{(\mathcal{M}_0)} &= \{k\}, \quad k \in \{1, ..., d^{(X_0)}\} && (51)\\
d^{(X_0)} &= \frac{(d-2)(d-1)}{2}. && (52)
\end{aligned}$$

Because functional modules with one element impose no constraints (see Equ. 34) and $\mathcal{M}_0$ is a flat modularization, there alway exists a model of $\mathbf{s}$ with modularization $\mathcal{M}_0$ and the moments defined in Equ. 35 and 36.

Secondly, $\mathcal{M}_\mathrm{L}$ denotes a modularization that is associated with a single large concise index set

$$\begin{aligned}
X_1^{(\mathcal{M}_\mathrm{L})} &= \bigcup_{c_0 \in \{1, ..., d^{(X_0)}\}} X_{c_0}^{(\mathcal{M}_0)} && (53)\\
d^{(X_\mathrm{L})} &= 1 && (54)
\end{aligned}$$

(see Fig. S2F). In contrast to all other modularizations, $\mathcal{M}_\mathrm{L}$ is not directly related to specific functional modules but certain distributions of observable states. That is, if the probability distribution of $s_d$ depends only on a binary interface variable, denoted by $y^{(d)}$, and this interface variable depends only on the linear sum of all interface variables of $\mathcal{M}_0$, i.e.

$$y^{(d)} = \begin{cases} 2, & \sum_{c_0=1}^{d-1} y^{(c_0)} = 2(d-1) \\ 1, & \text{otherwise,} \end{cases} \tag{55}$$

then any nested modularization $\mathcal{M}$ that contains the functional modules of $\mathcal{M}_0$ is a valid modularization of $\mathbf{s}$.[16] We obtain

$$\mathcal{M}_0 \leq \mathcal{M} \leq \mathcal{M}_\mathrm{L} \tag{56}$$

for all nested modularizations $\mathcal{M}$.

We also express the conditions for single functional modules (Equ. 28) with the notation of concise index sets to provide a unified framework for the next sections. For this purpose, we order the observables components such that $k < l$ for all $k \in S_1, l \notin S_1$.[17] Furthermore, we extend $\mathbf{B}^{(s)}$ (see Equ.27 and 39) to

$$B_{kl}^{(\mathrm{S})} = \frac{E[s_k s_l]}{E[s_k s_d]}, \quad k, l \in \{1, ..., d-1\}, \tag{57}$$

[14]If $\mathcal{M}_1 \leq \mathcal{M}_2$ and $\mathcal{M}_2 \leq \mathcal{M}_1$ then $\mathcal{M}_2 = \mathcal{M}_1$

[15]This is not an *only if* statement. For any $\mathcal{M}$ different from $\mathcal{M}_0$ and $\mathcal{M}_\mathrm{L}$ follows $\mathcal{M} \not\subseteq \mathcal{M}_\mathrm{L}$ and $\mathcal{M} \leq \mathcal{M}_\mathrm{L}$.

[16]This can be verified by setting the interface variable of each functional module that is an element of $\mathcal{M}$ but not an element of $\mathcal{M}_0$ to 2 if all its inputs are 2 and 1 otherwise.

[17]This is required because $\mathbf{B}^{(\mathrm{S})}$ is not symmetric.

which is square but not symmetric, and apply the same vectorization

$$b_{v(k,l)} = B^{(S)}_{kl}, \quad k \in \{1, ..., l-1\}, l \in \{2, ..., d-1\} \tag{58}$$

and concise index sets $X^{(\mathcal{M}_0)}$, $X^{(\mathcal{M})}$ and $X^{(\mathcal{M}_L)}$ as for multiple modularizations. Thus, **b**, has different values depending on the type of modularization, i.e., consisting of a single or multiple functional modules.[18]

The number of potential modularizations consisting of only a single function module grows approximately exponentially in $d$, i.e.,

$$\sum_{n=2}^{d-1} \binom{d-1}{n} = 2^{d-1} - d - 2, \tag{59}$$

resulting in more than $10^6$ potential modularizations for $d = 21$. To overcome this combinatorial explosion, we propose the following procedures. For flat modularization, we first identify all $d^{(a)}$ potential modularizations consisting of only a single function module and subsequently test for all $\leq 2^{d^{(a)}}$ potential flat modularizations obtained by combining these single functional modules. For nested modularization, we propose to start with an educated guess of a detailed nested modularization consisting of $d^{(b)}$ functional modules and test for all $2^{d^{(b)}}$ potential nested modularizations obtained by combining these functional modules.

#### 2.3 Statistical test assuming independent normal estimates

In this section we define a test statistic for nested modularizations given noisy unbiased estimates of **b** (in Equ. 40), denoted by $\hat{\mathbf{b}}$, that are distributed according to

$$\hat{\mathbf{b}} \sim \mathcal{N}(\mathbf{b}, \mathbf{\Sigma}), \tag{60}$$

where $\mathcal{N}$ denotes the normal distribution with mean **b** and arbitrary full-rank diagonal covariance matrix $\mathbf{\Sigma}$.

Let $\mathcal{M}$ denote a nested modularization associated with $d^{(X)}$ concise index sets and let $X_c^{(\mathcal{M})}, c \in \{1, ..., d^{(X)}\}$ denote the corresponding sets. Let $d^{(X_0)}$ denote the number of concise index sets associated with the modularization $\mathcal{M}_0$ and let $X_c^{(\mathcal{M}_0)}, c \in \{1, ..., d^{(X_0)}\}$ denote the corresponding sets. Then, according to the conditions in Equ. 48 and Theorem 12, the statistic

$$\mathcal{T}_{\mathcal{M}}(\hat{\mathbf{b}}, \mathbf{\Sigma}) = \frac{1}{2} \sum_{c_0=1}^{d^{(X_0)}} \frac{\left(\sum_{k_1 \in X_{c_0}^{(\mathcal{M}_0)}} \hat{b}_{k_1} (\Sigma_{k_1 k_1})^{-1}\right)^2}{\sum_{k_2 \in X_{c_0}^{(\mathcal{M}_0)}} (\Sigma_{k_2 k_2})^{-1}} - \frac{1}{2} \sum_{c=1}^{d^{(X)}} \frac{\left(\sum_{l_1 \in X_c^{(\mathcal{M})}} \hat{b}_{l_1} (\Sigma_{l_1 l_1})^{-1}\right)^2}{\sum_{l_2 \in X_c^{(\mathcal{M})}} (\Sigma_{l_2 l_2})^{-1}} \tag{61}$$

is distributed according to

$$\mathcal{T}_{\mathcal{M}} \sim \Gamma(\zeta_{\mathcal{M}}, 1) \tag{62}$$

$$\zeta_{\mathcal{M}} = \frac{1}{2}\left(d^{(X_0)} - d^{(X)}\right) \tag{63}$$

for $\mathcal{M} \neq \mathcal{M}_0$ under the two null hypotheses that (H1) the observable states **s** form a modularization $\mathcal{M}$ and (H2) there exists a model of a flat modularization $\mathcal{M}$ with the

[18] A test for single functional modules can also be based on $\mathbf{B}^{(\text{all})}$. However, a test based on $\mathbf{B}^{(S)}$ uses the simplest possible moments.

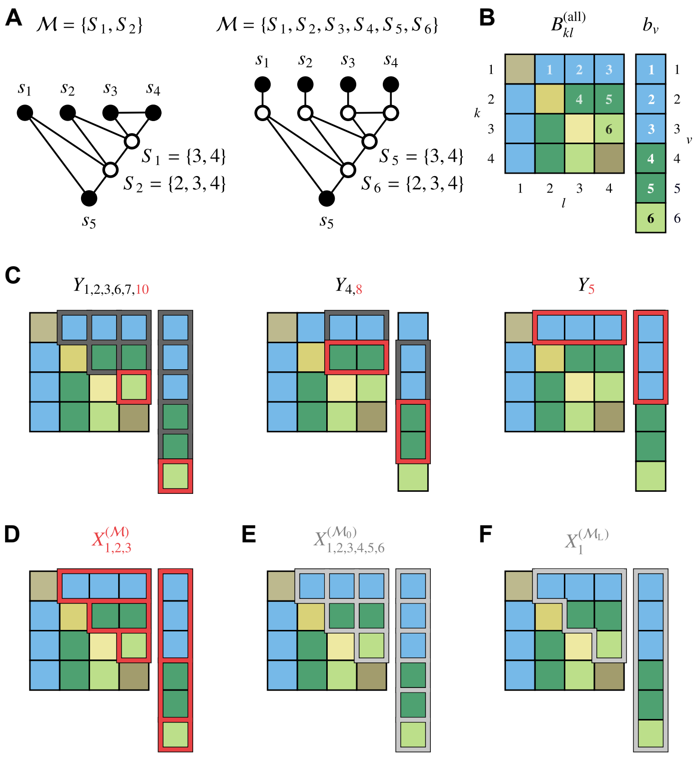


Figure S2: Nested modularizations. **A:** Two modularizations consisting of the functional modules $S_1 = \{3, 4\}$ and $S_2 = \{2, 3, 4\}$ (left) and $S_1 = \{1\}$, $S_2 = \{2\}$, $S_3 = \{3\}$, $S_4 = \{4\}$, $S_5 = \{3, 4\}$ and $S_6 = \{2, 3, 4\}$ (right), respectively. **B:** $\mathbf{B}^{(\text{all})}$ and its vectorization $\mathbf{b}$ associated with both modularizations shown in panel A. Matrix and vector elements with the same color have the same value. **C:** Redundant index sets for the modularizations shown in panel A. All elements within each boundary are elements of the same set. The non-empty redundant index sets for $\mathbf{b}$ are $Y_1 = \{1\}$, $Y_2 = \{2\}$, $Y_3 = \{3\}$, $Y_4 = \{2, 3\}$, $Y_5 = \{1, 2, 3\}$, $Y_6 = \{4\}$, $Y_7 = \{5\}$, $Y_8 = \{4, 5\}$, and $Y_{10} = \{6\}$. Red boundaries indicate sets selected as concise index sets. **D:** Concise index sets for the modularization $\mathcal{M}$ shown in panel A. **E:** Concise index sets for the modularization $\mathcal{M}_0$ and five observable components. A model with modularization $\mathcal{M}_0$ always exists. **F:** Concise index set for the modularization $\mathcal{M}_L$ and five observable components. There exists no distribution of observable states that corresponds to $\mathcal{M}_L$ for $\mathbf{B}^{(\text{all})}$ shown in panel B.

moments defined in Equ. 35 and 36. Here, we assume that $s_d$ is correlated with all other observable components and $\Gamma(\zeta_{\mathcal{M}}, 1)$ denotes the gamma distribution with shape parameter $\zeta_{\mathcal{M}}$ and scale parameter 1.

The test statistic $\mathcal{T}_{\mathcal{M}}$ is related to the posterior probability of a nested modularization $\mathcal{M}$ given the estimates $\hat{\mathbf{b}}$ if (i) the covariance matrix $\mathbf{\Sigma}$ is known and (ii) the prior distributions for $\mathbf{b}$ and $\mathcal{M}$ are non-informative. Then, the log odd of the posterior probability of $\mathcal{M}$ and the posterior probability of $\mathcal{M}_0$, i.e.,[19]

$$\text{logit}_{\mathcal{M}_0}(\mathcal{M}) = \log \frac{\text{P}(\mathcal{M}_0|\hat{\mathbf{b}}, \mathbf{\Sigma})}{\text{P}(\mathcal{M}|\hat{\mathbf{b}}, \mathbf{\Sigma})}, \tag{64}$$

[19]To simplify notation in the context of probabilistic inference (Lemma 13 and Equ. 64), we write $\text{P}(\mathcal{M})$ to denote a distribution evaluated for the particular value $\mathcal{M}$, provided that the interpretation is clear from the context.

can be derived analytically (see Lemma 13) and related to $\mathcal{T}_{\mathcal{M}}$ by

$$\mathcal{T}_{\mathcal{M}} = \text{logit}_{\mathcal{M}_0}(\mathcal{M}) + \zeta_{\mathcal{M}}. \tag{65}$$

Here, we call a prior distribution of $\mathcal{M}$ non-informative if the expected log odds of all $\mathcal{M}$ that fulfill the null hypotheses are identical.

For fixed $\mathcal{M}$, the ratio of the priors $\frac{\text{P}(\mathcal{M}_0)}{\text{P}(\mathcal{M})}$ is constant and finite (see Equ. 393) and, according to Bayes' theorem, $\mathcal{T}_{\mathcal{M}}$ is identical to the log-likelihood ratio for the two hypotheses $\mathcal{M}_0$ and $\mathcal{M}$ shifted by an irrelevant additive constant.[20] By the Neyman-Pearson theory, a statistical test based on a log-likelihood ratio is most powerful among all level $\alpha$ tests and its statistic is minimal sufficient.[21] Furthermore, if $\hat{\mathbf{b}}$ is obtained by sampling and converges asymptotically to a joint normal distribution, then the test is also consistent, i.e., its power for any incorrect flat $\mathcal{M}$ converges asymptotically to one.[22]

Finally, if $\mathcal{M}_1$ and $\mathcal{M}_2$ denote two modularizations such that $\mathcal{M}_1 \leq \mathcal{M}_2$,[23]

$$E[\text{logit}_{\mathcal{M}_0}(\mathcal{M}_1)] \leq E[\text{logit}_{\mathcal{M}_0}(\mathcal{M}_2)] \tag{66}$$

and, thus,

$$0 \leq E[\text{logit}_{\mathcal{M}_0}(\mathcal{M}_1)] \leq E[\text{logit}_{\mathcal{M}_0}(\mathcal{M}_2)] \leq E[\text{logit}_{\mathcal{M}_0}(\mathcal{M}_\text{L})]. \tag{67}$$

Fig. S3 illustrates statistical inference for the nested modularization shown in Fig. S2A.

#### 2.4 Asymptotic test for modularizations

The inference in the previous section is precise under the assumptions that $\hat{\mathbf{b}} \sim \mathcal{N}(\mathbf{b}, \mathbf{\Sigma})$, $s_d$ is correlated with all other observable components and $\mathbf{\Sigma}$ is known, invertible and diagonal. In this section, we derive an asymptotically conservative test that assumes only that the covariance matrix of all estimated sample moments ($\mathbf{\Sigma}'^{(\text{all})}$ in Equ. 153) is invertible.

Let $N$ denote half of the total number of i.i.d. samples of $s_k$, $k \in \{1, ..., d\}$ and let $\hat{s}_k^{(n)}, n \in \{1, ..., 2N\}$ denote the $n$-th sample. This definition has the advantage that $N$ denotes the total number of samples used to calculated sample moments and, thus, the corresponding means, variances and asymptotic distributions (e.g., the central limit theorem) are stated in their conventional form.

We estimate the sample means

$$\hat{b}_i^{(\nu(k,l))} = \frac{1}{N} \sum_{n=1}^{N} \hat{\Phi}_{ni}^{(\nu(k,l))} \tag{68}$$

of the numerators and denominators of $\mathbf{b}$ and their sample covariance matrices

$$\hat{\Sigma}'^{(\nu,\omega)}_{ij} = \frac{1}{N-1} \sum_{n=1}^{N} \left(\hat{\Phi}_{ni}^{(\nu)} - \hat{b}_i^{(\nu)}\right)\left(\hat{\Phi}_{nj}^{(\omega)} - \hat{b}_j^{(\omega)}\right) \tag{69}$$

$$\hat{\Sigma}'^{(\nu)}_{ij} = \hat{\Sigma}'^{(\nu,\nu)}_{ij}, \tag{70}$$

[20] For the log-likelihood ratio test $\mathcal{M}$ is considered as null hypothesis to be rejected because the alternative hypothesis $\mathcal{M}_0$ is always true. This null hypothesis is either H1 or H2.

[21] (Van der Vaart, 2000, chapter 16.1)

[22] According to Lemma 14

[23] According to Lemma 15.

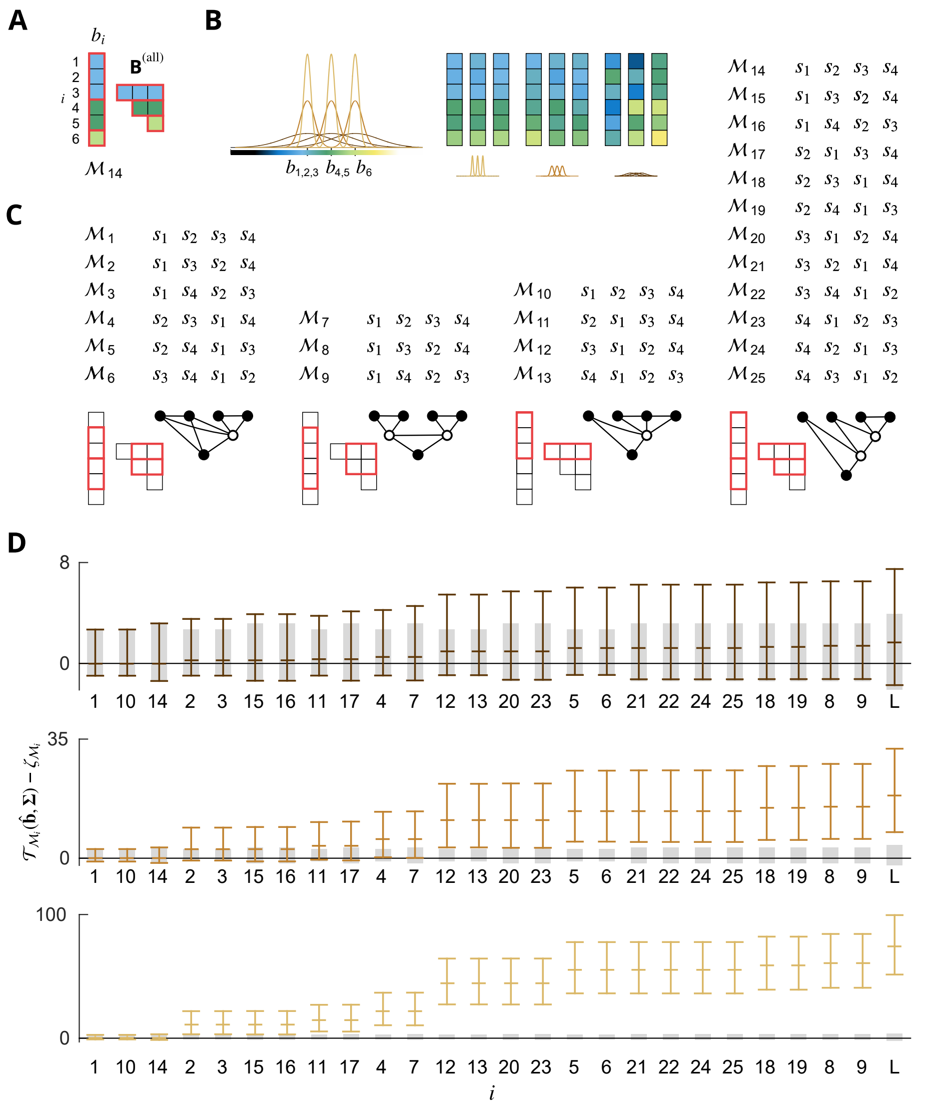

Figure S3: Statistical inference for nested modularizations. **A:** $\mathbf{B}^{(\text{all})}$ and its vectorization $\mathbf{b}$ associated with the modularizations shown in Fig. S2A. Matrix and vector elements with the same color have the same value. Red boundaries indicate concise index sets. **B:** Estimates $\hat{b}_i$ for $i \in \{1, ..., 6\}$ (right) distributed according to a joint normal distribution with means $b_i$ and standard deviations of either 0.15 (light brown lines), 0.3 (brown lines) or 1 (dark brown lines) times the difference $b_4 - b_1 = b_6 - b_4$ (left). All $\hat{b}_i$ are independent. Three samples for each of the three distributions are shown on the right. **C:** All modularizations for five observable components except $\mathcal{M}_0$ and $\mathcal{M}_\text{L}$. Concise index sets (red boundaries) with one element are not shown. **D:** Test statistic $\mathcal{T}_\mathcal{M}(\hat{\mathbf{b}}, \mathbf{\Sigma})$ for $\hat{\mathbf{b}} \sim \mathcal{N}(\mathbf{b}, \mathbf{\Sigma})$ and parameters $\mathbf{b}$ and $\mathbf{\Sigma}$ shown in panel B. Different values for the diagonal elements of $\mathbf{\Sigma}$ are color coded in light brown, brown and dark brown. Gray bars indicate the 2.5th and 97.5th percentiles of the distributions of the test statistics given the corresponding modularizations apply. Colored bars indicate the 2.5th percentile, the mean and the 97.5th percentile of the distributions of the test statistics for $\hat{\mathbf{b}}$ distributed according to panel B (same color code).

$i, j \in \{1, 2\}$, $v, \omega \in \{1, ..., d^{(X_0)}\}$, $k \in \{1, ..., l\}$, $l \in \{1, ..., d-1\}$ and $\omega(k, l) = v(k, l)$, where

$$\hat{\Phi}_{n1}^{(v(k,l))} = \hat{s}_k^{(2n-1)} \hat{s}_l^{(2n-1)} \hat{s}_d^{(2n-1)} \quad (71)$$

$$\hat{\Phi}_{n2}^{(v(k,l))} = \hat{s}_k^{(n)} \hat{s}_d^{(n)} \hat{s}_l^{(n+N)} \hat{s}_d^{(n+N)} \quad (72)$$

for modularizations consisting of multiple functional modules and

$$\hat{\Phi}_{n1}^{(v(k,l))} = \hat{s}_k^{(n)} \hat{s}_l^{(n)} \quad (73)$$

$$\hat{\Phi}_{n2}^{(v(k,l))} = \hat{s}_k^{(n)} \hat{s}_d^{(n)} \quad (74)$$

for modularizations consisting of a single functional module.[24]

We denote the corresponding population means and covariance matrices by

$$b_i^{(v)} = E\left[\hat{b}_i^{(v)}\right] \quad (75)$$

$$\Sigma'^{(v,\omega)}_{ij} = NE\left[\left(\hat{b}_i^{(v)} - b_i^{(v)}\right)\left(\hat{b}_j^{(\omega)} - b_j^{(\omega)}\right)\right] \quad (76)$$

$$\Sigma'^{(v)}_{ij} = \Sigma'^{(v,v)}_{ij} \quad (77)$$

for $i, j \in \{1, 2\}$, $v, \omega \in \{1, ..., d^{(X_0)}\}$, so that

$$b_1^{(v)} = E[s_k s_l s_d] \quad (78)$$

$$b_2^{(v)} = E[s_k s_d] E[s_l s_d] \quad (79)$$

for modularizations consisting of multiple functional modules (see Equ. 39),

$$b_1^{(v)} = E[s_k s_l] \quad (80)$$

$$b_2^{(v)} = E[s_k s_d] \quad (81)$$

for modularizations consisting of a single functional module (see Equ. 57) and

$$b_{v(k,l)} = \frac{b_1^{(v)}}{b_2^{(v)}}, \quad (82)$$

for both types of modularizations, where we assume $b_2^{(v)} \neq 0$, so that $b_{v(k,l)}$ is defined.[25] However, the derived test statistic is also asymptotically precise for the case $b_2^{(v)} = 0$ and the assumptions that all observable components are correlated with $s_d$ and $b_2^{(v)} \neq 0$ will be dropped below.[26]

According to the multidimensional central limit theorem (Van der Vaart, 2000), the distributions of $\hat{b}_1^{(v)}$ and $\hat{b}_2^{(v)}$ converge for invertible covariance matrix $\mathbf{\Sigma}'^{(v)}$ to the joint normal distribution

$$\boldsymbol{\rho}'^{(v)} = \sqrt{N}\left(\hat{\mathbf{b}}^{(v)} - \mathbf{b}^{(v)}\right) \xrightarrow{d} \mathcal{N}\left(\mathbf{0}, \mathbf{\Sigma}'^{(v)}\right) \quad (83)$$

as $N \to \infty$, where $\xrightarrow{d}$ denotes convergence in distribution and $v \in \{1, ..., d^{(X_0)}\}$.

[24]For $\mathcal{M}$ consisting of a single functional module, $N$ can also denote the total number of samples.

[25]Then, $\mu_{v(k,l)}$ in Equ. 107 is defined as well.

[26]See beginning of section *Sufficient and necessary conditions for models of functional modules.*

**Standard form of estimates**

For finite $N$ and normal $\hat{b}_1^{(v)}$ and $\hat{b}_2^{(v)}$ with covariance matrix

$$\mathbf{\Sigma}^{(v)} = \frac{\mathbf{\Sigma}'^{(v)}}{N}, \quad (84)$$

the ratio $\frac{\hat{b}_1^{(v)}}{\hat{b}_2^{(v)}}$ can be transformed into the standard form (Marsaglia et al., 2006)

$$\begin{aligned} \frac{\hat{b}_1^{(v)}}{\hat{b}_2^{(v)}} &= \Theta_3(\mathbf{\Sigma}^{(v)})\frac{\hat{\mu}_1^{(v)}(\mathbf{\Sigma}^{(v)})}{\hat{\mu}_2^{(v)}(\mathbf{\Sigma}^{(v)})} + \Theta_4(\mathbf{\Sigma}^{(v)}) && (85)\\ \hat{\mu}_1^{(v)}(\mathbf{\Sigma}^{(v)}) &= \mu_1^{(v)}(\mathbf{\Sigma}^{(v)}) + \rho_1^{(v)}(\mathbf{\Sigma}^{(v)}) && (86)\\ \hat{\mu}_2^{(v)}(\mathbf{\Sigma}^{(v)}) &= \mu_2^{(v)}(\mathbf{\Sigma}^{(v)}) + \rho_2^{(v)}(\mathbf{\Sigma}^{(v)}), && (87) \end{aligned}$$

where

$$\begin{aligned} \mu_1^{(v)}(\mathbf{\Sigma}^{(v)}) &= \Theta_1\left(b_1^{(v)}, b_2^{(v)}, \mathbf{\Sigma}^{(v)}\right) && (88)\\ \mu_2^{(v)}(\mathbf{\Sigma}^{(v)}) &= \Theta_2\left(b_2^{(v)}, \mathbf{\Sigma}^{(v)}\right) && (89) \end{aligned}$$

are non-negative constants and

$$\begin{aligned} \rho_1^{(v)}(\mathbf{\Sigma}^{(v)}) &= \Theta_1\left(\rho_1'^{(v)}, \rho_2'^{(v)}, N\mathbf{\Sigma}^{(v)}\right) && (90)\\ \rho_2^{(v)}(\mathbf{\Sigma}^{(v)}) &= \Theta_2\left(\rho_2'^{(v)}, N\mathbf{\Sigma}^{(v)}\right) && (91) \end{aligned}$$

are joint standard normal random variables[27] obtained by the functions

$$\begin{aligned} \Theta_1(x, y, \mathbf{\Sigma}^{(v)}) &= \left(\frac{x}{\sqrt{\Sigma_{11}^{(v)}}} - \frac{\Sigma_{12}^{(v)} y}{\sqrt{\Sigma_{11}^{(v)}}\Sigma_{22}^{(v)}}\right)\left(1 - \frac{\Sigma_{12}^{(v)2}}{\Sigma_{11}^{(v)}\Sigma_{22}^{(v)}}\right)^{-1/2} && (92)\\ \Theta_2(y, \mathbf{\Sigma}^{(v)}) &= \frac{y}{\sqrt{\Sigma_{22}^{(v)}}} && (93)\\ \Theta_3(\mathbf{\Sigma}^{(v)}) &= \sqrt{\frac{\Sigma_{11}^{(v)}}{\Sigma_{22}^{(v)}}\left(1 - \frac{\Sigma_{12}^{(v)2}}{\Sigma_{11}^{(v)}\Sigma_{22}^{(v)}}\right)} && (94)\\ \Theta_4(\mathbf{\Sigma}^{(v)}) &= \frac{\Sigma_{12}^{(v)}}{\Sigma_{22}^{(v)}} && (95) \end{aligned}$$

If the covariance matrix $\mathbf{\Sigma}^{(v)}$ is invertible then $\Sigma_{11}^{(v)}$, $\Sigma_{22}^{(v)}$ and $\Theta_3$ are non-zero. We obtain

$$\begin{aligned} \hat{\mu}_1^{(v)}(\mathbf{\Sigma}^{(v)}) &= \Theta_1\left(\hat{b}_1^{(v)}, \hat{b}_2^{(v)}, \mathbf{\Sigma}^{(v)}\right) && (96)\\ \hat{\mu}_2^{(v)}(\mathbf{\Sigma}^{(v)}) &= \Theta_2\left(\hat{b}_2^{(v)}, \mathbf{\Sigma}^{(v)}\right), && (97) \end{aligned}$$

which can be calculated given $\mathbf{\Sigma}^{(v)}$ without knowledge of $b_1^{(v)}$ and $b_2^{(v)}$.

[27] We only denote the dependence of $\hat{\boldsymbol{\mu}}^{(v)}$, $\boldsymbol{\mu}^{(v)}$ and $\boldsymbol{\rho}^{(v)}$ on $\mathbf{\Sigma}^{(v)}$ in contrast to the additional dependence on $\mathbf{b}^{(v)}$, $\boldsymbol{\rho}'^{(v)}$ and $N$.

Unfortunately, the mean of $\frac{\hat{b}_1^{(v)}}{\hat{b}_2^{(v)}}$ does not exists (Marsaglia et al., 2006) and cannot be used to estimate $b_v$. Instead, we define

$$\hat{b}_v(\mathbf{\Sigma}^{(v)}) = \begin{cases} b_v, & \left|\hat{\mu}_2^{(v)}(\mathbf{\Sigma}^{(v)})\right| < \theta_{\hat{\mu}} \\ \frac{\hat{b}_1^{(v)}}{\hat{b}_2^{(v)}}, & \text{otherwise} \end{cases} \tag{98}$$

with cutoff $\theta_{\hat{\mu}} > 0$ and set $\mathcal{T}_{\mathcal{M}} = 0$ if $\left(\left|\hat{\mu}_2^{(v)}(\mathbf{\Sigma}^{(v)})\right| < \theta_{\hat{\mu}} \text{ for all } v \in \{1, ..., d^{(\mathcal{X}_0)}\}\right)$.[28] Then, all moments of $\hat{b}_v$ exist.[29]

However, the distribution of this estimator is not normal, different $\hat{b}_v$ might be correlated and $\hat{b}_v$ is, in general, biased, i.e., $b_v \neq E[\hat{b}_v]$, despite that $\hat{b}_1^{(v)}$ and $\hat{b}_2^{(v)}$ are unbiased estimators of $b_1^{(v)}$ and $b_2^{(v)}$, respectively. In the following, we remedy this shortcomings that prevent precise inference as outlined in the previous section by introducing a more conservative test statistic.

**Asymptotic test assuming uncorrelated sample moment ratios**

For finite $N$ the variance of $\hat{b}_v$ is unknown even if $\mathbf{\Sigma}'^{(v)}$ is given, because it depends on the unknown mean $\mathbf{b}^{(v)}$ (see Equ. 98). In this section we introduce an approximation of the variance of $\hat{b}_v$ which is asymptotically correct as $N \to \infty$ and depends only on the sample estimates of $\mathbf{b}^{(v)}$ and $\mathbf{\Sigma}'^{(v)}$. We first derive a test statistic for uncorrelated $\hat{b}_v$ and extend the results in the final sections to correlated $\hat{b}_v$.

For large $N$, the distribution of $\frac{\hat{b}_1^{(v)}}{\hat{b}_2^{(v)}}$ will be sharply peaked around $\frac{b_1^{(v)}}{b_2^{(v)}}$. We make the ansatz to approximate the variance of $\frac{\hat{\mu}_1^{(v)}}{\hat{\mu}_2^{(v)}}$ by the variance of its second order Taylor series at the point $\left(\mu_1^{(v)}, \mu_2^{(v)}\right)$ under the assumption that the denominator has zero mass around zero (Stuart and Ord, 1994)[30]

$$\begin{aligned} \mathrm{VAR}\left[\frac{\hat{b}_1^{(v)}}{\hat{b}_2^{(v)}}\right] &= \mathrm{VAR}\left[\frac{\hat{\mu}_1^{(v)}}{\hat{\mu}_2^{(v)}}\right]\Theta_3(\mathbf{\Sigma}^{(v)})^2 && (99)\\ &\approx \left(\frac{1}{{\mu_2}^{(v)\,2}} + \frac{{\mu_1}^{(v)\,2}}{{\mu_2}^{(v)\,4}}\right)\Theta_3(\mathbf{\Sigma}^{(v)})^2 && (100)\\ &= \frac{1}{N}\left(\frac{1}{\Theta_2\left(b_2^{(v)}, \mathbf{\Sigma}'^{(v)}\right)^2} + \frac{\Theta_1\left(b_1^{(v)}, b_2^{(v)}, \mathbf{\Sigma}'^{(v)}\right)^2}{\Theta_2\left(b_2^{(v)}, \mathbf{\Sigma}'^{(v)}\right)^4}\right)\Theta_3(\mathbf{\Sigma}'^{(v)})^2, && (101) \end{aligned}$$

where the multiplicative factor $\Theta_3(\mathbf{\Sigma}^{(v)})^2 = \Theta_3(\mathbf{\Sigma}'^{(v)})^2$ follows from Equ. 85.

---

[28]Only ratios $\frac{\hat{b}_v - b_v}{\Sigma_{vv}^{1/2}}$ occur in the test statistic and $\hat{b}_v = b_v$ can be obtained without knowledge of $b_v$ by setting $(\Sigma_{vv})^{-1} = 0$ if $\left|\hat{\mu}_2^{(v)}\right| < \theta_{\hat{\mu}}$.

[29]If $\left|\hat{\mu}_2^{(v)}\right| \geq \theta_{\hat{\mu}}$, the absolute value of the denominator of the standard form in Equ. 85, i.e., $\hat{\mu}_2^{(v)}$, is bounded from below by $\theta_{\hat{\mu}}$ and the numerator $\hat{\mu}_1^{(v)}$ is normal.

[30]From the first order Taylor series for $f(x, y) = \frac{x}{y}$ around $(x_0, y_0)$ and $E[f(x, y)] \approx f(x_0, y_0)$ follows $\mathrm{VAR}[f(x, y)] \approx E\left[\left(f(x_0, y_0) + f_x(x_0, y_0)(x - x_0) + f_y(x_0, y_0)(y - y_0) - E[f(x, y)]\right)^2\right]$ and $\mathrm{VAR}[f(x, y)] \approx \frac{x_0^2}{y_0^2}\left(\frac{\mathrm{VAR}[x]}{x_0^2} - 2\frac{\mathrm{COV}[x, y]}{x_0 y_0} + \frac{\mathrm{VAR}[y]}{y_0^2}\right)$. (Stuart and Ord, 1994, p. 351)

Because $\mathbf{\Sigma}'^{(v)}$ is unknown, we substitute its sample covariance matrix $\hat{\mathbf{\Sigma}}'^{(v)}$ and define the asymptotically correct sample covariance matrix of $\hat{\mathbf{b}}$

$$\hat{\Sigma}^{(\infty)}_{v\omega}(\hat{\mathbf{\Sigma}}^{(v)}) = \begin{cases} \hat{\sigma}^{(\mathrm{T})}_{v}(\hat{\mathbf{\Sigma}}^{(v)})^{2}\;\Theta_3(\hat{\mathbf{\Sigma}}^{(v)})^{2}, & v=\omega \\ 0, & \text{otherwise} \end{cases} \tag{102}$$

$$\hat{\sigma}^{(\mathrm{T})}_{v}(\hat{\mathbf{\Sigma}}^{(v)})^{2} = \begin{cases} \hat{\sigma}_0, & \left|\hat{\mu}^{(v)}_2(\hat{\mathbf{\Sigma}}^{(v)})\right| < \theta_{\hat{\mu}} \\ \frac{1}{\hat{\mu}^{(v)}_2(\hat{\mathbf{\Sigma}}^{(v)})^2} + \frac{\hat{\mu}^{(v)}_1(\hat{\mathbf{\Sigma}}^{(v)})^2}{\hat{\mu}^{(v)}_2(\hat{\mathbf{\Sigma}}^{(v)})^4}, & \text{otherwise} \end{cases} \tag{103}$$

for $v,\omega \in \{1,...,d^{(X_0)}\}$, where $\hat{\sigma}_0 > 0$ is an arbitrary constant and $\hat{\mathbf{\Sigma}}^{(v)} = \hat{\mathbf{\Sigma}}'^{(v)}/N$. Here, we assume uncorrelated $\hat{b}_v$ and diagonal $\hat{\mathbf{\Sigma}}^{(\infty)}$. We extend the results in the final sections to correlated $\hat{b}_v$.

Asymptotically, $\hat{b}^{(v)}_1 \xrightarrow{p} b^{(v)}_1$, $\hat{b}^{(v)}_2 \xrightarrow{p} b^{(v)}_2$, $\hat{\mathbf{\Sigma}}'^{(v)} \xrightarrow{p} \mathbf{\Sigma}'^{(v)}$ and $(\hat{b}^{(v)}_1, \hat{b}^{(v)}_2, \hat{\mathbf{\Sigma}}'^{(v)}) \xrightarrow{p} (b^{(v)}_1, b^{(v)}_2, \mathbf{\Sigma}'^{(v)})$ as $N \to \infty$,[31] where $\xrightarrow{p}$ denotes convergence in probability. It follows from Equ. 100 and 101 and the continuous mapping theorem[32]

$$N\hat{\Sigma}^{(\infty)}_{vv}(\hat{\mathbf{\Sigma}}^{(v)}) \xrightarrow{p} \Sigma'_{vv} \tag{104}$$

$$\Sigma'_{vv} = \left( \frac{1}{\Theta_2\left(b^{(v)}_2, \mathbf{\Sigma}'^{(v)}\right)^2} + \frac{\Theta_1\left(b^{(v)}_1, b^{(v)}_2, \mathbf{\Sigma}'^{(v)}\right)^2}{\Theta_2\left(b^{(v)}_2, \mathbf{\Sigma}'^{(v)}\right)^4} \right) \Theta_3(\mathbf{\Sigma}'^{(v)})^2 \tag{105}$$

as $N \to \infty$. We define $\Sigma_{vv} = \Sigma'_{vv}/N$, so that $N\hat{\Sigma}^{(\infty)}_{vv}(\hat{\mathbf{\Sigma}}^{(v)}) \xrightarrow{p} N\Sigma_{vv}$ as $N \to \infty$.

We now verify that the second order Taylor series approximation in Equ. 100 and, thus, $\hat{\Sigma}^{(\infty)}_{vv}$ are indeed asymptotically correct. According to Lemma 16 (for $\lambda_v = 1$),

$$\frac{\hat{b}_v(\hat{\mathbf{\Sigma}}^{(v)}) - b_v}{\sqrt{\hat{\Sigma}^{(\infty)}_{vv}(\hat{\mathbf{\Sigma}}^{(v)})}} = \frac{\hat{\mu}_v(\hat{\mathbf{\Sigma}}^{(v)}) - \mu_v(\hat{\mathbf{\Sigma}}^{(v)})}{\hat{\sigma}^{(\mathrm{T})}_{v}(\hat{\mathbf{\Sigma}}^{(v)})}, \tag{106}$$

where we defined

$$\mu_v(\mathbf{\Sigma}^{(v)}) = \frac{\mu^{(v)}_1(\mathbf{\Sigma}^{(v)})}{\mu^{(v)}_2(\mathbf{\Sigma}^{(v)})} \tag{107}$$

$$\hat{\mu}_v(\mathbf{\Sigma}^{(v)}) = \begin{cases} \mu_v(\mathbf{\Sigma}^{(v)}), & \left|\hat{\mu}^{(v)}_2(\mathbf{\Sigma}^{(v)})\right| < \theta_{\hat{\mu}} \\ \frac{\hat{\mu}^{(v)}_1(\mathbf{\Sigma}^{(v)})}{\hat{\mu}^{(v)}_2(\mathbf{\Sigma}^{(v)})}, & \text{otherwise.} \end{cases} \tag{108}$$

According to Lemma 17 (for $\lambda_v = 1$),

$$\frac{\sqrt{N}(\hat{b}_v(\hat{\mathbf{\Sigma}}^{(v)}) - b_v)}{\sqrt{N\hat{\Sigma}^{(\infty)}_{vv}(\hat{\mathbf{\Sigma}}^{(v)})}} = \frac{\hat{\mu}_v(\hat{\mathbf{\Sigma}}^{(v)}) - \mu_v(\hat{\mathbf{\Sigma}}^{(v)})}{\hat{\sigma}^{(\mathrm{T})}_{v}(\hat{\mathbf{\Sigma}}^{(v)})} \xrightarrow{d} \mathcal{N}(0,1) \tag{109}$$

as $N \to \infty$. Therefore, the test statistic in Equ. 61 is asymptotically correct, i.e.,[33]

$$\mathcal{T}_{\mathcal{M}}(\hat{\mathbf{b}}, \hat{\mathbf{\Sigma}}^{(\infty)}) = \mathcal{T}_{\mathcal{M}}\left(\sqrt{N}\hat{\mathbf{b}}, N\hat{\mathbf{\Sigma}}^{(\infty)}\right) = \mathcal{T}_{\mathcal{M}}\left(\sqrt{N}(\hat{\mathbf{b}} - \mathbf{b}), N\hat{\mathbf{\Sigma}}^{(\infty)}\right) \xrightarrow{d} \Gamma(\zeta_{\mathcal{M}}, 1) \tag{110}$$

as $N \to \infty$ for $\mathcal{M} \neq \mathcal{M}_0$ under the two null hypotheses.

[31](Van der Vaart, 2000, theorem 2.7 (vi))

[32](Van der Vaart, 2000, theorem 2.3)

[33]By the continuous mapping theorem applied to Equ. 61, Equ. 105 and Equ. 109. According to Lemma 11, $\mathcal{T}_{\mathcal{M}}$ is independent of $\sqrt{N}\mathbf{b}$ when added to the first argument of $\mathcal{T}_{\mathcal{M}}$.

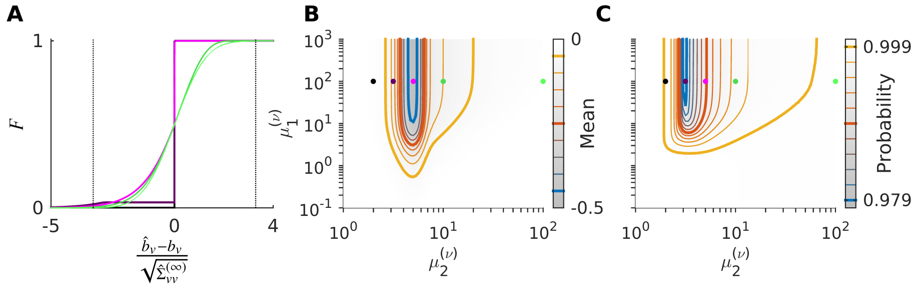


Figure S4: Cumulative distribution function $F$ (**A**), mean (**B**) and upper bound for the probability mass within the interval $\pm 3.2905$ (**C**) of $\frac{\hat{b}_v - b_v}{\sqrt{\hat{\Sigma}^{(\infty)}_{vv}}}$ for fixed $\boldsymbol{\mu}^{(v)}$ and $\theta_{\hat{\mu}} = 5$ as $N \to \infty$. For standard normal random variables the values in panel B and C are 0 and 0.999, respectively. The colors of the lines in panel A and dots in panel B and C correspond to $\mu_2^{(v)} \in \{10^{0.3}, 10^{0.5}, 10^{0.7}, 10^1, 10^2\}$ and $\mu_1^{(v)} = 10^2$. Dotted lines in panel A indicate the interval $\pm 3.2905$.

#### Conservative test assuming normal estimates and uncorrelated sample moment ratios

For noisy data and weak correlations (see Equ. 75) the distribution of $\frac{\hat{b}_v - b_v}{\sqrt{\hat{\Sigma}^{(\infty)}_{vv}}}$ converges slowly and might be far from standard normal even if the number of samples is large and $\sqrt{N}\left(\hat{\mathbf{b}}^{(v)} - \mathbf{b}^{(v)}\right)$ is approximately joint normal.

In particular, for weakly correlated observables $\mathbf{s}$ (see Equ. 75) the absolute values of $b_1^{(v)}$ and $b_2^{(v)}$ and, thus, $\mu_1^{(v)}$ and $\mu_2^{(v)}$ are small and $\hat{\mu}_1^{(v)}(\boldsymbol{\Sigma}^{(v)})/\hat{\mu}_2^{(v)}(\boldsymbol{\Sigma}^{(v)})$ (in Equ. 106) is not close to normal even if $\hat{\mu}_1^{(v)}(\boldsymbol{\Sigma}^{(v)})$ and $\hat{\mu}_2^{(v)}(\boldsymbol{\Sigma}^{(v)})$ are normal. Furthermore, the sample variance $\hat{\Sigma}^{(\infty)}_{vv}$ is a random variable that results in a non-normal distribution of $\frac{\hat{b}_v - b_v}{\sqrt{\hat{\Sigma}^{(\infty)}_{vv}}}$. The bias of $\frac{\hat{b}_v - b_v}{\sqrt{\hat{\Sigma}^{(\infty)}_{vv}}}$ for fixed $\mu_1^{(v)}$ and $\mu_2^{(v)}$ as $N \to \infty$ is shown in Fig. S4.

We now construct a conservative test statistic that applies to joint normal numerators and denominators of $\hat{b}_v$ , i.e., $\boldsymbol{\rho}'^{(v)} = \sqrt{N}\left(\hat{\mathbf{b}}^{(v)} - \mathbf{b}^{(v)}\right) \sim \mathcal{N}\left(\mathbf{0}, \boldsymbol{\Sigma}'^{(v)}\right)$ (see Equ. 83) and known covariance, i.e., $\hat{\boldsymbol{\Sigma}}'^{(v)} = \boldsymbol{\Sigma}'^{(v)}$ (see Equ. 70). For normal and i.i.d. samples[34]

$$\frac{\mathrm{VAR}\left[\hat{\Sigma}^{(v)\,\frac{1}{2}}_{ii}\right]}{\mathrm{VAR}[\hat{b}^{(v)}_i]} = 1 - c_4(N)^2 \tag{111}$$

$$c_4(N) = \sqrt{\frac{2}{N-1}}\,\frac{\Gamma\left(\frac{N}{2}\right)}{\Gamma\left(\frac{N-1}{2}\right)}, \tag{112}$$

where $i \in \{1, 2\}$ and $\Gamma$ denotes the Gamma function, with values less than $10^{-2}$ and $10^{-3}$ for more than 50 and 500 samples, respectively. Thus, for many datasets about 500 samples are sufficient to approximately fulfill the requirements above.

For the construction of the conservative test statistic, we apply four steps. First, we define a modified test statistic that is asymptotically more conservative than the gamma distribution $\Gamma(\zeta_{\mathcal{M}}, 1)$ of the original test statistic $\mathcal{T}_{\mathcal{M}}(\hat{\mathbf{b}}, \hat{\boldsymbol{\Sigma}}^{(\infty)})$ (see Equ. 110). Secondly, for finitely many samples, joint normal $\boldsymbol{\rho}'^{(v)}$ and known $\boldsymbol{\Sigma}'^{(v)}$, we define a finite volume

[34] According to Proposition 18.

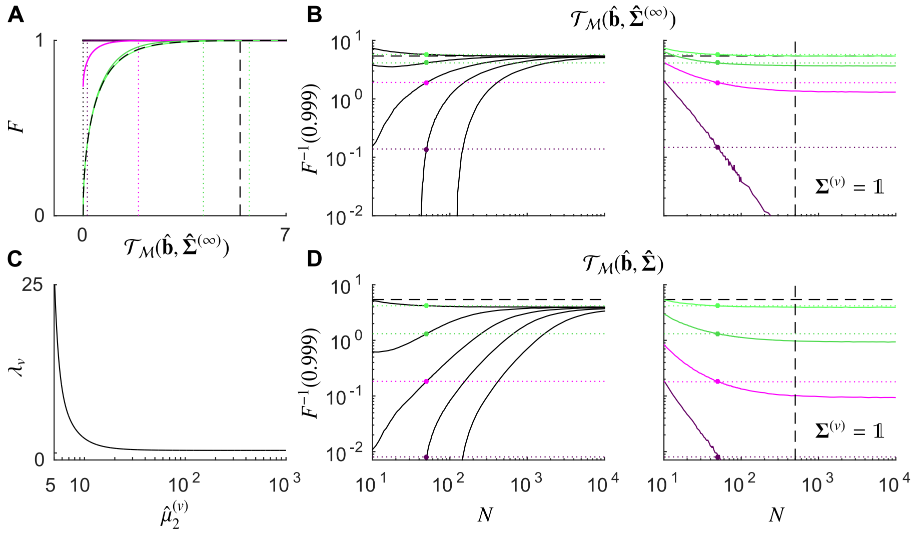


Figure S5: Convergence properties of the test statistics $\mathcal{T}_{\mathcal{M}}(\hat{\mathbf{b}}, \hat{\mathbf{\Sigma}}^{(\infty)})$ and $\mathcal{T}_{\mathcal{M}}(\hat{\mathbf{b}}, \hat{\mathbf{\Sigma}})$ for $\theta_{\hat{\mu}} = 5$, $\hat{\sigma}_0 \to \infty$, $d^{(\mathcal{X}_0)} = 2$, i.i.d. $\hat{\Phi}_n^{(v)} \sim \mathcal{N}\left(\mathbf{b}^{(v)}, \mathbf{\Sigma}'^{(v)}\right)$, $n \in \{1, ..., N\}$, $b_1^{(v)} = 10^2$, $b_2^{(v)} \in \{10^{0.3}, 10^{0.5}, 10^{0.7}, 10^1, 10^2\}$ and $\mathbf{\Sigma}'^{(v)} = \left(\begin{smallmatrix} 50 & 0 \\ 0 & 50 \end{smallmatrix}\right)$, $v \in \{1, 2\}$. These statistics correspond to the contributions of each of the two concise index sets to the test statistics of the modularizations $\mathcal{M}_1$,...,$\mathcal{M}_6$ shown in Fig. S3C. **A:** Cumulative distribution function $F$ of $\mathcal{T}_{\mathcal{M}}(\hat{\mathbf{b}}, \hat{\mathbf{\Sigma}}^{(\infty)})$ for $N = 50$ (solid lines). The color code for $\mathbf{b}^{(v)}$ is the same as the color code for $\boldsymbol{\mu}^{(v)}$ in Fig. S4B and C. Vertical dotted lines indicate the 99.9th percentiles of the cumulative distribution functions (same color code). The dashed curve and dashed vertical line correspond to the cumulative distribution function of the gamma distribution $\Gamma(\zeta_{\mathcal{M}}, 1)$ for $\zeta_{\mathcal{M}} = 0.5$ and its 99.9th percentile, respectively. **B:** (left) Dependence of the 99.9th percentiles of the cumulative distribution functions of $\mathcal{T}_{\mathcal{M}}(\hat{\mathbf{b}}, \hat{\mathbf{\Sigma}}^{(\infty)})$ on $N$. Colored dots and horizontal dotted lines indicate the 99.9th percentiles of the cumulative distribution functions in panel A (same color code). The dashed horizontal line indicates the 99.9th percentile of the gamma distribution $\Gamma(\zeta_{\mathcal{M}}, 1)$ for $\zeta_{\mathcal{M}} = 0.5$. (right) Same as left panel for $\mathbf{\Sigma}'^{(v)} = \left(\begin{smallmatrix} N & 0 \\ 0 & N \end{smallmatrix}\right)$, $v \in \{1, 2\}$. The distributions of $(\hat{\mathbf{b}}^{(v)} - \mathbf{b}^{(v)})$, $v \in \{1, 2\}$ are independent of $N$ and only the estimates $\hat{\mathbf{\Sigma}}^{(v)} = N^{-1}\hat{\mathbf{\Sigma}}'^{(v)}$, $v \in \{1, 2\}$ become more precise for larger $N$, i.e., $N^{-1}\hat{\mathbf{\Sigma}}'^{(v)} \xrightarrow{p} N^{-1}\mathbf{\Sigma}'^{(v)}$ as $N \to \infty$. The distributions of $\hat{b}_v$ converge to the distributions shown in Fig. S4A (same color code). The dashed vertical line indicates $N = 500$. **C:** $\lambda_v$ as a function of $\hat{\mu}_2^{(v)}$ for $\hat{\mu}_2^{(v)} \geq \theta_{\hat{\mu}}$ and $\theta_{\hat{\mu}} = 5$ (see Table 2 for parameters). **D:** Same as panel B for $\mathcal{T}_{\mathcal{M}}(\hat{\mathbf{b}}, \hat{\mathbf{\Sigma}})$. Error bars for numerical estimates in panels A, B and D are not shown.

$\mathcal{I}$ such that the modified test statistic conditioned on $\boldsymbol{\rho}^{(v)} \in \mathcal{I}$ (see Equ. 90 and 91) is more conservative than the gamma distribution $\Gamma(\zeta_{\mathcal{M}}, 1)$.[35] Finally, for $\boldsymbol{\rho}^{(v)}$ outside of $\mathcal{I}$, we assume that the null hypothesis is always (incorrectly) rejected and, thereby, obtain an upper bound for the $p$-value of the test statistic.

For the first step, we rescale $\hat{\Sigma}_{vv}^{(\infty)}$ (Equ. 102) by a function (see Fig. S5C)

$$\lambda_v\left(\hat{\mu}_2^{(v)}(\hat{\mathbf{\Sigma}}^{(v)})\right) = \begin{cases} 1, & \left|\hat{\mu}_2^{(v)}(\hat{\mathbf{\Sigma}}^{(v)})\right| < \theta_{\hat{\mu}} \\ \xi_0\left(1 + \sum_{n=1}^{5} \xi_n \left(\frac{\hat{\mu}_2^{(v)}(\hat{\mathbf{\Sigma}}^{(v)})}{6}\right)^{-2n}\right), & \text{otherwise,} \end{cases} \tag{113}$$

where the parameters $\xi_n \in \mathbb{R}$ are chosen such that $\xi_0 \geq 1$ and $\lambda_v \geq 1$, and define the

[35]The heavy tail of $\mathcal{T}_{\mathcal{M}}$ for finite $N$ prevents a similar approach for the entire sample space.

rescaled sample covariance matrix of $\hat{\mathbf{b}}$

$$\hat{\Sigma}_{v\omega}(\hat{\mathbf{\Sigma}}^{(v)}) = \begin{cases} \lambda_v\left(\hat{\mu}_2^{(v)}(\hat{\mathbf{\Sigma}}^{(v)})\right)\hat{\Sigma}_{vv}^{(\infty)}(\hat{\mathbf{\Sigma}}^{(v)}), & v = \omega \\ 0, & \text{otherwise} \end{cases} \tag{114}$$

for $v, \omega \in \{1, ..., d^{(X_0)}\}$. Asymptotically,[36]

$$\lambda_v\left(\hat{\mu}_2^{(v)}(\hat{\mathbf{\Sigma}}^{(v)})\right) \xrightarrow{p} \xi_0 \tag{115}$$

$$N\hat{\Sigma}_{vv}(\hat{\mathbf{\Sigma}}^{(v)}) \xrightarrow{p} \xi_0 N\hat{\Sigma}_{vv}^{(\infty)} \tag{116}$$

as $N \to \infty$, where the limit of $N\hat{\Sigma}_{vv}^{(\infty)}(\hat{\mathbf{\Sigma}}^{(v)})$ is given in Equ. 105. Analogous to $\hat{\Sigma}_{vv}^{(\infty)}$,[37]

$$\frac{\sqrt{N}(\hat{b}_v(\hat{\mathbf{\Sigma}}^{(v)}) - b_v)}{\sqrt{N\hat{\Sigma}_{vv}(\hat{\mathbf{\Sigma}}^{(v)})}} = \frac{\hat{\mu}_v(\hat{\mathbf{\Sigma}}^{(v)}) - \mu_v(\hat{\mathbf{\Sigma}}^{(v)})}{\sqrt{\lambda_v}\hat{\sigma}_v^{(\mathrm{T})}(\hat{\mathbf{\Sigma}}^{(v)})} \xrightarrow{d} \mathcal{N}(0, \xi_0^{-1}) \tag{117}$$

as $N \to \infty$ and[38]

$$\begin{aligned} \mathcal{T}_{\mathcal{M}}\left(\hat{\mathbf{b}}, \hat{\mathbf{\Sigma}}\right) &= \mathcal{T}_{\mathcal{M}}\left(\sqrt{N}\hat{\mathbf{b}}, N\hat{\mathbf{\Sigma}}\right) & (118)\\ &\xrightarrow{d} \mathcal{T}_{\mathcal{M}}\left(\sqrt{N}\hat{\mathbf{b}}, \xi_0\hat{\mathbf{\Sigma}}'^{(\infty)}\right) & (119)\\ &= \xi_0^{-1}\mathcal{T}_{\mathcal{M}}\left(\sqrt{N}\hat{\mathbf{b}}, N\hat{\mathbf{\Sigma}}^{(\infty)}\right) & (120)\\ &= \xi_0^{-1}\mathcal{T}_{\mathcal{M}}\left(\sqrt{N}(\hat{\mathbf{b}} - \mathbf{b}), N\hat{\mathbf{\Sigma}}^{(\infty)}\right) & (121)\\ &\xrightarrow{d} \Gamma(\zeta_{\mathcal{M}}, \xi_0^{-1}) & (122) \end{aligned}$$

as $N \to \infty$. Therefore, the test statistic $\mathcal{T}_{\mathcal{M}}(\hat{\mathbf{b}}, \hat{\mathbf{\Sigma}})$ is asymptotically more conservative than $\Gamma(\zeta_{\mathcal{M}}, 1)$, i.e., extreme values occur less likely for $\xi_0 \geq 1$ and $\mathcal{M} \neq \mathcal{M}_0$ under the two null hypotheses.

Secondly, we define a finite volume

$$\begin{aligned} \mathcal{I} &= I_1 \times I_2 & (123)\\ I_1 &= [-6, 6] & (124)\\ I_2 &= [-\delta_b, \delta_b], & (125) \end{aligned}$$

where $\delta_b = \theta_{\hat{\mu}} - 1$ and $\theta_{\hat{\mu}} \in \{5, 6, 7\}$. For joint standard normal $\boldsymbol{\rho}^{(v)}(\mathbf{\Sigma}^{(v)})$ (see Equ. 90 and 91) and given nominal significance level $\alpha^{(*)}$ there always exists a $\xi_0$ such that the distribution of the test statistic $\mathcal{T}_{\mathcal{M}}(\hat{\mathbf{b}}, \hat{\mathbf{\Sigma}})$ conditioned on ($\boldsymbol{\rho}^{(v)} \in \mathcal{I}$ for all $v \in \{1, ..., d^{(X_0)}\}$) is more conservative than the gamma distribution $\Gamma(\zeta_{\mathcal{M}}, 1)$.

More precisely, if the null hypothesis is true, the probability of sampling $\mathcal{T}_{\mathcal{M}}$ at least as extreme as that which was observed is referred to as the $p$-value.[39] Let $p\text{-value}|_{\boldsymbol{\rho}^{(v)} \in \mathcal{I}}$ and $p\text{-value}|_{\boldsymbol{\rho}^{(v)} \notin \mathcal{I}}$ denote the $p$-values for $\mathcal{T}_{\mathcal{M}}$ conditioned on ($\boldsymbol{\rho}^{(v)} \in \mathcal{I}$ for all $v \in \{1, ..., d^{(X_0)}\}$) and ($\boldsymbol{\rho}^{(v)} \notin \mathcal{I}$ for some $v \in \{1, ..., d^{(X_0)}\}$), respectively. The distribution of $\mathcal{T}_{\mathcal{M}}$ conditioned on ($\boldsymbol{\rho}^{(v)} \in \mathcal{I}$ for all $v \in \{1, ..., d^{(X_0)}\}$) consists of a continuous probability density function that is non-zero for all $\mathcal{T}_{\mathcal{M}} > 0$ and a point mass with value

[36] According to Lemma 19 and the continuous mapping theorem.

[37] According to Equ. 109, Equ. 110, Lemma 16 and Lemma 17.

[38] According to the continuous mapping theorem, Equ. 116, Equ. 61, Equ. 110 and Lemma 11.

[39] In the following, $\mathcal{T}_{\mathcal{M}}$ abbreviates $\mathcal{T}_{\mathcal{M}}(\hat{\mathbf{b}}, \hat{\mathbf{\Sigma}})$.

$\mathrm{P}\left(\left|\hat{\mu}_2^{(v)}(\boldsymbol{\Sigma}^{(v)})\right| < \theta_{\hat{\mu}} \text{ for all } v \in \{1, ..., d^{(X_0)}\}\right)$ at $\mathcal{T}_{\mathcal{M}} = 0$. Because $\mathcal{I}$ is bounded, $\mathcal{T}_{\mathcal{M}}$ conditioned on ($\boldsymbol{\rho}^{(v)} \in \mathcal{I}$ for all $v \in \{1, ..., d^{(X_0)}\}$) has bounded positive support and, thus, $p\text{-value}|_{\boldsymbol{\rho}^{(v)} \in \mathcal{I}} = 0$ for finite $\mathcal{T}_{\mathcal{M}}$. The smallest $\mathcal{T}_{\mathcal{M}}$ for which $p\text{-value}|_{\boldsymbol{\rho}^{(v)} \in \mathcal{I}} = 0$ scales linearly with $\xi_0^{-1}$ (see Equ. 122).

Furthermore, let $p^{(\Gamma)}$-value denote the $p$-value if $\mathcal{T}_{\mathcal{M}}$ were distributed according to the gamma distribution $\Gamma(\zeta_{\mathcal{M}}, 1)$, i.e.,

$$p^{(\Gamma)}\text{-value} \quad = \quad 1 - \frac{\gamma\left(\zeta_{\mathcal{M}}, \mathcal{T}_{\mathcal{M}}\right)}{\Gamma(\zeta_{\mathcal{M}})}, \tag{126}$$

where $\frac{\gamma(\zeta_{\mathcal{M}}, \mathcal{T}_{\mathcal{M}})}{\Gamma(\zeta_{\mathcal{M}})}$ is the cumulative distribution function of the gamma distribution, $\Gamma(\zeta_{\mathcal{M}})$ denotes the gamma function and $\gamma\left(\zeta_{\mathcal{M}}, \mathcal{T}_{\mathcal{M}}\right)$ denotes the lower incomplete gamma function. Because the gamma distribution has infinite positive support, $p^{(\Gamma)}\text{-value} > 0$ for all $\mathcal{T}_{\mathcal{M}} \geq 0$. Thus, for given $\alpha^{(*)}$, there always exist a large enough $\xi_0$ such that

$$p\text{-value}|_{\boldsymbol{\rho}^{(v)} \in \mathcal{I}} \leq p^{(\Gamma)}\text{-value} \tag{127}$$

for all $p^{(\Gamma)}\text{-value} \leq \alpha^{(*)}$ and $\mathcal{T}_{\mathcal{M}}$ conditioned on ($\boldsymbol{\rho}^{(v)} \in \mathcal{I}$ for all $v \in \{1, ..., d^{(X_0)}\}$) is more conservative than $\Gamma(\zeta_{\mathcal{M}}, 1)$. Table 2 shows suitable parameters $\xi_n$ for $\theta_{\hat{\mu}} \in \{5, 6, 7\}$ obtained by Numerical Analysis 22 to make $\mathcal{T}_{\mathcal{M}}$ efficient. According to Numerical Analysis 23, $p\text{-value}|_{\boldsymbol{\rho}^{(v)} \in \mathcal{I}} \leq p^{(\Gamma)}\text{-value}$ for $10^{-5} < \alpha^{(*)} < 0.05$.[40]

Finally, for samples outside of $\mathcal{I}$, we assume that the null hypothesis is always rejected and, thereby, obtain an upper bound for the $p$-value of the test statistic. More precisely, the $p$-value can be obtained by the corresponding $p$-values conditioned on ($\boldsymbol{\rho}^{(v)} \in \mathcal{I}$ for all $v \in \{1, ..., d^{(X_0)}\}$) and ($\boldsymbol{\rho}^{(v)} \notin \mathcal{I}$ for some $v \in \{1, ..., d^{(X_0)}\}$), i.e.,

$$p\text{-value} = p\text{-value}|_{\boldsymbol{\rho}^{(v)} \in \mathcal{I}} \, \mathrm{P}\left(\boldsymbol{\rho}^{(v)} \in \mathcal{I} \text{ for all } v \in \{1, ..., d^{(X_0)}\}\right) \tag{128}$$

$$+ \; p\text{-value}|_{\boldsymbol{\rho}^{(v)} \notin \mathcal{I}} \, \mathrm{P}\left(\boldsymbol{\rho}^{(v)} \notin \mathcal{I} \text{ for some } v \in \{1, ..., d^{(X_0)}\}\right), \tag{129}$$

where

$$\mathrm{P}\left(\boldsymbol{\rho}^{(v)} \notin \mathcal{I} \text{ for some } v \in \{1, ..., d^{(X_0)}\}\right) \quad = \quad 1 - \operatorname{erf}\left(\frac{\delta_b}{\sqrt{2}}\right)^{d^{(X_0)}} \cdot \operatorname{erf}\left(\frac{6}{\sqrt{2}}\right)^{d^{(X_0)}} \tag{130}$$

and erf denotes the error function.

If

$$p\text{-value}|_{\boldsymbol{\rho}^{(v)} \in \mathcal{I}} \leq p^{(\Gamma)}\text{-value} \tag{131}$$

for all $p^{(\Gamma)}\text{-value} \leq \alpha^{(*)}$, then

$$p\text{-value} \leq p^{(\Gamma)}\text{-value} \cdot \mathrm{P}\left(\boldsymbol{\rho}^{(v)} \in \mathcal{I} \text{ for all } v \in \{1, ..., d^{(X_0)}\}\right) \tag{132}$$
$$+ \, 1 \cdot \mathrm{P}\left(\boldsymbol{\rho}^{(v)} \notin \mathcal{I} \text{ for some } v \in \{1, ..., d^{(X_0)}\}\right)$$

for all $p\text{-value} \leq \alpha^{(*)}$, where $p\text{-value}|_{\boldsymbol{\rho}^{(v)} \notin \mathcal{I}}$ is replaced by its maximum value and samples $\boldsymbol{\rho}^{(v)} \notin \mathcal{I}$ for some $v$ are always rejected (type I error of one). This sets a lower

[40] The value $10^{-5}$ is chosen for numerical feasibility. However, the precise value of very small $p$-values is little informative, because, in general, sample distributions deviate at the far tails from their assumed asymptotic form.

bound, denoted by $\alpha^{(\min)}$, for possible nominal significance levels that depends on $\theta_{\hat{\mu}}$ and $d^{(X_0)}$ (see Table 3) such that

$$\alpha^{(*)} > \alpha^{(\min)} = \mathrm{P}\left(\boldsymbol{\rho}^{(v)} \notin \mathcal{I} \text{ for some } v \in \{1, ..., d^{(X_0)}\}\right). \tag{133}$$

For given $\alpha^{(*)}$,[41] $p$-value $\leq \alpha^{(*)}$ if

$$p^{(\Gamma)}\text{-value} \leq \alpha^{(\Gamma)} \tag{134}$$

$$\alpha^{(\Gamma)} = \frac{\alpha^{(*)} - \alpha^{(\min)}}{1 - \alpha^{(\min)}}, \tag{135}$$

where $\alpha^{(\Gamma)}$ denotes the corrected significance level.

**Correction for multiple testing**

If multiple modularizations $\mathcal{M}_i, i \in \{1, ..., d^{(\mathcal{H})}\}, d^{(\mathcal{H})} \in \mathbb{N}$ are tested on the same dataset, the random variables $\boldsymbol{\rho}^{(v)}$ and the condition ($\boldsymbol{\rho}^{(v)} \in \mathcal{I}$ for all $v \in \{1, ..., d^{(X_0)}\}$) are identical for all tests and only the significance levels for the $p^{(\Gamma)}$-values have to be corrected for multiple testing.

Let $p_i^{(\Gamma)}$-value denote the $p^{(\Gamma)}$-value (see Equ. 126) of the test statistic for the $i$-th modularization and let $\alpha_{d^{(\mathcal{H})}}^{(\Gamma)}$ denote the Bonferroni corrected significance level $\alpha^{(\Gamma)}$ (see Equ. 135), i.e.,

$$\alpha_{d^{(\mathcal{H})}}^{(\Gamma)} = \frac{\alpha^{(\Gamma)}}{d^{(\mathcal{H})}}. \tag{136}$$

Then, the overall probability of rejecting at least one true modularization out of $d^{(\text{true})}$ modularizations at significance level $\alpha_{d^{(\mathcal{H})}}^{(\Gamma)}$ is bounded by

$$\mathrm{P}\left(\bigcup_{i=1}^{d^{(\text{true})}} \left(p_i^{(\Gamma)}\text{-value} < \alpha_{d^{(\mathcal{H})}}^{(\Gamma)}\right)\right) \leq \alpha^{(*)}, \tag{137}$$

where the minimum value of $\alpha^{(*)}$ (according to Table 3) remains unchanged.

More precisely, analogous to the single hypothesis case (see Equ. 132),

$$\begin{aligned}\mathrm{P}\left(\bigcup_{i=1}^{d^{(\text{true})}} \left(p_i^{(\Gamma)}\text{-value} < \alpha_{d^{(\mathcal{H})}}^{(\Gamma)}\right)\right) \leq\ & \mathrm{P}\left(\bigcup_{i=1}^{d^{(\text{true})}} \left(p_i^{(\Gamma)}\text{-value} < \alpha_{d^{(\mathcal{H})}}^{(\Gamma)}\right) \middle| \boldsymbol{\rho}^{(v)} \in \mathcal{I} \text{ for all } v\right) \cdot \\ & \mathrm{P}\left(\boldsymbol{\rho}^{(v)} \in \mathcal{I} \text{ for all } v \in \{1, ..., d^{(X_0)}\}\right) + \\ & 1 \cdot \mathrm{P}\left(\boldsymbol{\rho}^{(v)} \notin \mathcal{I} \text{ for some } v \in \{1, ..., d^{(X_0)}\}\right),\end{aligned} \tag{138}$$

where we again assumed the worst case scenario for ($\boldsymbol{\rho}^{(v)} \notin \mathcal{I}$ for some $v \in \{1, ..., d^{(X_0)}\}$). If the $p_i$-value for $\mathcal{T}_{\mathcal{M}}$ conditioned on ($\boldsymbol{\rho}^{(v)} \in \mathcal{I}$ for some $v \in \{1, ..., d^{(X_0)}\}$), denoted by $p_i\text{-value}|_{\boldsymbol{\rho}^{(v)} \in \mathcal{I}}$, is bounded by $p_i^{(\Gamma)}$-value, i.e.,

$$p_i\text{-value}|_{\boldsymbol{\rho}^{(v)} \in \mathcal{I}} \leq p_i^{(\Gamma)}\text{-value} \tag{139}$$

[41] According to Numerical Analysis 22 and 23, Equ. 131 is estimated to be true for $10^{-5}\alpha^{(*)} < 0.05$.

for all $p_i^{(\Gamma)}$-value $\leq \alpha^{(*)}$, then,[42]

$$P\left(\bigcup_{i=1}^{d^{(\text{true})}} \left(p_i^{(\Gamma)}\text{-value} < \alpha^{(\Gamma)}_{d^{(\mathcal{H})}}\right) \middle| \boldsymbol{\rho}^{(v)} \in \mathcal{I} \text{ for all } v\right) \leq \tag{140}$$

$$P\left(\bigcup_{i=1}^{d^{(\text{true})}} \left(p_i\text{-value}|_{\boldsymbol{\rho}^{(v)}\in\mathcal{I}} < \alpha^{(\Gamma)}_{d^{(\mathcal{H})}}\right) \middle| \boldsymbol{\rho}^{(v)} \in \mathcal{I} \text{ for all } v\right)$$

$$\leq \sum_{i=1}^{d^{(\text{true})}} P\left( p_i\text{-value}|_{\boldsymbol{\rho}^{(v)}\in\mathcal{I}} < \alpha^{(\Gamma)}_{d^{(\mathcal{H})}} \middle| \boldsymbol{\rho}^{(v)} \in \mathcal{I} \text{ for all } v\right) \tag{141}$$

$$= d^{(\text{true})}\alpha^{(\Gamma)}_{d^{(\mathcal{H})}} = d^{(\text{true})}\frac{\alpha^{(\Gamma)}}{d^{(\mathcal{H})}} \leq \alpha^{(\Gamma)} \tag{142}$$

and[43]

$$P\left(\bigcup_{i=1}^{d^{(\text{true})}} \left(p_i^{(\Gamma)}\text{-value} < \alpha^{(\Gamma)}_{d^{(\mathcal{H})}}\right)\right) \leq \alpha^{(*)}. \tag{143}$$

For example, to test $d^{(\mathcal{H})} = 26$ modularizations for $d = 5$ observable components ($d^{(\mathcal{X}_0)} = 6$) and a significance level of $\alpha^{(*)} = 0.01$ we choose a cutoff $\theta_{\hat{\mu}} = 5$ resulting in $\alpha^{(\min)} = 3.8 \cdot 10^{-4}$ (see Table 3), $\alpha^{(\Gamma)} = 9.6 \cdot 10^{-3}$ and a Bonferroni corrected individual significance level $\alpha^{(\Gamma)}_{d^{(\mathcal{H})}} = 3.7 \cdot 10^{-4}$.

**Simple asymptotic test assuming arbitrarily correlated sample moment ratios**

For limited data, $\hat{b}_v$ for different $v$ will be estimated from the same samples and, in general, correlated. We define the full covariance matrices

$$\hat{\Sigma}_{v\omega}(\hat{\boldsymbol{\Sigma}}^{(v,\omega)}) = \sqrt{\lambda_v\left(\varrho^{(v)}\right)\lambda_\omega\left(\varrho^{(\omega)}\right)}\hat{\Sigma}^{(\infty)}_{v\omega}(\hat{\boldsymbol{\Sigma}}^{(v,\omega)}) \tag{144}$$

$$\hat{\Sigma}^{(\infty)}_{v\omega}(\hat{\boldsymbol{\Sigma}}^{(v,\omega)}) = \begin{cases} \hat{\sigma}_0\Theta_3(\hat{\boldsymbol{\Sigma}}^{(v,v)})\Theta_3(\hat{\boldsymbol{\Sigma}}^{(\omega,\omega)}), & (v = \omega) \wedge \left(\left|\varrho^{(v)}\right| < \theta_{\hat{\mu}}\right) \\ 0, & (v \neq \omega) \wedge \left(\left(\left|\varrho^{(v)}\right| < \theta_{\hat{\mu}}\right) \vee \left(\left|\varrho^{(\omega)}\right| < \theta_{\hat{\mu}}\right)\right) \\ \frac{1}{\hat{b}_2^{(v)}\hat{b}_2^{(\omega)}}\left(\hat{\Sigma}^{(v,w)}_{11} - \frac{\hat{b}_1^{(\omega)}}{\hat{b}_2^{(\omega)}}\hat{\Sigma}^{(v,w)}_{12} - \frac{\hat{b}_1^{(v)}}{\hat{b}_2^{(v)}}\hat{\Sigma}^{(v,w)}_{21} + \frac{\hat{b}_1^{(v)}\hat{b}_1^{(\omega)}}{\hat{b}_2^{(v)}\hat{b}_2^{(\omega)}}\hat{\Sigma}^{(v,w)}_{22}\right), & \text{otherwise} \end{cases} \tag{145}$$

$$\varrho^{(v)} = \frac{\hat{b}_2^{(v)}}{\sqrt{\hat{\Sigma}^{(v,v)}_{22}}} \tag{146}$$

for $v, \omega \in \{1, ..., d^{(\mathcal{X}_0)}\}$ and constant $\hat{\sigma}_0 > 0$.[44] These definitions differ from the previous definitions (Equ. 102 and 114) only for off-diagonal terms, i.e., for $v \neq \omega$.

Asymptotically,[45]

$$\sqrt{N}(\hat{\mathbf{b}} - \mathbf{b}) \xrightarrow{d} \mathcal{N}(\mathbf{0}, \boldsymbol{\Sigma}') \tag{147}$$

$$N\hat{\boldsymbol{\Sigma}}^{(\infty)} \xrightarrow{p} \boldsymbol{\Sigma}' \tag{148}$$

$$N\hat{\boldsymbol{\Sigma}} \xrightarrow{p} \xi_0\boldsymbol{\Sigma}' \tag{149}$$

[42] Equ. 141 follows from Boole's inequality. Equ. 142 follows from the uniform distribution of $p$-values.

[43] According to Equ. 142, Equ. 138 and Equ. 135.

[44] $\hat{\sigma}_0$ is defined in Equ. 103 and $\hat{\boldsymbol{\Sigma}}^{(v,\omega)} = \hat{\boldsymbol{\Sigma}}'^{(v,\omega)}/N$ (see Equ. 69 and 84).

[45] According to Lemma 20.

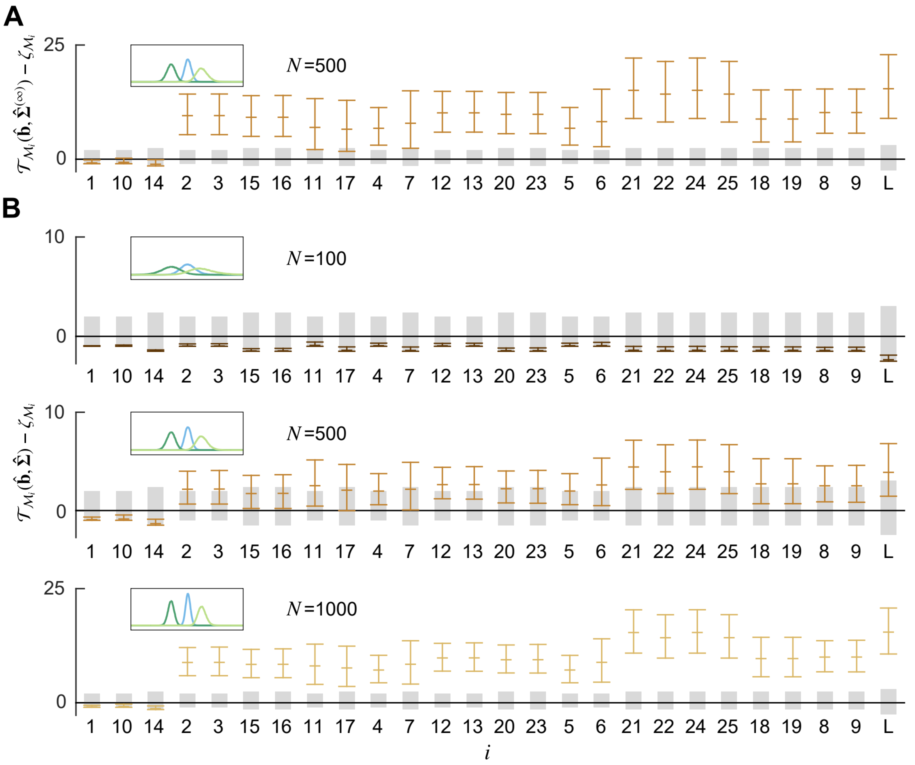


Figure S6: Asymptotic test assuming uncorrelated sample moment ratios for the modularizations shown in Fig. S3C and samples generated by the toy model in Section 2.5 that implements $\mathcal{M}_{14}$. **A:** Distribution of the statistic $\mathcal{T}_{\mathcal{M}}(\hat{\mathbf{b}}, \hat{\boldsymbol{\Sigma}}^{(\infty)})$. **B:** Distribution of the statistic $\mathcal{T}_{\mathcal{M}}(\hat{\mathbf{b}}, \hat{\boldsymbol{\Sigma}})$. Both statistics incorrectly assume uncorrelated $\hat{\mathbf{b}}$. Cutoff $\theta_{\hat{\mu}} = 5$ and $\hat{\sigma}_0 \to \infty$. Gray bars indicate the 0th and 95th percentiles of the gamma distribution $\Gamma(\zeta_{\mathcal{M}_i}, 1)$ shifted by $-\zeta_{\mathcal{M}_i}$. Colored bars indicate the 2.5th percentile, the mean and the 97.5th percentile of the distributions of the test statistics. Insets: Distributions of the elements of $\hat{\mathbf{b}}$ (color code as in Fig. S3A).

as $N \to \infty$, where $\boldsymbol{\Sigma}'$ has full rank and

$$\Sigma'_{v\omega} = \frac{1}{b_2^{(v)} b_2^{(\omega)}} \left( \Sigma'^{(v,w)}_{11} - \frac{b_1^{(\omega)}}{b_2^{(\omega)}} \Sigma'^{(v,w)}_{12} - \frac{b_1^{(v)}}{b_2^{(v)}} \Sigma'^{(v,w)}_{21} + \frac{b_1^{(v)} b_1^{(\omega)}}{b_2^{(v)} b_2^{(\omega)}} \Sigma'^{(v,w)}_{22} \right) \quad (150)$$

$$\Sigma'^{(v,\omega)}_{ij} = E\left[\rho'^{(v)}_i \rho'^{(\omega)}_j\right] \quad (151)$$

for $v, \omega \in \{1, ..., d^{(X_0)}\}$ and $i, j \in \{1, 2\}$.

For the toy example below, the off-diagonal terms of $\hat{\boldsymbol{\Sigma}}$ are positive and result in a more conservative test even without any corrections (see correct modularizations $\mathcal{M}_1$, $\mathcal{M}_{10}$ and $\mathcal{M}_{14}$ in Fig. S6). Asymptotically, exclusively positive correlations of moment ratios within the same concise index set always result in a more conservative test due to the invariance of the statistic to joint fluctuations[46] and, thus, an overestimation of the effective covariance of $\hat{\mathbf{b}}$. Numerical Analysis 23 corroborates that this holds also true for sufficiently many samples such that the moment estimates are approximately joint normal and $N\hat{\boldsymbol{\Sigma}} \approx N\boldsymbol{\Sigma}$.

[46] See Lemma 11

In order to correct for arbitrary correlations between sample moment ratios, we rescale the covariance matrix $\hat{\mathbf{\Sigma}}$ by a multiplicative factor $\lambda^{(\max)} \geq 1$ such that the distribution of $\mathcal{T}_{\mathcal{M}}(\hat{\mathbf{b}}, \lambda^{(\max)}\hat{\mathbf{\Sigma}})$ for correlated $\hat{\mathbf{b}}$ is asymptotically more conservative than the distribution of $\mathcal{T}_{\mathcal{M}}(\hat{\mathbf{b}}, \hat{\mathbf{\Sigma}})$ for uncorrelated $\hat{\mathbf{b}}$, which is $\Gamma(\zeta_{\mathcal{M}}, \xi_0^{-1})$. Because the test depends only on the norm $\sqrt{N}|\hat{\mathbf{b}} - \mathbf{b}|$ and $\sqrt{N}(\hat{\mathbf{b}} - \mathbf{b}) \xrightarrow{d} \mathcal{N}(\mathbf{0}, \mathbf{\Sigma}')$ for $N \to \infty$ (see Equ. 147), the smallest possible $\lambda^{(\max)}$ is the largest eigenvalue of $\mathbf{\Sigma}'$ normalized to unit diagonal. This matrix can be estimated by

$$\grave{\mathbf{\Sigma}}\hat{\mathbf{\Sigma}}\grave{\mathbf{\Sigma}}, \tag{152}$$

where the matrix $\grave{\mathbf{\Sigma}}$ has the same size as $\hat{\mathbf{\Sigma}}$, is diagonal and $\grave{\Sigma}_{vv} = \hat{\Sigma}_{vv}^{-1/2}$ if $v$ is an element of an index set of size larger than one and $\grave{\Sigma}_{vv} = 0$, otherwise.[47] As sample eigenvalues and eigenvectors are consistent estimators of the corresponding population eigenvalues and eigenvectors (Anderson, 2003), $\mathcal{T}_{\mathcal{M}}(\hat{\mathbf{b}}, \lambda^{(\max)}\hat{\mathbf{\Sigma}})$ is asymptotically more conservative than $\Gamma(\zeta_{\mathcal{M}}, \xi_0^{-1})$. If $\lambda^{(\max)} = 1$, $\mathcal{T}_{\mathcal{M}}(\hat{\mathbf{b}}, \hat{\mathbf{\Sigma}}) \sim \Gamma(\zeta_{\mathcal{M}}, \xi_0^{-1})$. Numerical Analysis 23 corroborates that this holds also true for sufficiently many samples such that the moment estimates are approximately joint normal and $N\hat{\mathbf{\Sigma}} \approx N\mathbf{\Sigma}$. Results for the toy example below are shown in Fig. S7.

The precise identification of nested modularizations in Chapter 2.2 is based on the assumption that $s_d$ is correlated with all other observable components, so that the denominator of $b_v$ is non-zero, i.e., $b_2^{(v)} \neq 0$ for all $v \in \{1, ..., d^{(X_0)}\}$. In contrast, all tests derived in this chapter are still valid if any $b_2^{(v)} = 0$ as long as the corrected significance levels $\alpha^{(\Gamma)}$ are used (see Equ. 135) and the estimates $\hat{b}_2^{(v)}$, $\hat{\Sigma}'^{(v)}_{11}$ and $\hat{\Sigma}'^{(v)}_{22}$ are non-zero, which is almost surely the case. For $b_2^{(v)} = 0$, the probability that $\hat{b}_v$ is clamped to $b_v$ is large enough such that the test statistics remain conservative.[48]

Furthermore, a finite mean of $\hat{b}_v$ is not required but convenient to summarize the data. In particular, for $b_2^{(v)} \neq 0$, $\theta_{\hat{\mu}} > 0$ and joint normal $\hat{\Phi}_{n1}^{(v)}$ and $\hat{\Phi}_{n2}^{(v)}$, the mean of $\hat{b}_v$ is finite.[49] Finally, the clamping of $\hat{b}_v$ to the unknown $b_v$ if $\left(\left|\varrho^{(v)}\right| < \theta_{\hat{\mu}}\right)$ (see Equ.98) can be implemented by taking the limit $\hat{\sigma}_0 \to \infty$.[50]

In summary, the only assumption required for $\mathcal{T}_{\mathcal{M}}(\hat{\mathbf{b}}, \lambda^{(\max)}\hat{\mathbf{\Sigma}})$ to be asymptotically more conservative than the gamma distribution $\Gamma(\zeta_{\mathcal{M}}, \xi_0^{-1})$, in addition to the definitions in Chapter 1, is an invertible covariance matrix[51]

$$\mathbf{\Sigma}'^{(\mathrm{all})} = \begin{pmatrix} \mathbf{\Sigma}'^{(1,1)} & \cdots & \mathbf{\Sigma}'^{(1,d^{(X_0)})} \\ \vdots & \ddots & \vdots \\ \mathbf{\Sigma}'^{(d^{(X_0)},1)} & \cdots & \mathbf{\Sigma}'^{(d^{(X_0)},d^{(X_0)})} \end{pmatrix}. \tag{153}$$

For uncorrelated $\hat{b}_v$ and, thus, diagonal $\hat{\mathbf{\Sigma}}$ and $\lambda^{(\max)} = 1$, $\mathcal{T}_{\mathcal{M}}(\hat{\mathbf{b}}, \hat{\mathbf{\Sigma}}) \xrightarrow{d} \Gamma(\zeta_{\mathcal{M}}, \xi_0^{-1})$

[47] $\mathcal{T}_{\mathcal{M}}$ is independent of sample moment ratios associated with concise index sets of size one.

[48] For $b_2^{(v)} = 0$, all $\hat{b}_v$ within $\mathcal{I}$ are clamped to $b_v$.

[49] For normal i.i.d. samples, $\hat{b}_2^{(v)}$ and $\hat{\Sigma}_{22}^{(v,v)}$ are independent, $\hat{b}_1^{(v)}$ and $\hat{\Sigma}_{22}^{(v,v)}$ are independent and $\frac{\hat{b}_1^{(v)}}{\sqrt{\hat{\Sigma}_{22}^{(v,v)}}}$ has a non-central $t$-distribution with finite mean. From $|\hat{b}_v| = \left|\frac{\hat{b}_1^{(v)}}{\hat{b}_2^{(v)}}\right| \leq \left|\frac{\hat{b}_1^{(v)}}{\theta_{\hat{\mu}}\sqrt{\hat{\Sigma}_{22}^{(v,v)}}}\right|$ follows $\hat{b}_v$ has finite mean.

[50] More precisely, the limits $N \to \infty$ and $\hat{\sigma}_0 \to \infty$ can be exchanged. This is because all test statistics remain finite as $\hat{\sigma}_0 \to \infty$, $\hat{\sigma}_0$ has only an impact on the test statistics if $\left(\left|\varrho^{(v)}\right| < \theta_{\hat{\mu}}\right)$ and $\mathrm{P}(\left(\left|\varrho^{(v)}\right| < \theta_{\hat{\mu}}\right)$ is independent of $\hat{\sigma}_0$. Because $\lim_{N\to\infty} \mathrm{P}(\left(\left|\varrho^{(v)}\right| < \theta_{\hat{\mu}}\right) = 0$, all test statistics converge in probability to the corresponding statistics for $\left(\left|\varrho^{(v)}\right| \geq \theta_{\hat{\mu}}\right)$ after taking the limit $\hat{\sigma}_0 \to \infty$.

[51] Required for Lemma 20.

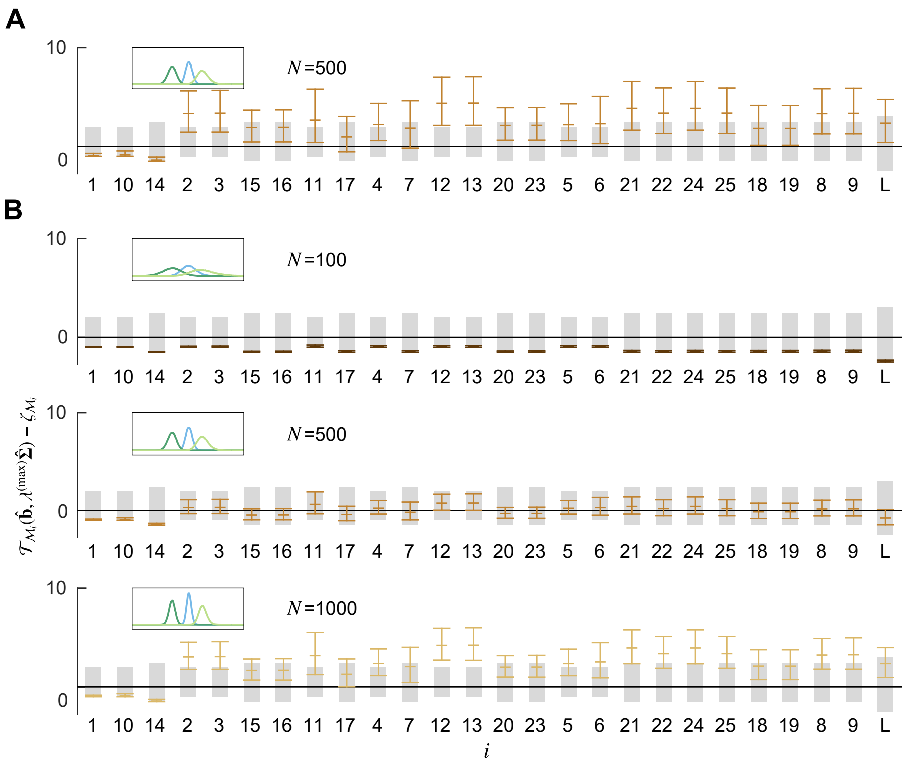


Figure S7: Simple asymptotic test assuming arbitrarily correlated sample moment ratios for the modularizations shown in Fig. S3C and samples generated by the toy model in Section 2.5 that implements $\mathcal{M}_{14}$. **A:** Distribution of the statistic $\mathcal{T}_{\mathcal{M}}(\hat{\mathbf{b}}, \lambda^{(\max)}\hat{\boldsymbol{\Sigma}})$ for $\lambda_v = 1$, $v \in \{1, ..., d^{(X_0)}\}$, so that $\hat{\boldsymbol{\Sigma}} = \hat{\boldsymbol{\Sigma}}^{(\infty)}$. **B:** Distribution of the statistic $\mathcal{T}_{\mathcal{M}}(\hat{\mathbf{b}}, \lambda^{(\max)}\hat{\boldsymbol{\Sigma}})$ for $\lambda_v$, $v \in \{1, ..., d^{(X_0)}\}$ chosen according to Table 2 (see Equ. 113). Cutoff $\theta_{\hat{\mu}} = 5$ and $\hat{\sigma}_0 \to \infty$. Gray bars indicate the 0th and 95th percentiles of the gamma distribution $\Gamma(\zeta_{\mathcal{M}_i}, 1)$ shifted by $-\zeta_{\mathcal{M}_i}$. Colored bars indicate the 2.5th percentile, the mean and the 97.5th percentile of the distributions of the test statistics. Insets: Distributions of the elements of $\hat{\mathbf{b}}$ (color code as in Fig. S3A).

as $N \to \infty$. For sufficiently large $N$ such that the estimates of the numerator and denominator of all $\hat{b}_v$ are joint normal and $\hat{\boldsymbol{\Sigma}}'^{(\text{all})} \approx \boldsymbol{\Sigma}'^{(\text{all})}$, we consolidate numerically that $\mathcal{T}_{\mathcal{M}}(\hat{\mathbf{b}}, \lambda^{(\max)}\hat{\boldsymbol{\Sigma}})$ is more conservative than the gamma distribution $\Gamma(\zeta_{\mathcal{M}}, 1)$ even for finite $N$.

**Advanced asymptotic test assuming arbitrarily correlated sample moment ratios**

The impact of dependencies between observables within functional modules on the asymptotic test can be taken into account precisely by simply decorrelating the corresponding sample moment ratios. However, the sample moment ratios related to different concise index sets are in general also correlated. Moreover, decorrelation is not possible without violating the independence assumption of modules, i.e., observables inside and outside of modules have to be independent when conditioned on the interface variable. As before, we correct for these intra-module correlations by introducing a multiplicative scaling factor $\lambda^{(\max)}$ that is large enough such that the test statistic applied to the rescaled covariance matrix of the partially decorrelated sample moment ratios is asymptotically more conservative than $\Gamma(\zeta_{\mathcal{M}}, \xi_0^{-1})$.

More precisely, for each concise index set $X_c^{(\mathcal{M})}, c \in \{1, ..., d^{(\mathcal{X})}\}$, let $\hat{\mathbf{b}}[X_c^{(\mathcal{M})}] \in \mathbb{R}^{|X_c^{(\mathcal{M})}|}$ denote the sub-vector $\hat{b}_k$, $k \in X_c^{(\mathcal{M})}$. For each pair $c_1, c_2 \in \{1, ..., d^{(\mathcal{X})}\}$, let $\hat{\mathbf{\Sigma}}[X_{c_1}^{(\mathcal{M})}, X_{c_2}^{(\mathcal{M})}] \in \mathbb{R}^{|X_{c_1}^{(\mathcal{M})}| \times |X_{c_2}^{(\mathcal{M})}|}$ denote the sub-matrix $\hat{\Sigma}_{k_1 k_2}$, $k_1 \in X_{c_1}^{(\mathcal{M})}$, $k_2 \in X_{c_2}^{(\mathcal{M})}$. Furthermore, let

$$\hat{\mathbf{\Sigma}}[X_c^{(\mathcal{M})}, X_c^{(\mathcal{M})}] = \mathbf{Q}^{(c)} \mathbf{\Lambda}^{(c)} \mathbf{Q}^{(c)^T} \tag{154}$$

denote the eigendecomposition of $\hat{\mathbf{\Sigma}}[X_c^{(\mathcal{M})}, X_c^{(\mathcal{M})}]$, where the columns of $\mathbf{Q}^{(c)}$ are the eigenvectors and the elements on the diagonal of $\mathbf{\Lambda}^{(c)}$ are the eigenvalues of $\hat{\mathbf{\Sigma}}[X_c^{(\mathcal{M})}, X_c^{(\mathcal{M})}]$.

We define

$$\acute{\mathbf{b}} = \begin{pmatrix} \acute{\mathbf{b}}[X_1^{(\mathcal{M})}] \\ \vdots \\ \acute{\mathbf{b}}[X_{d^{(\mathcal{X})}}^{(\mathcal{M})}] \end{pmatrix} \qquad \acute{\mathbf{\Sigma}} = \begin{pmatrix} \acute{\mathbf{\Sigma}}[X_1^{(\mathcal{M})}, X_1^{(\mathcal{M})}] & \dots & \acute{\mathbf{\Sigma}}[X_1^{(\mathcal{M})}, X_{d^{(\mathcal{X})}}^{(\mathcal{M})}] \\ \vdots & \ddots & \vdots \\ \acute{\mathbf{\Sigma}}[X_{d^{(\mathcal{X})}}^{(\mathcal{M})}, X_1^{(\mathcal{M})}] & \dots & \acute{\mathbf{\Sigma}}[X_{d^{(\mathcal{X})}}^{(\mathcal{M})}, X_{d^{(\mathcal{X})}}^{(\mathcal{M})}] \end{pmatrix}, \tag{155}$$

where

$$\acute{\mathbf{b}}[X_c^{(\mathcal{M})}] = \mathbf{R}^{(c)^{-1}} \mathbf{Q}^{(c)^T} \hat{\mathbf{b}}[X_c^{(\mathcal{M})}] \tag{156}$$

$$\acute{\mathbf{\Sigma}}[X_{c_1}^{(\mathcal{M})}, X_{c_2}^{(\mathcal{M})}] = \begin{cases} \mathbf{R}^{(c_1)^{-2}} \mathbf{\Lambda}^{(c_1)}, & c_1 = c_2 \\ \mathbf{0} \in \mathbb{R}^{|X_{c_1}^{(\mathcal{M})}| \times |X_{c_2}^{(\mathcal{M})}|}, & \text{otherwise} \end{cases} \tag{157}$$

$$R_{k_1 k_2}^{(c)} = \begin{cases} r_{k_1}^{(c)}, & (k_1 = k_2) \\ 0, & \text{otherwise} \end{cases} \tag{158}$$

$$r_{k_1}^{(c)} = \sum_{k_2 \in \{1, ..., |X_c^{(\mathcal{M})}|\}} Q_{k_2 k_1}^{(c)} \tag{159}$$

$k_1, k_2 \in \{1, ..., |X_c^{(\mathcal{M})}|\}$, $\acute{\mathbf{\Sigma}}$ is diagonal and we assume all $\mathbf{R}^{(c)}$ are invertible. For diagonal $\hat{\mathbf{\Sigma}}$, $\acute{\mathbf{\Sigma}} = \hat{\mathbf{\Sigma}}$ and $\acute{\mathbf{b}} = \hat{\mathbf{b}}$.

Furthermore, we define $\lambda^{(\max)}$ as the largest eigenvalue of the matrix

$$\mathbf{M} = \begin{pmatrix} \mathbf{M}[X_1^{(\mathcal{M})}, X_1^{(\mathcal{M})}] & \dots & \mathbf{M}[X_1^{(\mathcal{M})}, X_{d^{(\mathcal{X})}}^{(\mathcal{M})}] \\ \vdots & \ddots & \vdots \\ \mathbf{M}[X_{d^{(\mathcal{X})}}^{(\mathcal{M})}, X_1^{(\mathcal{M})}] & \dots & \mathbf{M}[X_{d^{(\mathcal{X})}}^{(\mathcal{M})}, X_{d^{(\mathcal{X})}}^{(\mathcal{M})}] \end{pmatrix}, \tag{160}$$

where

$$\mathbf{M}[X_{c_1}^{(\mathcal{M})}, X_{c_2}^{(\mathcal{M})}] = \begin{cases} \mathbf{\Lambda}^{(c_1)^{-\frac{1}{2}}} \mathbf{Q}^{(c_1)^T} \hat{\mathbf{\Sigma}}[X_{c_1}^{(\mathcal{M})}, X_{c_2}^{(\mathcal{M})}] \mathbf{Q}^{(c_2)} \mathbf{\Lambda}^{(c_2)^{-\frac{1}{2}}}, & \\ & (|X_{c_1}^{(\mathcal{M})}| > 1) \wedge (|X_{c_2}^{(\mathcal{M})}| > 1) \\ \mathbf{0} \in \mathbb{R}^{|X_{c_1}^{(\mathcal{M})}| \times |X_{c_2}^{(\mathcal{M})}|}, & \text{otherwise} \end{cases} \tag{161}$$

for $c_1, c_2 \in \{1, ..., d^{(\mathcal{X})}\}$. $\acute{\mathbf{b}}$, $\acute{\mathbf{\Sigma}}$ and $\lambda^{(\max)}$ depend on $\mathcal{M}$ in contrast to $\hat{\mathbf{b}}$ and $\hat{\mathbf{\Sigma}}$.

Because sample eigenvalues and eigenvectors are consistent estimators of the corresponding population eigenvalues and eigenvectors (Anderson, 2003), it follows from Theorem 21 that $\mathcal{T}_{\mathcal{M}}(\acute{\mathbf{b}}, \lambda^{(\max)} \acute{\mathbf{\Sigma}})$ is asymptotically more conservative than $\Gamma(\zeta_{\mathcal{M}}, \xi_0^{-1})$ If $\lambda^{(\max)} = 1$, $\mathcal{T}_{\mathcal{M}}(\acute{\mathbf{b}}, \acute{\mathbf{\Sigma}}) \sim \Gamma(\zeta_{\mathcal{M}}, \xi_0^{-1})$. Numerical Analysis 23 corroborates that this holds also true for sufficiently many samples such that the moment estimates are approximately joint normal and $N\hat{\mathbf{\Sigma}} \approx N\mathbf{\Sigma}$. Results for the toy example below are shown in Fig. S8.

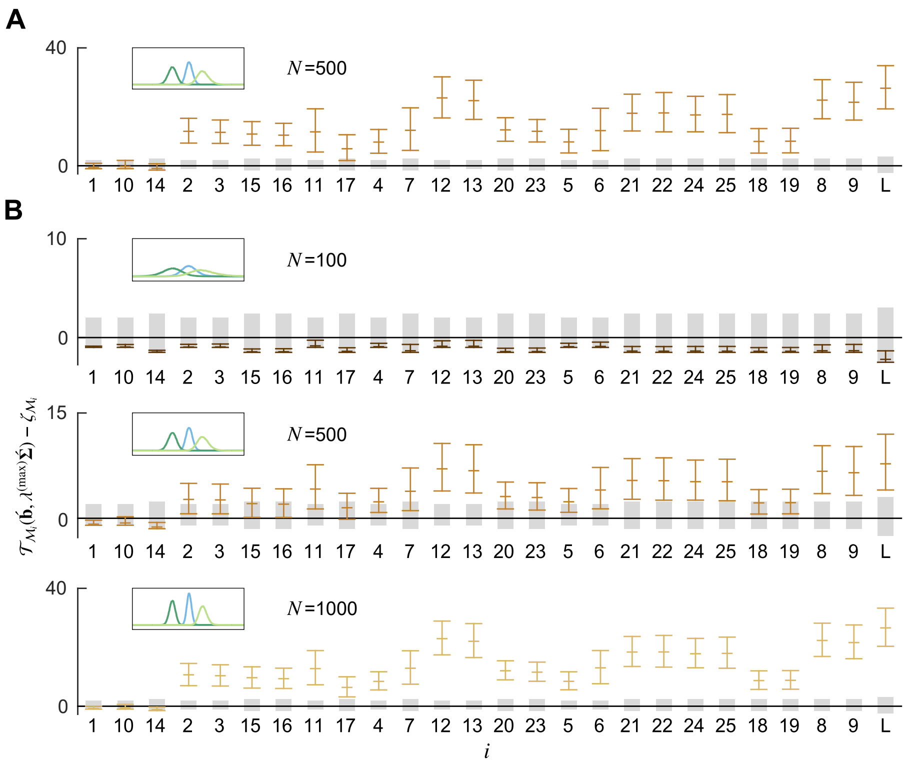


Figure S8: Advanced asymptotic test assuming arbitrarily correlated sample moment ratios for the modularizations shown in Fig. S3C and samples generated by the toy model in Section 2.5 that implements $\mathcal{M}_{14}$. **A:** Distribution of the statistic $\mathcal{T}_{\mathcal{M}}(\acute{\mathbf{b}}, \lambda^{(\max)}\acute{\mathbf{\Sigma}})$ for $\lambda_v = 1$, $v \in \{1, ..., d^{(X_0)}\}$, so that $\hat{\mathbf{\Sigma}} = \hat{\mathbf{\Sigma}}^{(\infty)}$. **B:** Distribution of the statistic $\mathcal{T}_{\mathcal{M}}(\acute{\mathbf{b}}, \lambda^{(\max)}\acute{\mathbf{\Sigma}})$ for $\lambda_v$, $v \in \{1, ..., d^{(X_0)}\}$ chosen according to Table 2 (see Equ. 113). Cutoff $\theta_{\hat{\mu}} = 5$ and $\hat{\sigma}_0 \to \infty$. Gray bars indicate the 0th and 95th percentiles of the gamma distribution $\Gamma(\zeta_{\mathcal{M}_i}, 1)$ shifted by $-\zeta_{\mathcal{M}_i}$. Colored bars indicate the 2.5th percentile, the mean and the 97.5th percentile of the distributions of the test statistics. Insets: Distributions of the elements of $\hat{\mathbf{b}}$ (color code as in Fig. S3A).

In conclusion, the only assumptions required for $\mathcal{T}_{\mathcal{M}}(\acute{\mathbf{b}}, \lambda^{(\max)}\acute{\mathbf{\Sigma}})$ to be asymptotically more conservative than the gamma distribution $\Gamma(\zeta_{\mathcal{M}}, \xi_0^{-1})$, in addition to the definitions in Chapter 1, are invertible $\mathbf{R}^{(c)}$ and an invertible covariance matrix $\mathbf{\Sigma}'^{(\mathrm{all})}$.[52] For uncorrelated $\hat{b}_v$, $\lambda^{(\max)} = 1$ and $\mathcal{T}_{\mathcal{M}}(\acute{\mathbf{b}}, \acute{\mathbf{\Sigma}}) \xrightarrow{d} \Gamma(\zeta_{\mathcal{M}}, \xi_0^{-1})$ as $N \to \infty$.

#### 2.5 Toy example

We implement the modularization shown in Fig. S2A ($\mathcal{M}_{14}$ in Fig. S3C) for $d = 5$ observable components $s_i \in \{1, 2\}, i \in \{1, ..., d-1\}$, $s_d \in \{1.2, 1.4, 2.2, 2.4\}$ and $d^{(\mathcal{M})} = 2$ modules. The binary interface variables $y^{(c)} \in \{1, 2\}$, $c \in \{1, 2\}$ are distributed according to

[52]Required for Lemma 20 and Theorem 21.

$$P(s_4 = 2) = 0.5 \tag{162}$$

$$P(s_3 = 2|s_4) = \begin{cases} 0.85, & s_4 = 2 \\ 0.15, & \text{otherwise}, \end{cases} \tag{163}$$

$$P(s_2 = 2|y^{(1)}) = \begin{cases} 0.85, & y^{(1)} = 2 \\ 0.15, & \text{otherwise}, \end{cases} \tag{164}$$

$$P(s_1 = 2|y^{(2)}) = \begin{cases} 0.85, & y^{(2)} = 2 \\ 0.15, & \text{otherwise}, \end{cases} \tag{165}$$

$$P(y^{(1)} = 2|s_3, s_4) = \begin{cases} 1, & s_3 = s_4 = 2 \\ 0, & \text{otherwise}, \end{cases} \tag{166}$$

$$P(y^{(2)} = 2|s_2, y^{(1)}) = \begin{cases} 0.85, & y^{(1)} = s_2 = 2 \\ 1, & y^{(1)} \neq s_2 \\ 0, & \text{otherwise}, \end{cases} \tag{167}$$

$$s_5 = 0.2 s_1 + y^{(2)}, \tag{168}$$

so that all elements of $\mathbf{s}$ are correlated, which increase the difficulty of the identification task. Fig. S6 shows the distribution of $\mathcal{T}_{\mathcal{M}}(\hat{\mathbf{b}}, \hat{\mathbf{\Sigma}})$ calculated for $N \in \{100, 500, 1000\}$ samples normalized to zero mean and unit variance. All elements of $\hat{\mathbf{b}}$ are positively correlated with Pearson's correlation coefficients larger than 0.4. Furthermore, $\Theta_3(\mathbf{\Sigma}^{(v)}) \in [1.18, 1.24]$ and $\Theta_4(\mathbf{\Sigma}^{(v)}) \in [-10^{-3}, 10^{-3}]$ for all $v \in \{1, ..., d^{(X_0)}\}$.

For the correct modularizations $\mathcal{M}_1$, $\mathcal{M}_{10}$ and $\mathcal{M}_{14}$, the uncorrected test statistic is more conservative than the corresponding gamma distributions even though it incorrectly assumes uncorrelated $\hat{\mathbf{b}}$. Results for the simple and the advanced asymptotic test (both assuming arbitrarily correlated components of $\hat{\mathbf{b}}$) are shown in Fig. S7 and Fig. S8, respectively. For the toy example, the correlation corrected test has larger statistical power than the simple test.

### 3 Appendix

**Lemma 1.** *Let $P_n(\mathbf{x}), n \in \mathbb{N}$ and $Q_m(\mathbf{z}), m \in \mathbb{N}$ denote all monomials in $\mathbf{x}$ and $\mathbf{z}$ (in arbitrary order), respectively. Let $y$ denote a discrete random variable with $L$ values. Assume the matrix of mixed moments $M_{ij} = E[P_i(\mathbf{x})Q_j(\mathbf{z})]$, $i, j \in \{1, ..., L\}$ has full rank.*

*If $\mathbf{x}$ is independent of $\mathbf{z}$ given $y$, then*

$$E[P_n(\mathbf{x})Q_m(\mathbf{z})] = \sum_{j=1}^{L} C_{mj} E[P_n(\mathbf{x})Q_j(\mathbf{z})] \tag{169}$$

*for all $n, m \in \mathbb{N}$ and $C_{mj} \in \mathbb{R}$.*

*Proof.* Let $y \in \{1, ..., L\}$ with probabilities $\mathrm{P}(y = i)$ for $i \in \{1, ..., L\}$.[53] According to the basic properties of conditional expectations, $\mathbf{x}$ is independent of $\mathbf{z}$ given $y$ only if

$$E[P_n(\mathbf{x})Q_m(\mathbf{z})] = \sum_{i=1}^{L} E[P_n(\mathbf{x})|y = i] E[Q_m(\mathbf{z})|y = i] \mathrm{P}(y = i). \tag{170}$$

Furthermore, all conditional expectations exists because all moments exist (Kleiber and Stoyanov, 2013).

Each mixed moment can be written as a scalar product

$$E[P_n(\mathbf{x})Q_m(\mathbf{z})] = \boldsymbol{\mu}^{(n)^T} \cdot \boldsymbol{\nu}^{(m)} \tag{171}$$

$$\tag{172}$$

of $L$ dimensional vectors $\boldsymbol{\mu}^{(n)}$ and $\boldsymbol{\nu}^{(m)}$ with components

$$\mu_i^{(n)} = E[P_n(\mathbf{x})|y = i] \tag{173}$$

$$\nu_i^{(m)} = E[Q_m(\mathbf{z})|y = i] \mathrm{P}(y = i). \tag{174}$$

Because $\mathbf{M}$ has full rank, all $\boldsymbol{\nu}^{(j)}$ for $j \in \{1, ..., L\}$ have to be linearly independent and can be chosen as a vector basis in $\mathbb{R}^m$. Let $C_{mj}$ denote the corresponding coefficients such that $\boldsymbol{\nu}^{(m)} = \sum_{j=1}^{L} C_{mj} \boldsymbol{\nu}^{(j)}$. It follows,

$$E[P_n(\mathbf{x})Q_m(\mathbf{z})] = \boldsymbol{\mu}^{(n)^T} \cdot \boldsymbol{\nu}^{(m)} = \sum_{j=1}^{L} C_{mj} \boldsymbol{\mu}^{(n)^T} \cdot \boldsymbol{\nu}^{(j)} = \sum_{j=1}^{L} C_{mj} E[P_n(\mathbf{x})Q_j(\mathbf{z})] \tag{175}$$

for all $n, m \in \mathbb{N}$. □

[53] The symbol P denotes probabilities, $p$ denotes probability density functions and $P_n$ denotes monomials.

**Lemma 2.** *Let* $P_n(\mathbf{x})$, $n \in \{1, ..., n_{max}\}$ *for* $n_{max} \geq 2$ *and* $Q_m(\mathbf{z})$, $m \in \{1, ..., m_{max}\}$ *for* $m_{max} \geq 2$ *denote arbitrary monomials in* $\mathbf{x}$ *and* $\mathbf{z}$*, respectively. Assume the matrix of mixed moments* $M_{ij} = E[P_i(\mathbf{x})Q_j(\mathbf{z})]$, $i, j \in \{1, 2\}$ *has full rank.*

*Then,*

$$E[P_n(\mathbf{x})Q_m(\mathbf{z})] = \sum_{j=1}^{2} C_{mj} E[P_n(\mathbf{x})Q_j(\mathbf{z})] \tag{176}$$

*for all* $n \in \{1, ..., n_{max}\}$*, all* $m \in \{1, ..., m_{max}\}$ *and* $C_{mj} \in \mathbb{R}$ *if and only if*

$$E[P_n(\mathbf{x})Q_m(\mathbf{z})] = \sum_{i,j=1}^{2} E[P_n(\mathbf{x})Q_j(\mathbf{z})] \left(M^{-1}\right)_{ji} E[P_i(\mathbf{x})Q_m(\mathbf{z})] \tag{177}$$

*for all* $n \in \{1, ..., n_{max}\}$ *and all* $m \in \{1, ..., m_{max}\}$.

*Proof.* First, $\mathbf{M}^{-1}$ exists because $\mathbf{M}$ has full rank. Then, the *if* part follows by choosing

$$C_{mj} = \sum_{i=1}^{2} \left(M^{-1}\right)_{ji} E[P_i(\mathbf{x})Q_m(\mathbf{z})] \tag{178}$$

for $j \in \{1, 2\}$. The *only if* part follows from Equ. 176

$$E[P_i(\mathbf{x})Q_m(\mathbf{z})] = \sum_{j=1}^{2} C_{mj} E[P_i(\mathbf{x})Q_j(\mathbf{z})] \tag{179}$$

$$= \sum_{j=1}^{2} M_{ij} C_{mj} \tag{180}$$

$$\sum_{i=1}^{2} \left(M^{-1}\right)_{ji} E[P_i(\mathbf{x})Q_m(\mathbf{z})] = C_{mj} \tag{181}$$

for $i \in \{1, 2\}$. Inserting $C_{mj}$ into Equ. 176 results in

$$E[P_n(\mathbf{x})Q_m(\mathbf{z})] = \sum_{i,j=1}^{2} E[P_n(\mathbf{x})Q_j(\mathbf{z})] \left(M^{-1}\right)_{ji} E[P_i(\mathbf{x})Q_m(\mathbf{z})] \tag{182}$$

for all $n \in \{1, ..., n_{max}\}$ and all $m \in \{1, ..., m_{max}\}$. □

**Proposition 3.** *Let* $f, g : \mathbb{R}^{d^{(x)}} \times \mathbb{R}^{d^{(z)}} \rightarrow \mathbb{R}$ *denote two Borel measurable bounded functions. Let* $Q_m(\mathbf{z}), m \in \mathbb{N}$ *denote all monomials in* $\mathbf{z}$*.*

*If*

$$E\left[f(\mathbf{x}, \mathbf{z}) Q_m(\mathbf{z})\right] = E\left[g(\mathbf{x}, \mathbf{z}) Q_m(\mathbf{z})\right] \tag{183}$$

*for all* $m \in \mathbb{N}$*, then*

$$E\left[f(\mathbf{x}, \mathbf{z}) | \, \mathbf{z}\right] = E\left[g(\mathbf{x}, \mathbf{z}) | \, \mathbf{z}\right]. \tag{184}$$

[54]

*Proof.* The functions $f$ and $g$ are bounded and, thus, their expectations and conditional expectations exist ($\mathbf{x}$ and $\mathbf{z}$ are bounded). Define $w(\mathbf{x}, \mathbf{z}) = f(\mathbf{x}, \mathbf{z}) - g(\mathbf{x}, \mathbf{z})$, so that $w$ and $\mathbf{z}$ are bounded random variables with values in $\mathbb{R}$ and $\mathbb{R}^{d^{(z)}}$. Equ. 183 and the linearity of expectations imply

$$E[w(\mathbf{x}, \mathbf{z}) \mathcal{P}(\mathbf{z})] = 0 \tag{185}$$

for any polynomial $\mathcal{P}$ on $\mathbb{R}^{d^{(z)}}$. Because the set of all such polynomials is dense in $L^1(K)$ for any compact $K \subset \mathbb{R}^{d^{(z)}}$ (Driver, 2003) and $w$ and $\mathbf{z}$ are bounded, it follows, $E[w(\mathbf{x}, \mathbf{z}) h(\mathbf{z})] = 0$ for any $h \in L^1(K)$. Thus, $E[w(\mathbf{x}, \mathbf{z}) \mathbf{1}_{\mathbf{z} \in B}] = 0$ for any Borel set $B \in \mathcal{B}^d$, where $\mathbf{1}$ denotes the indicator function, and

$$E[w(\mathbf{x}, \mathbf{z}) | B] \, \mathrm{P}((\mathbf{x}, \mathbf{z}) \in B) = E[w(\mathbf{x}, \mathbf{z}) \mathbf{1}_{\mathbf{z} \in B}] = 0 \tag{186}$$

implies $E[w(\mathbf{x}, \mathbf{z}) | B] = 0$ for any $B$ with non-zero measure. Thus, $E[w(\mathbf{x}, \mathbf{z}) | \mathbf{z}] = 0$ and $E[f(\mathbf{x}, \mathbf{z}) | \mathbf{z}] = E[g(\mathbf{x}, \mathbf{z}) | \mathbf{z}]$. □

[54]We consider propositions as results for which no originality is claimed.

**Proposition 4.** *Let* $\mathbf{x}_k \in \mathbb{R}^{d^{(x)}}, k \in \mathbb{N}$ *denote an infinite sequence of uniformly bounded random variables with probability measures* $\mathrm{P}_k$ *on the Borel sets of* $\mathbb{R}^{d^{(x)}}$.

*Then, there exists a random variable* $\mathbf{x}$ *with probability measure* $\mathrm{P}$ *and a subsequence* $\mathrm{P}_{n_k}$ *for indices* $n_1 < n_2 < n_3 \ldots$ *such that*

$$\lim_{k\to\infty} \int g d\mathrm{P}_{n_k} = \int g d\mathrm{P} \tag{187}$$

*for all continuous bounded functions* $g : \mathbb{R}^{d^{(x)}} \to \mathbb{R}, \mathbf{x} \mapsto g(\mathbf{x})$.

*Proof.* A family $\Pi$ of probability measures defined on the Borel sets of $\mathbb{R}^{d^{(x)}}$ is called tight if, for each $\varepsilon > 0$, there exists a compact set $K_\varepsilon$ such that $\mathrm{P}(K_\varepsilon) > 1 - \varepsilon$ for all $\mathrm{P} \in \Pi$. Because $\mathbf{x}_k$ are uniformly bounded with joint compact support, denoted by $K$, it follows from $\mathrm{P}_k(K) = 1$ that the probability measures $\mathrm{P}_k$ are tight.

According to Prokohorov's theorem (Billingsley, 2013)[55], if a family of probability measures $\Pi$ on the Borel sets of $\mathbb{R}^{d^{(x)}}$ is tight, then it is relatively compact in the sense that for each sequence $\mathrm{P}_1, \mathrm{P}_2, ...$ in $\Pi$ there exists a probability measure P, not necessarily in $\Pi$, and a subsequence $\mathrm{P}_{n_k}$ that converges to P in the sense that

$$\lim_{k\to\infty} \int g d\mathrm{P}_{n_k} = \int g d\mathrm{P} \tag{188}$$

for all continuous bounded functions $g : \mathbb{R}^{d^{(x)}} \to \mathbb{R}, \mathbf{x} \mapsto g(\mathbf{x})$. □

**Proposition 5.** *Let* $\mathbf{x}_k \in \mathbb{R}^{d^{(x)}}, k \in \mathbb{N}$ *denote an infinite sequence of uniformly bounded random variables with probability measures* $\mathrm{P}_k$ *on the Borel sets of* $\mathbb{R}^{d^{(x)}}$. *Let* $P_n(\mathbf{x}_k), n \in \mathbb{N}$ *denote all monomials in* $\mathbf{x}_k$.

*If*

$$\lim_{k\to\infty} E[P_n(\mathbf{x}_k)] = \mu_n \tag{189}$$

*for all* $n \in \mathbb{N}$ *and* $\mu_n \in \mathbb{R}$, *then there exists a random variable* $\mathbf{x}$ *with moments*

$$E[P_n(\mathbf{x})] = \mu_n \tag{190}$$

*for all* $n \in \mathbb{N}$.

*Proof.* Let $g_n(\mathbf{x}_k)$, $n \in \mathbb{N}$ be continuous and bounded functions that agree with $P_n(\mathbf{x}_k)$ on the joint compact support of all $\mathrm{P}_k$, denoted by $K$, so that $E[P_n(\mathbf{x}_k)] = E[g_n(\mathbf{x}_k)]$. Then, it follows from Proposition 4, that there exists a random variable $\mathbf{x}$ with probability measure P and a subsequence $\mathrm{P}_{n_k}$, where $n_1 < n_2 < ...$ is an increasing sequence of indices, such that

$$\lim_{k\to\infty} E[g_n(\mathbf{x}_{n_k})] = E[g_n(\mathbf{x})]. \tag{191}$$

A sequence converges only if all its subsequences converge to the same limit and

$$\mu_n = \lim_{k\to\infty} E[P_n(\mathbf{x}_k)] = \lim_{k\to\infty} E[P_n(\mathbf{x}_{n_k})] = \lim_{k\to\infty} E[g_n(\mathbf{x}_{n_k})] = E[g_n(\mathbf{x})]. \tag{192}$$

Because P is supported on a subset of $K$,[56] $E[P_n(\mathbf{x})] = E[g_n(\mathbf{x})] = \mu_n$ for all $n \in \mathbb{N}$. □

[55] Theorem 6.1 (Prokohorov's theorem) and Theorem 2.1 (Portmanteau Theorem) in (Billingsley, 2013).

[56] Theorem 2.1 (iii) (Portmanteau Theorem) for $F = K$ in (Billingsley, 2013)

**Theorem 6.** *Let $P_n(\mathbf{x}), n \in \mathbb{N}$ and $Q_m(\mathbf{z}), m \in \mathbb{N}$ denote all monomials in $\mathbf{x}$ and $\mathbf{z}$ (in arbitrary order). Assume the matrix of mixed moments $M_{ij} = E[P_i(\mathbf{x})Q_j(\mathbf{z})]$, $i, j \in \{1, 2\}$ has full rank.*

*Then, there exists a discrete random variable $y \in \{1, 2\}$ such that $\mathbf{x}$ is independent of $\mathbf{z}$ given $y$ if and only if*

$$E[P_n(\mathbf{x})Q_m(\mathbf{z})] = \sum_{j=1}^{2} C_{mj} E[P_n(\mathbf{x})Q_j(\mathbf{z})] \tag{193}$$

*for all $n, m \in \mathbb{N}$ and $C_{mj} \in \mathbb{R}$.*

*Proof.* The *only if* part follows from Lemma 1. To prove the *if* part, we infer from Lemma 2 and Equ. 193

$$E[P_n(\mathbf{x})Q_m(\mathbf{z})] \quad = \quad \sum_{i,j=1}^{2} E[P_n(\mathbf{x})Q_j(\mathbf{z})] \left(M^{-1}\right)_{ji} E[P_i(\mathbf{x})Q_m(\mathbf{z})] \tag{194}$$

for all $n, m \in \mathbb{N}$. In the remainder of this section, we show that $\mathbf{M}^{-1}$ can be decomposed into $2 \times 2$ matrices $\mathbf{U}$, $\mathbf{S}$ and $\mathbf{V}$, where

$$\mathbf{M}^{-1} \quad = \quad \mathbf{USV}^T \tag{195}$$

and $\mathbf{S}$ is a diagonal matrix with non-negative elements and unit trace, so that

$$E[P_n(\mathbf{x})Q_m(\mathbf{z})] \quad = \quad \sum_{l=1}^{2} E\left[P_n(\mathbf{x})\left(\sum_{j=1}^{2} U_{jl} Q_j(\mathbf{z})\right)\right] \cdot S_{ll} \cdot \tag{196}$$

$$E\left[Q_m(\mathbf{z})\left(\sum_{i=1}^{2} V_{il} P_i(\mathbf{x})\right)\right] \tag{197}$$

for all $n, m \in \mathbb{N}$. We define

$$\mathrm{P}(y = l) \quad = \quad S_{ll} \tag{198}$$

$$\tag{199}$$

for $l \in \{1, 2\}$ as the probabilities of a discrete random variable $y \in \{1, 2\}$ and

$$E[P_n(\mathbf{x})|y = l] \quad = \quad E\left[P_n(\mathbf{x}) \sum_{j=1}^{2} U_{jl} Q_j(\mathbf{z})\right] \tag{200}$$

$$E[Q_m(\mathbf{z})]|y = l] \quad = \quad E\left[Q_m(\mathbf{z}) \sum_{i=1}^{2} V_{il} P_i(\mathbf{x})\right] \tag{201}$$

as the moments of the probability distributions of $\mathbf{x}$ and $\mathbf{z}$ given $y = l$ resulting in

$$E[P_n(\mathbf{x})Q_m(\mathbf{z})] \quad = \quad \sum_{l=1}^{2} E[P_n(\mathbf{x})|y = l] E[Q_m(\mathbf{z})]|y = l] \mathrm{P}(y = l). \tag{202}$$

These moments uniquely determine the probability distributions of bounded random variables (Kleiber and Stoyanov, 2013). Thus, not only the moments but also the distributions of $\mathbf{x}$ and $\mathbf{z}$ on both sides of Equ. 202 are identical and the same conditional independence statements hold, i.e. $\mathbf{x}$ is independent of $\mathbf{z}$ given $y$.

More precisely, Proposition 3 applied to Equ. 194 implies

$$E[P_n(\mathbf{x})Q_m(\mathbf{z})] = E\left[\left(\sum_{i,j=1}^{2} E[P_n(\mathbf{x})Q_j(\mathbf{z})]\left(M^{-1}\right)_{ji} P_i(\mathbf{x})\right) Q_m(\mathbf{z})\right] \tag{203}$$

$$E[P_n(\mathbf{x})|\mathbf{z}] = \sum_{i,j=1}^{2} E[P_n(\mathbf{x})Q_j(\mathbf{z})]\left(M^{-1}\right)_{ji} E[P_i(\mathbf{x})|\mathbf{z}] \tag{204}$$

$$= \sum_{j=1}^{2} E[P_n(\mathbf{x})Q_j(\mathbf{z})]u_j(\mathbf{z}), \tag{205}$$

where

$$u_j(\mathbf{z}) = E\left[\sum_{i=1}^{2}\left(M^{-1}\right)_{ji} P_i(\mathbf{x})\middle|\mathbf{z}\right]. \tag{206}$$

This equation has to hold for constant $P_n(\mathbf{x})$ resulting in

$$E[1|\mathbf{z}] = 1 = \sum_{j=1}^{2} E[Q_j(\mathbf{z})]u_j(\mathbf{z}) \tag{207}$$

for any Borel set in $\sigma(\mathbf{z})$ with non-zero measure, where $\sigma(\mathbf{z})$ denotes the $\sigma$-algebra generated by $\mathbf{z}$. Equ. 207 implies that at least one $E[Q_j(\mathbf{z})] \neq 0$ and all points $\mathbf{u}(\mathbf{z}) = (u_1(\mathbf{z}), u_2(\mathbf{z}))$ lie on a line. Without loss of generality we assume $E[Q_2(\mathbf{z})] \neq 0$.[57]

Because $\mathbf{x}$ is bounded, the conditional expectation $u_1(\mathbf{z})$ is bounded by $U_{11} \leq u_1(\mathbf{z}) \leq U_{12}$ for all $\mathbf{z}$, where we define

$$U_{11} = \inf_{B\in\sigma(\mathbf{z})} u_1(B) \tag{208}$$

$$U_{12} = \sup_{B\in\sigma(\mathbf{z})} u_1(B) \tag{209}$$

$$U_{21} = \frac{1 - E[Q_1(\mathbf{z})]U_{11}}{E[Q_2(\mathbf{z})]} \tag{210}$$

$$U_{22} = \frac{1 - E[Q_1(\mathbf{z})]U_{12}}{E[Q_2(\mathbf{z})]} \tag{211}$$

and the supremum and infimum are taken over all Borel sets $B$ with non-zero measure. Then, all $\mathbf{u}(\mathbf{z})$ are in the convex hull of $(U_{11}, U_{21})$ and $(U_{12}, U_{22})$. We insert these bounds into Equ. 205 and define

$$E[P_n(\mathbf{x})|y = l] = E\left[P_n(\mathbf{x})\sum_{j=1}^{2} U_{jl}Q_j(\mathbf{z})\right]. \tag{212}$$

Each sequence $E[P_n(\mathbf{x})|y = l]$ with index $n$ has to be a moment sequence of a random variable. This is because the supremum and infimum in Equ. 208 and 209 occur either at a certain Borel set $B_l \in \sigma(\mathbf{z})$ or at the limit point of a sequence of Borel sets $B_{kl} \in \sigma(\mathbf{z})$ indexed by $k \in \mathbb{N}$. In the former case, $E[P_n(\mathbf{x})|y = l]$ are by definition the moments $E[P_n(\mathbf{x})|B_l]$ of the random variable $\mathbf{x}$ conditioned on $\mathbf{z} \in B_l$. In the latter case, each sequence of Borel sets implies a sequence of distributions of $\mathbf{x}_{kl}$ conditioned on $\mathbf{z} \in B_{kl}$.

[57]Otherwise $E[Q_1(\mathbf{z})] \neq 0$ and we apply the limits in Equ. 208 and 209 to $u_2(B)$ in Equ. 210 and 211.

Because $\lim_{k\to\infty} u_j(\mathbf{x}_{kl}) = U_{jl}$ the conditional expectation in Equ. 205 converges to $\lim_{k\to\infty} E[P_n(\mathbf{x}_{kl})|B_{kl}] = E[P_n(\mathbf{x})|y = l]$. Then, by Proposition 5 there exists a random variable $\mathbf{x}$ with moments $E[P_n(\mathbf{x})|y = l]$. From Equ. 212 follows that the probability measure of this random variable is given by

$$\mathrm{P}_{1l}(B) = \int_B \sum_{j=1}^{2} U_{jl} Q_j(\mathbf{z}) d\mathrm{P} \tag{213}$$

$$E[P_n(\mathbf{x})|y = l] = \int P_n(\mathbf{x}) d\mathrm{P}_{1l} \tag{214}$$

for all $B \in \sigma(\mathbf{x})$, where $\sigma(\mathbf{x})$ denotes the $\sigma$-algebra generated by $\mathbf{x}$..

Furthermore, the matrix $\mathbf{U}$ has full rank and, thus, $\mathbf{U}^{-1}$ exists. Otherwise, Equ. 208 to 211 imply $U_{j1} = U_{j2}$, $u_j(\mathbf{z})$ is constant, from Equ. 205 follows

$$E[P_i(\mathbf{x})] = E[E[P_i(\mathbf{x})|\mathbf{z}]] = \sum_{j=1}^{2} E[P_i(\mathbf{x}) Q_j(\mathbf{z})] u_j = E[P_i(\mathbf{x})|\mathbf{z}] \tag{215}$$

and the matrix

$$M_{ij} = E[P_i(\mathbf{x}) Q_j(\mathbf{z})] = E[E[P_i(\mathbf{x})|\mathbf{z}] Q_j(\mathbf{z})] = E[P_i(\mathbf{x})] E[Q_j(\mathbf{z})] \tag{216}$$

has rank one violating the premise. Then, from Equ. 205 follows

$$E[P_n(\mathbf{x})|\mathbf{z}] = \sum_{i,j,l=1}^{2} E[P_n(\mathbf{x}) Q_j(\mathbf{z})] U_{jl} (U^{-1})_{li} u_i(\mathbf{z}) \tag{217}$$

$$= \sum_{l=1}^{2} E[P_n(\mathbf{x})|y = l] \bar{u}_l(\mathbf{z}) \tag{218}$$

$$\bar{u}_l(\mathbf{z}) = \sum_{i=1}^{2} (U^{-1})_{li} u_i(\mathbf{z}). \tag{219}$$

By definition, all $\mathbf{u}(\mathbf{z})$ are in the convex hull of $(U_{11}, U_{21})$ and $(U_{12}, U_{22})$. Thus, there exists a function $d : \mathbb{R}^{d^{(\mathbf{z})}} \to [0, 1]$ such that

$$\mathbf{u}(\mathbf{z}) = \mathbf{U} \begin{pmatrix} d(\mathbf{z}) \\ 1 - d(\mathbf{z}) \end{pmatrix} \tag{220}$$

$$\bar{\mathbf{u}}(\mathbf{z}) = \begin{pmatrix} d(\mathbf{z}) \\ 1 - d(\mathbf{z}) \end{pmatrix}, \tag{221}$$

$\bar{u}_l(\mathbf{z}) \in [0, 1]$ and $\sum_l \bar{u}_l(\mathbf{z}) = 1$. Furthermore, $E[\bar{u}_l(\mathbf{z})] > 0$ because $\bar{u}_l(\mathbf{z})$ is non-negative and non-constant, i.e. by definition it has to take the values 0 and 1 for the supremum and infimum of $u_l(\mathbf{z})$.

Then,

$$E[P_n(\mathbf{x}) Q_m(\mathbf{z})] = E[E[P_n(\mathbf{x})|\mathbf{z}] Q_m(\mathbf{z})] \tag{222}$$

$$= \sum_{l=1}^{2} E[P_n(\mathbf{x})|y = l] E[\bar{u}_l(\mathbf{z}) Q_m(\mathbf{z})]. \tag{223}$$

For every non-negative measurable function $f$ we can construct the Borel measure $\int_B f dP$, where $B \in \mathcal{B}^d$. Because $\bar{u}_l(\mathbf{z})$ are non-negative measurable functions with finite and positive $E[\bar{u}_l(\mathbf{z})]$ we define

$$P_{2l}(B) = \frac{1}{E[\bar{u}_l(\mathbf{z})]} \int_B \bar{u}_l(\mathbf{z}) dP \tag{224}$$

and

$$\begin{aligned} E[Q_m(\mathbf{z})|y = l] &= \frac{E\left[Q_m(\mathbf{z})\bar{u}_l(\mathbf{z})\right]}{E[\bar{u}_l(\mathbf{z})]} = \int Q_m(\mathbf{z}) dP_{2l} &(225)\\ P(y = l) &= S_{ll} = E[\bar{u}_l(\mathbf{z})], &(226) \end{aligned}$$

where $\sum_l P(y = l) = 1$ and $P(y = l) > 0$. Defining $\mathbf{V} = (\mathbf{S}^{-1}\mathbf{U}^{-1}\mathbf{M}^{-1})^T$ we obtain

$$\begin{aligned} E[Q_m(\mathbf{z})|y = l] &= \frac{E\left[Q_m(\mathbf{z})\bar{u}_l(\mathbf{z})\right]}{E[\bar{u}_l(\mathbf{z})]} &(227)\\ &= E\left[Q_m(\mathbf{z}) \sum_{i=1}^{2} V_{il} P_i(\mathbf{x})\right]. &(228) \end{aligned}$$

For $E[P_n(\mathbf{x})|y = l]$ defined in Equ. 212, we obtain the desired factorization of mixed moments (see Equ. 202)

$$\begin{aligned} E[P_n(\mathbf{x})Q_m(\mathbf{z})] &= E[E[P_n(\mathbf{x})|\mathbf{z}]Q_m(\mathbf{z})] &(229)\\ &= \sum_{l=1}^{2} E[P_n(\mathbf{x})|y = l] E[\bar{u}_l(\mathbf{z})Q_m(\mathbf{z})] &(230)\\ &= \sum_{l=1}^{2} E[P_n(\mathbf{x})|y = l] E[Q_m(\mathbf{z})|y = l] P(y = l) &(231) \end{aligned}$$

for all $n, m \in \mathbb{N}$, where the conditional expectations are taken with respect to the distributions defined in Equ. 213, 224 and 226.

In general, this factorization is not unique. The moments in Equ. 205 may define a conditional probability distribution of $\mathbf{x}$ given $y$ for points $\mathbf{u}(\mathbf{z})$ outside of the convex hull of $(U_{11}, U_{21})$ and $(U_{12}, U_{22})$ resulting in a different valid factorization. □

**Lemma 7.** *Let* $P_n(\mathbf{x})$*,* $n \in \{1, ..., n_{max}\}$*,* $n_{max} \geq 2$ *denote monomials in* $\mathbf{x}$*, where* $P_1(\mathbf{x}) = 1$ *and* $P_n(\mathbf{x}) \in \{1, x_1, ..., x_{d^{(x)}}\}$*. Let* $Q_m(\mathbf{z})$*,* $m \in \{1, ..., m_{max}\}$*,* $m_{max} \geq 2$ *denote monomials in* $\mathbf{z}$*, where* $Q_1(\mathbf{z}) = 1$ *and* $Q_m(\mathbf{z}) \in \{1, z_1, ..., z_{d^{(z)}}\}$*. Assume the matrix of mixed moments* $M_{ij} = E[P_i(\mathbf{x})Q_j(\mathbf{z})]$*,* $i, j \in \{1, 2\}$ *is diagonal and has full rank.*

*Then, there exist bounded random variables* $\tilde{\mathbf{x}} \in \mathbb{R}^{d^{(x)}}$*,* $\tilde{\mathbf{z}} \in \mathbb{R}^{d^{(z)}}$ *and* $\tilde{y} \in \{1, 2\}$ *such that* $\tilde{\mathbf{x}}$ *is independent of* $\tilde{\mathbf{z}}$ *given* $\tilde{y}$ *and*

$$E[P_n(\tilde{\mathbf{x}})Q_m(\tilde{\mathbf{z}})] = E[P_n(\mathbf{x})Q_m(\mathbf{z})] \tag{232}$$

*for all* $n \in \{1, ..., n_{max}\}$ *and all* $m \in \{1, ..., m_{max}\}$ *if and only if*

$$E[P_n(\mathbf{x})Q_m(\mathbf{z})] = \sum_{j=1}^{2} C_{mj} E[P_n(\mathbf{x})Q_j(\mathbf{z})] \tag{233}$$

*for all* $n \in \{1, ..., n_{max}\}$*, all* $m \in \{1, ..., m_{max}\}$ *and* $C_{mj} \in \mathbb{R}$*.*

*Proof.* The *only if* part follows from Lemma 1. That is, if $\mathbf{M}$ has full rank, then there exist random variables $\tilde{\mathbf{x}}$, $\tilde{\mathbf{z}}$ and $\tilde{y}$ so that $\tilde{\mathbf{x}}$ is independent of $\tilde{\mathbf{z}}$ given $\tilde{y}$ only if

$$E[P_n(\tilde{\mathbf{x}})Q_m(\tilde{\mathbf{z}})] = \sum_{j=1}^{2} C_{mj} E[P_n(\tilde{\mathbf{x}})Q_j(\tilde{\mathbf{z}})] \tag{234}$$

for all $n \in \{1, ..., n_{\max}\}$ and all $m \in \{1, ..., m_{\max}\}$. Substituting $E[P_n(\mathbf{x})Q_m(\mathbf{z})]$ for $E[P_n(\tilde{\mathbf{x}})Q_m(\tilde{\mathbf{z}})]$ results in Equ. 233.

To prove the *if* part, we infer from Lemma 2 and Equ. 233

$$E[P_n(\mathbf{x})Q_m(\mathbf{z})] = \sum_{i,j=1}^{2} E[P_n(\mathbf{x})Q_j(\mathbf{z})] \left(M^{-1}\right)_{ji} E[P_i(\mathbf{x})Q_m(\mathbf{z})] \tag{235}$$

for all $n \in \{1, ..., n_{\max}\}$ and all $m \in \{1, ..., m_{\max}\}$. In the remainder of this section, we construct random variables $\tilde{\mathbf{x}}$, $\tilde{\mathbf{z}}$ and $\tilde{y}$ so that $\tilde{\mathbf{x}}$ is independent of $\tilde{\mathbf{z}}$ given $\tilde{y}$ and show

$$E[P_n(\tilde{\mathbf{x}})Q_m(\tilde{\mathbf{z}})] = E[P_n(\mathbf{x})Q_m(\mathbf{z})] \tag{236}$$

for all $n \in \{1, ..., n_{\max}\}$ and all $m \in \{1, ..., m_{\max}\}$.

More precisely, we decomposed $\mathbf{M}^{-1}$ into the $2 \times 2$ matrices $\mathbf{U}$, $\mathbf{S}$ and $\mathbf{V}$ such that $\mathbf{M}^{-1} = \mathbf{U}\mathbf{S}\mathbf{V}^T$,

$$\mathbf{U} = \mathbf{M}^{-1}\mathbf{V} \tag{237}$$

$$\mathbf{S} = 0.5\mathbb{1} \tag{238}$$

$$\mathbf{V} = \begin{pmatrix} 1 & 1 \\ -1 & 1 \end{pmatrix}, \tag{239}$$

and $\mathbf{V}\mathbf{S}\mathbf{V}^T = \mathbb{1}$. We define the random variable $\tilde{y} \in \{1, 2\}$ with probabilities

$$\mathrm{P}(\tilde{y} = l) = S_{ll} \tag{240}$$

for $l \in \{1, 2\}$.

We define the random variable $\tilde{\mathbf{x}} \in \mathbb{R}^{d^{(x)}}$ such that

$$\tilde{\mathbf{x}} \sim \mathcal{U}(\mathbf{\Delta}^{(l)}) \tag{241}$$

given $\tilde{y} = l$, where $\mathcal{U}(\boldsymbol{\Delta}^{(l)})$ denotes the multivariate uniform distribution supported on the box

$$\{(\tilde{x}_1, ..., \tilde{x}_{d^{(x)}}) | \min\{0, \Delta_k^{(l)}\} \leq \tilde{x}_k \leq \max\{0, \Delta_k^{(l)}\}\}, \tag{242}$$

where $\boldsymbol{\Delta}^{(l)}$ denotes a vector with elements

$$\Delta_k^{(l)} = 2\sum_{j=1}^{2} E[x_k Q_j(\mathbf{z})] U_{jl}, \quad k \in \{1, ..., d^{(x)}\}. \tag{243}$$

The expected values of $P_n(\tilde{\mathbf{x}})$ for $\tilde{\mathbf{x}} \sim \mathcal{U}(\boldsymbol{\Delta}^{(l)})$ given $\tilde{y} = l$ are denoted by $E[P_n(\tilde{\mathbf{x}})|\tilde{y} = l]$, where $E[P_n(\tilde{\mathbf{x}})|\tilde{y} = l] = 0.5\Delta_k^{(l)}$ if $P_n(\tilde{\mathbf{x}}) = \tilde{x}_k$.

We define the random variable $\tilde{\mathbf{z}} \in \mathbb{R}^{d^{(z)}}$ such that

$$\tilde{\mathbf{z}} \sim \mathcal{U}(\boldsymbol{\theta}^{(l)}) \tag{244}$$

given $\tilde{y} = l$, where $\boldsymbol{\theta}^{(l)}$ denotes a vector with elements

$$\theta_k^{(l)} = 2\sum_{i=1}^{2} V_{il} E[z_k P_i(\mathbf{x})], \quad k \in \{1, ..., d^{(z)}\}. \tag{245}$$

The expected values of $Q_m(\tilde{\mathbf{z}})$ for $\tilde{\mathbf{z}} \sim \mathcal{U}(\boldsymbol{\theta}^{(l)})$ given $\tilde{y} = l$ are denoted by $E[Q_m(\tilde{\mathbf{z}})|\tilde{y} = l]$, where $E[Q_m(\tilde{\mathbf{z}})|\tilde{y} = l] = 0.5\theta_k^{(l)}$ if $Q_m(\tilde{\mathbf{z}}) = \tilde{z}_k$.

For all $n \in \{1, ..., n_{\max}\}$ such that $P_n(\mathbf{x}) \neq 1$ and all $m \in \{1, ..., m_{\max}\}$ such that $Q_m(\mathbf{z}) \neq 1$

$$E[P_n(\tilde{\mathbf{x}})|\tilde{y} = l] = E\left[\sum_{j=1}^{2} P_n(\mathbf{x}) Q_j(\mathbf{z}) U_{jl}\right] \tag{246}$$

$$E[Q_m(\tilde{\mathbf{z}})|\tilde{y} = l] = E\left[\sum_{i=1}^{2} V_{il} P_i(\mathbf{x}) Q_m(\mathbf{z})\right] \tag{247}$$

and

$$\begin{aligned}
E[P_n(\tilde{\mathbf{x}}) Q_m(\tilde{\mathbf{z}})] &= \sum_{l=1}^{2} E[P_n(\tilde{\mathbf{x}})|\tilde{y} = l] E[Q_m(\tilde{\mathbf{z}})|\tilde{y} = l] \mathrm{P}(\tilde{y} = l) && (248)\\
&= \sum_{l=1}^{2} E\left[\sum_{j=1}^{2} P_n(\mathbf{x}) Q_j(\mathbf{z}) U_{jl}\right] S_{ll} E\left[\sum_{i=1}^{2} V_{il} P_i(\mathbf{x}) Q_m(\mathbf{z})\right] && (249)\\
&= \sum_{i,j=1}^{2} E[P_n(\mathbf{x}) Q_j(\mathbf{z})] \left(M^{-1}\right)_{ji} E[P_i(\mathbf{x}) Q_m(\mathbf{z})] && (250)\\
&= E[P_n(\mathbf{x}) Q_m(\mathbf{z})]. && (251)
\end{aligned}$$

For all $n \in \{1, ..., n_{\max}\}$ such that $P_n(\mathbf{x}) = P_n(\tilde{\mathbf{x}}) = 1$ and all $m \in \{1, ..., m_{\max}\}$ such that $Q_m(\mathbf{z}) \neq 1$

$$\begin{aligned} E[P_n(\tilde{\mathbf{x}})Q_m(\tilde{\mathbf{z}})] &= \sum_{l=1}^{2} 1 \cdot E[Q_m(\tilde{\mathbf{z}})|\tilde{y} = l]\mathrm{P}(\tilde{y} = l) && (252)\\ &= \sum_{l=1}^{2} S_{ll} E\left[\sum_{i=1}^{2} V_{il} P_i(\mathbf{x}) Q_m(\mathbf{z})\right] && (253)\\ &= \frac{1}{2} E\left[\sum_{i=1}^{2} (V_{i1} + V_{i2}) P_i(\mathbf{x}) Q_m(\mathbf{z})\right] && (254)\\ &= E\left[P_1(\mathbf{x}) Q_m(\mathbf{z})\right] && (255)\\ &= E\left[P_n(\mathbf{x}) Q_m(\mathbf{z})\right], && (256) \end{aligned}$$

where we used $V_{11} + V_{12} = 2$ and $V_{21} + V_{22} = 0$.

For all $n \in \{1, ..., n_{\max}\}$ such that $P_n(\mathbf{x}) \neq 1$ and all $m \in \{1, ..., m_{\max}\}$ such that $Q_m(\mathbf{z}) = Q_m(\tilde{\mathbf{z}}) = 1$

$$\begin{aligned} E[P_n(\tilde{\mathbf{x}})Q_m(\tilde{\mathbf{z}})] &= \sum_{l=1}^{2} E[P_n(\tilde{\mathbf{x}})|\tilde{y} = l] \cdot 1 \cdot \mathrm{P}(\tilde{y} = l) && (257)\\ &= \sum_{l=1}^{2} E\left[\sum_{j=1}^{2} P_n(\mathbf{x}) Q_j(\mathbf{z}) U_{jl}\right] S_{ll} && (258)\\ &= \frac{1}{2} E\left[\sum_{i,j=1}^{2} P_n(\mathbf{x}) Q_j(\mathbf{z}) \left(M^{-1}\right)_{ji} (V_{i1} + V_{i2})\right] && (259)\\ &= \sum_{j=1}^{2} E\left[P_n(\mathbf{x}) Q_j(\mathbf{z})\right] \left(M^{-1}\right)_{j1} && (260)\\ &= E\left[P_n(\mathbf{x}) Q_1(\mathbf{z})\right] && (261)\\ &= E\left[P_n(\mathbf{x}) Q_m(\mathbf{z})\right], && (262) \end{aligned}$$

where we used that $\mathbf{M}^{-1}$ is a diagonal matrix and $\left(M^{-1}\right)_{11} = E[P_1(\mathbf{x})Q_1(\mathbf{z})]^{-1} = 1$.

For all $n \in \{1, ..., n_{\max}\}$ such that $P_n(\mathbf{x}) = P_n(\tilde{\mathbf{x}}) = 1$ and all $m \in \{1, ..., m_{\max}\}$ such that $Q_m(\mathbf{z}) = Q_m(\tilde{\mathbf{z}}) = 1$

$$\begin{aligned} E[P_n(\tilde{\mathbf{x}})Q_m(\tilde{\mathbf{z}})] &= \sum_{l=1}^{2} 1 \cdot 1 \cdot \mathrm{P}(\tilde{y} = l) && (263)\\ &= \sum_{l=1}^{2} S_{ll} = 1 && (264)\\ &= E\left[P_n(\mathbf{x}) Q_m(\mathbf{z})\right]. && (265) \end{aligned}$$

Therefore,

$$E[P_n(\tilde{\mathbf{x}})Q_m(\tilde{\mathbf{z}})] = E\left[P_n(\mathbf{x}) Q_m(\mathbf{z})\right] \quad (266)$$

for all $n \in \{1, ..., n_{\max}\}$ and all $m \in \{1, ..., m_{\max}\}$. □

**Lemma 8.** *Let $P_n(\mathbf{x})$, $n \in \{1, ..., n_{max}\}$, $n_{max} \geq 2$ denote monomials in $\mathbf{x}$, where $P_1(\mathbf{x}) = 1$ and $P_n(\mathbf{x}) \in \{1, x_1, ..., x_{d^{(x)}}\}$. Let $Q_m(\mathbf{z})$, $m \in \{1, ..., m_{max}\}$, $m_{max} \geq 2$ denote monomials in $\mathbf{z}$, where $Q_1(\mathbf{z}) = 1$ and $z_1$ occurs in each $Q_m(\mathbf{z})$ for $m > 1$ with degree one. Assume the matrix of mixed moments $M_{ij} = E[P_i(\mathbf{x})Q_j(\mathbf{z})]$, $i, j \in \{1, 2\}$ is diagonal and has full rank.*

*Then, there exist bounded random variables $\tilde{\mathbf{x}} \in \mathbb{R}^{d^{(x)}}$, $\tilde{\mathbf{z}} \in \mathbb{R}^{d^{(z)}}$ and $\tilde{y} \in \{1, 2\}$ such that $\tilde{\mathbf{x}}$ is independent of $\tilde{\mathbf{z}}$ given $\tilde{y}$ and*

$$E[P_n(\tilde{\mathbf{x}})Q_m(\tilde{\mathbf{z}})] = E[P_n(\mathbf{x})Q_m(\mathbf{z})] \tag{267}$$

*for all $n \in \{1, ..., n_{max}\}$ and all $m \in \{1, ..., m_{max}\}$ if and only if*

$$E[P_n(\mathbf{x})Q_m(\mathbf{z})] = \sum_{j=1}^{2} C_{mj} E[P_n(\mathbf{x})Q_j(\mathbf{z})] \tag{268}$$

*for all $n \in \{1, ..., n_{max}\}$, all $m \in \{1, ..., m_{max}\}$ and $C_{mj} \in \mathbb{R}$.*

*Proof.* The *only if* part follows from Lemma 1. That is, if $\mathbf{M}$ has full rank, then there exist random variables $\tilde{\mathbf{x}}$, $\tilde{\mathbf{z}}$ and $\tilde{y}$ so that $\tilde{\mathbf{x}}$ is independent of $\tilde{\mathbf{z}}$ given $\tilde{y}$ only if

$$E[P_n(\tilde{\mathbf{x}})Q_m(\tilde{\mathbf{z}})] = \sum_{j=1}^{2} C_{mj} E[P_n(\tilde{\mathbf{x}})Q_j(\tilde{\mathbf{z}})] \tag{269}$$

for all $n \in \{1, ..., n_{\max}\}$ and all $m \in \{1, ..., m_{\max}\}$. Substituting $E[P_n(\mathbf{x})Q_m(\mathbf{z})]$ for $E[P_n(\tilde{\mathbf{x}})Q_m(\tilde{\mathbf{z}})]$ results in Equ. 268.

To prove the *if* part, we infer from Lemma 2 and Equ. 268

$$E[P_n(\mathbf{x})Q_m(\mathbf{z})] = \sum_{i,j=1}^{2} E[P_n(\mathbf{x})Q_j(\mathbf{z})] \left(M^{-1}\right)_{ji} E[P_i(\mathbf{x})Q_m(\mathbf{z})] \tag{270}$$

for all $n \in \{1, ..., n_{\max}\}$ and all $m \in \{1, ..., m_{\max}\}$. In the remainder of this section, we construct random variables $\tilde{\mathbf{x}}$, $\tilde{\mathbf{z}}$ and $\tilde{y}$ so that $\tilde{\mathbf{x}}$ is independent of $\tilde{\mathbf{z}}$ given $\tilde{y}$. Subsequently, we show

$$E[P_n(\tilde{\mathbf{x}})Q_m(\tilde{\mathbf{z}})] = E[P_n(\mathbf{x})Q_m(\mathbf{z})] \tag{271}$$

for all $n \in \{1, ..., n_{\max}\}$ and all $m \in \{1, ..., m_{\max}\}$.

More precisely, we decomposed $\mathbf{M}^{-1}$ into the $2 \times 2$ matrices $\mathbf{U}$, $\mathbf{S}$ and $\mathbf{V}$ such that $\mathbf{M}^{-1} = \mathbf{U}\mathbf{S}\mathbf{V}^T$,

$$\mathbf{U} = \mathbf{M}^{-1}\mathbf{V} \tag{272}$$

$$\mathbf{S} = 0.5\mathbb{1} \tag{273}$$

$$\mathbf{V} = \begin{pmatrix} 1 & 1 \\ -1 & 1 \end{pmatrix}, \tag{274}$$

and $\mathbf{V}\mathbf{S}\mathbf{V}^T = \mathbb{1}$. We define a random variable $\tilde{y} \in \{1, 2\}$ with probabilities

$$\mathrm{P}(\tilde{y} = l) = S_{ll}, \quad l \in \{1, 2\}. \tag{275}$$

Furthermore, we define

$$\Delta_{kl} = \sum_{j=1}^{2} E[x_k Q_j(\mathbf{z})] U_{jl}, \quad k \in \{1, ..., d^{(x)}\} \tag{276}$$

and a discrete random vector $\tilde{\mathbf{x}} \in \mathbb{R}^{d^{(x)}}$, where

$$\begin{aligned} \tilde{x}_k &\in \{2\Delta_{k1} - \Delta_{k2}, 2\Delta_{k2} - \Delta_{k1}\} && (277) \\ \mathrm{P}\,(\tilde{x}_k = (2\Delta_{k1} - \Delta_{k2}) | \,\tilde{y} = l) &= 1 - \frac{l}{3} && (278) \\ \mathrm{P}\,(\tilde{x}_k = (2\Delta_{k2} - \Delta_{k1}) | \,\tilde{y} = l) &= \frac{l}{3} && (279) \\ & && (280) \end{aligned}$$

for all $k \in \{1, ..., d^{(x)}\}$ for which $\Delta_{k1} \neq \Delta_{k2}$, and $\tilde{x}_k = \Delta_{k1}$ for all $k \in \{1, ..., d^{(x)}\}$ for which $\Delta_{k1} = \Delta_{k2}$. With this choice, (i) all $\tilde{x}_k$ are either constant or binary random variables,[58] and (ii) if $P_n(\tilde{\mathbf{x}}) = \tilde{x}_k$, then the expected value of $P_n(\tilde{\mathbf{x}})$ given $\tilde{y} = l$, denoted by $E[P_n(\tilde{\mathbf{x}})|\tilde{y} = l]$, is

$$E[P_n(\tilde{\mathbf{x}})|\tilde{y} = l] = \Delta_{kl} = \sum_{j=1}^{2} E[x_k Q_j(\mathbf{z})] U_{jl}. \tag{281}$$

We define $\tilde{\mathbf{z}} \in \mathbb{R}^{d^{(z)}}$ such that

$$\tilde{z}_r = \begin{cases} \sum_{i=1}^{2} V_{il} P_i(\mathbf{x}) z_1, & r = 1 \\ z_r, & \text{otherwise} \end{cases} \tag{282}$$

given $\tilde{y} = l$, which is a Borel measurable function of $\mathbf{x}$ and $\mathbf{z}$ and, thus, a random variable. The expected values of $Q_m(\tilde{\mathbf{z}})$ for the distribution of $\tilde{\mathbf{z}}$ given $\tilde{y} = l$ (Equ. 282) are denoted by $E[Q_m(\tilde{\mathbf{z}})|\tilde{y} = l]$.

Then, for all $n \in \{1, ..., n_{\max}\}$ such that $P_n(\mathbf{x}) \neq 1$ and all $m \in \{1, ..., m_{\max}\}$ such that $Q_m(\mathbf{z}) \neq 1$

$$\begin{aligned} E[P_n(\tilde{\mathbf{x}})|\tilde{y} = l] &= E\left[\sum_{j=1}^{2} P_n(\mathbf{x}) Q_j(\mathbf{z}) U_{jl}\right] && (283) \\ E[Q_m(\tilde{\mathbf{z}})|\tilde{y} = l] &= E\left[\sum_{i=1}^{2} V_{il} P_i(\mathbf{x}) Q_m(\mathbf{z})\right] && (284) \end{aligned}$$

and

$$\begin{aligned} E[P_n(\tilde{\mathbf{x}}) Q_m(\tilde{\mathbf{z}})] &= \sum_{l=1}^{2} E[P_n(\tilde{\mathbf{x}})|\tilde{y} = l] E[Q_m(\tilde{\mathbf{z}})|\tilde{y} = l] \mathrm{P}(\tilde{y} = l) && (285) \\ &= \sum_{l=1}^{2} E\left[\sum_{j=1}^{2} P_n(\mathbf{x}) Q_j(\mathbf{z}) U_{jl}\right] S_{ll} E\left[\sum_{i=1}^{2} V_{il} P_i(\mathbf{x}) Q_m(\mathbf{z})\right] && (286) \\ &= \sum_{i,j=1}^{2} E[P_n(\mathbf{x}) Q_j(\mathbf{z})] \left(M^{-1}\right)_{ji} E[P_i(\mathbf{x}) Q_m(\mathbf{z})] && (287) \\ &= E[P_n(\mathbf{x}) Q_m(\mathbf{z})]. && (288) \end{aligned}$$

Finally, analogous to Equ. 252-265 in Lemma 7 follows

$$E[P_n(\tilde{\mathbf{x}}) Q_m(\tilde{\mathbf{z}})] = E\left[P_n(\mathbf{x}) Q_m(\mathbf{z})\right] \tag{289}$$

for all $n \in \{1, ..., n_{\max}\}$ and all $m \in \{1, ..., m_{\max}\}$. □

[58] This allows for nesting multiple functional modules so that some observables within a module are interface variables of other modules.

**Lemma 9.** *Let* $\mathbf{x} \in \mathbb{R}^{d^{(x)}}$ *denote a bounded random variable and let* $\mathbf{z} \in \mathbb{R}^{d^{(z)}}$ *denote a non-negative bounded random variable. Let* $P_n(\mathbf{x})$*,* $n \in \{1, ..., n_{max}\}$*,* $n_{max} \geq 2$ *denote arbitrary monomials in* $\mathbf{x}$*. Let* $Q_m(\mathbf{z})$*,* $m \in \{1, ..., m_{max}\}$*,* $m_{max} \geq 2$ *denote monomials in* $\mathbf{z}$ *such that* $z_1$ *occurs in each* $Q_m(\mathbf{z})$ *with degree one. Assume the matrix of mixed moments* $M_{ij} = E[P_i(\mathbf{x})Q_j(\mathbf{z})]$*,* $i, j \in \{1, 2\}$ *has full rank.*

*Then, there exist bounded random variables* $\tilde{\mathbf{x}} \in \mathbb{R}^{d^{(x)}}$*,* $\tilde{\mathbf{z}} \in \mathbb{R}^{d^{(z)}}$ *and* $\tilde{y} \in \{1, 2\}$ *such that* $\tilde{\mathbf{x}}$ *is independent of* $\tilde{\mathbf{z}}$ *given* $\tilde{y}$ *and*

$$E[P_n(\tilde{\mathbf{x}})Q_m(\tilde{\mathbf{z}})] = E[P_n(\mathbf{x})Q_m(\mathbf{z})] \tag{290}$$

*for all* $n \in \{1, ..., n_{max}\}$ *and all* $m \in \{1, ..., m_{max}\}$ *if and only if*

$$E[P_n(\mathbf{x})Q_m(\mathbf{z})] = \sum_{j=1}^{2} C_{mj} E[P_n(\mathbf{x})Q_j(\mathbf{z})] \tag{291}$$

*for all* $n \in \{1, ..., n_{max}\}$*, all* $m \in \{1, ..., m_{max}\}$ *and* $C_{mj} \in \mathbb{R}$*.*

*Proof.* The *only if* part follows from Lemma 1. That is, if $\mathbf{M}$ has full rank, then there exist random variables $\tilde{\mathbf{x}}$, $\tilde{\mathbf{z}}$ and $\tilde{y}$ so that $\tilde{\mathbf{x}}$ is independent of $\tilde{\mathbf{z}}$ given $\tilde{y}$ only if

$$E[P_n(\tilde{\mathbf{x}})Q_m(\tilde{\mathbf{z}})] = \sum_{j=1}^{2} C_{mj} E[P_n(\tilde{\mathbf{x}})Q_j(\tilde{\mathbf{z}})] \tag{292}$$

for all $n \in \{1, ..., n_{\max}\}$ and all $m \in \{1, ..., m_{\max}\}$. Substituting $E[P_n(\mathbf{x})Q_m(\mathbf{z})]$ for $E[P_n(\tilde{\mathbf{x}})Q_m(\tilde{\mathbf{z}})]$ results in Equ. 291.

To prove the *if* part, we infer from Lemma 2 and Equ. 291

$$E[P_n(\mathbf{x})Q_m(\mathbf{z})] = \sum_{i,j=1}^{2} E[P_n(\mathbf{x})Q_j(\mathbf{z})] \left(M^{-1}\right)_{ji} E[P_i(\mathbf{x})Q_m(\mathbf{z})] \tag{293}$$

for all $n \in \{1, ..., n_{\max}\}$ and all $m \in \{1, ..., m_{\max}\}$. In the remainder of this section, we construct random variables $\tilde{\mathbf{x}}$, $\tilde{\mathbf{z}}$ and $\tilde{y}$ so that $\tilde{\mathbf{x}}$ is independent of $\tilde{\mathbf{z}}$ given $\tilde{y}$. Subsequently, we show

$$E[P_n(\tilde{\mathbf{x}})Q_m(\tilde{\mathbf{z}})] = E[P_n(\mathbf{x})Q_m(\mathbf{z})] \tag{294}$$

for all $n \in \{1, ..., n_{\max}\}$ and all $m \in \{1, ..., m_{\max}\}$.

More precisely, we define a random variable $\tilde{y} \in \{1, 2\}$ with probabilities

$$\mathrm{P}(\tilde{y} = l) = \frac{1}{2}, \tag{295}$$

where $l \in \{1, 2\}$.

For every non-negative measurable function $f$ we can construct the Borel measure $\int_B f d\mathrm{P}$, where $B \in \mathcal{B}^d$. Because $Q_l(\mathbf{z})$ are non-negative measurable functions with finite and positive $E[Q_l(\mathbf{z})]$ (otherwise $Q_l(\mathbf{z})$ has to be zero for all $\mathbf{z}$ and $\mathbf{M}$ has not full rank) we define

$$\mathrm{P}_l(B) = \frac{1}{E[Q_l(\mathbf{z})]} \int_B Q_l(\mathbf{z}) d\mathrm{P} \tag{296}$$

and a random variable $\tilde{\mathbf{x}} \in \mathbb{R}^{d^{(x)}}$ with these distributions so that

$$E[P_n(\tilde{\mathbf{x}})|\tilde{y} = l] = \frac{E\left[P_n(\mathbf{x})Q_l(\mathbf{z})\right]}{E[Q_l(\mathbf{z})]} = \int P_n(\tilde{\mathbf{x}}) d\mathrm{P}_l \tag{297}$$

given $\tilde{y} = l$.

Finally, we define $\tilde{\mathbf{z}} \in \mathbb{R}^{d^{(z)}}$ such that

$$\tilde{z}_r = \begin{cases} \sum_{i=1}^{2} 2E[Q_l(\mathbf{z})] \left(M^{-1}\right)_{li} P_i(\mathbf{x}) z_1, & r = 1 \\ z_r, & \text{otherwise} \end{cases} \tag{298}$$

given $\tilde{y} = l$, which is a Borel measurable function of $\mathbf{x}$ and $\mathbf{z}$ and, thus, a random variable. The expected values of $Q_m(\tilde{\mathbf{z}})$ for the distribution of $\tilde{\mathbf{z}}$ given $\tilde{y} = l$ (Equ. 298) are denoted by $E[Q_m(\tilde{\mathbf{z}})|\tilde{y} = l]$. Then,

$$E[P_n(\tilde{\mathbf{x}})Q_m(\tilde{\mathbf{z}})] = \sum_{l=1}^{2} E[P_n(\tilde{\mathbf{x}})|\tilde{y} = l] E[Q_m(\tilde{\mathbf{z}})|\tilde{y} = l] \mathrm{P}(\tilde{y} = l) \tag{299}$$

$$= \sum_{l=1}^{2} \frac{E\left[P_n(\mathbf{x})Q_l(\mathbf{z})\right]}{E[Q_l(\mathbf{z})]} \cdot \tag{300}$$

$$\sum_{i=1}^{2} 2E[Q_l(\mathbf{z})] \left(M^{-1}\right)_{li} E[P_i(\mathbf{x})Q_m(\mathbf{z})] \cdot \frac{1}{2} \tag{301}$$

$$= \sum_{i,l=1}^{2} E[P_n(\mathbf{x})Q_l(\mathbf{z})] \left(M^{-1}\right)_{li} E[P_i(\mathbf{x})Q_m(\mathbf{z})] \tag{302}$$

$$= E[P_n(\mathbf{x})Q_m(\mathbf{z})] \tag{303}$$

for all $n \in \{1, ..., n_{\max}\}$ and all $m \in \{1, ..., m_{\max}\}$. □

**Theorem 10.** *Let $\mathcal{M}$ denote a flat modularization. Furthermore, let all observable components be correlated with $s_d$, i.e. $E[s_k s_d] \neq 0$ for all $k \in \{1, ..., d-1\}$, and normalized, i.e. $E[s_d] = 0$ and $E[s_k] = 0$ for all $k \in \{1, ..., d-1\}$.*

*Then, there exist bounded random variables $\tilde{\mathbf{s}} \in \mathbb{R}^d$ and $\tilde{y}^{(c)} \in \{1, 2\}$ such that (i) the elements of $\tilde{\mathbf{s}}$ inside of the c-th functional module of $\mathcal{M}$, denoted by $\tilde{\mathbf{x}}^{(c)}$, are independent of the elements of $\tilde{\mathbf{s}}$ outside of the c-th functional module of $\mathcal{M}$, denoted by $\tilde{\mathbf{z}}^{(c)}$, given $\tilde{y}^{(c)}$ for all $c \in \{1, ..., d^{(\mathcal{M})}\}$ and (ii)*

$$E[\tilde{s}_k \tilde{s}_d] = E[s_k s_d], \quad E[\tilde{s}_k] = E[s_k], \quad E[\tilde{s}_d] = E[s_d] \tag{304}$$

*for all $k \in \{1, ..., d-1\}$ and*

$$E[\tilde{x}_k^{(c)} \tilde{z}_1^{(c)} \tilde{z}_l^{(c)}] = E[x_k^{(c)} z_1^{(c)} z_l^{(c)}] \tag{305}$$

*for all $k \in \{1, ..., |S_c|\}$, all $l \in \{2, ..., d - |S_c|\}$ and all $c \in \{1, ..., d^{(\mathcal{M})}\}$ if and only if*

$$B_l^{(c)} = \frac{E[x_k^{(c)} z_1^{(c)} z_l^{(c)}]}{E[x_k^{(c)} z_1^{(c)}] E[z_1^{(c)} z_l^{(c)}]} \tag{306}$$

*for all $k \in \{1, ..., |S_c|\}$, all $l \in \{2, ..., d - |S_c|\}$, all $c \in \{1, ..., d^{(\mathcal{M})}\}$ and $B_l^{(c)} \in \mathbb{R}$.*

*Proof.* All $S_c$ are non-empty, because all $\mathbf{x}^{(c)}$ have by definition at least one element. The *only if* part follows from Lemma 1. For the $c$-th functional module, let

$$P_n(\tilde{\mathbf{x}}^{(c)}) = \begin{cases} 1, & n = 1 \\ \tilde{x}_{n-1}^{(c)}, & \text{otherwise} \end{cases}, \quad n \in \{1, ..., d^{(\mathrm{x},c)} + 1\} \tag{307}$$

$$Q_m(\tilde{\mathbf{z}}^{(c)}) = \begin{cases} 1, & m = 1 \\ \tilde{z}_1^{(c)}, & m = 2 \\ \tilde{z}_1^{(c)} \tilde{z}_{m-1}^{(c)}, & \text{otherwise,} \end{cases}, \quad m \in \{1, ..., d^{(\mathrm{z},c)} + 1\}, \tag{308}$$

where $d^{(\mathrm{x},c)} = |S_c|$ and $d^{(\mathrm{z},c)} = d - |S_c|$. Then, the matrix $M_{ij}^{(c)} = E[P_i(\tilde{\mathbf{x}}^{(c)}) Q_j(\tilde{\mathbf{z}}^{(c)})]$, $i, j \in \{1, 2\}$ is diagonal and has full rank, i.e.

$$\mathbf{M}^{(c)} = \begin{pmatrix} 1 & 0 \\ 0 & E[\tilde{x}_1^{(c)} \tilde{z}_1^{(c)}] \end{pmatrix}, \tag{309}$$

because, according to Equ. 304, $E[\tilde{s}_d] = E[s_d] = 0$ and

$$E[\tilde{s}_k] = E[s_k] = 0 \tag{310}$$

$$E[\tilde{s}_k \tilde{s}_d] = E[s_k s_d] \neq 0 \tag{311}$$

for all $k \in \{1, ..., d-1\}$. According to Lemma 1, there exist random variables $\tilde{\mathbf{x}}^{(c)}$, $\tilde{\mathbf{z}}^{(c)}$ and $\tilde{y}^{(c)}$ so that $\tilde{\mathbf{x}}^{(c)}$ is independent of $\tilde{\mathbf{z}}^{(c)}$ given $\tilde{y}^{(c)}$ only if

$$E[P_n(\tilde{\mathbf{x}}^{(c)}) Q_m(\tilde{\mathbf{z}}^{(c)})] = \sum_{j=1}^{2} C_{mj}^{(c)} E[P_n(\tilde{\mathbf{x}}^{(c)}) Q_j(\tilde{\mathbf{z}}^{(c)})] \tag{312}$$

for all $n \in \{1, ..., d^{(\mathrm{x},c)} + 1\}$, all $m \in \{1, ..., d^{(\mathrm{z},c)} + 1\}$ and $C_{mj}^{(c)} \in \mathbb{R}$. From Equ. 309 and Lemma 2 follows that Equ. 312 holds if and only if

$$E[\tilde{x}_k^{(c)} \tilde{z}_1^{(c)} \tilde{z}_l^{(c)}] = \frac{E[\tilde{x}_k^{(c)}] E[\tilde{z}_1^{(c)} \tilde{z}_l^{(c)}]}{M_{11}} + \frac{E[\tilde{x}_k^{(c)} \tilde{z}_1^{(c)}] E[\tilde{x}_1^{(c)} \tilde{z}_1^{(c)} \tilde{z}_l^{(c)}]}{M_{22}} \tag{313}$$

$$\frac{E[\tilde{x}_k^{(c)} \tilde{z}_1^{(c)} \tilde{z}_l^{(c)}]}{E[\tilde{x}_k^{(c)} \tilde{z}_1^{(c)}]} = \frac{E[\tilde{x}_1^{(c)} \tilde{z}_1^{(c)} \tilde{z}_l^{(c)}]}{E[\tilde{x}_1^{(c)} \tilde{z}_1^{(c)}]} = \frac{E[\tilde{x}_{k_2}^{(c)} \tilde{z}_1^{(c)} \tilde{z}_l^{(c)}]}{E[\tilde{x}_{k_2}^{(c)} \tilde{z}_1^{(c)}]} \tag{314}$$

for all $k, k_2 \in \{1, ..., d^{(x,c)}\}$ and all $l \in \{2, ..., d^{(z,c)}\}$.[59] Dividing both sides of the last equation by $E[z_1^{(c)} z_l^{(c)}]$ and substituting

$$E[\tilde{x}_k^{(c)} \tilde{z}_1^{(c)} \tilde{z}_l^{(c)}] = E[x_k^{(c)} z_1^{(c)} z_l^{(c)}] \tag{315}$$

$$E[\tilde{x}_k^{(c)} \tilde{z}_1^{(c)}] = E[x_k^{(c)} z_1^{(c)}] \tag{316}$$

results in the condition

$$B_l^{(c)} = \frac{E[x_k^{(c)} z_1^{(c)} z_l^{(c)}]}{E[x_k^{(c)} z_1^{(c)}] E[z_1^{(c)} z_l^{(c)}]} \tag{317}$$

for all $k \in \{1, ..., d^{(x,c)}\}$, all $l \in \{2, ..., d^{(z,c)}\}$ and $B_l^{(c)} \in \mathbb{R}$. These conditions have to hold for all $c \in \{1, ..., d^{(\mathcal{M})}\}$, which concludes the proof of the *only if* part.

To prove the *if* part, we apply the following iterative method to construct bounded random variables $\tilde{\mathbf{s}}^{(t)} \in \mathbb{R}^d$, $t \in \{1, ..., d^{(\mathcal{M})}\}$ from $\tilde{\mathbf{s}}^{(t-1)}$, where $\tilde{\mathbf{s}}^{(0)} = \mathbf{s}$. Let $\tilde{\mathbf{x}}^{(c,t)}$ and $\tilde{\mathbf{z}}^{(c,t)}$ denote the elements of $\tilde{\mathbf{s}}^{(t)}$ inside and outside of the $c$-th functional module of $\mathcal{M}$, where $\tilde{z}_1^{(c,t)}$ denotes $\tilde{s}_d^{(t)}$. We show that the following two conditions are fulfilled for the construction outlined below.

1. $\tilde{\mathbf{x}}^{(c,t)}$ is independent of $\tilde{\mathbf{z}}^{(c,t)}$ given $\tilde{y}^{(c)}$ for all $c \in \{1, ..., t\}$. That is, $\tilde{\mathbf{s}}^{(t)}$ consists of the same $t-1$ functional modules as $\tilde{\mathbf{s}}^{(t-1)}$ for all $t > 1$.

2. $$E[\tilde{s}_k^{(t)} \tilde{s}_d^{(t)}] = E[\tilde{s}_k^{(t-1)} \tilde{s}_d^{(t-1)}], \quad E[\tilde{s}_k^{(t)}] = E[\tilde{s}_k^{(t-1)}], \quad E[\tilde{s}_d^{(t)}] = E[\tilde{s}_d^{(t-1)}] \tag{318}$$

   for all $k \in \{1, ..., d-1\}$ and all $t \in \{1, ..., d^{(\mathcal{M})}\}$, and

   $$E[P_n(\tilde{\mathbf{x}}^{(c,t)}) Q_m(\tilde{\mathbf{z}}^{(c,t)})] = E[P_n(\tilde{\mathbf{x}}^{(c,t-1)}) Q_m(\tilde{\mathbf{z}}^{(c,t-1)})] \tag{319}$$

   for all $n \in \{1, ..., d^{(x,c)} + 1\}$, all $m \in \{1, ..., d^{(z,c)} + 1\}$, all $c \in \{1, ..., d^{(\mathcal{M})}\}$ and all $t \in \{1, ..., d^{(\mathcal{M})}\}$. That is, all $\tilde{\mathbf{s}}^{(t)}$ have the same moments as $\mathbf{s}$ in Equ. 304 and 305.

Then, defining $\tilde{\mathbf{s}} = \tilde{\mathbf{s}}^{(d^{(\mathcal{M})})}$, condition 1 implies that (i) $\tilde{\mathbf{x}}^{(c)}$ is independent of $\tilde{\mathbf{z}}^{(c)}$ given $\tilde{y}^{(c)}$ for all $c \in \{1, ..., d^{(\mathcal{M})}\}$ and condition 2 implies that (ii)

$$E[P_n(\tilde{\mathbf{x}}^{(c)}) Q_m(\tilde{\mathbf{z}}^{(c)})] = E[P_n(\mathbf{x}^{(c)}) Q_m(\mathbf{z})^{(c)})] \tag{320}$$

for all $n \in \{1, ..., d^{(x,c)} + 1\}$, all $m \in \{1, ..., d^{(z,c)} + 1\}$ and all $c \in \{1, ..., d^{(\mathcal{M})}\}$, which concludes the proof.

We first show that conditions 1 and 2 hold for $t = 1$. As already shown for the *only if* part, the conditions for $\tilde{\mathbf{s}}^{(t-1)} = \tilde{\mathbf{s}}^{(0)} = \mathbf{s}$ in Equ. 306 (and Equ. 317) hold if and only if the conditions in Equ. 312 hold for all $c \in \{1, ..., d^{(\mathcal{M})}\}$. Then, according to Lemma 8, for the choice $c_t = t$, there exist bounded random variables $\tilde{\mathbf{s}}^{(t)} \in \mathbb{R}^d$, $\tilde{\mathbf{x}}^{(c_t,t)} \in \mathbb{R}^{d^{(x,c_t)}}$, $\tilde{\mathbf{z}}^{(c_t,t)} \in \mathbb{R}^{d^{(z,c_t)}}$ and $\tilde{y}^{(c_t)} \in \{1, 2\}$ so that $\tilde{\mathbf{x}}^{(c_t,t)}$ is independent of $\tilde{\mathbf{z}}^{(c_t,t)}$ given $\tilde{y}^{(c_t)}$ and

$$E[P_n(\tilde{\mathbf{x}}^{(c_t,t)}) Q_m(\tilde{\mathbf{z}}^{(c_t,t)})] = E[P_n(\tilde{\mathbf{x}}^{(c_t,t-1)}) Q_m(\tilde{\mathbf{z}}^{(c_t,t-1)})] \tag{321}$$

for all $n \in \{1, ..., d^{(x,c_t)} + 1\}$ and all $m \in \{1, ..., d^{(z,c_t)} + 1\}$. Lemma 8 does not guarantee the remaining conditions in Equ. 318 and 319, which, because of the flat modularization, are only conditions for $\tilde{\mathbf{z}}^{(c_t,t)}$. That is,

$$E[\tilde{z}_r^{(c_t,t)}] = E[\tilde{z}_r^{(c_t,t-1)}] \tag{322}$$

$$E[\tilde{z}_1^{(c_t,t)} \tilde{z}_r^{(c_t,t)} \tilde{z}_{r_2}^{(c_t,t)}] = E[\tilde{z}_1^{(c_t,t-1)} \tilde{z}_r^{(c_t,t-1)} \tilde{z}_{r_2}^{(c_t,t-1)}] \tag{323}$$

[59] Equ. 312 is always true for $n \in \{1, 2\}$ or $m \in \{1, 2\}$ as can be verified using Lemma 2.

for all $r \in \{2, ..., d^{(z,c_t)}\}$ and all $r_2 \in R^{(r)}$, where $R^{(r)} \subseteq \{2, ..., d^{(z,c_t)}\}$ depends on $r$.

These conditions can be fulfilled by the same construction of $\tilde{\mathbf{s}}^{(t)}$ as used to prove the existence of $\tilde{\mathbf{s}}$ in Lemma 8. That is, we define the random variable $\tilde{\mathbf{z}}^{(c_t,t)} \in \mathbb{R}^{d^{(z,c_t)}}$ by (see Equ. 282)

$$\tilde{z}_r^{(c_t,t)} = \begin{cases} \sum_{i=1}^{2} V_{il} P_i(\tilde{\mathbf{x}}^{(c_t,t-1)}) \tilde{z}_1^{(c_t,t-1)}, & r = 1 \\ \tilde{z}_r^{(c_t,t-1)}, & \text{otherwise} \end{cases} \tag{324}$$

$$\mathbf{V} = \begin{pmatrix} 1 & 1 \\ -1 & 1 \end{pmatrix} \tag{325}$$

given $\tilde{y}^{(c_t)} = l$, which implies the conditions in Equ. 322 and 323 (see Equ. 252 to 256) and, thus, concludes the proof of conditions 1 and 2 for $t = 1$.

Furthermore, condition 2 holds also for $t > 1$. More precisely, condition 2 for $t = 1$ implies that all conditions for $\tilde{\mathbf{s}}^{(t-1)}$ (i.e. $\mathbf{s}$) in Equ. 306 hold for $\tilde{\mathbf{s}}^{(t)}$ as well and $\mathbf{M}^{(c)}$ is diagonal and has full rank for all $c \in \{1, ..., d^{(\mathcal{M})}\}$. Then, the steps in the previous paragraph can be repeated to iteratively prove condition 2 and the existence of $\tilde{\mathbf{s}}^{(t)}$ for all $t \in \{2, ..., d^{(\mathcal{M})}\}$.

Finally, condition 1 for $t > 1$ is not directly implied by Lemma 8, which guarantees only that $\tilde{\mathbf{s}}^{(t)}$ consists of the $c_t$-th functional module and not all $t-1$ modules within $\tilde{\mathbf{s}}^{(t-1)}$. However, condition 1 for $t > 1$ is fulfilled by the construction of $\tilde{\mathbf{s}}^{(t)}$ as used to prove the existence of $\tilde{\mathbf{s}}$ in Lemma 8 (see above). More precisely, the construction of $\tilde{\mathbf{s}}^{(t)}$ for $t > 1$ and $c_t = t$ implies the following conditional independence statements.

- $\tilde{z}_r^{(c_t,t)}$ depends only on $\tilde{z}_r^{(c_t,t-1)}$ for all $r \in \{2, ..., d^{(z,c_t)}\}$ (see Equ. 324).
- $\tilde{z}_1^{(c_t,t)}$ depends only on $\tilde{\mathbf{x}}^{(c_t,t-1)}$, $\tilde{z}_1^{(c_t,t-1)}$ and $\tilde{y}^{(c_t)}$ (see Equ. 324).
- $\tilde{\mathbf{x}}^{(c_t,t)}$ depends only on $\tilde{y}^{(c_t)}$ (see construction in Lemma 8).
- $\tilde{y}^{(c_t)}$ does not directly depend on $\tilde{\mathbf{s}}^{(t-1)}$ (only through $\tilde{z}_1^{(c_t,t)}$ in Equ. 324, see construction in Lemma 8).

Then, if $\tilde{z}_r^{(c_t,t-1)}, r \in \{2, ..., d^{(z,c_t)}\}$ is independent of $\tilde{z}_1^{(c_t,t-1)}$ and $\tilde{\mathbf{x}}^{(c_t,t-1)}$ given $\tilde{y}^{(c)}$, where $c \in \{1, ..., t-1\}$, the dependencies between variables can be described by the following Bayesian network.

$$\tilde{z}_r^{(c_t,t)} \leftarrow \tilde{z}_r^{(c_t,t-1)} \leftarrow \tilde{y}^{(c)} \rightarrow (\tilde{z}_1^{(c_t,t-1)}, \tilde{\mathbf{x}}^{(c_t,t-1)}) \rightarrow \tilde{z}_1^{(c_t,t)} \leftarrow \tilde{y}^{(c_t)} \rightarrow \tilde{\mathbf{x}}^{(c_t,t)}, \tag{326}$$

where we omitted $r \in \{2, ..., d^{(z,c_t)}\}$ for notational clarity.

Now, let $\tilde{\mathbf{x}}^{(c,t-1)}$ be independent of $\tilde{\mathbf{z}}^{(c,t-1)}$ given $\tilde{y}^{(c)}$, where $c \in \{1, ..., t-1\}$. Because of the nested modularization, all elements of $\tilde{\mathbf{x}}^{(c,t-1)}$ correspond to elements of $\tilde{z}_r^{(c_t,t-1)}, r \in \{2, ..., d^{(z,c_t)}\}$ and all elements of $\tilde{\mathbf{x}}^{(c_t,t-1)}$ correspond to elements of $\tilde{z}_r^{(c,t-1)}, r \in \{2, ..., d^{(z,c)}\}$. Thus, the dependencies between variables can be described by the following Bayesian network.

$$\tilde{\mathbf{x}}^{(c,t)} \leftarrow \tilde{\mathbf{x}}^{(c,t-1)} \leftarrow \tilde{y}^{(c)} \rightarrow (\tilde{z}_1^{(c_t,t-1)}, \tilde{z}_r^{(c,t-1)}) \rightarrow (\tilde{z}_1^{(c_t,t)}, \tilde{z}_r^{(c,t)}), \tag{327}$$

where we again omitted $r \in \{2, ..., d^{(z,c_t)}\}$ for notational clarity. By definition, $\tilde{z}_1^{(c,t)} = \tilde{z}_1^{(c_t,t)}$ and it follows

$$\tilde{\mathbf{x}}^{(c,t)} \leftarrow \tilde{y}^{(c)} \rightarrow \tilde{\mathbf{z}}^{(c,t)}. \tag{328}$$

Each variable in a Bayesian network is conditionally independent of its non-descendants given its parent variables. Thus, $\tilde{\mathbf{x}}^{(c,t)}$ is independent of $\tilde{\mathbf{z}}^{(c,t)}$ given $\tilde{y}^{(c)}$ for all $c \in \{1, ..., t\}$ (where the case $c = t$ is implied by Lemma 8), which concludes the proof of condition 1 for $t > 1$. □

**Lemma 11.** *Let $\mathcal{M}$ denote a nested modularization. Let $\hat{\mathbf{b}} = \mathbf{b} + \boldsymbol{\rho}$ for $\boldsymbol{\rho} \in \mathbb{R}^{d^{(X_0)}}$ and $\mathbf{b} \in \mathbb{R}^{d^{(X_0)}}$ such that $b_k = b_l$ for all $k, l \in X_c^{(\mathcal{M})}$ and all $c \in \{1, ..., d^{(X)}\}$.*[60] *Furthermore, let $\boldsymbol{\Sigma} \in \mathbb{R}^{d^{(X_0)} \times d^{(X_0)}}$ such that all diagonal elements are non-zero.*

*Then,*

$$\mathcal{T}_{\mathcal{M}} = \frac{1}{2} \sum_{c_0=1}^{d^{(X_0)}} \frac{\left(\sum_{k_1 \in X_{c_0}^{(\mathcal{M}_0)}} \hat{b}_{k_1} (\Sigma_{k_1 k_1})^{-1}\right)^2}{\sum_{k_2 \in X_{c_0}^{(\mathcal{M}_0)}} (\Sigma_{k_2 k_2})^{-1}} - \frac{1}{2} \sum_{c=1}^{d^{(X)}} \frac{\left(\sum_{l_1 \in X_c^{(\mathcal{M})}} \hat{b}_{l_1} (\Sigma_{l_1 l_1})^{-1}\right)^2}{\sum_{l_2 \in X_c^{(\mathcal{M})}} (\Sigma_{l_2 l_2})^{-1}} \tag{329}$$

*is precisely*

$$\mathcal{T}_{\mathcal{M}} = \frac{1}{2} \sum_{c=1}^{d^{(X)}} \left( \sum_{c_0 \in K^{(c)}} \frac{\left(\sum_{k_1 \in X_{c_0}^{(\mathcal{M}_0)}} \rho_{k_1} (\Sigma_{k_1 k_1})^{-1}\right)^2}{\sum_{k_2 \in X_{c_0}^{(\mathcal{M}_0)}} (\Sigma_{k_2 k_2})^{-1}} - \frac{\left(\sum_{l_1 \in X_c^{(\mathcal{M})}} \rho_{l_1} (\Sigma_{l_1 l_1})^{-1}\right)^2}{\sum_{l_2 \in X_c^{(\mathcal{M})}} (\Sigma_{l_2 l_2})^{-1}} \right) \tag{330}$$

*for all $\mathcal{M} \neq \mathcal{M}_0$, i.e., $\mathcal{T}_{\mathcal{M}}$ is independent of* $\mathbf{b}$.

*Proof.* Because $X_c^{(\mathcal{M})}$ and $X_c^{(\mathcal{M}_0)}$ are concise index sets they consist of the same indices, i.e.

$$\bigcup_{c_0 \in \{1, ..., d^{(X_0)}\}} X_{c_0}^{(\mathcal{M}_0)} = \bigcup_{c \in \{1, ..., d^{(X)}\}} X_c^{(\mathcal{M})}, \tag{331}$$

and each index occurs in exactly one index set associated with $\mathcal{M}$ and $\mathcal{M}_0$, respectively. Furthermore, $\mathcal{M}$ is nested and $X_c^{(\mathcal{M}_0)}$ are sets consisting of only one observable component. Therefore, for all $c \in \{1, ..., d^{(X)}\}$ there exists a set $K^{(c)} \subseteq \{1, ..., d^{(X_0)}\}$ such that

$$X_c^{(\mathcal{M})} = \bigcup_{c_0 \in K^{(c)}} X_{c_0}^{(\mathcal{M}_0)}. \tag{332}$$

By assumption, $b_k = b_l = b_c^{(\mathcal{M})}$ for all $k \in X_{c_0}^{(\mathcal{M}_0)}$, all $l \in X_c^{(\mathcal{M})}$, all $c_0 \in K^{(c)}$, all $c \in \{1, ..., d^{(X)}\}$ and $b_c^{(\mathcal{M})} \in \mathbb{R}$. It follows

$$\mathcal{T}_{\mathcal{M}} = \frac{1}{2} \sum_{c=1}^{d^{(X)}} \left( \sum_{c_0 \in K^{(c)}} \frac{\left(\sum_{k_1 \in X_{c_0}^{(\mathcal{M}_0)}} (b_c^{(\mathcal{M})} + \rho_{k_1}) (\Sigma_{k_1 k_1})^{-1}\right)^2}{\sum_{k_2 \in X_{c_0}^{(\mathcal{M}_0)}} (\Sigma_{k_2 k_2})^{-1}} - \frac{\left(\sum_{l_1 \in X_c^{(\mathcal{M})}} (b_c^{(\mathcal{M})} + \rho_{l_1}) (\Sigma_{l_1 l_1})^{-1}\right)^2}{\sum_{l_2 \in X_c^{(\mathcal{M})}} (\Sigma_{l_2 l_2})^{-1}} \right) \tag{333}$$

[60] $\mathbf{b}$ not necessarily denotes the mean of the random variable $\hat{\mathbf{b}}$ and can even depend on $\hat{\mathbf{b}}$ as long as $b_k = b_l$ for all $k, l \in X_c^{(\mathcal{M})}$ and all $c \in \{1, ..., d^{(X)}\}$.

and

$$\mathcal{T}_{\mathcal{M}} = \frac{1}{2}\sum_{c=1}^{d^{(\mathcal{X})}}\left[\sum_{c_0\in K^{(c)}}\left(\left(b_c^{(\mathcal{M})}\right)^2\sum_{k_1\in X_{c_0}^{(\mathcal{M}_0)}}\left(\Sigma_{k_1k_1}\right)^{-1}+\right.\right. \tag{334}$$

$$\left.2b_c^{(\mathcal{M})}\sum_{k_2\in X_{c_0}^{(\mathcal{M}_0)}}\rho_{k_2}\left(\Sigma_{k_2k_2}\right)^{-1}+\frac{\left(\sum_{k_3\in X_{c_0}^{(\mathcal{M}_0)}}\rho_{k_3}\left(\Sigma_{k_3k_3}\right)^{-1}\right)^2}{\sum_{k_4\in X_{c_0}^{(\mathcal{M}_0)}}\left(\Sigma_{k_4k_4}\right)^{-1}}\right)- \tag{335}$$

$$\left(\left(b_c^{(\mathcal{M})}\right)^2\sum_{l_1\in X_c^{(\mathcal{M})}}\left(\Sigma_{l_1l_1}\right)^{-1}+\right. \tag{336}$$

$$\left.\left.2b_c^{(\mathcal{M})}\sum_{l_2\in X_c^{(\mathcal{M})}}\rho_{l_2}\left(\Sigma_{l_2l_2}\right)^{-1}+\frac{\left(\sum_{l_3\in X_c^{(\mathcal{M})}}\rho_{l_3}\left(\Sigma_{l_3l_3}\right)^{-1}\right)^2}{\sum_{l_4\in X_c^{(\mathcal{M})}}\left(\Sigma_{l_4l_4}\right)^{-1}}\right)\right] \tag{337}$$

$$= \frac{1}{2}\sum_{c=1}^{d^{(\mathcal{X})}}\left(\sum_{c_0\in K^{(c)}}\frac{\left(\sum_{k_1\in X_{c_0}^{(\mathcal{M}_0)}}\rho_{k_1}\left(\Sigma_{k_1k_1}\right)^{-1}\right)^2}{\sum_{k_2\in X_{c_0}^{(\mathcal{M}_0)}}\left(\Sigma_{k_2k_2}\right)^{-1}}-\frac{\left(\sum_{l_1\in X_c^{(\mathcal{M})}}\rho_{l_1}\left(\Sigma_{l_1l_1}\right)^{-1}\right)^2}{\sum_{l_2\in X_c^{(\mathcal{M})}}\left(\Sigma_{l_2l_2}\right)^{-1}}\right), \tag{338}$$

which is independent of **b**. □

**Theorem 12.** *Let $\mathcal{M}$ denote a nested modularization. Furthermore, let $\hat{\mathbf{b}} \sim \mathcal{N}(\mathbf{b}, \mathbf{\Sigma})$, where $\mathcal{N}$ denotes the normal distribution with covariance matrix $\mathbf{\Sigma}$ and mean $\mathbf{b} \in \mathbb{R}^{d^{(X_0)}}$ such that $b_k = b_l$ for all $k, l \in X_c^{(\mathcal{M})}$ and all $c \in \{1, ..., d^{(X)}\}$.*

*If $\mathbf{\Sigma}$ if diagonal and has full-rank, then*

$$\mathcal{T}_{\mathcal{M}}(\hat{\mathbf{b}}, \mathbf{\Sigma}) = \frac{1}{2} \sum_{c_0=1}^{d^{(X_0)}} \frac{\left(\sum_{k_1 \in X_{c_0}^{(\mathcal{M}_0)}} \hat{b}_{k_1} (\Sigma_{k_1 k_1})^{-1}\right)^2}{\sum_{k_2 \in X_{c_0}^{(\mathcal{M}_0)}} (\Sigma_{k_2 k_2})^{-1}} - \frac{1}{2} \sum_{c=1}^{d^{(X)}} \frac{\left(\sum_{l_1 \in X_c^{(\mathcal{M})}} \hat{b}_{l_1} (\Sigma_{l_1 l_1})^{-1}\right)^2}{\sum_{l_2 \in X_c^{(\mathcal{M})}} (\Sigma_{l_2 l_2})^{-1}} \tag{339}$$

*is distributed according to*

$$\mathcal{T}_{\mathcal{M}} \sim \Gamma(\zeta_{\mathcal{M}}, 1) \tag{340}$$

$$\zeta_{\mathcal{M}} = \frac{1}{2}\left(d^{(X_0)} - d^{(X)}\right) \tag{341}$$

*for all $\mathcal{M} \neq \mathcal{M}_0$, where $\Gamma(\zeta_{\mathcal{M}}, 1)$ denotes the gamma distribution with shape parameter $\zeta_{\mathcal{M}}$ and scale parameter 1.*

*Proof.* We define $\boldsymbol{\rho} = \hat{\mathbf{b}} - \mathbf{b}$, substitute $\hat{\mathbf{b}}$ and obtain

$$\mathcal{T}_{\mathcal{M}} = \frac{1}{2} \sum_{c_0=1}^{d^{(X_0)}} \frac{\left(\sum_{k_1 \in X_{c_0}^{(\mathcal{M}_0)}} (b_{k_1} + \rho_{k_1}) (\Sigma_{k_1 k_1})^{-1}\right)^2}{\sum_{k_2 \in X_{c_0}^{(\mathcal{M}_0)}} (\Sigma_{k_2 k_2})^{-1}} - \tag{342}$$

$$\frac{1}{2} \sum_{c=1}^{d^{(X)}} \frac{\left(\sum_{l_1 \in X_c^{(\mathcal{M})}} (b_{l_1} + \rho_{l_1}) (\Sigma_{l_1 l_1})^{-1}\right)^2}{\sum_{l_2 \in X_c^{(\mathcal{M})}} (\Sigma_{l_2 l_2})^{-1}}. \tag{343}$$

According to Lemma 11, $\mathcal{T}_{\mathcal{M}}$ is independent of $\mathbf{b}$ and

$$\mathcal{T}_{\mathcal{M}} = \frac{1}{2} \sum_{c=1}^{d^{(X)}} \left( \sum_{c_0 \in K^{(c)}} \frac{\left(\sum_{k_1 \in X_{c_0}^{(\mathcal{M}_0)}} \rho_{k_1} (\Sigma_{k_1 k_1})^{-1}\right)^2}{\sum_{k_2 \in X_{c_0}^{(\mathcal{M}_0)}} (\Sigma_{k_2 k_2})^{-1}} - \frac{\left(\sum_{l_1 \in X_c^{(\mathcal{M})}} \rho_{l_1} (\Sigma_{l_1 l_1})^{-1}\right)^2}{\sum_{l_2 \in X_c^{(\mathcal{M})}} (\Sigma_{l_2 l_2})^{-1}} \right). \tag{344}$$

Because $X_c^{(\mathcal{M})}$ and $X_{c_0}^{(\mathcal{M}_0)}$ are concise index sets they consist of the same indices, i.e.

$$\bigcup_{c_0 \in \{1,...,d^{(X_0)}\}} X_{c_0}^{(\mathcal{M}_0)} = \bigcup_{c \in \{1,...,d^{(X)}\}} X_c^{(\mathcal{M})}, \tag{345}$$

and each index occurs in exactly one index set associated with $\mathcal{M}$ and $\mathcal{M}_0$, respectively. Furthermore, $\mathcal{M}$ is nested and $X_{c_0}^{(\mathcal{M}_0)}$, $c_0 \in \{1, ..., d^{(X_0)}\}$ are sets consisting of only one observable component. Therefore, for all $c \in \{1, ..., d^{(X)}\}$ there exists a set $K^{(c)} \subseteq \{1, ..., d^{(X_0)}\}$ such that[61]

$$X_c^{(\mathcal{M})} = \bigcup_{c_0 \in K^{(c)}} X_{c_0}^{(\mathcal{M}_0)}. \tag{346}$$

We define $\boldsymbol{\rho}^{(c)} \in \mathbb{R}^{d^{(X_0)}}$, where

$$\rho_{c_0}^{(c)} = \begin{cases} \dfrac{\sum_{k_1 \in X_{c_0}^{(\mathcal{M}_0)}} \rho_{k_1} (\Sigma_{k_1 k_1})^{-1}}{\left(\sum_{k_2 \in X_{c_0}^{(\mathcal{M}_0)}} (\Sigma_{k_2 k_2})^{-1}\right)^{1/2}}, & c_0 \in K^{(c)} \\ 0, & \text{otherwise,} \end{cases} \tag{347}$$

[61]The theorem still holds if $\mathcal{M}_0$ is replaced by any other modularization that is a refinement of $\mathcal{M}$. In case of $\mathcal{M}_0$, $K^{(c)} = X_c^{(\mathcal{M})}$, because $X_{c_0}^{(\mathcal{M}_0)} = \{c_0\}$ (see Equ. 50).

so that $\rho_{c_0}^{(c)}$ is standard normal for all $c_0 \in K^{(c)}$ and 0 otherwise.[62] Then,

$$\mathcal{T}_{\mathcal{M}} = \frac{1}{2}\sum_{c=1}^{d^{(X)}} \boldsymbol{\rho}^{(c)\,T}\,\mathbf{A}^{(c)}\boldsymbol{\rho}^{(c)} \tag{348}$$

$$\mathbf{A}^{(c)} = \mathbb{1} - \frac{\mathbf{a}^{(c)}\;\mathbf{a}^{(c)\,T}}{\mathbf{a}^{(c)\,T}\;\mathbf{a}^{(c)}} \tag{349}$$

$$a_{c_0}^{(c)} = \begin{cases} \sum_{k_2 \in X_{c_0}^{(\mathcal{M}_0)}} \left(\Sigma_{k_2 k_2}\right)^{-1/2}, & c_0 \in K^{(c)} \\ 0, & \text{otherwise,} \end{cases} \tag{350}$$

where $\mathbf{a}^{(c)} \in \mathbb{R}^{d^{(X_0)}}$. For each $c \in \{1, ..., d^{(X)}\}$ the matrix $\mathbf{A}^{(c)}$ projects $\boldsymbol{\rho}^{(c)}$ into a $|K^{(c)}| - 1$ dimensional sub-space perpendicular to $\mathbf{a}^{(c)}$ such that $\boldsymbol{\rho}^{(c)T}\mathbf{A}^{(c)}\boldsymbol{\rho}^{(c)}$ is the sum of the squares of $|K^{(c)}| - 1$ independent standard normal random variables. Thus, $\mathcal{T}_{\mathcal{M}}$ is the sum of the squares of $d^{(X_0)} - d^{(X)}$ independent standard normal random variables and distributed according to (Mathai and Provost, 1992)

$$\mathcal{T}_{\mathcal{M}} \sim \Gamma(\zeta_{\mathcal{M}}, 1) \tag{351}$$

$$\zeta_{\mathcal{M}} = \frac{1}{2}\left(d^{(X_0)} - d^{(X)}\right) \tag{352}$$

$$\mathrm{E}[\mathcal{T}_{\mathcal{M}}] = \zeta_{\mathcal{M}} \tag{353}$$

$$\mathrm{Var}[\mathcal{T}_{\mathcal{M}}] = \zeta_{\mathcal{M}}, \tag{354}$$

where $\Gamma(\zeta_{\mathcal{M}}, 1)$ denotes the gamma distribution with shape parameter $\zeta_{\mathcal{M}}$ and scale parameter 1.[63] □

[62] The theorem still holds if $\mathcal{M}_0$ is replaced by any other modularization that is a refinement of $\mathcal{M}$. In case of $\mathcal{M}_0$, Equ. 350 simplifies to $a_{c_0}^{(c)} = \left(\Sigma_{c_0 c_0}\right)^{-1/2}$ if $c_0 \in K^{(c)}$ and zero otherwise and Equ. 347 simplifies to $\rho_{c_0}^{(c)} = \rho_{c_0} a_{c_0}^{(c)}$ if $c_0 \in K^{(c)}$ and zero otherwise.

[63] The sum of independent normal random variables results in a chi-square distribution. Rescaling with a factor of 1/2 results in a gamma distribution. See also (Mathai and Provost, 1992, Equ. 4.1.2).

**Lemma 13.** *Let $\mathcal{M}$ denote a nested modularization. Furthermore, let $\hat{\mathbf{b}} \sim \mathcal{N}(\mathbf{b}, \mathbf{\Sigma})$, where $\mathcal{N}$ denotes the normal distribution with arbitrary full-rank diagonal covariance matrix $\mathbf{\Sigma}$ and mean $\mathbf{b} \in \mathbb{R}^{d^{(X_0)}}$ such that $b_i = b_j$ for all $i, j \in X_c^{(\mathcal{M})}$ and all $c \in \{1, ..., d^{(X)}\}$.*

*If the prior distribution of $\mathbf{b}$ is non-informative, then*

$$\mathcal{T}_{\mathcal{M}} = \text{logit}_{\mathcal{M}_0}(\mathcal{M}) + \zeta_{\mathcal{M}} \tag{355}$$

$$\zeta_{\mathcal{M}} = \frac{1}{2}\left(d^{(X_0)} - d^{(X)}\right) \tag{356}$$

*for log odd*

$$\text{logit}_{\mathcal{M}_0}(\mathcal{M}) = \log \frac{\text{P}(\mathcal{M}_0|\hat{\mathbf{b}}, \mathbf{\Sigma})}{\text{P}(\mathcal{M}|\hat{\mathbf{b}}, \mathbf{\Sigma})}. \tag{357}$$

*Proof.* Let $p(\hat{\mathbf{b}}, \mathbf{\Sigma}|\mathcal{M})$ denote the joint probability density function of $\hat{\mathbf{b}}$ and $\mathbf{\Sigma}$ conditioned on $\mathcal{M}$.[64] The posterior probability of $\mathcal{M}$ given $\hat{\mathbf{b}}$ and $\mathbf{\Sigma}$ can be inferred from Bayes' theorem, i.e.,

$$\begin{aligned} \text{P}(\mathcal{M}|\hat{\mathbf{b}}, \mathbf{\Sigma}) &\propto p(\hat{\mathbf{b}}|\mathcal{M}, \mathbf{\Sigma})\text{P}(\mathcal{M}|\mathbf{\Sigma}) & (358)\\ &= \int p(\hat{\mathbf{b}}, \mathbf{b}|\mathcal{M}, \mathbf{\Sigma})\text{P}(\mathcal{M}|\mathbf{\Sigma})\text{d}\mathbf{b} & (359)\\ &= \int p(\hat{\mathbf{b}}|\mathbf{b}, \mathcal{M}, \mathbf{\Sigma})p(\mathbf{b}|\mathcal{M}, \mathbf{\Sigma})\text{P}(\mathcal{M}|\mathbf{\Sigma})\text{d}\mathbf{b}, & (360) \end{aligned}$$

where

$$p(\hat{\mathbf{b}}|\mathbf{b}, \mathcal{M}, \mathbf{\Sigma}) = p_{\mathcal{N}}(\hat{\mathbf{b}}|\mathbf{b}, \mathbf{\Sigma}) \tag{361}$$

and $p_{\mathcal{N}}(\hat{\mathbf{b}}|\mathbf{b}, \mathbf{\Sigma})$ denotes the probability density function of a normally distributed random variable $\hat{\mathbf{b}}$ with mean $\mathbf{b}$ and covariance matrix $\mathbf{\Sigma}$ defined by

$$p_{\mathcal{N}}(\hat{\mathbf{b}}|\mathbf{b}, \mathbf{\Sigma}) = \frac{1}{|2\pi\mathbf{\Sigma}|^{1/2}} \, e^{-\frac{1}{2}\sum_{i,j=1}^{d^{(X_0)}}(\hat{b}_i - b_i)(\Sigma^{-1})_{ij}(\hat{b}_j - b_j)}. \tag{362}$$

We first define the priors $p(\mathbf{b}|\mathcal{M}, \mathbf{\Sigma})$ and $\text{P}(\mathcal{M}|\mathbf{\Sigma})$ and calculate the posterior probability $\text{P}(\mathcal{M}|\hat{\mathbf{b}}, \mathbf{\Sigma})$ in Equ. 360 by integrating over $\mathbf{b}$. Subsequently, we modify both priors to become non-informative and obtain Equ. 355, where we call $\text{P}(\mathcal{M}|\mathbf{\Sigma})$ non-informative if

$$E[\text{logit}_{\mathcal{M}_0}(\mathcal{M})] = 0 \tag{363}$$

for all $\mathcal{M}$ that are consistent with $\mathbf{b}$, i.e., $b_i = b_j$ for all $i, j \in X_c^{(\mathcal{M})}$ and all $c \in \{1, ..., d^{(X)}\}$.

More precisely, from $\mathcal{M}_0 \leq \mathcal{M}$ follows $X_{c_0}^{(\mathcal{M}_0)} \subseteq X_c^{(\mathcal{M})}$ for all $c_0 \in \{1, ..., d^{(X_0)}\}$ and some $c \in \{1, ..., d^{(X)}\}$. Furthermore, each concise index set associated with $\mathcal{M}$ consists of one or multiple concise index sets associated with $\mathcal{M}_0$. Thus, for all $c \in \{1, ..., d^{(X)}\}$ there exists a set $K^{(c)} \subseteq \{1, ..., d^{(X_0)}\}$ such that

$$X_c^{(\mathcal{M})} = \bigcup_{c_0 \in K^{(c)}} X_{c_0}^{(\mathcal{M}_0)}. \tag{364}$$

By assumption, $b_i = b_j = b_c^{(\mathcal{M})}$ for all $i \in X_{c_0}^{(\mathcal{M}_0)}$, all $j \in X_c^{(\mathcal{M})}$, all $c_0 \in K^{(c)}$, all $c \in \{1, ..., d^{(X)}\}$ and $b_c^{(\mathcal{M})} \in \mathbb{R}$.

[64]To simplify notation, we simply write $p(\hat{\mathbf{b}})$ to denote the probability density function over the random variable $\hat{\mathbf{b}}$. Thus, the symbol $p$ is overloaded and used for different density functions, provided that the interpretation is clear from the context.

Then, the likelihood of $\mathbf{b}$ can be expressed in terms of $\mathbf{b}^{(\mathcal{M})}$

$$
\begin{aligned}
p(\hat{\mathbf{b}}|\mathbf{b},\mathcal{M},\boldsymbol{\Sigma}) &= \frac{1}{|2\pi\boldsymbol{\Sigma}|^{1/2}}\, e^{-\frac{1}{2}\sum_{i,j=1}^{d^{(\mathcal{X}_0)}}(\hat{b}_i-b_i)(\Sigma^{-1})_{ij}(\hat{b}_j-b_j)} && (365)\\
&= \frac{1}{|2\pi\boldsymbol{\Sigma}|^{1/2}}\, e^{-\frac{1}{2}\sum_{k,l=1}^{d^{(\mathcal{X})}}\sum_{i\in K^{(k)}}\sum_{j\in K^{(l)}}(\hat{b}_i-b_k^{(\mathcal{M})})(\Sigma^{-1})_{ij}(\hat{b}_j-b_l^{(\mathcal{M})})} && (366)\\
&= \frac{1}{|2\pi\boldsymbol{\Sigma}|^{1/2}}\, e^{-\frac{1}{2}\sum_{k,l}\sum_{i,j}(\hat{b}_i-b_k^{(\mathcal{M})})(\Sigma^{-1})_{ij}(\hat{b}_j-b_l^{(\mathcal{M})})}, && (367)
\end{aligned}
$$

where we omit for notational clarity in Equ. 367 and the rest of this section the limits and sets of summations provided that $k,l\in\{1,...,d^{(\mathcal{X})}\}$, $i\in K^{(k)}$ and $j\in K^{(l)}$. Furthermore, the exponent in Equ. 367 can be rearranged so that $p(\hat{\mathbf{b}}|\mathbf{b},\mathcal{M},\boldsymbol{\Sigma})$ can be expressed in terms of the probability density function of the normally distributed $\mathbf{b}^{(\mathcal{M})}$, i.e.,

$$
\begin{aligned}
p(\hat{\mathbf{b}}|\mathbf{b},\mathcal{M},\boldsymbol{\Sigma}) &= \frac{1}{|2\pi\boldsymbol{\Sigma}|^{1/2}} e^{-\frac{1}{2}\sum_{k,l}\alpha_{kl}-\frac{1}{2}\sum_{k,l}b_k^{(\mathcal{M})}(\Sigma'^{-1})_{kl}b_l^{(\mathcal{M})}+\frac{1}{2}\sum_l\beta_l b_l^{(\mathcal{M})}+\frac{1}{2}\sum_k b_k^{(\mathcal{M})}\beta'_k} && (368)\\
&= \frac{1}{|2\pi\boldsymbol{\Sigma}|^{1/2}} e^{-\frac{1}{2}\sum_{k,l}\left(\alpha_{kl}-\gamma_k(\Sigma'^{-1})_{kl}\gamma_l\right)}\, e^{-\frac{1}{2}\sum_{k,l}(b_k^{(\mathcal{M})}-\gamma_k)(\Sigma'^{-1})_{kl}(b_l^{(\mathcal{M})}-\gamma_l)} \\
&= c_c\, p_{\mathcal{N}}(\mathbf{b}^{(\mathcal{M})}|\gamma,\boldsymbol{\Sigma}') && (369)
\end{aligned}
$$

with prefactor

$$
c_c = \frac{|2\pi\boldsymbol{\Sigma}'|^{1/2}}{|2\pi\boldsymbol{\Sigma}|^{1/2}} e^{-\frac{1}{2}\sum_{k,l}\left(\alpha_{kl}-\gamma_k(\Sigma'^{-1})_{kl}\gamma_l\right)}, \qquad (370)
$$

where we define[65]

$$
\begin{aligned}
(\Sigma'^{-1})_{kl} &= \sum_{i,j}(\Sigma^{-1})_{ij} = (\Sigma'^{-1})_{lk} && (371)\\
\alpha_{kl} &= \sum_{i,j}\hat{b}_i(\Sigma^{-1})_{ij}\hat{b}_j && (372)\\
\beta_l &= \sum_k\sum_{i,j}\hat{b}_i(\Sigma^{-1})_{ij} && (373)\\
\beta'_k &= \sum_l\sum_{i,j}(\Sigma^{-1})_{ij}\hat{b}_j = \beta_k && (374)\\
\gamma_k &= \sum_l\beta_l\Sigma'_{lk}, && (375)
\end{aligned}
$$

use the relations

$$
\begin{aligned}
\sum_{k,l}\gamma_k(\Sigma'^{-1})_{kl}b_l^{(\mathcal{M})} &= \sum_{k,l}\sum_o\beta_o\Sigma'_{ok}(\Sigma'^{-1})_{kl}b_l^{(\mathcal{M})} = \sum_l\sum_o\beta_o\delta_{ol}b_l^{(\mathcal{M})} = \sum_l\beta_l b_l^{(\mathcal{M})} && (376)\\
\sum_{k,l}b_k^{(\mathcal{M})}(\Sigma'^{-1})_{kl}\gamma_l &= \sum_{k,l}\sum_o b_k^{(\mathcal{M})}(\Sigma'^{-1})_{kl}\Sigma'_{lo}\beta_o = \sum_k\sum_o b_k^{(\mathcal{M})}\delta_{ko}\beta_o = \sum_k b_k^{(\mathcal{M})}\beta_k, && (377)
\end{aligned}
$$

and denote the Kronecker delta function by $\delta_{ol}$, which is 1 if $o=l$ and zero otherwise.

We choose a normally distributed prior for $\mathbf{b}^{(\mathcal{M})}$

$$
p(\mathbf{b}|\mathcal{M},\boldsymbol{\Sigma}) = p_{\mathcal{N}}(\mathbf{b}^{(\mathcal{M})}|\mathbf{0},\boldsymbol{\Sigma}^{(b)}), \quad \boldsymbol{\Sigma}^{(b)} = \sigma^2\mathbb{1}, \quad \sigma>0 \qquad (378)
$$

[65] $\beta'_k=\beta_k$ (Equ. 374) and $\boldsymbol{\Sigma}'^{-1}=(\boldsymbol{\Sigma}'^{-1})^T$ because if $\boldsymbol{\Sigma}$ is non-singular and symmetric then so is $\boldsymbol{\Sigma}^{-1}$.

which is a conjugate prior for the likelihood of $\mathbf{b}$. Then, the product

$$p(\hat{\mathbf{b}}|\mathbf{b},\mathcal{M},\mathbf{\Sigma})p(\mathbf{b}|\mathcal{M},\mathbf{\Sigma}) = c_c p_{\mathcal{N}}(\mathbf{b}^{(\mathcal{M})}|\gamma,\mathbf{\Sigma}')p_{\mathcal{N}}(\mathbf{b}^{(\mathcal{M})}|\mathbf{0},\mathbf{\Sigma}^{(b)}) \quad (379)$$

can be obtained from the relation

$$\begin{aligned}
p_{\mathcal{N}}(\mathbf{m}|\mathbf{m}_1,\mathbf{\Sigma}_1)p_{\mathcal{N}}(\mathbf{m}|\mathbf{m}_2,\mathbf{\Sigma}_2) &= c_d p_{\mathcal{N}}(\mathbf{m}|\mathbf{m}_3,\mathbf{\Sigma}_3) && (380)\\
c_d &= p_{\mathcal{N}}(\mathbf{m}_1|\mathbf{m}_2,\mathbf{\Sigma}_1+\mathbf{\Sigma}_2) && (381)\\
\mathbf{\Sigma}_3 &= (\mathbf{\Sigma}_1^{-1}+\mathbf{\Sigma}_2^{-1})^{-1} && (382)\\
\mathbf{m}_3 &= \mathbf{\Sigma}_3(\mathbf{\Sigma}_1^{-1}\mathbf{m}_1+\mathbf{\Sigma}_2^{-1}\mathbf{m}_2). && (383)
\end{aligned}$$

Integrating over $\mathbf{m}$ in Equ. 380 results in $c_d$, and, thus integrating over $\mathbf{b}$ in Equ. 360 results in the posterior probability

$$\begin{aligned}
\mathrm{P}(\mathcal{M}|\hat{\mathbf{b}},\mathbf{\Sigma}) &\propto c_c c_d \mathrm{P}(\mathcal{M}|\mathbf{\Sigma}) && (384)\\
&\propto \frac{|2\pi\mathbf{\Sigma}'|^{1/2}}{|2\pi(\mathbf{\Sigma}'+\mathbf{\Sigma}^{(b)})|^{1/2}} e^{-\frac{1}{2}\gamma^T[(\mathbf{\Sigma}'+\mathbf{\Sigma}^{(b)})^{-1}-\mathbf{\Sigma}'^{-1}]\gamma}\mathrm{P}(\mathcal{M}|\mathbf{\Sigma}) && (385)\\
&= \frac{1}{|\mathbb{1}+\mathbf{\Sigma}^{(b)}\mathbf{\Sigma}'^{-1}|^{1/2}} e^{-\frac{1}{2}\gamma^T[(\mathbf{\Sigma}'+\mathbf{\Sigma}^{(b)})^{-1}-\mathbf{\Sigma}'^{-1}]\gamma}\mathrm{P}(\mathcal{M}|\mathbf{\Sigma}), && (386)
\end{aligned}$$

where the terms $|2\pi\mathbf{\Sigma}|^{1/2}$ and $\alpha_{kl}$ have been dropped because their multiplicative contribution to the posterior is independent of the modularization $\mathcal{M}$. For diagonal covariance matrices $\mathbf{\Sigma}$ and $\mathbf{\Sigma}^{(b)}$ we obtain

$$\begin{aligned}
-\gamma^T[(\mathbf{\Sigma}'+\mathbf{\Sigma}^{(b)})^{-1}-\mathbf{\Sigma}'^{-1}]\gamma &= -\sum_k \gamma_k^2\left[\frac{1}{\Sigma'_{kk}+\sigma^2}-(\Sigma'^{-1})_{kk}\right] && (387)\\
&= -\sum_k \beta_k^2\left[\frac{(\Sigma'_{kk})^2}{\Sigma'_{kk}+\sigma^2}-\Sigma'_{kk}\right] && (388)\\
&= \sum_k \beta_k^2\frac{\Sigma'_{kk}\sigma^2}{\Sigma'_{kk}+\sigma^2} && (389)\\
&= \sum_k \frac{\beta_k^2}{1/\sigma^2+(\Sigma'^{-1})_{kk}} && (390)
\end{aligned}$$

and

$$\mathrm{P}(\mathcal{M}|\hat{\mathbf{b}},\mathbf{\Sigma}) \propto \frac{1}{\prod_k(1+\sigma^2(\Sigma'^{-1})_{kk})^{1/2}} e^{\frac{1}{2}\sum_k \frac{\beta_k^2}{1/\sigma^2+(\Sigma'^{-1})_{kk}}}\mathrm{P}(\mathcal{M}|\mathbf{\Sigma}). \quad (391)$$

The prior $p(\mathbf{b}|\mathcal{M},\mathbf{\Sigma})$ becomes non-informative for $\sigma^2 \to \infty$. However, for finite $\mathrm{P}(\mathcal{M}|\mathbf{\Sigma})$ the term before the exponential function in the last equation converges to zero with a leading term proportional to $\sigma^{-d^{(X)}}$ and only $\mathcal{M}$ with the smallest number of index sets $d^{(X)}$ have non-zero posterior probability. To obtain non-informative priors for $\mathcal{M}$, we demand

$$E[\mathrm{logit}_{\mathcal{M}_0}(\mathcal{M})] = 0 \quad (392)$$

for all $\mathcal{M}$ that are consistent with $\mathbf{b}$, i.e., $b_i = b_j$ for all $i, j \in X_c^{(\mathcal{M})}$ and all $c \in \{1, ..., d^{(X)}\}$.

With the ansatz

$$\mathrm{P}(\mathcal{M}|\boldsymbol{\Sigma}) \quad \propto \quad f^{(\mathrm{pr})}(\mathcal{M}) \prod_{k=1}^{d^{(X)}} (1 + \sigma^2 (\Sigma'^{-1})_{kk})^{1/2}, \tag{393}$$

where $f^{(\mathrm{pr})}(\mathcal{M})$ denotes an arbitrary positive non-zero real-valued function of $\mathcal{M}$, we obtain for diagonal covariance matrices $\boldsymbol{\Sigma}$

$$\mathrm{logit}_{\mathcal{M}_0}(\mathcal{M}) \quad = \quad \frac{1}{2} \sum_{k,i} \frac{(\hat{b}_i (\Sigma^{-1})_{ii})^2}{1/\sigma^2 + (\Sigma^{-1})_{ii}} + \log\left(f^{(\mathrm{pr})}(\mathcal{M}_0)\right) - \tag{394}$$

$$\frac{1}{2} \sum_{k} \frac{(\sum_i \hat{b}_i (\Sigma^{-1})_{ii})^2}{1/\sigma^2 + \sum_i (\Sigma^{-1})_{ii}} - \log\left(f^{(\mathrm{pr})}(\mathcal{M})\right). \tag{395}$$

According to Theorem 12,

$$\mathrm{logit}_{\mathcal{M}_0}(\mathcal{M}) \quad = \quad \frac{1}{2} \sum_{k,i} \frac{((b_k^{(\mathcal{M})} + \rho_i)(\Sigma^{-1})_{ii})^2}{1/\sigma^2 + (\Sigma^{-1})_{ii}} + \log\left(f^{(\mathrm{pr})}(\mathcal{M}_0)\right) - \tag{396}$$

$$\frac{1}{2} \sum_{k} \frac{(\sum_i (b_k^{(\mathcal{M})} + \rho_i)(\Sigma^{-1})_{ii})^2}{1/\sigma^2 + \sum_i (\Sigma^{-1})_{ii}} - \log\left(f^{(\mathrm{pr})}(\mathcal{M})\right) \tag{397}$$

$$\lim_{\sigma^2 \to \infty} \mathrm{logit}_{\mathcal{M}_0}(\mathcal{M}) \quad = \quad \mathcal{T}_{\mathcal{M}} + \log\left(f^{(\mathrm{pr})}(\mathcal{M}_0)\right) - \log\left(f^{(\mathrm{pr})}(\mathcal{M})\right) \tag{398}$$

and for non-informative priors $p(\mathbf{b}|\mathcal{M}, \boldsymbol{\Sigma})$ follows

$$E[\mathrm{logit}_{\mathcal{M}_0}(\mathcal{M})] \quad = \quad E[\mathcal{T}_{\mathcal{M}}] + \log\left(f^{(\mathrm{pr})}(\mathcal{M}_0)\right) - \log\left(f^{(\mathrm{pr})}(\mathcal{M})\right) \tag{399}$$

$$= \quad \frac{1}{2}\left(d^{(X_0)} - d^{(X)}\right) + \log\left(f^{(\mathrm{pr})}(\mathcal{M}_0)\right) - \log\left(f^{(\mathrm{pr})}(\mathcal{M})\right). \tag{400}$$

Thus, the priors for $\mathcal{M}$ are only non-informative if we identify

$$f^{(\mathrm{pr})}(\mathcal{M}) = e^{-\frac{d^{(X)}}{2}} \tag{401}$$

so that

$$E[\mathrm{logit}_{\mathcal{M}_0}(\mathcal{M})] = 0 \tag{402}$$

for all $\mathcal{M}$ that are consistent with $\mathbf{b}$, i.e., $b_i = b_j$ for all $i, j \in X_c^{(\mathcal{M})}$ and all $c \in \{1, ..., d^{(X)}\}$. Because the covariance matrix $\boldsymbol{\Sigma}$ is diagonal and has full-rank, $(\Sigma^{-1})_{ii} = (\Sigma_{ii})^{-1}$ and we obtain

$$\mathcal{T}_{\mathcal{M}} \quad = \quad \mathrm{logit}_{\mathcal{M}_0}(\mathcal{M}) + \zeta_{\mathcal{M}} \tag{403}$$

$$\zeta_{\mathcal{M}} \quad = \quad \frac{1}{2}\left(d^{(X_0)} - d^{(X)}\right). \tag{404}$$

□

**Lemma 14.** *Let $\mathcal{M}$ denote a flat modularization. Furthermore, let $\sqrt{N}\left(\hat{\mathbf{b}} - \mathbf{b}\right) \xrightarrow{d} \mathcal{N}(\mathbf{0}, \mathbf{\Sigma}')$ as $N \to \infty$ for $\mathbf{b} \in \mathbb{R}^{d^{(X_0)}}$ and diagonal full-rank covariance matrix $\mathbf{\Sigma}'$.*

*Then, the statistical test based on $\mathcal{T}_{\mathcal{M}}(\hat{\mathbf{b}}, \mathbf{\Sigma})$ is consistent.*

*Proof.* In general, the observable states form multiple modularizations. E.g., if the observable states form a flat modularization $\mathcal{M}_1$, then any modularization $\mathcal{M}_2 \subset \mathcal{M}_1 = \{S_1, ..., S_{d^{(\mathcal{M})}}\}$ and $\mathcal{M}_0$ are true modularizations as well. Let $\mathcal{H} = \{\mathcal{M}_1, \mathcal{M}_2, ...\}$ denote the finite set of all true flat modularizations. We test for $\mathcal{M} \in \mathcal{H}$ versus $\mathcal{M} \notin \mathcal{H}$ based on $\mathcal{T}_{\mathcal{M}}$.

A sequence of statistical tests is consistent if it is asymptotically of level $\alpha$ for all levels $\alpha \in (0, 1]$ and the power of all alternative hypotheses converges to 1.[66] By assumption, $\sqrt{N}\left(\hat{\mathbf{b}} - \mathbf{b}\right) \xrightarrow{d} \mathcal{N}(\mathbf{0}, \mathbf{\Sigma})$ as $N \to \infty$ and, by Theorem 12,

$$\mathcal{T}_{\mathcal{M}}(\hat{\mathbf{b}}, \mathbf{\Sigma}) \xrightarrow{d} \Gamma(\xi, \mathcal{M}) \tag{405}$$

as $N \to \infty$ for all $\mathcal{M} \in \mathcal{H}$ and $\zeta_{\mathcal{M}} = \frac{1}{2}\left(d^{(X_0)} - d^{(X)}\right)$. Therefore, for given $\alpha \in (0, 1]$, we can always choose a sufficiently large $t > 0$ such that the power of the test is asymptotically of level $\alpha$, i.e.,

$$\lim_{N\to\infty} \mathrm{P}(\mathcal{T}_{\mathcal{M}} > t | \mathcal{M}_1) \leq \alpha \tag{406}$$

for all $\mathcal{M} \in \mathcal{H}$. We now show, for any fixed $t > 0$, that the power of the test converges for all incorrect alternatives to one, i.e., $\lim_{N\to\infty} \mathrm{P}(\mathcal{T}_{\mathcal{M}} > t | \mathcal{M}_1) = 1$ for all $\mathcal{M} \notin \mathcal{H}$.

For fixed $N$, let $\boldsymbol{\rho}' = \sqrt{N}(\hat{\mathbf{b}} - \mathbf{b})$ and $\boldsymbol{\epsilon} = \mathbf{b} - \bar{\mathbf{b}}$, where $\bar{\mathbf{b}} \in \mathbb{R}^{d^{(X_0)}}$ denotes an arbitrary common mean consistent with $\mathcal{M}_1$, i.e., $\bar{b}_k = \bar{b}_l$ for all $k, l \in X_c^{(\mathcal{M}_1)}$ and all $c \in \{1, ..., d^{(X_1)}\}$. Then,[67]

$$\begin{aligned}
\mathcal{T}_{\mathcal{M}}\left(\hat{\mathbf{b}}, \mathbf{\Sigma}\right) &= \mathcal{T}_{\mathcal{M}}\left(\sqrt{N}\hat{\mathbf{b}}, N\mathbf{\Sigma}\right) && (407)\\
&= \mathcal{T}_{\mathcal{M}}\left(\sqrt{N}(\hat{\mathbf{b}} - \bar{\mathbf{b}}), \mathbf{\Sigma}'\right) && (408)\\
&= \mathcal{T}_{\mathcal{M}}\left(\boldsymbol{\rho}' + \sqrt{N}\boldsymbol{\epsilon}, \mathbf{\Sigma}'\right) && (409)
\end{aligned}$$

and, analogous to Theorem 12 (see Equ. 348),

$$\mathcal{T}_{\mathcal{M}}(\hat{\mathbf{b}}, \mathbf{\Sigma}) = \frac{1}{2}\sum_{c=1}^{d^{(X)}} \left(\boldsymbol{\rho}^{(c)} + \sqrt{N}\boldsymbol{\epsilon}^{(c)}\right)^T \mathbf{A}^{(c)} \left(\boldsymbol{\rho}^{(c)} + \sqrt{N}\boldsymbol{\epsilon}^{(c)}\right) \tag{410}$$

$$\rho_{c_0}^{(c)} = \begin{cases} \rho'_{c_0} a_{c_0}^{(c)}, & c_0 \in X_c^{(\mathcal{M})} \\ 0, & \text{otherwise,} \end{cases} \tag{411}$$

$$\epsilon_{c_0}^{(c)} = \begin{cases} \epsilon_{c_0} a_{c_0}^{(c)}, & c_0 \in X_c^{(\mathcal{M})} \\ 0, & \text{otherwise,} \end{cases} \tag{412}$$

$$\mathbf{A}^{(c)} = \mathbb{1} - \frac{\mathbf{a}^{(c)}\, \mathbf{a}^{(c)\,T}}{\mathbf{a}^{(c)\,T}\, \mathbf{a}^{(c)}} \tag{413}$$

$$a_{c_0}^{(c)} = \begin{cases} \left(\Sigma'_{c_0 c_0}\right)^{-1/2}, & c_0 \in X_c^{(\mathcal{M})} \\ 0, & \text{otherwise,} \end{cases} \tag{414}$$

[66](Van der Vaart, 2000, definition 14.2)

[67]According to Lemma 11, $\mathcal{T}_{\mathcal{M}}$ is independent of arbitrary common means $\bar{\mathbf{b}}$.

where $\mathbf{a}^{(c)} \in \mathbb{R}^{d^{(X_0)}}$.

A flat modularization $\mathcal{M}$ is incorrect, i.e., $\mathcal{M} \notin \mathcal{H}$, if and only if there exists a $c' \in \{1, ..., d^{(X_1)}\}$ and a deviation $\boldsymbol{\epsilon}^{(c')}$ such that $d^{(c')} = \left|\mathbf{A}^{(c')}\boldsymbol{\epsilon}^{(c')}\right| > 0$. That is, $\boldsymbol{\epsilon}^{(c')}$ cannot be exclusively a deviation by a common mean, otherwise $\mathcal{M} \in \mathcal{H}$. For notational clarity and without loss of generality, we assume $\boldsymbol{\epsilon}^{(c)} = \mathbf{0}$ for all $c \neq c'$. Then,

$$
\begin{aligned}
\mathcal{T}_{\mathcal{M}} &= \frac{1}{2}\sum_{c=1}^{d^{(X)}}(\boldsymbol{\rho}^{(c)} + \sqrt{N}\boldsymbol{\epsilon}^{(c)})^T\mathbf{A}^{(c)}(\boldsymbol{\rho}^{(c)} + \sqrt{N}\boldsymbol{\epsilon}^{(c)}) && (415)\\
&= \frac{1}{2}\left(N\boldsymbol{\epsilon}^{(c')^T}\mathbf{A}^{(c')}\boldsymbol{\epsilon}^{(c')} + 2\sqrt{N}\boldsymbol{\epsilon}^{(c')^T}\mathbf{A}^{(c')}\boldsymbol{\rho}^{(c')} + 2\mathcal{T}^{(0)}_{\mathcal{M}}\right) && (416)\\
&= \frac{1}{2}\left(N\boldsymbol{\epsilon}^{(c')^T}\mathbf{A}^{(c')^T}\mathbf{A}^{(c')}\boldsymbol{\epsilon}^{(c')} + 2\sqrt{N}\boldsymbol{\epsilon}^{(c')^T}\mathbf{A}^{(c')}\boldsymbol{\rho}^{(c')} + 2\mathcal{T}^{(0)}_{\mathcal{M}}\right) && (417)\\
&= N\left(\frac{d^{(c')^2}}{2} + \frac{1}{\sqrt{N}}\boldsymbol{\epsilon}^{(c')^T}\mathbf{A}^{(c')}\boldsymbol{\rho}^{(c')} + \frac{1}{N}\mathcal{T}^{(0)}_{\mathcal{M}}\right), && (418)
\end{aligned}
$$

where

$$
\mathcal{T}^{(0)}_{\mathcal{M}} = \frac{1}{2}\sum_{c=1}^{d^{(X)}}\boldsymbol{\rho}^{(c)^T}\mathbf{A}^{(c)}\boldsymbol{\rho}^{(c)} \tag{419}
$$

and $\mathcal{T}^{(0)}_{\mathcal{M}} \xrightarrow{d} \Gamma(\xi, \mathcal{M})$ as $N \to \infty$.

For sequences of random variables $\hat{\mathbf{b}}$, $\boldsymbol{\rho}^{(c)}$ and $\mathcal{T}_{\mathcal{M}}$ indexed by $N$,[68]

$$
\frac{1}{N}\mathcal{T}_{\mathcal{M}} \xrightarrow{d} \frac{d^{(c')^2}}{2} \tag{420}
$$

as $N \to \infty$ and

$$
\begin{aligned}
\lim_{N\to\infty} \mathrm{P}(\mathcal{T}_{\mathcal{M}} > t|\mathcal{M}_1) &= \lim_{N\to\infty} \mathrm{P}\left(\tfrac{1}{N}\mathcal{T}_{\mathcal{M}} > \tfrac{t}{N}\middle|\mathcal{M}_1\right) && (421)\\
&\geq \mathrm{P}\left(\tfrac{d^{(c')^2}}{2} > \tfrac{d^{(c')^2}}{4}\middle|\mathcal{M}_1\right) && (422)\\
&= 1 && (423)
\end{aligned}
$$

for all $t > 0$ and all $\mathcal{M} \notin \mathcal{H}$. □

[68] According to the continuous mapping theorem (Van der Vaart, 2000, theorem 2.3).

**Lemma 15.** *Let $\mathcal{M}_1$ and $\mathcal{M}_2$ denote two nested modularizations. Furthermore, let $\hat{\mathbf{b}} \sim \mathcal{N}(\mathbf{b}, \boldsymbol{\Sigma})$, where $\mathcal{N}$ denotes the normal distribution with arbitrary full-rank diagonal covariance matrix $\boldsymbol{\Sigma}$ and arbitrary mean $\mathbf{b} \in \mathbb{R}^{d^{(X_0)}}$. If $\mathcal{M}_1 \leq \mathcal{M}_2$, then*

$$E\left[\mathcal{T}_{\mathcal{M}_1}(\hat{\mathbf{b}}, \boldsymbol{\Sigma}) + \frac{d^{(X_1)}}{2}\right] \leq E\left[\mathcal{T}_{\mathcal{M}_2}(\hat{\mathbf{b}}, \boldsymbol{\Sigma}) + \frac{d^{(X_2)}}{2}\right]. \tag{424}$$

*Proof.* We define $\boldsymbol{\rho} = \hat{\mathbf{b}} - \mathbf{b}$, so that $\boldsymbol{\rho} \sim \mathcal{N}(\mathbf{0}, \boldsymbol{\Sigma})$, and obtain from Equ. 61

$$E\left[\mathcal{T}_{\mathcal{M}_2} + \frac{d^{(X_2)}}{2}\right] - E\left[\mathcal{T}_{\mathcal{M}_1} + \frac{d^{(X_1)}}{2}\right] = \frac{1}{2}\sum_{c_1=1}^{d^{(X_1)}} \frac{\left(\sum_{l_1 \in X_{c_1}^{(\mathcal{M}_1)}} b_{l_1} \, (\Sigma_{l_1 l_1})^{-1}\right)^2}{\sum_{l_2 \in X_{c_1}^{(\mathcal{M}_1)}} (\Sigma_{l_2 l_2})^{-1}} - \tag{425}$$

$$\frac{1}{2}\sum_{c_2=1}^{d^{(X_2)}} \frac{\left(\sum_{l_3 \in X_{c_2}^{(\mathcal{M}_2)}} b_{l_3} \, (\Sigma_{l_3 l_3})^{-1}\right)^2}{\sum_{l_4 \in X_{c_2}^{(\mathcal{M}_2)}} (\Sigma_{l_4 l_4})^{-1}}, \tag{426}$$

where the contributions of the noise $\boldsymbol{\rho}$ to the expectation values are canceled by the terms $d^{(X_1)}/2$ and $d^{(X_2)}/2$.

From $\mathcal{M}_1 \leq \mathcal{M}_2$ follows $X_{c_1}^{(\mathcal{M}_1)} \subseteq X_{c_2}^{(\mathcal{M}_2)}$ for all $c_1 \in \{1, ..., d^{(X_1)}\}$ and some $c_2 \in \{1, ..., d^{(X_2)}\}$. Furthermore, each concise index set associated with $\mathcal{M}_2$ consists of one or multiple concise index sets associated with $\mathcal{M}_1$. We define $K^{(c_2)} \subseteq \{1, ..., d^{(X_1)}\}, c_2 \in \{1, ..., d^{(X_2)}\}$ such that

$$X_{c_2}^{(\mathcal{M}_2)} = \bigcup_{c_1 \in K^{(c_2)}} X_{c_1}^{(\mathcal{M}_1)}. \tag{427}$$

Then,

$$E\left[\mathcal{T}_{\mathcal{M}_2} + \frac{d^{(X_2)}}{2}\right] - E\left[\mathcal{T}_{\mathcal{M}_1} + \frac{d^{(X_1)}}{2}\right] = \frac{1}{2}\sum_{c_2=1}^{d^{(X_2)}}\left(\sum_{c_1 \in K^{(c_2)}} \frac{\varphi_{c_1}^2}{\phi_{c_1}} - \frac{\left(\sum_{c_1 \in K^{(c_2)}} \varphi_{c_1}\right)^2}{\sum_{c_1 \in K^{(c_2)}} \phi_{c_1}}\right) \tag{428}$$

$$\varphi_{c_1} = \sum_{l_1 \in X_{c_1}^{(\mathcal{M}_1)}} b_{l_1} \, (\Sigma_{l_1 l_1})^{-1} \tag{429}$$

$$\phi_{c_1} = \sum_{l_2 \in X_{c_1}^{(\mathcal{M}_1)}} (\Sigma_{l_2 l_2})^{-1}. \tag{430}$$

According to Jensen's inequality

$$\left(\frac{\sum_{c_1} \phi_{c_1} \frac{\varphi_{c_1}}{\phi_{c_1}}}{\sum_{c_1} \phi_{c_1}}\right)^2 \leq \frac{\sum_{c_1} \phi_{c_1} \left(\frac{\varphi_{c_1}}{\phi_{c_1}}\right)^2}{\sum_{c_1} \phi_{c_1}} \tag{431}$$

$$\frac{\left(\sum_{c_1} \varphi_{c_1}\right)^2}{\sum_{c_1} \phi_{c_1}} \leq \sum_{c_1} \frac{\varphi_{c_1}^2}{\phi_{c_1}}, \tag{432}$$

where $\phi_{c_1} > 0$, and

$$E\left[\mathcal{T}_{\mathcal{M}_2} + \frac{d^{(X_2)}}{2}\right] - E\left[\mathcal{T}_{\mathcal{M}_1} + \frac{d^{(X_1)}}{2}\right] \geq 0 \tag{433}$$

$$E\left[\mathcal{T}_{\mathcal{M}_1} + \frac{d^{(X_1)}}{2}\right] \leq E\left[\mathcal{T}_{\mathcal{M}_2} + \frac{d^{(X_2)}}{2}\right]. \tag{434}$$

□

**Lemma 16.** *Let* $\boldsymbol{\Sigma}^{(v)} \in \mathbb{R}^{2\times 2}$ *such that* $\Sigma_{11}^{(v)}$, $\Sigma_{22}^{(v)}$ *and* $\Theta_3(\boldsymbol{\Sigma}^{(v)})$ *are non-zero. Furthermore, let* $b_2^{(v)} \neq 0$ *and* $\theta_{\hat{\mu}} > 0$.

*Then,*

$$\frac{\hat{b}_v(\boldsymbol{\Sigma}^{(v)}) - b_v}{\sqrt{\hat{\Sigma}_{vv}(\boldsymbol{\Sigma}^{(v)})}} = \frac{\hat{\mu}_v(\boldsymbol{\Sigma}^{(v)}) - \mu_v(\boldsymbol{\Sigma}^{(v)})}{\sqrt{\lambda_v}\hat{\sigma}_v^{(T)}(\boldsymbol{\Sigma}^{(v)})}. \tag{435}$$

*Proof.* Because $b_2^{(v)} \neq 0$ and $\Sigma_{22}^{(v)} \neq 0$, it follows $\mu_2^{(v)} \neq 0$ (Equ. 89).

For $\left|\hat{\mu}_2^{(v)}\right| < \theta_{\hat{\mu}}$, it follows $\hat{b}_v = b_v$ and $\hat{\mu}_v = \mu_v$ (Equ. 98 and 108). Thus,

$$\frac{\hat{b}_v - b_v}{\sqrt{\hat{\Sigma}_{vv}}} = \frac{\hat{\mu}_v - \mu_v}{\sqrt{\lambda_v}\hat{\sigma}_v^{(\mathrm{T})}} = 0, \tag{436}$$

because both denominators are non-zero (Equ. 113 and 114).

For $\left|\hat{\mu}_2^{(v)}\right| \geq \theta_{\hat{\mu}}$, it follows $\hat{b}_2^{(v)} \neq 0$, because $\Sigma_{22}^{(v)} \neq 0$ (Equ. 97). We obtain from Equ. 107, 108, 85 and 98

$$\begin{aligned}
\hat{\mu}_v - \mu_v &= \frac{\hat{\mu}_1^{(v)}}{\hat{\mu}_2^{(v)}} - \frac{\mu_1^{(v)}}{\mu_2^{(v)}} && (437)\\
&= \frac{1}{\Theta_3(\boldsymbol{\Sigma}^{(v)})}\left(\frac{\hat{b}_1^{(v)}}{\hat{b}_2^{(v)}} - \Theta_4(\boldsymbol{\Sigma}^{(v)})\right) - \frac{\mu_1^{(v)}}{\mu_2^{(v)}} && (438)\\
&= \frac{1}{\Theta_3(\boldsymbol{\Sigma}^{(v)})}\left[\hat{b}_v - \left(\Theta_4(\boldsymbol{\Sigma}^{(v)}) + \Theta_3(\boldsymbol{\Sigma}^{(v)})\frac{\mu_1^{(v)}}{\mu_2^{(v)}}\right)\right], && (439)
\end{aligned}$$

where all denominators are non-zero. Substituting

$$\begin{aligned}
\Theta_4(\boldsymbol{\Sigma}^{(v)}) + \Theta_3(\boldsymbol{\Sigma}^{(v)})\frac{\mu_1^{(v)}}{\mu_2^{(v)}} &= \Theta_4(\boldsymbol{\Sigma}^{(v)}) + \Theta_3(\boldsymbol{\Sigma}^{(v)})\frac{\Theta_1\left(b_1^{(v)}, b_2^{(v)}, \boldsymbol{\Sigma}^{(v)}\right)}{\Theta_2\left(b_2^{(v)}, \boldsymbol{\Sigma}^{(v)}\right)} && (440)\\
&= \frac{\Sigma_{12}^{(v)}}{\sqrt{\Sigma_{11}^{(v)}\Sigma_{22}^{(v)}}}\sqrt{\frac{\Sigma_{11}^{(v)}}{\Sigma_{22}^{(v)}}} + \frac{b_1^{(v)}}{b_2^{(v)}} - \frac{\Sigma_{12}^{(v)}}{\sqrt{\Sigma_{11}^{(v)}\Sigma_{22}^{(v)}}}\sqrt{\frac{\Sigma_{11}^{(v)}}{\Sigma_{22}^{(v)}}} && (441)\\
&= b_v, && (442)
\end{aligned}$$

where we used Equ. 88, 89, 94, 95 and 82, and dividing both sides by $\sqrt{\lambda_v}\hat{\sigma}_v^{(\mathrm{T})}$ results in

$$\frac{\hat{b}_v - b_v}{\Theta_3(\boldsymbol{\Sigma}^{(v)})\sqrt{\lambda_v}\hat{\sigma}_v^{(\mathrm{T})}} = \frac{\hat{b}_v - b_v}{\hat{\Sigma}_{vv}^{1/2}} = \frac{\hat{\mu}_v - \mu_v}{\sqrt{\lambda_v}\hat{\sigma}_v^{(\mathrm{T})}}, \tag{443}$$

where we used Equ. 114. □

**Lemma 17.** *Let* $\mathbf{\Sigma}'^{(v)} \in \mathbb{R}^{2\times 2}$ *such that* $\Sigma'^{(v)}_{22} \neq 0$. *Furthermore, let* $b_2^{(v)} \neq 0$ *and* $\theta_{\hat{\mu}} > 0$. *Then,*

$$\frac{\hat{\mu}_v(\hat{\mathbf{\Sigma}}^{(v)}) - \mu_v(\hat{\mathbf{\Sigma}}^{(v)})}{\sqrt{\lambda_v}\hat{\sigma}_v^{(T)}(\hat{\mathbf{\Sigma}}^{(v)})} \xrightarrow{d} \frac{\rho_1{}^{(v)}(\mathbf{\Sigma}^{(v)}) - \frac{\Theta_1\left(b_1^{(v)}, b_2^{(v)}, \mathbf{\Sigma}'^{(v)}\right)}{\Theta_2\left(b_2^{(v)}, \mathbf{\Sigma}'^{(v)}\right)}\rho_2{}^{(v)}(\mathbf{\Sigma}^{(v)})}{\sqrt{\xi_0\left(1 + \left(\frac{\Theta_1\left(b_1^{(v)}, b_2^{(v)}, \mathbf{\Sigma}'^{(v)}\right)}{\Theta_2\left(b_2^{(v)}, \mathbf{\Sigma}'^{(v)}\right)}\right)^2\right)}} \sim \mathcal{N}(0, \xi_0^{-1}) \tag{444}$$

*as* $N \to \infty$.

*Proof.* From Equ. 86 to 89 we obtain

$$\hat{\mu}_1^{(v)}(\hat{\mathbf{\Sigma}}^{(v)}) = \sqrt{N}\Theta_1\left(b_1^{(v)}, b_2^{(v)}, \hat{\mathbf{\Sigma}}'^{(v)}\right) + \rho_1^{(v)}(\hat{\mathbf{\Sigma}}^{(v)}) \tag{445}$$

$$\hat{\mu}_2^{(v)}(\hat{\mathbf{\Sigma}}^{(v)}) = \sqrt{N}\Theta_2\left(b_2^{(v)}, \hat{\mathbf{\Sigma}}'^{(v)}\right) + \rho_2^{(v)}(\hat{\mathbf{\Sigma}}^{(v)}). \tag{446}$$

From Equ. 108 it follows, that the two sequences of random variables $\hat{\mu}_v(\hat{\mathbf{\Sigma}}^{(v)})$ and $\frac{\hat{\mu}_1^{(v)}(\hat{\mathbf{\Sigma}}^{(v)})}{\hat{\mu}_2^{(v)}(\hat{\mathbf{\Sigma}}^{(v)})}$ have identical terms for the same index $N$ if $\left|\hat{\mu}_2^{(v)}(\hat{\mathbf{\Sigma}}^{(v)})\right| \geq \theta_{\hat{\mu}}$. Furthermore, for all $N$ and all $\varepsilon > 0$

$$P\left(\left|\frac{\hat{\mu}_v - \mu_v}{\sqrt{\lambda_v}\hat{\sigma}_v^{(\mathrm{T})}} - \frac{\frac{\hat{\mu}_1^{(v)}}{\hat{\mu}_2^{(v)}} - \mu_v}{\hat{\sigma}^{(\mathrm{abb})}}\right| > \varepsilon\right) \leq P\left(\left|\hat{\mu}_2^{(v)}\right| < \theta_{\hat{\mu}}\right) \tag{447}$$

$$= P\left(\left|\sqrt{N}\Theta_2 + \rho_2^{(v)}\right| < \theta_{\hat{\mu}}\right) \tag{448}$$

$$= P\left(\left|\Theta_2 + \frac{1}{\sqrt{N}}\rho_2^{(v)}\right| < \frac{\theta_{\hat{\mu}}}{\sqrt{N}}\right), \tag{449}$$

where

$$\hat{\sigma}^{(\mathrm{abb})} = \sqrt{\xi_0\left(1 + \sum_{n=1}^{5}\xi_n\left(\frac{\hat{\mu}_2^{(v)}}{6}\right)^{-2n}\right)}\sqrt{\frac{1}{\hat{\mu}_2^{(v)\,2}} + \frac{\hat{\mu}_1^{(v)\,2}}{\hat{\mu}_2^{(v)\,4}}}. \tag{450}$$

Because $b_2^{(v)} \neq 0$ and $\Sigma'^{(v)}_{22} \neq 0$, it follows $\Theta_2\left(b_2^{(v)}, \mathbf{\Sigma}'^{(v)}\right) \neq 0$ (Equ. 89). We define the constant $\theta' = \frac{\left|\Theta_2\left(b_2^{(v)}, \mathbf{\Sigma}'^{(v)}\right)\right|}{2} > 0$ and obtain

$$P\left(\left|\Theta_2 + \frac{1}{\sqrt{N}}\rho_2^{(v)}\right| < \frac{\theta_{\hat{\mu}}}{\sqrt{N}}\right) \leq P\left(\left|\Theta_2 + \frac{1}{\sqrt{N}}\rho_2^{(v)}\right| < \theta'\right) \tag{451}$$

for large enough $N > \left(\frac{\theta_{\hat{\mu}}}{\theta'}\right)^2$.

Asymptotically, $\hat{\mathbf{\Sigma}}'^{(v)} \xrightarrow{d} \mathbf{\Sigma}'^{(v)}$ as $N \to \infty$ and according to the continuous mapping theorem[69]

$$\left(\Theta_2\left(b_2^{(v)}, \hat{\mathbf{\Sigma}}'^{(v)}\right), \rho_2^{(v)}(\hat{\mathbf{\Sigma}}^{(v)}), \frac{1}{\sqrt{N}}\right) \xrightarrow{d} \left(\Theta_2\left(b_2^{(v)}, \mathbf{\Sigma}'^{(v)}\right), \rho_2^{(v)}(\mathbf{\Sigma}^{(v)}), 0\right) \tag{452}$$

$$\left|\Theta_2\left(b_2^{(v)}, \hat{\mathbf{\Sigma}}'^{(v)}\right) + \frac{1}{\sqrt{N}}\rho_2^{(v)}(\hat{\mathbf{\Sigma}}^{(v)})\right| \xrightarrow{d} \left|\Theta_2\left(b_2^{(v)}, \mathbf{\Sigma}'^{(v)}\right)\right| \tag{453}$$

[69](Van der Vaart, 2000, theorem 2.3)

as $N \to \infty$, where $\boldsymbol{\rho}^{(v)}(\boldsymbol{\Sigma}^{(v)}) \sim \mathcal{N}(\boldsymbol{0}_2, \mathbb{1}_2)$ (see Equ. 90 and 91). By the portmanteau lemma,[70] convergence in distribution in Equ. 453 implies

$$\lim_{N\to\infty} \mathrm{P}\left(\left|\Theta_2\left(b_2^{(v)}, \hat{\boldsymbol{\Sigma}}'^{(v)}\right) + \frac{1}{\sqrt{N}}\rho_2^{(v)}(\hat{\boldsymbol{\Sigma}}^{(v)})\right| \le \theta'\right) = \mathrm{P}\left(\left|\Theta_2\left(b_2^{(v)}, \boldsymbol{\Sigma}'^{(v)}\right)\right| \le \theta'\right) \tag{454}$$

and it follows

$$\lim_{N\to\infty} \mathrm{P}\left(\left|\frac{\hat{\mu}_v - \mu_v}{\sqrt{\lambda_v}\hat{\sigma}_v^{(\mathrm{T})}} - \frac{\frac{\hat{\mu}_1^{(v)}}{\hat{\mu}_2^{(v)}} - \mu_v}{\hat{\sigma}^{(\mathrm{abb})}}\right| > \varepsilon\right) \le \lim_{N\to\infty} \mathrm{P}\left(\left|\Theta_2 + \frac{1}{\sqrt{N}}\rho_2^{(v)}\right| \le \theta'\right) \tag{455}$$

$$= \mathrm{P}\left(\left|\Theta_2\left(b_2^{(v)}, \boldsymbol{\Sigma}'^{(v)}\right)\right| \le \theta'\right) \tag{456}$$

$$= \mathrm{P}\left(2\theta' \le \theta'\right) = 0 \tag{457}$$

for all $\varepsilon > 0$, i.e., the absolute difference of the two sequences converges in probability to zero. This implies that the two sequences converge in distribution[71] and

$$\frac{\hat{\mu}_v - \mu_v}{\sqrt{\lambda_v}\hat{\sigma}_v^{(\mathrm{T})}} \xrightarrow{d} \mathcal{N}(0, \xi_0^{-1}) \tag{458}$$

as $N \to \infty$ if

$$\frac{\frac{\hat{\mu}_1^{(v)}}{\hat{\mu}_2^{(v)}} - \mu_v}{\hat{\sigma}^{(\mathrm{abb})}} \xrightarrow{d} \mathcal{N}(0, \xi_0^{-1}) \tag{459}$$

as $N \to \infty$. We conclude by proofing Equ. 459.

From Equ. 103, 107, 108 and 113 we obtain for $\hat{\mu}_2^{(v)}(\hat{\boldsymbol{\Sigma}}^{(v)}) \neq 0$ [72]

$$\frac{\frac{\hat{\mu}_1^{(v)}}{\hat{\mu}_2^{(v)}} - \mu_v}{\hat{\sigma}^{(\mathrm{abb})}} = \frac{\frac{\hat{\mu}_1^{(v)}}{\hat{\mu}_2^{(v)}} - \frac{\mu_1^{(v)}}{\mu_2^{(v)}}}{\sqrt{\xi_0\left(1 + \sum_{n=1}^5 \xi_n \left(\frac{\hat{\mu}_2^{(v)}}{6}\right)^{-2n}\right)}\sqrt{\frac{1}{\hat{\mu}_2^{(v)\,2}} + \frac{\hat{\mu}_1^{(v)\,2}}{\hat{\mu}_2^{(v)\,4}}}} \tag{460}$$

$$= \frac{\frac{\hat{\mu}_1^{(v)}|\hat{\mu}_2^{(v)}|}{\hat{\mu}_2^{(v)}} - \frac{\mu_1^{(v)}|\hat{\mu}_2^{(v)}|}{\mu_2^{(v)}}}{\sqrt{\xi_0\left(1 + \sum_{n=1}^5 \xi_n \left(\frac{\hat{\mu}_2^{(v)}}{6}\right)^{-2n}\right)}\sqrt{1 + \frac{\hat{\mu}_1^{(v)\,2}}{\hat{\mu}_2^{(v)\,2}}}} \tag{461}$$

$$= \frac{|\hat{\mu}_2^{(v)}|}{\hat{\mu}_2^{(v)}} \frac{\left(\sqrt{N}\Theta_1 + \rho_1^{(v)}\right) - \left(\frac{\sqrt{N}\Theta_1}{\sqrt{N}\Theta_2}\sqrt{N}\Theta_2 + \frac{\sqrt{N}\Theta_1}{\sqrt{N}\Theta_2}\rho_2^{(v)}\right)}{\sqrt{\xi_0\left(1 + \sum_{n=1}^5 \xi_n \left(\frac{\hat{\mu}_2^{(v)}}{6}\right)^{-2n}\right)}\sqrt{1 + \frac{\hat{\mu}_1^{(v)\,2}}{\hat{\mu}_2^{(v)\,2}}}}. \tag{462}$$

According to the continuous mapping theorem, we obtain

$$\left(\Theta_1\left(b_1^{(v)}, b_2^{(v)}, \hat{\boldsymbol{\Sigma}}'^{(v)}\right), \Theta_2\left(b_2^{(v)}, \hat{\boldsymbol{\Sigma}}'^{(v)}\right), \boldsymbol{\rho}^{(v)}(\hat{\boldsymbol{\Sigma}}^{(v)}), \frac{1}{\sqrt{N}}\right) \tag{463}$$

$$\xrightarrow{d} \left(\Theta_1\left(b_1^{(v)}, b_2^{(v)}, \boldsymbol{\Sigma}'^{(v)}\right), \Theta_2\left(b_2^{(v)}, \boldsymbol{\Sigma}'^{(v)}\right), \boldsymbol{\rho}^{(v)}(\boldsymbol{\Sigma}^{(v)}), 0\right) \tag{464}$$

[70](Van der Vaart, 2000, theorem 2.2 (i))

[71](Van der Vaart, 2000, theorem 2.7 (iv))

[72]$\hat{\mu}_2^{(v)}(\hat{\boldsymbol{\Sigma}}^{(v)}) = 0$ has mass zero.

and

$$\frac{\frac{\hat{\mu}_1^{(v)}}{\hat{\mu}_2^{(v)}} - \mu_v}{\hat{\sigma}^{(\text{abb})}} \xrightarrow{d} \pm \frac{\rho_1{}^{(v)}(\mathbf{\Sigma}^{(v)}) - \frac{\Theta_1\left(b_1^{(v)}, b_2^{(v)}, \mathbf{\Sigma}'^{(v)}\right)}{\Theta_2\left(b_2^{(v)}, \mathbf{\Sigma}'^{(v)}\right)} \rho_2{}^{(v)}(\mathbf{\Sigma}^{(v)})}{\sqrt{\xi_0 \left(1 + \left(\frac{\Theta_1\left(b_1^{(v)}, b_2^{(v)}, \mathbf{\Sigma}'^{(v)}\right)}{\Theta_2\left(b_2^{(v)}, \mathbf{\Sigma}'^{(v)}\right)}\right)^2\right)}} \tag{465}$$

$$\xrightarrow{d} \mathcal{N}(0, \xi_0^{-1}) \tag{466}$$

as $N \to \infty$, where the sign in Equ. 465 is the same as the sign of $\Theta_2\left(b_2^{(v)}, \mathbf{\Sigma}'^{(v)}\right)$. □

**Proposition 18.** *Let $\hat{b}_i^{(v)}$ and $\hat{\Sigma}_{ii}^{\prime(v)}$, $i \in \{1,2\}$ denote the sample mean and the sample variance of $N > 1$ i.i.d. normal random samples.*

*Then,*

$$\frac{VAR\left[\hat{\Sigma}_{ii}^{(v)\,\frac{1}{2}}\right]}{VAR[\hat{b}_i^{(v)}]} = 1 - c_4\left(N\right)^2, \; where \qquad c_4\left(N\right) = \sqrt{\frac{2}{N-1}}\frac{\Gamma\left(\frac{N}{2}\right)}{\Gamma\left(\frac{N-1}{2}\right)}, \tag{467}$$

*$\hat{\Sigma}_{ii}^{(v)} = \hat{\Sigma}_{ii}^{\prime(v)}/N$ and $\Gamma$ denotes the Gamma function, with values less than $10^{-2}$ and $10^{-3}$ for more than 50 and 500 samples, respectively.*

*Proof.* The sample variance $\hat{\Sigma}_{ii}^{\prime(v)}$ is an unbiased estimator of the population variance denoted by $\Sigma_{ii}^{\prime(v)}$, i.e. $E[\hat{\Sigma}_{ii}^{\prime(v)}] = \Sigma_{ii}^{\prime(v)}$, and the variance of the sample mean is given by $\mathrm{VAR}[\hat{b}_i^{(v)}] = \frac{1}{N}\Sigma_{ii}^{\prime(v)}$. Then,

$$\frac{\mathrm{VAR}\left[\hat{\Sigma}_{ii}^{(v)\,\frac{1}{2}}\right]}{\mathrm{VAR}[\hat{b}_i^{(v)}]} = \frac{\mathrm{VAR}\left[\hat{\Sigma}_{ii}^{\prime(v)\,1/2}\right]}{N\mathrm{VAR}[\hat{b}_i^{(v)}]} \tag{468}$$

$$= \frac{E\left[\hat{\Sigma}_{ii}^{\prime(v)}\right] - E\left[\hat{\Sigma}_{ii}^{\prime(v)\,1/2}\right]^2}{N\mathrm{VAR}[\hat{b}_i^{(v)}]} \tag{469}$$

$$= \frac{\Sigma_{ii}^{\prime(v)} - E\left[\hat{\Sigma}_{ii}^{\prime(v)\,1/2}\right]^2}{\Sigma_{ii}^{\prime(v)}} \tag{470}$$

$$= 1 - \frac{E\left[\hat{\Sigma}_{ii}^{\prime(v)\,1/2}\right]^2}{\Sigma_{ii}^{\prime(v)}} \tag{471}$$

and

$$\lim_{N\to\infty}\frac{\mathrm{VAR}\left[\hat{\Sigma}_{ii}^{(v)\,\frac{1}{2}}\right]}{\mathrm{VAR}[\hat{b}_i^{(v)}]} = 1 - \lim_{N\to\infty}\frac{E\left[\hat{\Sigma}_{ii}^{\prime(v)\,1/2}\right]^2}{\Sigma_{ii}^{\prime(v)}} = 1 - 1 = 0, \tag{472}$$

because $\hat{\Sigma}_{ii}^{\prime(v)\,1/2}$ is an asymptotically consistent estimator of $\Sigma_{ii}^{\prime(v)\,1/2}$. According to (Kenney and Keeping, 1951),[73] for i.i.d. normal random samples

$$E\left[\hat{\Sigma}_{ii}^{\prime(v)\,1/2}\right] = c_4\left(N\right)\Sigma_{ii}^{\prime(v)\,1/2}, \tag{473}$$

and

$$\frac{\mathrm{VAR}\left[\hat{\Sigma}_{ii}^{(v)\,\frac{1}{2}}\right]}{\mathrm{VAR}[\hat{b}_i^{(v)}]} = 1 - c_4\left(N\right)^2 \tag{474}$$

with values less than $10^{-2}$ and $10^{-3}$ for more than 50 and 500 samples, respectively.

□

[73]Weisstein, Eric W. "Standard Deviation Distribution." From MathWorld–A Wolfram Web Resource.

**Lemma 19.** *Let* $b_2^{(v)} \neq 0$, $\hat{\Sigma}_{22}'^{(v)} > 0$ *and* $\theta_{\hat{\mu}} > 0$.
*Then,*

$$\lambda_v\left(\hat{\mu}_2^{(v)}(\hat{\mathbf{\Sigma}}^{(v)})\right) \overset{p}{\to} \xi_0 \tag{475}$$

*as* $N \to \infty$.

*Proof.* According to Equ. 97,[74]

$$\hat{\mu}_2^{(v)}(\hat{\mathbf{\Sigma}}^{(v)}) = \Theta_2\left(\hat{b}_2^{(v)}, \hat{\mathbf{\Sigma}}^{(v)}\right) = \frac{1}{\sqrt{N}}\Theta_2\left(\hat{b}_2^{(v)}, \hat{\mathbf{\Sigma}}'^{(v)}\right). \tag{476}$$

Asymptotically, $\hat{b}_2^{(v)} \overset{p}{\to} b_2^{(v)}$, $\hat{\mathbf{\Sigma}}'^{(v)} \overset{p}{\to} \mathbf{\Sigma}'^{(v)}$, $\frac{1}{\sqrt{N}} \overset{p}{\to} 0$ and $\left(\frac{1}{\sqrt{N}}, \hat{b}_2^{(v)}, \hat{\mathbf{\Sigma}}'^{(v)}\right) \overset{p}{\to} \left(0, b_2^{(v)}, \mathbf{\Sigma}'^{(v)}\right)$ as $N \to \infty$.[75] The continuous mapping theorem implies

$$\frac{1}{\sqrt{N}}\left|\hat{\mu}_2^{(v)}(\hat{\mathbf{\Sigma}}^{(v)})\right| \overset{p}{\to} \left|\Theta_2\left(b_2^{(v)}, \mathbf{\Sigma}'^{(v)}\right)\right| > 0 \tag{477}$$

$$\frac{1}{\sqrt{N}}\theta_{\hat{\mu}} \overset{p}{\to} 0 \tag{478}$$

as $N \to \infty$ and, thus,

$$\lim_{N\to\infty} \mathrm{P}\left(\left|\hat{\mu}_2^{(v)}(\hat{\mathbf{\Sigma}}^{(v)})\right| < \theta_{\hat{\mu}}\right) = 0. \tag{479}$$

According to Equ. 113,

$$\lambda_v\left(\hat{\mu}_2^{(v)}(\hat{\mathbf{\Sigma}}^{(v)})\right) = \begin{cases} 1, & \left|\hat{\mu}_2^{(v)}(\hat{\mathbf{\Sigma}}^{(v)})\right| < \theta_{\hat{\mu}} \\ \xi_0\left(1 + \sum_{n=1}^{5}\xi_n\left(\frac{\hat{\mu}_2^{(v)}(\hat{\mathbf{\Sigma}}^{(v)})}{6}\right)^{-2n}\right), & \text{otherwise.} \end{cases} \tag{480}$$

For all $N$ and all $\varepsilon > 0$,

$$\mathrm{P}\left(\left|\lambda_v - \xi_0\left(1 + \sum_{n=1}^{5}\xi_n\left(\frac{\hat{\mu}_2^{(v)}(\hat{\mathbf{\Sigma}}^{(v)})}{6}\right)^{-2n}\right)\right| > \varepsilon\right) \leq \mathrm{P}\left(\left|\hat{\mu}_2^{(v)}(\hat{\mathbf{\Sigma}}^{(v)})\right| < \theta_{\hat{\mu}}\right) \tag{481}$$

and

$$\lambda_v\left(\hat{\mu}_2^{(v)}(\hat{\mathbf{\Sigma}}^{(v)})\right) \overset{p}{\to} \xi_0\left(1 + \sum_{n=1}^{5}\xi_n\left(\frac{\hat{\mu}_2^{(v)}(\hat{\mathbf{\Sigma}}^{(v)})}{6}\right)^{-2n}\right) \tag{482}$$

as $N \to \infty$. It follows from the continuous mapping theorem and Equ. 477,

$$\xi_0\left(1 + \sum_{n=1}^{5}\xi_n\left(\frac{\hat{\mu}_2^{(v)}(\hat{\mathbf{\Sigma}}^{(v)})}{6}\right)^{-2n}\right) = \xi_0\left(1 + \sum_{n=1}^{5}\frac{1}{(N)^n}\xi_n\left(\frac{\left(\frac{1}{\sqrt{N}}\left|\hat{\mu}_2^{(v)}(\hat{\mathbf{\Sigma}}^{(v)})\right|\right)^2}{6^2}\right)^{-n}\right) \tag{483}$$

$$\overset{p}{\to} \xi_0 \tag{484}$$

and $\lambda_v\left(\hat{\mu}_2^{(v)}(\hat{\mathbf{\Sigma}}^{(v)})\right) \overset{p}{\to} \xi_0$ as $N \to \infty$. □

---

[74] $\lambda_v$, $\hat{\mu}_2^{(v)}$, $\hat{\mathbf{\Sigma}}'^{(v)}$ and $\xi_0$ are defined in Equ. 113, Equ. 97, Equ. 70 and Table 2, respectively.
[75] (Van der Vaart, 2000, theorem 2.7 (vi))

**Lemma 20.** *Let*

$$\Sigma'^{(v,\omega)}_{ij} = E\left[\rho'^{(v)}_i \rho'^{(\omega)}_j\right] \quad (485)$$

$$\Sigma^{(v,\omega)}_{ij} = \frac{1}{N}\Sigma'^{(v,\omega)}_{ij} \quad (486)$$

*for* $i, j \in \{1, 2\}$ *and* $v, \omega \in \{1, ..., d^{(X_0)}\}$. *Assume the covariance matrix*

$$\mathbf{\Sigma}'^{(all)} = \begin{pmatrix} \mathbf{\Sigma}'^{(1,1)} & \cdots & \mathbf{\Sigma}'^{(1,d^{(X_0)})} \\ \vdots & \ddots & \vdots \\ \mathbf{\Sigma}'^{(d^{(X_0)},1)} & \cdots & \mathbf{\Sigma}'^{(d^{(X_0)},d^{(X_0)})} \end{pmatrix} \quad (487)$$

*is invertible. Furthermore, let* $b_2^{(v)} \neq 0$ *for all* $v \in \{1, ..., d^{(X_0)}\}$ *and* $\theta_{\hat{\mu}} > 0$.

*Then,*

$$\sqrt{N}(\hat{\mathbf{b}} - \mathbf{b}) \xrightarrow{d} \mathcal{N}(\mathbf{0}, \mathbf{\Sigma}') \quad (488)$$

$$N\hat{\mathbf{\Sigma}}^{(\infty)}(\hat{\mathbf{\Sigma}}^{(v,\omega)}) \xrightarrow{p} \mathbf{\Sigma}' \quad (489)$$

$$N\hat{\mathbf{\Sigma}}(\hat{\mathbf{\Sigma}}^{(v,\omega)}) \xrightarrow{p} \xi_0\mathbf{\Sigma}' \quad (490)$$

$$\Sigma'_{v\omega} = \frac{1}{b_2^{(v)}b_2^{(\omega)}}\left(\Sigma'^{(v,w)}_{11} - \frac{b_1^{(\omega)}}{b_2^{(\omega)}}\Sigma'^{(v,w)}_{12} - \frac{b_1^{(v)}}{b_2^{(v)}}\Sigma'^{(v,w)}_{21} + \frac{b_1^{(v)}b_1^{(\omega)}}{b_2^{(v)}b_2^{(\omega)}}\Sigma'^{(v,w)}_{22}\right) \quad (491)$$

*for* $v, \omega \in \{1, ..., d^{(X_0)}\}$ *as* $N \to \infty$ *and* $\mathbf{\Sigma}'$ *has full rank.*

*Proof.* We abbreviate $\mathbf{\Sigma}'^{(v,v)}$ as $\mathbf{\Sigma}'^{(v)}$. The covariance matrix $\mathbf{\Sigma}'^{(\text{all})}$ is positive semidefinite and, thus, invertible if and only if it is positive definite.[76] Then, $\Sigma'^{(v)}_{11}$, $\Sigma'^{(v)}_{22}$, and $\Theta_3(\mathbf{\Sigma}'^{(v)})$ have to be larger than zero for all $v \in \{1, ..., d^{(X_0)}\}$. We first prove Equ. 488, subsequently, we prove Equ. 489 and 490 and, finally, we show that $\mathbf{\Sigma}'$ has full rank.

For finite $N$,[77]

$$\hat{b}_v(\mathbf{\Sigma}^{(v)}) = \begin{cases} b_v, & \left|\hat{\mu}_2^{(v)}(\mathbf{\Sigma}^{(v)})\right| < \theta_{\hat{\mu}} \\ \frac{\hat{b}_1^{(v)}}{\hat{b}_2^{(v)}}, & \text{otherwise} \end{cases} \quad (492)$$

for $v \in \{1, ..., d^{(X_0)}\}$ and cutoff $\theta_{\hat{\mu}} > 0$. According to the multidimensional central limit theorem (Van der Vaart, 2000), the distributions of $\hat{\mathbf{b}}^{(v)}$ (Equ. 98) converge for invertible covariance matrices $\mathbf{\Sigma}'^{(\text{all})}$ to the joint normal distribution

$$\begin{pmatrix} \boldsymbol{\rho}'^{(1)} \\ \vdots \\ \boldsymbol{\rho}'^{(d^{(X_0)})} \end{pmatrix} = \sqrt{N}\left[\begin{pmatrix} \hat{\mathbf{b}}^{(1)} \\ \vdots \\ \hat{\mathbf{b}}^{(d^{(X_0)})} \end{pmatrix} - \begin{pmatrix} \mathbf{b}^{(1)} \\ \vdots \\ \mathbf{b}^{(d^{(X_0)})} \end{pmatrix}\right] \xrightarrow{d} \mathcal{N}\left(\mathbf{0}, \mathbf{\Sigma}'^{(\text{all})}\right) \quad (493)$$

as $N \to \infty$ (compare Equ. 83).

Asymptotically, $\hat{\mu}_2^{(v)}(\mathbf{\Sigma}^{(v)})$ is normal, has unit variance and a diverging mean (Equ. 87), so that for any finite $\theta_{\hat{\mu}}$

$$\lim_{N\to\infty} \mathrm{P}\left[\left|\hat{\mu}_2^{(v)}(\mathbf{\Sigma}^{(v)})\right| < \theta_{\hat{\mu}}\right] = 0. \quad (494)$$

[76](Horn and Johnson, 2012, p. 431, Corollary 7.1.7)

[77]See Equ. 98.

We show below that this implies

$$\sqrt{N}(\hat{b}_v(\mathbf{\Sigma}^{(v)}) - b_v) \xrightarrow{p} \sqrt{N}\left(\frac{\hat{b}_1^{(v)}}{\hat{b}_2^{(v)}} - b_v\right) \tag{495}$$

as $N \to \infty$ for all $v \in \{1, ..., d^{(X_0)}\}$. Then, also the corresponding joint vectors converge in probability[78] and, thus, in distribution.[79] As[80]

$$\sqrt{N}\left(\frac{\hat{b}_1^{(v)}}{\hat{b}_2^{(v)}} - b_v\right) = \frac{1}{\hat{b}_2^{(v)}}\sqrt{N}\left(\hat{b}_1^{(v)} - \frac{b_1^{(v)}}{b_2^{(v)}}\hat{b}_2^{(v)}\right) \tag{496}$$

$$= \frac{1}{\hat{b}_2^{(v)}}\left(\sqrt{N}b_1^{(v)} + \rho_1'^{(v)} - \frac{b_1^{(v)}}{b_2^{(v)}}\sqrt{N}b_2^{(v)} - \frac{b_1^{(v)}}{b_2^{(v)}}\rho_2'^{(v)}\right) \tag{497}$$

$$= \frac{1}{\hat{b}_2^{(v)}}\left(\rho_1'^{(v)} - \frac{b_1^{(v)}}{b_2^{(v)}}\rho_2'^{(v)}\right) \tag{498}$$

and $\boldsymbol{\rho}'^{(v)}$ converges in distribution to a joint normal with zero mean as $N \to \infty$ also $\sqrt{N}\left(\frac{\hat{b}_1^{(v)}}{\hat{b}_2^{(v)}} - b_v\right)$ converges in probability to the joint normal $\frac{1}{b_2^{(v)}}\left(\rho_1'^{(v)} - \frac{b_1^{(v)}}{b_2^{(v)}}\rho_2'^{(v)}\right)$ with zero mean and covariance matrix

$$N\text{COV}\left[\frac{\hat{b}_1^{(v)}}{\hat{b}_2^{(v)}}, \frac{\hat{b}_1^{(\omega)}}{\hat{b}_2^{(\omega)}}\right] \xrightarrow{d} \frac{1}{b_2^{(v)}}\frac{1}{b_2^{(\omega)}}\text{COV}\left[\rho_1'^{(v)} - \frac{b_1^{(v)}}{b_2^{(v)}}\rho_2'^{(v)}, \rho_1'^{(\omega)} - \frac{b_1^{(\omega)}}{b_2^{(\omega)}}\rho_2'^{(\omega)}\right] \tag{499}$$

$$\xrightarrow{d} \Sigma'_{v\omega} \tag{500}$$

as $N \to \infty$ for

$$\Sigma'_{v\omega} = \frac{1}{b_2^{(v)}b_2^{(\omega)}}\left(\Sigma_{11}'^{(v,w)} - \frac{b_1^{(\omega)}}{b_2^{(\omega)}}\Sigma_{12}'^{(v,w)} - \frac{b_1^{(v)}}{b_2^{(v)}}\Sigma_{21}'^{(v,w)} + \frac{b_1^{(v)}b_1^{(\omega)}}{b_2^{(v)}b_2^{(\omega)}}\Sigma_{22}'^{(v,w)}\right), \tag{501}$$

which concludes the proof of Equ. 488.

We now show that

$$\lim_{N\to\infty} \text{P}\left[\left|\hat{\mu}_2^{(v)}(\mathbf{\Sigma}^{(v)})\right| < \theta_{\hat{\mu}}\right] = 0 \tag{502}$$

implies

$$\sqrt{N}(\hat{b}_v(\mathbf{\Sigma}^{(v)}) - b_v) \xrightarrow{p} \sqrt{N}\left(\frac{\hat{b}_1^{(v)}}{\hat{b}_2^{(v)}} - b_v\right) \tag{503}$$

for $v \in \{1, ..., d^{(X_0)}\}$. That is, we show for all $\varepsilon > 0$

$$\lim_{N\to\infty} \text{P}\left(\left|\sqrt{N}\left(\hat{b}_v(\mathbf{\Sigma}^{(v)}) - \frac{\hat{b}_1^{(v)}}{\hat{b}_2^{(v)}}\right)\right| > \varepsilon\right) = 0. \tag{504}$$

[78](Van der Vaart, 2000, theorem 2.7 (vi))

[79]If one of the joint vectors converges in distribution as is shown below.

[80]For the definition of $b_v$ see Equ. 82 and $\sqrt{N}\hat{b}_i^{(v)} = \sqrt{N}b_i^{(v)} + \rho_i'^{(v)}, i \in \{1, 2\}$ precisely (see Equ. 83).

For finite $N$, $\hat{b}_v(\mathbf{\Sigma}^{(v)}) = \frac{\hat{b}_1^{(v)}}{\hat{b}_2^{(v)}}$ if $\left|\hat{\mu}_2^{(v)}(\mathbf{\Sigma}^{(v)})\right| \geq \theta_{\hat{\mu}}$, i.e., $\hat{b}_v(\mathbf{\Sigma}^{(v)})$ is not clamped. For all $\varepsilon > 0$,[81]

$$\mathrm{P}\left(\left|\sqrt{N}\left(\hat{b}_v(\mathbf{\Sigma}^{(v)}) - \frac{\hat{b}_1^{(v)}}{\hat{b}_2^{(v)}}\right)\right| > \varepsilon\right) \leq \mathrm{P}\left[\left|\hat{\mu}_2^{(v)}(\mathbf{\Sigma}^{(v)})\right| < \theta_{\hat{\mu}}\right] \tag{505}$$

$$= \mathrm{P}\left[\left|\Theta_2\left(b_2^{(v)}, \mathbf{\Sigma}'^{(v)}\right) + \frac{\rho_2^{(v)}(\mathbf{\Sigma}^{(v)})}{\sqrt{N}}\right| < \frac{\theta_{\hat{\mu}}}{\sqrt{N}}\right]. \tag{506}$$

The term $\Theta_2\left(b_2^{(v)}, \mathbf{\Sigma}'^{(v)}\right) \neq 0$, because $b_2^{(v)} \neq 0$ and $\Sigma_{22}'^{(v)} \neq 0$ (Equ. 93). We define the constant $\theta' = \frac{\left|\Theta_2\left(b_2^{(v)}, \mathbf{\Sigma}'^{(v)}\right)\right|}{2} > 0$. For large enough $N > \left(\frac{\theta_{\hat{\mu}}}{\theta'}\right)^2$,

$$\mathrm{P}\left(\left|\Theta_2\left(b_2^{(v)}, \mathbf{\Sigma}'^{(v)}\right) + \frac{\rho_2^{(v)}(\mathbf{\Sigma}^{(v)})}{\sqrt{N}}\right| < \frac{\theta_{\hat{\mu}}}{\sqrt{N}}\right) \leq \mathrm{P}\left(\left|\Theta_2\left(b_2^{(v)}, \mathbf{\Sigma}'^{(v)}\right) + \frac{\rho_2^{(v)}(\mathbf{\Sigma}^{(v)})}{\sqrt{N}}\right| < \theta'\right). \tag{507}$$

According to the continuous mapping theorem,[82]

$$\left|\Theta_2\left(b_2^{(v)}, \mathbf{\Sigma}'^{(v)}\right) + \frac{1}{\sqrt{N}}\rho_2^{(v)}(\mathbf{\Sigma}^{(v)})\right| \xrightarrow{d} \left|\Theta_2\left(b_2^{(v)}, \mathbf{\Sigma}'^{(v)}\right)\right| \tag{508}$$

as $N \to \infty$. By the portmanteau lemma,[83] convergence in distribution implies

$$\lim_{N\to\infty} \mathrm{P}\left(\left|\Theta_2\left(b_2^{(v)}, \mathbf{\Sigma}'^{(v)}\right) + \frac{1}{\sqrt{N}}\rho_2^{(v)}(\mathbf{\Sigma}^{(v)})\right| \leq \theta'\right) = \mathrm{P}\left(\left|\Theta_2\left(b_2^{(v)}, \mathbf{\Sigma}'^{(v)}\right)\right| \leq \theta'\right) \tag{509}$$

and taking the limit on both sides of Equ. 505 and 506 implies

$$\lim_{N\to\infty} \mathrm{P}\left(\left|\sqrt{N}\left(\hat{b}_v(\mathbf{\Sigma}^{(v)}) - \frac{\hat{b}_1^{(v)}}{\hat{b}_2^{(v)}}\right)\right| > \varepsilon\right) \leq \tag{510}$$

$$\lim_{N\to\infty} \mathrm{P}\left(\left|\Theta_2\left(b_2^{(v)}, \mathbf{\Sigma}'^{(v)}\right) + \frac{\rho_2^{(v)}(\mathbf{\Sigma}^{(v)})}{\sqrt{N}}\right| < \theta'\right)$$

$$= \mathrm{P}\left(\left|\Theta_2\left(b_2^{(v)}, \mathbf{\Sigma}'^{(v)}\right)\right| \leq \theta'\right) \tag{511}$$

$$= \mathrm{P}\left(2\theta' \leq \theta'\right) = 0 \tag{512}$$

for all $\varepsilon > 0$, i.e., convergence in probability, which concludes the proof of Equ. 503.

We now prove Equ. 489 and 490. The covariance $\mathbf{\Sigma}'$ can be estimated analogous to Equ. 101 by the covariance of the second order Taylor series of $\frac{\hat{b}_1^{(v)}}{\hat{b}_2^{(v)}}$ and $\frac{\hat{b}_1^{(\omega)}}{\hat{b}_2^{(\omega)}}$ under the assumption that the denominators have zero mass around zero (Stuart and Ord, 1994)[84],

[81] See Equ. 87 and Equ. 89.

[82] (Van der Vaart, 2000, theorem 2.3)

[83] (Van der Vaart, 2000, theorem 2.2 (i))

[84] From the first order Taylor series for $f(x, y) = \frac{x}{y}$ around $(x_0, y_0)$ and $E[f(x, y)] \approx f(x_0, y_0)$ follows $\mathrm{COV}[f(x, y), f(u, v)] \approx E\left[\left(f_x(x_0, y_0)(x - x_0) + f_y(x_0, y_0)(y - y_0)\right)\left(f_u(u_0, v_0)(u - u_0) + f_v(u_0, v_0)(v - v_0)\right)\right]$ and $\mathrm{COV}[f(x, y), f(u, v)] \approx \frac{x_0 u_0}{y_0 v_0}\left(\frac{\mathrm{COV}[x,u]}{x_0 u_0} - \frac{\mathrm{COV}[x,v]}{x_0 v_0} - \frac{\mathrm{COV}[y,u]}{y_0 u_0} + \frac{\mathrm{COV}[y,v]}{y_0 v_0}\right)$. (Stuart and Ord, 1994, p. 351)

i.e.,

$$\mathrm{COV}\left[\frac{\hat{b}_1^{(v)}}{\hat{b}_2^{(v)}}, \frac{\hat{b}_1^{(\omega)}}{\hat{b}_2^{(\omega)}}\right] \approx \frac{b_1^{(v)}b_1^{(\omega)}}{b_2^{(v)}b_2^{(\omega)}}\left(\frac{\mathrm{COV}[\hat{b}_1^{(v)}, \hat{b}_1^{(\omega)}]}{b_1^{(v)}b_1^{(\omega)}} - \frac{\mathrm{COV}[\hat{b}_1^{(v)}, \hat{b}_2^{(\omega)}]}{b_1^{(v)}b_2^{(\omega)}} \right. \tag{513}$$

$$\left. - \frac{\mathrm{COV}[\hat{b}_2^{(v)}, \hat{b}_1^{(\omega)}]}{b_2^{(v)}b_1^{(\omega)}} + \frac{\mathrm{COV}[\hat{b}_2^{(v)}, \hat{b}_2^{(\omega)}]}{b_2^{(v)}b_2^{(\omega)}}\right)$$

$$= \frac{b_1^{(v)}b_1^{(\omega)}}{b_2^{(v)}b_2^{(\omega)}}\left(\frac{\Sigma_{11}^{(v,w)}}{b_1^{(v)}b_1^{(\omega)}} - \frac{\Sigma_{12}^{(v,w)}}{b_1^{(v)}b_2^{(\omega)}} \right. \tag{514}$$

$$\left. - \frac{\Sigma_{21}^{(v,w)}}{b_2^{(v)}b_1^{(\omega)}} + \frac{\Sigma_{22}^{(v,w)}}{b_2^{(v)}b_2^{(\omega)}}\right)$$

for $v, \omega \in \{1, ..., d^{(X_0)}\}$, where[85]

$$\mathbf{\Sigma}^{(v,\omega)} = \frac{1}{N}\mathbf{\Sigma}'^{(v,\omega)} = \mathrm{COV}[\hat{b}_1^{(v)}, \hat{b}_1^{(\omega)}]. \tag{515}$$

Because $\mathbf{\Sigma}'^{(v,\omega)}$ and $\mathbf{b}^{(v)}, \mathbf{b}^{(\omega)}$ are unknown, we substitute the corresponding sample estimates and define (see Equ. 144 and 145)

$$\hat{\Sigma}_{v\omega}^{(\infty)}(\hat{\mathbf{\Sigma}}^{(v,\omega)}) = \begin{cases} \hat{\sigma}_0\Theta_3(\hat{\mathbf{\Sigma}}^{(v,v)})\Theta_3(\hat{\mathbf{\Sigma}}^{(\omega,\omega)}), & (v = \omega) \wedge \left(\left|\hat{\mu}_2^{(v)}\right| < \theta_{\hat{\mu}}\right) \\ 0, & (v \neq \omega) \wedge \left(\left(\left|\hat{\mu}_2^{(v)}\right| < \theta_{\hat{\mu}}\right) \vee \left(\left|\hat{\mu}_2^{(\omega)}\right| < \theta_{\hat{\mu}}\right)\right) \\ \frac{1}{\hat{b}_2^{(v)}\hat{b}_2^{(\omega)}}\left(\hat{\Sigma}_{11}^{(v,w)} - \frac{\hat{b}_1^{(\omega)}}{\hat{b}_2^{(\omega)}}\hat{\Sigma}_{12}^{(v,w)} - \right. & \\ \left. \frac{\hat{b}_1^{(v)}}{\hat{b}_2^{(v)}}\hat{\Sigma}_{21}^{(v,w)} + \frac{\hat{b}_1^{(v)}\hat{b}_1^{(\omega)}}{\hat{b}_2^{(v)}\hat{b}_2^{(\omega)}}\hat{\Sigma}_{22}^{(v,w)}\right), & \text{otherwise} \end{cases} \tag{516}$$

for arbitrary $\hat{\sigma}_0 > 0$ (the same $\hat{\sigma}_0$ as in Equ. 103), where the factor $\Theta_3(\hat{\mathbf{\Sigma}}^{(v,v)})\Theta_3(\hat{\mathbf{\Sigma}}^{(\omega,\omega)})$ in Equ. 516 ensures for $v = \omega$ consistency with $\hat{\Sigma}_{vv}^{(\infty)}(\hat{\mathbf{\Sigma}}^{(v)})$ as defined in Equ. 102. More, precisely, $\hat{\Sigma}_{v\omega}^{(\infty)}(\hat{\mathbf{\Sigma}}^{(v,\omega)}) = \hat{\Sigma}_{vv}^{(\infty)}(\hat{\mathbf{\Sigma}}^{(v)})$ for $v = \omega$ and $\hat{\mathbf{\Sigma}}^{(v,v)} = \hat{\mathbf{\Sigma}}^{(v)}$ (see Equ. 102 and 103).[86]

Asymptotically,[87]

$$\lim_{N\to\infty} \Pr\left[\left(\left|\hat{\mu}_2^{(v)}(\hat{\mathbf{\Sigma}}^{(v,v)})\right| < \theta_{\hat{\mu}}\right) \vee \left(\left|\hat{\mu}_2^{(\omega)}(\hat{\mathbf{\Sigma}}^{(\omega,\omega)})\right| < \theta_{\hat{\mu}}\right)\right] = 0 \tag{517}$$

and, thus, $\hat{\Sigma}_{v\omega}^{(\infty)}(\hat{\mathbf{\Sigma}}^{(v,\omega)})$ converges in distribution to the formula in the second case of Equ. 516. Furthermore,

$$\left(\hat{\mathbf{b}}^{(v)}, \hat{\mathbf{\Sigma}}'^{(v,\omega)}\right) \xrightarrow{p} \left(\mathbf{b}^{(v)}, \mathbf{\Sigma}'^{(v,\omega)}\right) \tag{518}$$

as $N \to \infty$ and, according to the continuous mapping theorem,[88]

$$N\hat{\Sigma}_{v\omega}^{(\infty)}(\hat{\mathbf{\Sigma}}^{(v,\omega)}) \xrightarrow{p} \frac{1}{b_2^{(v)}b_2^{(\omega)}}\left(\Sigma_{11}'^{(v,w)} - \frac{b_1^{(\omega)}}{b_2^{(\omega)}}\Sigma_{12}'^{(v,w)} - \frac{b_1^{(v)}}{b_2^{(v)}}\Sigma_{21}'^{(v,w)} + \frac{b_1^{(v)}b_1^{(\omega)}}{b_2^{(v)}b_2^{(\omega)}}\Sigma_{22}'^{(v,w)}\right) \tag{519}$$

$$= \Sigma_{v\omega}' \tag{520}$$

[85] The last step follows from the definition of $\hat{b}_1^{(v)}$ and $\hat{b}_2^{(v)}$ in Equ. 68 and Equ. 98.

[86] This can be shown by substituting Equ. 85 and Equ. 97.

[87] The proof is analogous to the proof from Equ. 502 to Equ. 512.

[88] Convergence in distribution to a constant implies convergence in probability. (Van der Vaart, 2000, theorem 2.7)

as $N \to \infty$.[89] By definition

$$\hat{\Sigma}_{\nu\omega}(\hat{\mathbf{\Sigma}}^{(\nu,\omega)}) = \sqrt{\lambda_\nu \left(\hat{\mu}_2^{(\nu)}(\hat{\mathbf{\Sigma}}^{(\nu,\nu)})\right) \lambda_\omega \left(\hat{\mu}_2^{(\omega)}(\hat{\mathbf{\Sigma}}^{(\omega,\omega)})\right)} \hat{\Sigma}_{\nu\omega}^{(\infty)}(\hat{\mathbf{\Sigma}}^{(\nu,\omega)}) \tag{521}$$

for $\nu, \omega \in \{1, ..., d^{(X_0)}\}$ and we obtain[90]

$$\begin{aligned}
\lambda_\nu \left(\hat{\mu}_2^{(\nu)}(\hat{\mathbf{\Sigma}}^{(\nu,\nu)})\right) &\xrightarrow{p} \xi_0 && (522)\\
N\hat{\Sigma}_{\nu\omega}(\hat{\mathbf{\Sigma}}^{(\nu,\omega)}) &\xrightarrow{p} \xi_0 N \hat{\Sigma}_{\nu\omega}^{(\infty)}(\hat{\mathbf{\Sigma}}^{(\nu,\omega)}) && (523)\\
&\xrightarrow{p} \xi_0 \Sigma'_{\nu\omega} && (524)
\end{aligned}$$

as $N \to \infty$, which concludes the proof of Equ. 489 and 490.

Finally, we show $\mathbf{\Sigma}'$ has full rank. We define

$$\begin{aligned}
\mathbf{U} &= \begin{pmatrix} u'_1 & u''_1 & 0 & 0 & \cdots & 0 & 0 \\ 0 & 0 & u'_2 & u''_2 & & 0 & 0 \\ \vdots & & & & \ddots & & \vdots \\ 0 & 0 & 0 & 0 & \cdots & u'_{d^{(X_0)}} & u''_{d^{(X_0)}} \end{pmatrix} && (525)\\
u'_\nu &= \frac{1}{b_2^{(\nu)}} && (526)\\
u''_\nu &= -\frac{b_1^{(\nu)}}{b_2^{(\nu)2}}, && (527)
\end{aligned}$$

where $u'_\nu \neq 0$ for all $\nu \in \{1, ..., d^{(X_0)}\}$, and obtain $\mathbf{\Sigma}' = \mathbf{U}\mathbf{\Sigma}'^{(\mathrm{all})}\mathbf{U}^T$. The rank of the square submatrix consisting of all $u'_\nu$ on the diagonal is $d^{(X_0)}$. Because the rank of a matrix is at least as large as the rank of any submatrix, $\mathbf{U}$ has rank $d^{(X_0)}$ as well. Let $\mathbf{Q}^{(\mathrm{all})}\mathbf{\Lambda}^{(\mathrm{all})}\mathbf{Q}^{(\mathrm{all})^T}$ denote the eigendecomposition of the covariance matrix $\mathbf{\Sigma}'^{(\mathrm{all})}$. Then,

$$\mathbf{\Sigma}' = \left(\mathbf{U}\mathbf{Q}^{(\mathrm{all})}\mathbf{\Lambda}^{(\mathrm{all})^{1/2}}\right)\left(\mathbf{U}\mathbf{Q}^{(\mathrm{all})}\mathbf{\Lambda}^{(\mathrm{all})^{1/2}}\right)^T, \tag{528}$$

$\mathbf{Q}^{(\mathrm{all})}\mathbf{\Lambda}^{(\mathrm{all})^{1/2}}$ has, by assumption, full rank and, thus, $\left(\mathbf{U}\mathbf{Q}^{(\mathrm{all})}\mathbf{\Lambda}^{(\mathrm{all})^{1/2}}\right)$ has the same rank as $\mathbf{U}$.[91] For any real matrix $\mathbf{A}$, $\mathrm{rank}(\mathbf{A}) = \mathrm{rank}(\mathbf{A}\mathbf{A}^T)$ (Mirsky, 2012) and we conclude $\mathbf{\Sigma}'$ has full rank $d^{(X_0)}$. □

[89]For $\nu = \omega$, Equ. 519 is identical to Equ. 105.

[90]According to Lemma 19 and the continuous mapping theorem.

[91]According to Sylvester's rank inequality (Mirsky, 2012).

**Theorem 21.** *Let* $\hat{\mathbf{b}} \sim \mathcal{N}(\mathbf{b}, \mathbf{\Sigma})$ *for full-rank covariance matrix* $\mathbf{\Sigma}$ *and mean* $\mathbf{b} \in \mathbb{R}^{d^{(X_0)}}$ *such that* $b_{k_1} = b_{k_2}$ *for all* $k_1, k_2 \in X_c^{(\mathcal{M})}$ *and all* $c \in \{1, ..., d^{(X)}\}$*. Furthermore, let all* $\mathbf{R}^{(c)}$ *for* $c \in \{1, ..., d^{(X)}\}$ *be invertible.*

*Then,* $\mathcal{T}_{\mathcal{M}}(\acute{\mathbf{b}}, \lambda^{(max)}\acute{\mathbf{\Sigma}})$ *is more conservative than the gamma distribution* $\Gamma(\zeta_{\mathcal{M}}, 1)$ *with shape parameter* $\zeta_{\mathcal{M}} = \frac{1}{2}\left(d^{(X_0)} - d^{(X)}\right)$ *and scale parameter 1. If* $\lambda^{(max)} = 1$*,* $\mathcal{T}_{\mathcal{M}}(\acute{\mathbf{b}}, \acute{\mathbf{\Sigma}}) \sim \Gamma(\zeta_{\mathcal{M}}, 1)$*.*

*Proof.* For each concise index set $X_c^{(\mathcal{M})}$, $c \in \{1, ..., d^{(X)}\}$, let $\hat{\mathbf{b}}[X_c^{(\mathcal{M})}] \in \mathbb{R}^{|X_c^{(\mathcal{M})}|}$ denote the sub-vector $\hat{b}_k$, $k \in X_c^{(\mathcal{M})}$ and let $\mathbf{b}[X_c^{(\mathcal{M})}] \in \mathbb{R}^{|X_c^{(\mathcal{M})}|}$ denote the sub-vector $b_k$, $k \in X_c^{(\mathcal{M})}$. All components of $\mathbf{b}[X_c^{(\mathcal{M})}]$ have the same value, denoted by $b^{(c)}$, because $b_{k_1} = b_{k_2}$ for all $k_1, k_2 \in X_c^{(\mathcal{M})}$ and all $c \in \{1, ..., d^{(X)}\}$. For each pair $c_1, c_2 \in \{1, ..., d^{(X)}\}$, let $\mathbf{\Sigma}[X_{c_1}^{(\mathcal{M})}, X_{c_2}^{(\mathcal{M})}] \in \mathbb{R}^{|X_{c_1}^{(\mathcal{M})}| \times |X_{c_2}^{(\mathcal{M})}|}$ denote the sub-matrix $\Sigma_{k_1 k_2}$, $k_1 \in X_{c_1}^{(\mathcal{M})}$, $k_2 \in X_{c_2}^{(\mathcal{M})}$ and let

$$\mathbf{\Sigma}[X_c^{(\mathcal{M})}, X_c^{(\mathcal{M})}] = \mathbf{Q}^{(c)}\mathbf{\Lambda}^{(c)}\mathbf{Q}^{(c)^T} \tag{529}$$

denote its eigendecomposition, where $\mathbf{Q}^{(c)}$ is an orthogonal matrix whose columns are the eigenvectors and $\mathbf{\Lambda}^{(c)}$ is a diagonal matrix whose entries are the eigenvalues of $\mathbf{\Sigma}[X_c^{(\mathcal{M})}, X_c^{(\mathcal{M})}]$. A positive semidefinite matrix is invertible if and only if it is positive definite.[92] Because $\mathbf{\Sigma}$ has full-rank, it is positive definite. Then, all sub-matrices $\mathbf{\Sigma}[X_c^{(\mathcal{M})}, X_c^{(\mathcal{M})}]$, $c \in \{1, ..., d^{(X)}\}$ are positive definite and have only non-zero eigenvalues. Therefore, $\mathbf{\Lambda}^{(c)}$ and $\acute{\mathbf{\Sigma}}[X_c^{(\mathcal{M})}, X_c^{(\mathcal{M})}]$ have full-rank and are invertible (see Equ. 157).

For each $c \in \{1, ..., d^{(X)}\}$, we define $\boldsymbol{\rho}^{(c)} = \hat{\mathbf{b}}[X_c^{(\mathcal{M})}] - \mathbf{b}[X_c^{(\mathcal{M})}]$. For $\mathbf{\Lambda}^{(c)}$ and $\mathbf{Q}^{(c)}$ obtained by the eigendecomposition,

$$\mathbf{\Lambda}^{(c_1)^{-\frac{1}{2}}}\mathbf{Q}^{(c_1)^T}\boldsymbol{\rho}^{(c)} \tag{530}$$

is joint standard normal and[93]

$$\begin{aligned}
\acute{\mathbf{b}}[X_c^{(\mathcal{M})}] &= \mathbf{R}^{(c)^{-1}}\mathbf{Q}^{(c)^T}\hat{\mathbf{b}}[X_c^{(\mathcal{M})}] && (531)\\
&= \mathbf{R}^{(c)^{-1}}\mathbf{Q}^{(c)^T}\mathbf{b}[X_c^{(\mathcal{M})}] + \mathbf{R}^{(c)^{-1}}\mathbf{Q}^{(c)^T}\boldsymbol{\rho}^{(c)} && (532)\\
&= \mathbf{R}^{(c)^{-1}}\mathbf{r}^{(c)}b^{(c)} + \left(\mathbf{R}^{(c)^{-1}}\mathbf{\Lambda}^{(c)^{\frac{1}{2}}}\right)\left(\mathbf{\Lambda}^{(c)^{-\frac{1}{2}}}\mathbf{Q}^{(c)^T}\boldsymbol{\rho}^{(c)}\right) && (533)\\
&\sim \mathcal{N}(\mathbf{b}^{(c)}, \mathbf{R}^{(c)^{-2}}\mathbf{\Lambda}^{(c)}) && (534)\\
&\sim \mathcal{N}(\mathbf{b}^{(c)}, \acute{\mathbf{\Sigma}}[X_c^{(\mathcal{M})}, X_c^{(\mathcal{M})}]). && (535)
\end{aligned}$$

Although all components of $\acute{\mathbf{b}}[X_c^{(\mathcal{M})}]$ are mutually independent, they are still potentially correlated with components of $\acute{\mathbf{b}}[X_{c_1}^{(\mathcal{M})}]$ for $c_1 \neq c$. Moreover, decorrelation is not possible without violating the independence assumption of modules, i.e., observables inside and outside of modules have to be independent when conditioned on the interface variable.[94] Let $\mathring{\mathbf{\Sigma}}$ denote the full potentially non-diagonal covariance matrix of $\acute{\mathbf{b}}$, so that the diagonal elements of $\mathring{\mathbf{\Sigma}}$ and (the diagonal) $\acute{\mathbf{\Sigma}}$ are identical. To compensate for the additional off-diagonal terms of $\mathring{\mathbf{\Sigma}}$, we introduce $\lambda$ large enough such that the test statistic $\mathcal{T}_{\mathcal{M}}(\acute{\mathbf{b}}, \lambda\acute{\mathbf{\Sigma}})$ is more conservative than $\Gamma(\zeta_{\mathcal{M}}, 1)$.

More precisely,[95] only the length of projections of $\acute{\boldsymbol{\rho}} = \acute{\mathbf{b}} - \mathbf{b}$ into certain subspaces of $\acute{\boldsymbol{\rho}}$ contribute to $\mathcal{T}_{\mathcal{M}}$. We rescale $\mathring{\boldsymbol{\rho}} = \lambda^{-1/2}\acute{\boldsymbol{\rho}}$ by a common factor $\lambda \geq 1$ such

[92](Horn and Johnson, 2012, p. 431, Corollary 7.1.7)

[93]See Equ. 156 to 159.

[94]In case of a global eigendecomposition not restricted to each functional index set $c$, $\acute{\mathbf{b}}^{(c)}$ in Equ. 531 would, in general, depend on contributions from other functional modules, i.e., $\hat{\mathbf{b}}^{(c_1)}$ for $c_1 \neq c$.

[95]According to Theorem 12 and Equ. 348.

that for any fixed projection, denoted by $\mathbf{A}$, the distribution of the lengths after the projection, i.e., $|\mathbf{A}\mathring{\boldsymbol{\rho}}|$ for the dependent $\mathring{\boldsymbol{\rho}} \sim \mathcal{N}(\mathbf{0}, \lambda^{-1}\mathring{\boldsymbol{\Sigma}})$, is more conservative than the corresponding distribution of $|\mathbf{A}\breve{\boldsymbol{\rho}}|$ for an independent $\breve{\boldsymbol{\rho}} \sim \mathcal{N}(\mathbf{0}, \acute{\boldsymbol{\Sigma}})$. Then, also the test statistic $\mathcal{T}_{\mathcal{M}}(\mathring{\boldsymbol{\rho}}, \mathring{\boldsymbol{\Sigma}})$ is more conservative than $\mathcal{T}_{\mathcal{M}}(\breve{\boldsymbol{\rho}}, \mathring{\boldsymbol{\Sigma}})$ if both statistics depend on the same covariance $\mathring{\boldsymbol{\Sigma}}$ and, thus, implement the same projection $\mathbf{A}$. Because

$$\mathcal{T}_{\mathcal{M}}(\mathring{\boldsymbol{\rho}}, \mathring{\boldsymbol{\Sigma}}) = \mathcal{T}_{\mathcal{M}}(\lambda^{-1/2}\acute{\boldsymbol{\rho}}, \mathring{\boldsymbol{\Sigma}}) = \mathcal{T}_{\mathcal{M}}(\acute{\boldsymbol{\rho}}, \lambda\mathring{\boldsymbol{\Sigma}}) = \mathcal{T}_{\mathcal{M}}\left(\acute{\mathbf{b}}, \lambda\acute{\boldsymbol{\Sigma}}\right) \tag{536}$$

and

$$\mathcal{T}_{\mathcal{M}}(\breve{\boldsymbol{\rho}}, \mathring{\boldsymbol{\Sigma}}) = \mathcal{T}_{\mathcal{M}}(\breve{\boldsymbol{\rho}}, \acute{\boldsymbol{\Sigma}}) \sim \Gamma(\zeta_{\mathcal{M}}, 1), \tag{537}$$

it follows $\mathcal{T}_{\mathcal{M}}\left(\acute{\mathbf{b}}, \lambda\acute{\boldsymbol{\Sigma}}\right)$ is more conservative than $\Gamma(\zeta_{\mathcal{M}}, 1)$.

The smallest $\lambda$ with this property is the largest eigenvalue $\lambda^{(\max)}$ of the covariance matrix $\mathbf{M}$ of the observables whitened independently for each $c \in \{1, ..., d^{(\mathcal{X})}\}$ (see Equ. 160 and 161). As $\mathcal{T}_{\mathcal{M}}$ does not dependent on $X_c^{(\mathcal{M})}$ if $|X_c^{(\mathcal{M})}| = 1$, the corresponding entries in $\mathbf{M}$ are removed before the eigenvalue calculation by setting $\mathbf{M}[X_{c_1}^{(\mathcal{M})}, X_{c_2}^{(\mathcal{M})}] = 0$ if either $|X_{c_1}^{(\mathcal{M})}| = 1$ or $|X_{c_2}^{(\mathcal{M})}| = 1$. If $\lambda^{(\max)} = 1$, $\acute{\mathbf{b}} \sim \mathcal{N}(\mathbf{b}, \acute{\boldsymbol{\Sigma}})$.

We conclude that $\mathcal{T}_{\mathcal{M}}(\acute{\mathbf{b}}, \lambda^{(\max)}\acute{\boldsymbol{\Sigma}})$ is more conservative than the gamma distribution $\Gamma(\zeta_{\mathcal{M}}, 1)$ with shape parameter $\zeta_{\mathcal{M}} = \frac{1}{2}\left(d^{(\mathcal{X}_0)} - d^{(\mathcal{X})}\right)$ and scale parameter 1. If $\lambda^{(\max)} = 1$, $\mathcal{T}_{\mathcal{M}}(\acute{\mathbf{b}}, \acute{\boldsymbol{\Sigma}}) \sim \Gamma(\zeta_{\mathcal{M}}, 1)$. □

**Numerical Analysis 22.**

*Let* $\hat{\mathbf{b}}^{(v)} \sim \mathcal{N}\left(\mathbf{b}^{(v)}, \mathbf{\Sigma}^{(v)}\right)$ *for* $v \in \{1, ..., d^{(\mathcal{X}_0)}\}$ *and* $\mathbf{\Sigma}$ *be diagonal. Let* $\theta_{\hat{\mu}} \in \{5, 6, 7\}$ *and* $\hat{\sigma}_0 \to \infty$. *Furthermore, let* $p^{(\Gamma(\zeta_M,1))}$*-value denote the p-value of the gamma distribution* $\Gamma(1/2, 2)$ *for shape parameter* $\frac{1}{2}$ *and scale parameter 2 and p-value*$|_{\rho^{(v)}\in\mathcal{I}}$ *denote the p-value of the distribution of*

$$\tilde{\rho} = \frac{\left(\hat{b}_v(\mathbf{\Sigma}^{(v)}) - b_v\right)^2}{\hat{\Sigma}_{vv}(\mathbf{\Sigma}^{(v)})} \tag{538}$$

*conditioned on* $(\boldsymbol{\rho}^{(v)} \in \mathcal{I}$ *for all* $v \in \{1, ..., d^{(\mathcal{X}_0)}\})$.

*We consolidate with numerical methods that*

$$p\text{-}value|_{\rho^{(v)}\in\mathcal{I}} \leq p^{(\Gamma(1/2,2))}\text{-}value \tag{539}$$

*for all* $p^{(\Gamma(1/2,2))}$*-value* $\leq 0.95$.[96]

We first transform $\hat{\mathbf{b}}^{(v)}$ into standard form[97]

$$\frac{\hat{b}_v(\mathbf{\Sigma}^{(v)}) - b_v}{\sqrt{\hat{\Sigma}_{vv}(\mathbf{\Sigma}^{(v)})}} = \frac{\hat{\mu}_v(\mathbf{\Sigma}^{(v)}) - \mu_v(\mathbf{\Sigma}^{(v)})}{\sqrt{\lambda_v}\hat{\sigma}_v^{(\mathrm{T})}(\mathbf{\Sigma}^{(v)})} \tag{540}$$

for $v \in \{1, ..., d^{(\mathcal{X}_0)}\}$. A distribution with cumulative distribution function $F^{(\mathrm{a})}$ is more conservative than a distribution with cumulative distribution function $F^{(\mathrm{b})}$ if for the corresponding inverse cumulative distribution functions

$$F^{(\mathrm{a})^{-1}} \leq F^{(\mathrm{b})^{-1}} \tag{541}$$

for all $F^{(\mathrm{b})}$.

In order to obtain precise bounds for $F^{(\mathrm{a})^{-1}}$, we split $\mathcal{I}$ into $800 \times 800$ suitably located and disjoint 2D bins filling all $\mathcal{I}$. For each 2D bin, we calculate the probability mass for the joint standard normal $\hat{\boldsymbol{\mu}}^{(v)}$ and construct two distribution functions each consisting of $800 \times 800$ point masses. That is, for the first (second) distribution we introduce for each 2D bin a point mass containing the total probability mass of that bin located at one of its corners, i.e., the corner with the largest (smallest) value of the random variable for which $F^{(\mathrm{a})^{-1}}$ is calculated. The inverse cumulative distribution functions for both of these distributions can be calculated precisely, the first (second) resulting in a precise upper (lower) bound for $F^{(\mathrm{a})^{-1}}$.[98] By construction, the upper bound of $F^{(\mathrm{a})^{-1}}$ is above $F^{(\mathrm{b})^{-1}}$ for $F^{(\mathrm{b})}$ close to zero. Therefore, we constrain the range of investigated $F^{(\mathrm{b})}$ values to above $(1 - 0.95)$.

Let $F^{(1)^{-1}}$ denote the upper bound of the inverse cumulative distribution function of $\tilde{\rho}$ conditioned on $(\boldsymbol{\rho}^{(v)} \in \mathcal{I}$ for all $v \in \{1, ..., d^{(\mathcal{X}_0)}\})$. Furthermore, let $F^{(\Gamma(1/2,2))^{-1}}$ denote the inverse cumulative distribution function of $\Gamma(1/2, 2)$. To consolidate Equ. 539, we verify for $162 \times 121$ pairs of $(\mu_1^{(v)}, \mu_2^{(v)})$, i.e.,

$$\mu_1^{(v)} \in \{10^k | k \in \{-\infty, -1, -1 + \tfrac{1}{40}, -1 + \tfrac{2}{40}, ..., 3\}\} \tag{542}$$

$$\mu_2^{(v)} \in \{10^k | k \in \{0, \tfrac{1}{40}, \tfrac{2}{40}, ..., 3\}\}, \tag{543}$$

[96] The chi-squared distribution $\chi^2(1)$ is a special case of the gamma distribution $\Gamma(1/2, 2)$.

[97] According to Lemma 16.

[98] Except for the bin that contains the single maximum of the investigated random variable, which in general lies not at one of its corners. As this error is local, and small compared to differences between compared inverse cumulative distribution functions (or their bounds) it can be neglected.

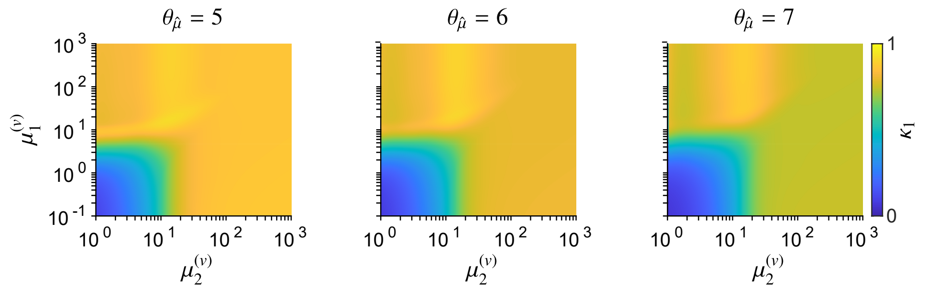


Figure S9: $\kappa_1$ for $\theta_{\hat{\mu}} = 5$ (left panel), $\theta_{\hat{\mu}} = 6$ (middle panel) and $\theta_{\hat{\mu}} = 7$ (right panel).

that the upper bound $F^{(1)^{-1}}$ is smaller than $F^{(\Gamma(1/2,2))^{-1}}$ for all $F^{(\Gamma(1/2,2))} \geq (1 - 0.95)$. Fig. S9 shows

$$\kappa_1(\mu_1^{(v)}, \mu_2^{(v)}) = \max \frac{F^{(1)^{-1}}}{F^{(\Gamma(1/2,2))^{-1}}}, \tag{544}$$

as a function of $\mu_1^{(v)}$ and $\mu_2^{(v)}$, where the maximum is taken over all $\tilde{\rho}$ such that $F^{(\Gamma(1/2,2))} \geq (1 - 0.95)$. For all tested $\mu_1^{(v)}$ and $\mu_2^{(v)}$, $\kappa_1$ is smaller than one.[99]

[99]The author gratefully acknowledges the computational and data resources provided by the Leibniz Supercomputing Centre (www.lrz.de).

**Numerical Analysis 23.**

*Let* $d^{(X_0)} = 2$ *and*

$$\begin{pmatrix}\hat{\mathbf{b}}^{(1)}\\ \hat{\mathbf{b}}^{(2)}\end{pmatrix} \sim \mathcal{N}\left(\begin{pmatrix}\mathbf{b}^{(1)}\\ \mathbf{b}^{(2)}\end{pmatrix}, \mathbf{\Sigma}^{(all)}\right) \tag{545}$$

$$\mathbf{\Sigma}^{(all)} = \begin{pmatrix}\mathbf{\Sigma}^{(1,1)} & \mathbf{\Sigma}^{(1,2)}\\ \mathbf{\Sigma}^{(2,1)} & \mathbf{\Sigma}^{(2,2)}\end{pmatrix}. \tag{546}$$

*Let* $\theta_{\hat{\mu}} \in \{5, 6, 7\}$ *and* $\hat{\sigma}_0 \to \infty$. *Furthermore, let* $p^{(\Gamma(1,1))}$*-value denote the p-value of the gamma distribution* $\Gamma(1/2, 1)$ *for shape parameter* $\frac{1}{2}$ *and scale parameter 1 and p-value*$|_{\boldsymbol{\rho}^{(v)}\in\mathcal{I}}$ *denote the p-value of the distribution of* $\mathcal{T}_{\mathcal{M}}$ *conditioned on* ($\boldsymbol{\rho}^{(v)} \in \mathcal{I}$ *for all* $v \in \{1, 2\}$).

*We consolidate with numerical methods that if* $b_1 = b_2$, $\Theta_4(\mathbf{\Sigma}^{(1)}) = \Theta_4(\mathbf{\Sigma}^{(2)})$ *and* $\Theta_3(\mathbf{\Sigma}^{(1)}) = \Theta_3(\mathbf{\Sigma}^{(2)})$*, then*

$$p\text{-}value|_{\boldsymbol{\rho}^{(v)}\in\mathcal{I}} \leq p^{(\Gamma(1/2,1))}\text{-}value \tag{547}$$

*for* $\mathcal{T}_{\mathcal{M}}(\hat{\mathbf{b}}, \lambda^{(max)}\hat{\mathbf{\Sigma}})$, $\mathcal{T}_{\mathcal{M}}(\acute{\mathbf{b}}, \lambda^{(max)}\acute{\mathbf{\Sigma}})$ *and* $10^{-5} < p^{(\Gamma(1/2,1))}$*-value* $< 0.05$. *If in addition* $\hat{b}_1$ *and* $\hat{b}_2$ *are non-negatively correlated, then the inequality holds also for* $\mathcal{T}_{\mathcal{M}}(\hat{\mathbf{b}}, \hat{\mathbf{\Sigma}})$ *and* $10^{-5} < p^{(\Gamma(1/2,1))}$*-value* $< 0.05$.

We first transform $\hat{\mathbf{b}}^{(v)}$ for $v \in \{1, 2\}$ into standard form[100]

$$\frac{\hat{b}_v(\mathbf{\Sigma}^{(v,v)}) - b_v}{\sqrt{\hat{\Sigma}_{vv}(\mathbf{\Sigma}^{(v,v)})}} = \frac{\hat{\mu}_v(\mathbf{\Sigma}^{(v,v)}) - \mu_v(\mathbf{\Sigma}^{(v,v)})}{\sqrt{\lambda_v}\hat{\sigma}_v^{(\mathrm{T})}(\mathbf{\Sigma}^{(v,v)})}. \tag{548}$$

In general, $\mathbf{\Sigma}^{(1,2)}$ has non-zero elements and

$$\begin{pmatrix}\hat{\boldsymbol{\mu}}^{(1)}\\ \hat{\boldsymbol{\mu}}^{(2)}\end{pmatrix} \sim \mathcal{N}\left(\begin{pmatrix}\mu_1^{(1)}\\ \mu_2^{(1)}\\ \mu_1^{(2)}\\ \mu_2^{(2)}\end{pmatrix}, \begin{pmatrix}1 & 0 & c_{11} & c_{12}\\ 0 & 1 & c_{21} & c_{22}\\ c_{11} & c_{21} & 1 & 0\\ c_{12} & c_{12} & 0 & 1\end{pmatrix}\right), \tag{549}$$

for $c_{ij} \in [-1, 1]$ and $i, j \in \{1, 2\}$. By assumption $b_1 = b_2$, $\Theta_4(\mathbf{\Sigma}^{(1)}) = \Theta_4(\mathbf{\Sigma}^{(2)})$ and $\Theta_3(\mathbf{\Sigma}^{(1)}) = \Theta_3(\mathbf{\Sigma}^{(2)})$, so that[101]

$$\mu_1^{(1)} = \mu_2^{(1)}\frac{\mu_1^{(2)}}{\mu_2^{(2)}}. \tag{550}$$

The constraints $\Theta_4(\mathbf{\Sigma}^{(1)}) = \Theta_4(\mathbf{\Sigma}^{(2)})$ and $\Theta_3(\mathbf{\Sigma}^{(1)}) = \Theta_3(\mathbf{\Sigma}^{(2)})$ are chosen for computational feasibility.[102]

A distribution with cumulative distribution function $F^{(\mathrm{a})}$ is more conservative than a distribution with cumulative distribution function $F^{(\mathrm{b})}$ if for the corresponding inverse cumulative distribution functions

$$F^{(\mathrm{a})^{-1}} \leq F^{(\mathrm{b})^{-1}} \tag{551}$$

[100] According to Lemma 16.

[101] See Equ. 440.

[102] The author gratefully acknowledges the computational and data resources provided by the Leibniz Supercomputing Centre (www.lrz.de).

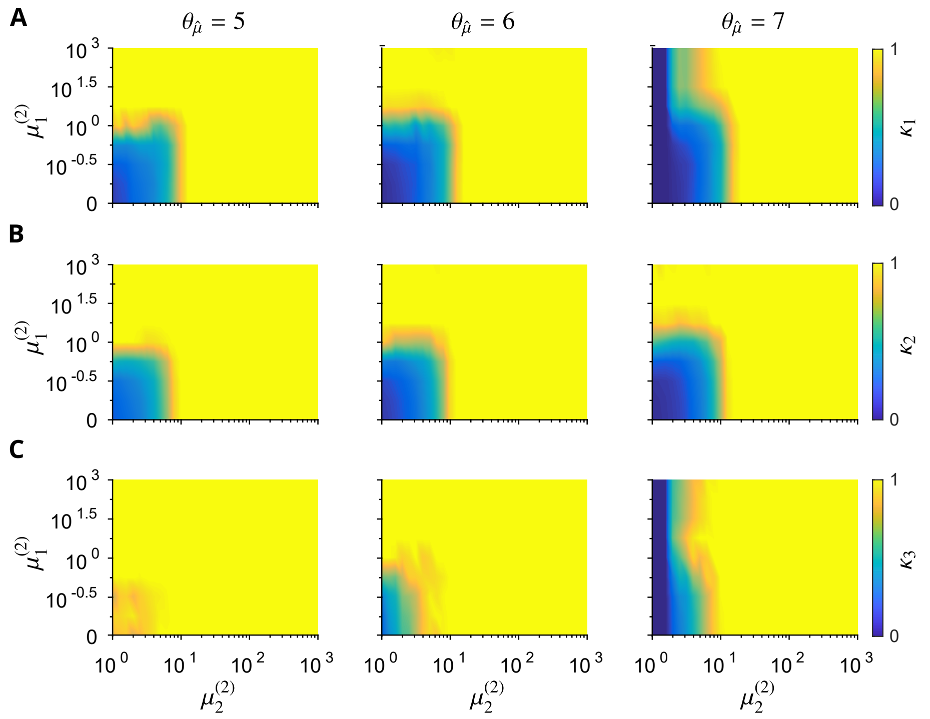


Figure S10: $\kappa_1$ (**A**), $\kappa_2$ (**B**) and $\kappa_3$ (**C**) for $\theta_{\hat{\mu}} = 5$ (left panels), $\theta_{\hat{\mu}} = 6$ (middle panels) and $\theta_{\hat{\mu}} = 7$ (right panels).

for all $F^{(b)}$.

In order to obtain precise bounds for $F^{(a)^{-1}}$, we split $\mathcal{I} \times \mathcal{I}$ into $n \times n \times n \times n$ suitably located and disjoint 4D bins filling all $\mathcal{I} \times \mathcal{I}$. For each 4D bin, we calculate the probability mass for the joint standard normal $(\hat{\boldsymbol{\mu}}^{(1)}, \hat{\boldsymbol{\mu}}^{(2)})$ and construct two distribution functions each consisting of $n \times n \times n \times n$ point masses. That is, for the first (second) distribution we introduce for each 4D bin a point mass containing the total probability mass of that bin located at one of its corners, i.e., the corner with the largest (smallest) value of the random variable for which $F^{(a)^{-1}}$ is calculated. The inverse cumulative distribution functions for both of these distributions can be calculated precisely, the first (second) resulting in a precise upper (lower) bound for $F^{(a)^{-1}}$.[103]

By construction, the upper bound of $F^{(a)^{-1}}$ is above $F^{(b)^{-1}}$ for $F^{(b)}$ close to zero. Therefore, we constrain the range of investigated $F^{(b)}$ to above $(1-0.05)$. For computational feasibility, we start with $n = 8$ and increase $n$ by two if $F^{(a)^{-1}} > F^{(b)^{-1}}$ for some $(1-10^{-5}) \geq F^{(b)} \geq (1-0.05)$.[104] The largest value for $n$ obtain by this procedure is 40. Let $F^{(1)^{-1}}$, $F^{(2)^{-1}}$ and $F^{(3)^{-1}}$ denote the upper bounds of the inverse cumulative distribution functions of $\mathcal{T}_{\mathcal{M}}(\hat{\mathbf{b}}, \hat{\boldsymbol{\Sigma}})$, $\mathcal{T}_{\mathcal{M}}(\hat{\mathbf{b}}, \lambda^{(\max)}\hat{\boldsymbol{\Sigma}})$ and $\mathcal{T}_{\mathcal{M}}(\acute{\mathbf{b}}\lambda^{(\max)}\acute{\boldsymbol{\Sigma}})$, respectively. Furthermore, let $F^{(\Gamma(1/2,1))^{-1}}$ denote the inverse cumulative distribution function of $\Gamma(1/2, 1)$.

[103]Except for the bin that contains the single maximum of the investigated random variable, which in general lies not at one of its corners. As this error is local, and small compared to differences between compared inverse cumulative distribution functions (or their bounds) it can be neglected.

[104]The precise value of very small $p$-values is little informative, because, in general, sample distributions deviate at the far tails from their assumed asymptotic form.

We choose $c_{ij} \in \{-0.99, -0.66, -0.33, 0, 0.33, 0.66, 0.99\}$ for $i, j \in \{1, 2\}$ and generate joint normal moments for all combinations of $c_{ij}$ that result in positive semi-definite covariance matrices.[105] For $\mathcal{T}_{\mathcal{M}}(\hat{\mathbf{b}}, \hat{\boldsymbol{\Sigma}})$, we consider only combinations of $c_{ij}$ that result in non-negatively correlated $\hat{b}_1$ and $\hat{b}_2$. We verify for $9 \times 33 \times 33$ values of $(\mu_1^{(1)}, \mu_2^{(1)}, \mu_1^{(2)}, \mu_2^{(2)})$, i.e.,

$$\mu_1^{(2)} \in \{10^k | k \in \{-\infty, -1, -0.5, 0, ..., 2, 3\}\} \tag{552}$$

$$\mu_2^{(1)}, \mu_2^{(2)} \in \{10^k | k \in \{0, \tfrac{1}{160}, \tfrac{1}{40}, 0.1, 0.2, 0.3, ..., 3\}\}, \tag{553}$$

that the upper bounds $F^{(1)^{-1}}$, $F^{(2)^{-1}}$ and $F^{(3)^{-1}}$ are smaller than $F^{(\Gamma(1/2,1))^{-1}}$ for all $(1 - 10^{-5}) \geq F^{(\Gamma(1/2,1))} \geq (1 - 0.05)$. Fig. S10 shows

$$\kappa_n(\mu_1^{(2)}, \mu_2^{(2)}) = \max \frac{F^{(n)^{-1}}}{F^{(\Gamma(1/2,1))^{-1}}}, \tag{554}$$

for $n \in \{1, 2, 3\}$ as a function of $\mu_1^{(2)}$ and $\mu_2^{(2)}$, where the maximum is taken over all tested $c_{ij}$, $\mu_2^{(1)}$ and $\mathcal{T}_{\mathcal{M}}$ such that $(1 - 10^{-5}) \geq F^{(\Gamma(1/2,1))} \geq (1 - 0.05)$. For all parameter values, $\kappa_n < 1$ for $n \in \{1, 2, 3\}$.[106]

[105] By applying the Cholesky decomposition, resulting in 569 combinations.

[106] The author gratefully acknowledges the computational and data resources provided by the Leibniz Supercomputing Centre (www.lrz.de).

### References


Anderson, T. (2003). Wiley series in probability and statistics. 3. hoboken.

Billingsley, P. (2013). *Convergence of probability measures*. John Wiley & Sons.

Driver, B. K. (2003). *Analysis tools with applications*. Springer.

Horn, R. A. and Johnson, C. R. (2012). *Matrix analysis*. Cambridge university press.

Kenney, J. and Keeping, E. (1951). The distribution of the standard deviation. *Mathematics of Statistics, Pt*, 2:170–173.

Kleiber, C. and Stoyanov, J. (2013). Multivariate distributions and the moment problem. *Journal of Multivariate Analysis*, 113:7–18.

Marsaglia, G. et al. (2006). Ratios of normal variables. *Journal of Statistical Software*, 16(4):1–10.

Mathai, A. M. and Provost, S. B. (1992). *Quadratic forms in random variables: theory and applications*. M. Dekker New York.

Mirsky, L. (2012). *An introduction to linear algebra*. Courier Corporation.

Stuart, A. and Ord, K. (1994). Kendalls advanced theory of statistics arnold.

Van der Vaart, A. W. (2000). *Asymptotic statistics*, volume 3. Cambridge university press.
